# Transcriptomic response to different heme sources in *Trypanosoma cruzi*

**DOI:** 10.1101/2025.07.14.664770

**Authors:** Evelyn Tevere, María G. Mediavilla, Cecilia B. Di Capua, Marcelo L. Merli, Carlos Robello, Luisa Berná, Julia A. Cricco

**Author notes:** Corresponding authors: Julia A. Cricco, Luisa Berná. Equal contribution as first authors (Evelyn Tevere, María G. Mediavilla).

## Abstract

Heme is an essential molecule for most organisms, yet some parasites, like *Trypanosoma cruzi*, the causative agent of Chagas disease, cannot synthesize it. These parasites must acquire heme from their hosts, making this process critical for their survival. In the midgut of the insect vector, *T. cruzi* epimastigotes are exposed to both hemoglobin (Hb) and free heme resulting from its degradation. Despite the importance of this nutrient, how different heme sources influence parasite gene expression remains poorly understood.

Here, we showed that heme restitution either as hemin or Hb to heme-starved parasites induces an early and distinct transcriptional response in *T. cruzi* epimastigotes. Using RNA sequencing at 4- and 24-hours post-supplementation, we identified gene subsets commonly or uniquely regulated by each heme source, including genes putatively linked to heme acquisition and metabolism. We also presented here the first studies focused on CRAL/TRIO domain-containing protein (*Tc*CRAL/TRIO), a novel heme responsive hemoprotein identified from this study.

Our results provide a more detailed picture of *T. cruzi* biology and highlights heme acquisition as a promising point of vulnerability. These findings may ultimately contribute to the identification of potential molecular targets for the development of new therapeutic strategies against Chagas disease.

## INTRODUCTION

*Trypanosoma cruzi* is the causative agent of Chagas disease, a human neglected disease. This parasite presents a digenetic life cycle comprising two hosts (a mammalian host and a hematophagous triatomine vector) and, at least, four well defined developmental stages.

Trypanosomatids are aerobic organisms that rely on hemoproteins for key metabolic processes, including ergosterol biosynthesis, fatty acid desaturation, and mitochondrial respiration [1]. However, *T. cruzi*, as well as other trypanosomatids, cannot synthesize heme and must acquire it from its hosts [2, 3]. This parasite can incorporate the cofactor during its replicative life-cycle stages, epimastigotes and amastigotes [4]. In the triatomine midgut, hemoglobin (Hb, a heme-bound tetrameric protein), serves as a significant heme source, while free heme (product of protein degradation)is directly bioavailable for epimastigotes [5]. However, free heme also poses a significant threat due to its capacity to catalyze the formation of reactive oxygen species (ROS), which can damage cellular components [6]. To counteract the toxic effects of ROS and enhance its survival, *T. cruzi* has evolved several defense mechanisms, including the trypanothione system—a functional analog of the glutathione system in most eukaryotes—along with robust DNA repair pathways and unique DNA polymerases [7]. Previously, we studied the participation of *Tc*HRG (former *Tc*HTE) in epimastigote heme uptake both as free heme (added as hemin) [4, 8] and as Hb [9]. Our findings revealed that the parasite modulates heme uptake in response to intracellular heme levels to balance the supply of this essential cofactor for hemoproteins while mitigating its potential toxicity. Specifically, heme starvation triggers an increase in *Tc*HRG at mRNA and protein levels which are subsequently downregulated within the first 24 hours of hemin or Hb restoration [8, 9]. However, the precise mechanisms underlying the regulation of *Tc*HRG gene expression, the intracellular trafficking of heme, and its utilization as a cofactor or as an iron source remain to be elucidated.

In this study, we used transcriptome analysis as a tool to investigate the molecular response of the parasite to heme stress, analyzing the replenishment of the heme source, provided as either hemin or Hb, in cultures previously starved of this cofactor. The analysis of the differentially expressed genes (DEGs) over time indicates that *T. cruzi* exhibits both shared and distinct transcriptional responses to different sources of heme. A core set of genes commonly modulated by hemin and Hb suggests a conserved heme-responsive program, involving redox balance and metabolic adaptations. Hemin triggers extensive remodeling of gene expression related to structural components, glycolysis, and signaling pathways. In contrast, Hb supplementation elicits a more limited but distinct transcriptional shift, marked by the induction of proteolytic enzymes and modulation of signaling components.

Additionally, the analysis of the heme-supplemented samples (hemin or Hb) vs. heme-deprived samples (no heme addition after starvation) revealed a small but consistent set of DEGs, most of them downregulated. Notably, a CRAL/TRIO domain-containing protein (named here *Tc*CRAL/TRIO), consistently repressed by heme, emerged as a key candidate for further study. Biochemical assays confirmed *Tc*CRAL/TRIO as a hemoprotein, suggesting a key role in heme homeostasis in *T. cruzi*.

In summary, this study contributes to a better understanding of the biology of *T. cruzi* regarding heme homeostasis in the epimastigote stage and gives significant information to renew the dissection of heme uptake, metabolism and distribution.

## RESULTS

### 1 – *Tc*HRG expression presents a fine-tuned response to heme

To gain insight and identify genes and pathways involved in heme homeostasis, we investigated *T. cruzi* transcriptional response to heme by RNA-sequencing (RNA-seq) using free heme (added as hemin) and Hb as heme sources. We chose *TcHRG* as a model gene [8, 9] and performed a heme-response time course experiment in epimastigotes. Briefly, epimastigotes routinely cultured in the presence of 5 µM hemin were subjected to heme starvation (LIT+10% FBS without the addition of a heme source) for 48 h. Then, parasites were transferred to fresh media supplemented with 5 µM hemin, 1.25 µM Hb (equivalent to 5 µM heme as Hb) or without heme (heme-deprived samples, identified as no heme in figures). The starvation step ensured uniform starting conditions for assessing the effect of heme restitution for both sources. Samples were collected immediately before repletion (t_0_) and at different time intervals over 24 (for mRNA) and 48 (for protein) hours (Figure 1A).

**Figure 1.**
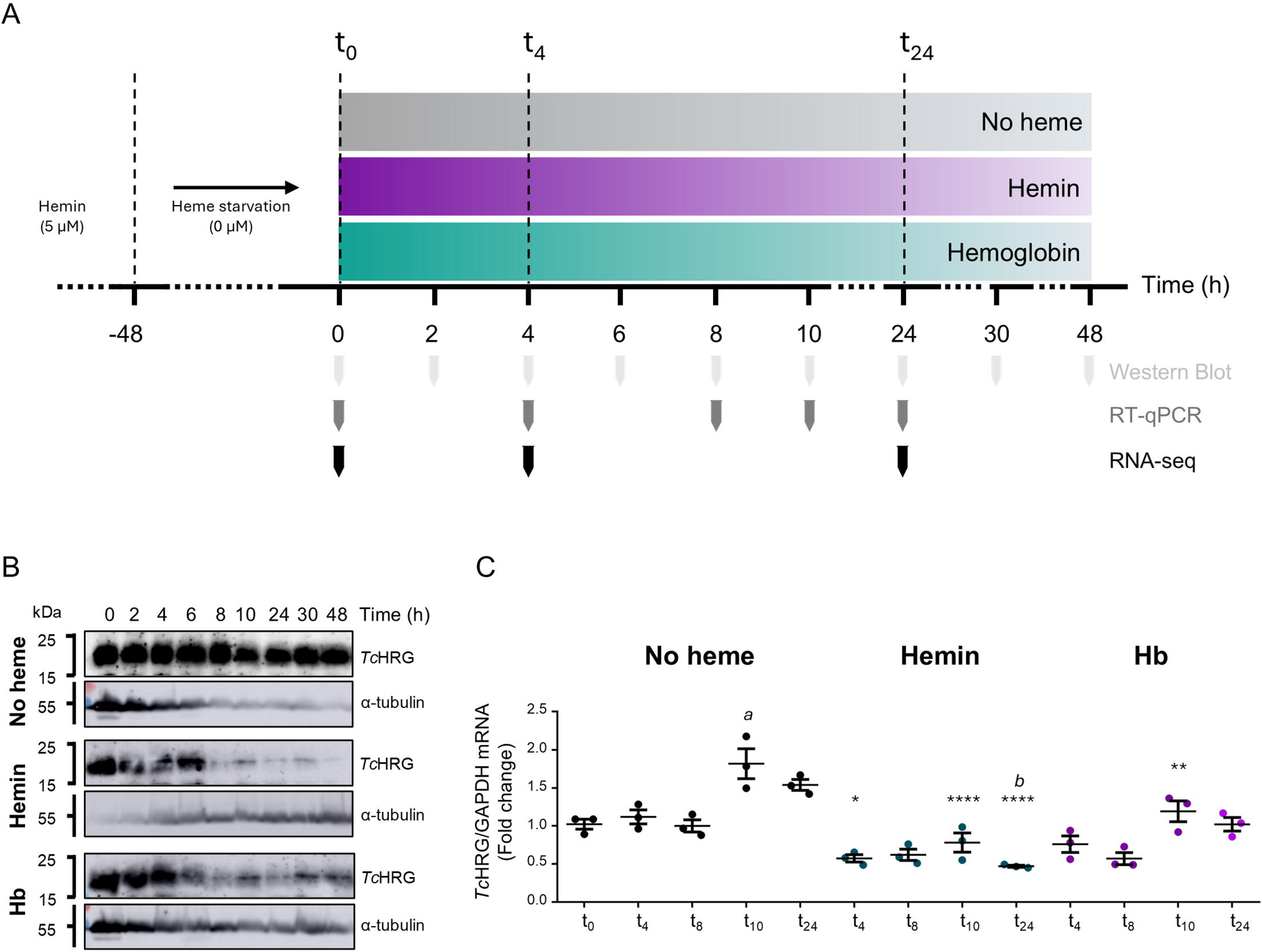
**(A)** Experimental design scheme. Epimastigotes cultured in LIT+10% FBS + 5 µM hemin were subjected to heme starvation for 48 h. Parasites were then transferred to fresh media supplemented with 5 µM hemin, 1.25 µM hemoglobin (Hb), or without heme. The time points for sample collection used in *Tc*HRG analysis by western blot and RT-qPCR as well as for the samples included in the RNA-seq analysis are indicated. **(B)** Western blot analysis of *Tc*HRG protein levels and α-tubulin in the three conditions analyzed (no heme, 5 µM hemin, and 1.25 µM Hb) at different time points. **(C)** RT-qPCR analysis of *Tc*HRG mRNA levels in the three conditions: no heme (black dots), 5 µM hemin (blue dots), and 1.25 µM Hb (purple dots), at different time points post-treatment, using GAPDH as the reference gene. Data are presented as the mean ± SD of three biological replicas. Statistical significance was determined by one-way ANOVA followed by Tukey’s multiple comparisons test (****: p < 0.0001; ***: p < 0.001; **: p < 0.01; *: p < 0.05, significantly different from no heme condition at the same time of incubation. *a*: p < 0.001; *b*: p < 0.05, significantly different against no heme t_0_ sample).

Western blotting using anti-*Tc*HRG polyclonal antibodies [8] showed that *Tc*HRG protein signals remained stable in heme-deprived parasites but decreased markedly 6– 8 hours after supplementation with either hemin or Hb, being almost undetectable in hemin-treated and notably reduced in Hb-supplemented samples (Figure 1B).

*TcHRG* transcript levels were analyzed by RT-qPCR (Figure 1C). In heme-deprived epimastigotes, they remained stable up to 8 hours, followed by an increase at 10 and 24 hours after medium renewal. Upon heme reintroduction, transcript levels dropped approximately 50% and 25% after 4 hours of treatment with hemin and Hb, respectively. These changes in mRNA levels mirrored the protein expression patterns, with a more marked response to hemin than to Hb, consistent with our previous observations [9].

Based on these observations, we conducted the RNA-seq assay using the experimental scheme shown in Figure 1A. Samples were collected at three time points: after heme starvation (t_0_), 4 hours (t_4_), and 24 hours post heme refeeding (t_24_). Heme-deprived cultures were included to account for experimental noise arising only from medium replacement and the dilution steps used to keep the parasites in logarithmic growth.

### 2 – Heme restitution induces rapid and mild changes in epimastigotes’ transcriptome

A global comparison between heme-supplemented (hemin or Hb) and heme-deprived conditions, irrespective of time, identified 51 DEGs (Table S1). In this initial analysis, *TcHRG* showed a log_2_ (fold change) (logFC) of −0.48, consistent with our previous results [8, 9]. Based on this and considering the moderate variations detected in the overall transcriptional response, we established a logFC threshold of |0.40| as a bona fide cutoff for subsequent transcriptomic analyses.

Applying this threshold and a statistical significance criterion of adjusted p-value (padj) < 0.05, we identified 20 DEGs (Figure 2). Although the overall number of DEGs is relatively small, the affected genes suggest potential transcriptional adjustments related to protein processing, metabolism, and electron transport. The specific contributions of these pathways will be explored in detail in the following sections.

**Figure 2.**
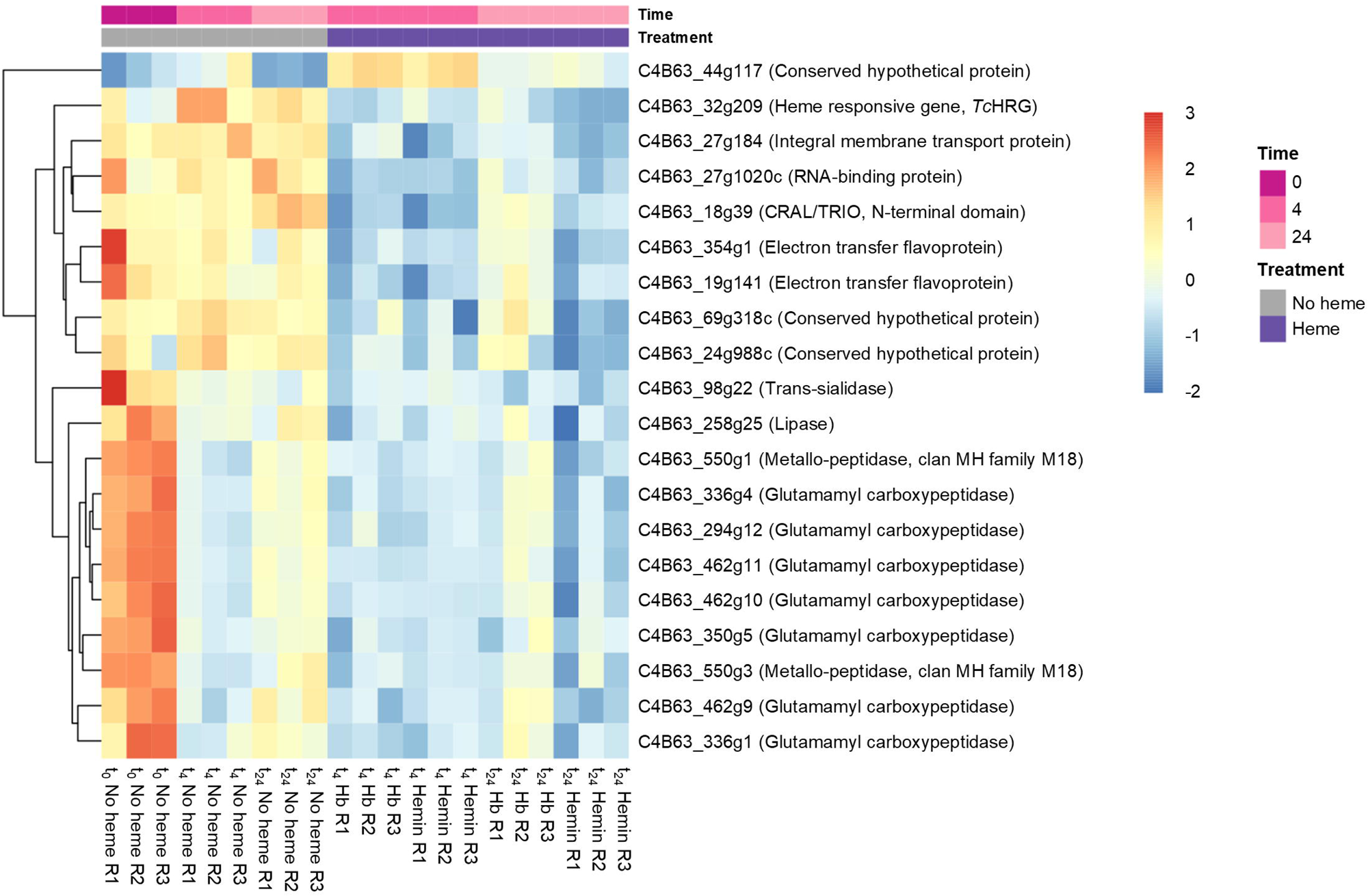
Heatmap illustrating the expression patterns of differentially expressed genes in heme-supplemented (hemin or hemoglobin, Hb) and heme-deprived (no heme) conditions, irrespective of time. Gene IDs and annotations listed on the right correspond to the *Trypanosoma cruzi* Dm28c 2018 genome sequence and annotation (*TritrypDB*). Sample nomenclature is as follows (example): “t_0_ No heme R1” refers to a No heme sample from experiment replica 1 at time 0 h.

To further dissect the transcriptional response to heme over time, we analyzed DEGs at t_4_ and t_24_ against baseline levels (t_0_) in heme-deprived, hemin, and Hb conditions (Figure 3A and Table S2).

**Figure 3.**
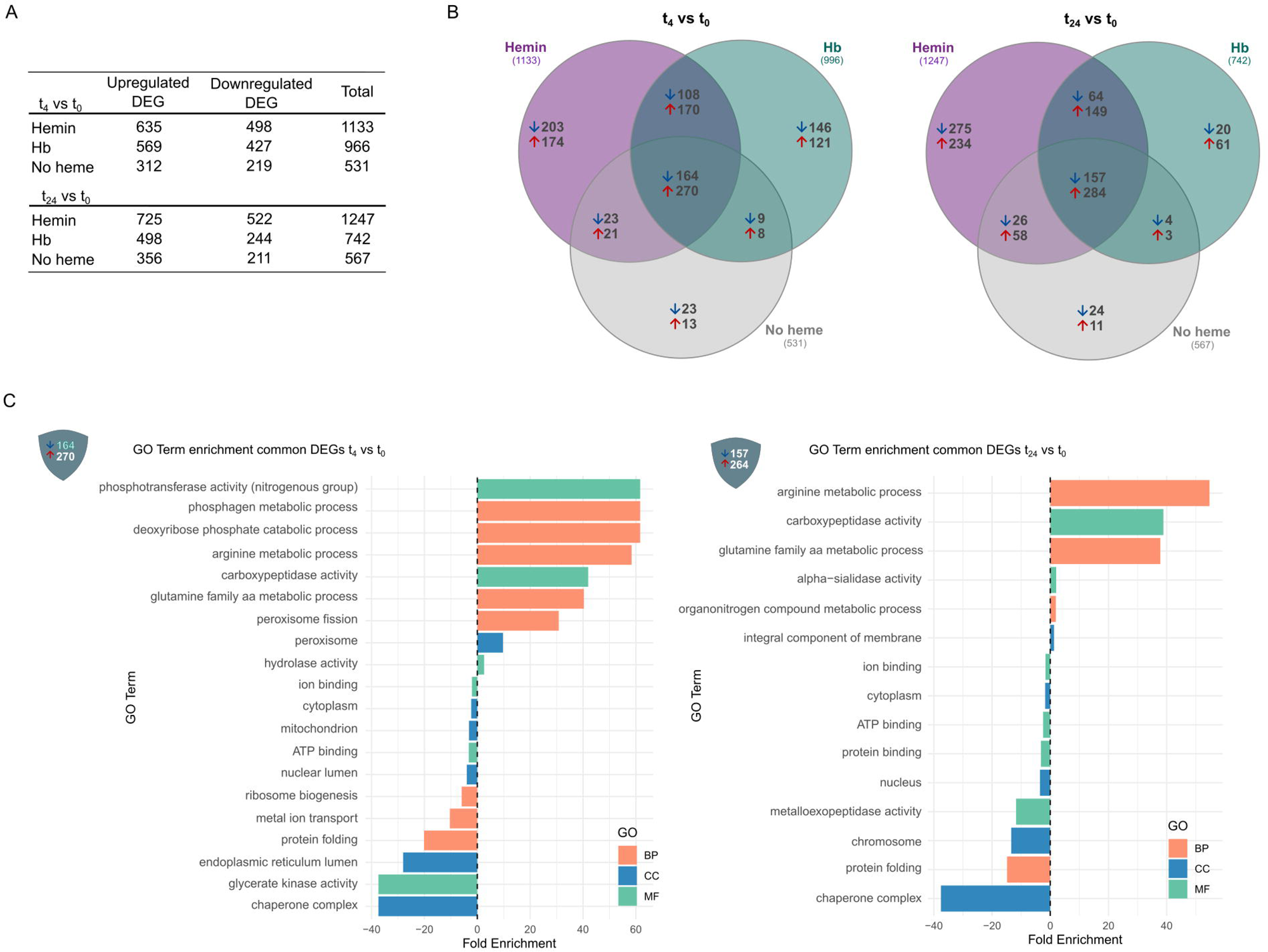
**(A)** Differentially expressed genes (DEGs) at t_4_ and t_24_ after media renewal compared to baseline levels (t_0_) in each of the three culture conditions: no heme, hemin, and hemoglobin (Hb). **(B)** Venn diagrams showing DEGs among the no heme, hemin, and Hb conditions at t_4_ vs. t_0_ and t_24_ vs. t_0_. **(C)** Gene Ontology (GO) enrichment analysis of genes shared among the three conditions at t_4_ vs. t_0_ and t_24_ vs. t_0_. BP: Biological Process. CC: Cellular Component. MF: Molecular Function.

As mentioned earlier, very few DEGs exhibited a logFC greater than |1| (doubling or halving their expressions) (Table S2) and more than 75% showed moderate expression changes, with logFC values between |0.4| and |0.7| (Figure S1). Still, transcriptional changes were already detectable at t_4_, indicating an early adjustment to heme restoration.

A subset of DEGs shared across all conditions was identified, comprising 434 genes (t_4_ vs. t_0_) and 441 genes (t_24_ vs. t_0_) (Figure 3B, Table S2). These genes were excluded from further analysis, as they likely represent a general response to media renewal and parasite dilution rather than a specific effect of variations in heme availability (Figure 3C).

### 2a – Genes involved in the core transcriptional response to heme

We first identified a set of genes specifically modulated by both hemin and Hb (hemin∩Hb), which points to a shared transcriptional response to heme. This group included 108 downregulated and 170 upregulated genes in the t_4_ vs. t_0_ dataset, and 64 downregulated and 149 upregulated genes in the t_24_ vs. t_0_ dataset (Table 1, Table S3). Gene Ontology (GO) analysis (Figure 4A, Table S3) of downregulated genes at t_4_ vs. t_0_ revealed enrichment in molecular function terms such as alanine metabolic process and electron transfer activities. For example, we observed the downregulation of genes encoding subunits VI and VIII of cytochrome c oxidase and two electron transfer flavoproteins. In the t_24_ vs. t_0_ comparison, terms were associated with phosphotransferase activity, transporter activity, and membrane components.

**Figure 4.**
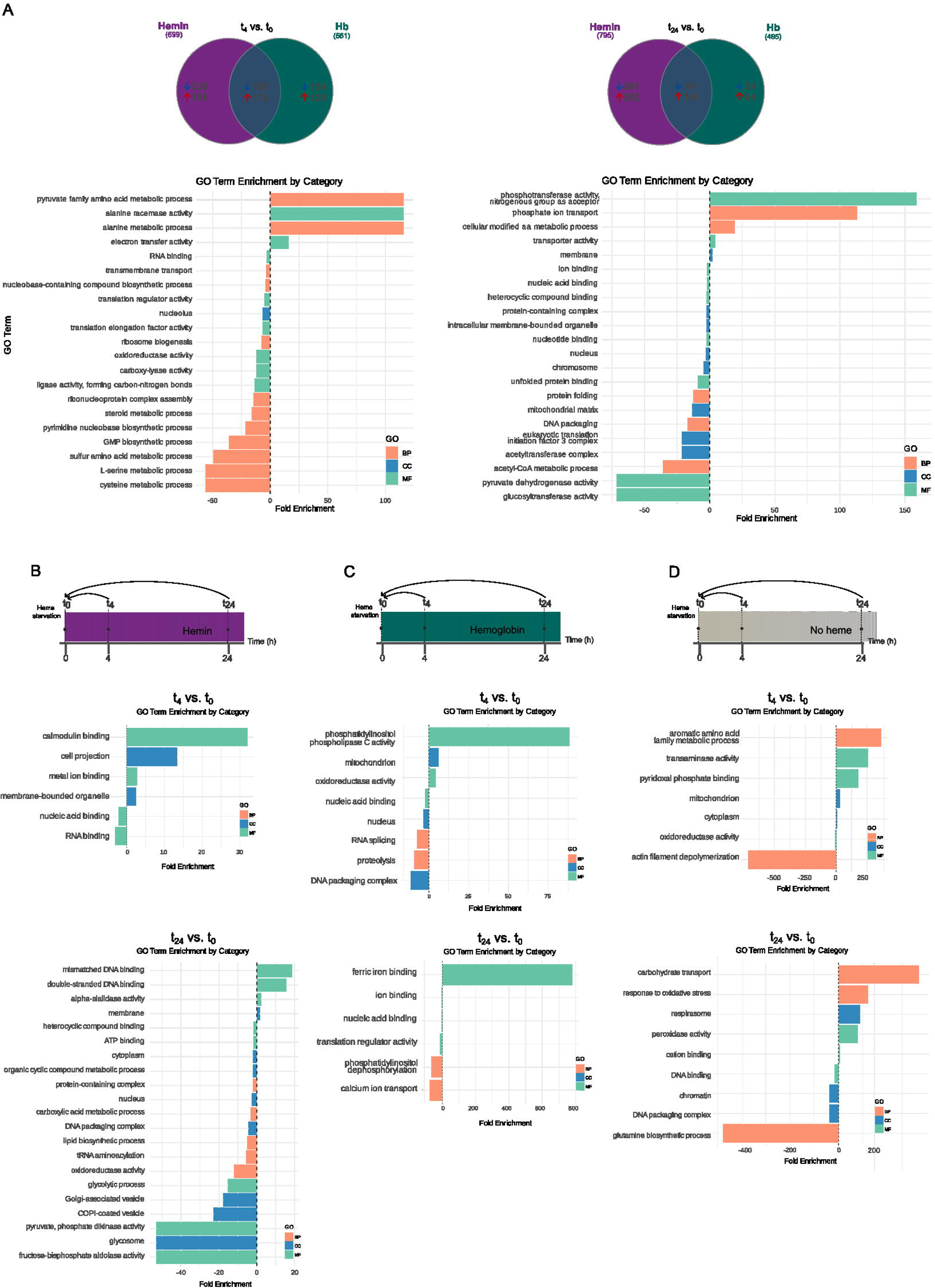
Differentially expressed genes (DEGs) in no heme, hemin, hemoglobin (Hb) conditions, and merged heme (hemin and Hb) conditions. **(A)** Common genes detected simultaneously in hemin- and Hb-supplemented conditions, along with Gene Ontology (GO) analysis of the response to heme. **(B)** GO analysis of DEGs in the hemin condition at t_4_ vs. t_0_ and t_24_ vs. t_0_. **(C)** GO analysis of DEGs in the Hb condition at t_4_ vs. t_0_ and t_24_ vs. t_0_. **(D)** GO analysis of DEGs in the no heme condition at t_4_ vs. t_0_ and t_24_ vs. t_0_. BP: Biological Process. CC: Cellular Component. MF: Molecular Function.

**Table 1.** Differentially expressed genes uniquely modulated under the no heme, hemin, or hemoglobin (Hb) conditions. These genes were identified in the t_4_ vs. t_0_ and t_24_ vs. t_0_ comparisons, excluding those whose expression changes were attributed to media renewal (i.e., genes common to all conditions).

|  | Upregulated<br>DEGs | Downregulated<br>DEGs | Total |
| --- | --- | --- | --- |
| $t_4$ vs. $t_0$ | | | |
| No heme | 13 | 23 | 36 |
| Hemin | 174 | 203 | 377 |
| Hb | 121 | 146 | 267 |
| $t_{24}$ vs. $t_0$ | | | |
| No heme | 11 | 24 | 35 |
| Hemin | 234 | 275 | 509 |
| Hb | 61 | 20 | 81 |

It is also worth mentioning the downregulation of genes involved in arginine metabolism and fatty acid metabolism. We identified two and six copies of the arginine permease gene (t_4_ and t_24_, respectively) and three copies of the arginine kinase gene (t_24_). Also, analysis at t_4_ revealed a downregulation of genes encoding for cytochrome b5-dependent oleate desaturase, cytochrome b5–which acts as an electron donor for several desaturases–and cytochrome b5 domain-containing protein 1.

Regarding upregulated genes, the t_4_ vs. t_0_ dataset showed enrichment in GO terms including serine and cysteine metabolism, transmembrane transport, ribosome biogenesis, pyrimidine biosynthesis, steroid metabolism, oxidoreductase activity, and nucleolus components, among others. In the t_24_ vs. t_0_ dataset, enriched processes included terms such as protein folding, acetyl-CoA metabolism, and DNA packaging. Many of the identified genes were associated with cellular components from the mitochondrial matrix and nucleus, including nucleosome assembly proteins, eukaryotic translation initiation factor 3, ruvB-like DNA helicase, and histone genes. Additional upregulated genes were involved in binding to small molecules and unfolded protein binding, as well as displaying hexosyl transferase and pyruvate dehydrogenase activities. Terms involving translation regulation were significantly enriched at both time points. Notably, a gene encoding the glycolytic enzyme enolase was upregulated at t_24_. A thorough analysis of the upregulated hemin∩Hb DEGs revealed several genes involved in trypanothione biosynthesis and ROS detoxification. These include a diphthine synthase gene (t_4_), 19 cystathionine beta synthase (CBS) genes (t_4_), a trypanothione reductase gene (t_4_), a spermidine synthase gene (t_4_ and t_24_), a pyridoxal kinase gene (t_24_), and a putrescine-cadaverine transporter gene (t_24_). Diphthine synthase produces S-adenosylhomocysteine, a homocysteine precursor, while CBS catalyzes its condensation with serine to form cystathionine, an intermediate in glutathione biosynthesis. CBS requires pyridoxal phosphate as a cofactor, which is obtained through the phosphorylation of pyridoxal by pyridoxal kinase. Both glutathione and spermidine are essential to trypanothione biosynthesis, and finally, trypanothione reductase helps to regenerate the reduced form of this compound. Putrescine, on the other hand, serves as a substrate for spermidine synthase in the production of spermidine. Also, we found upregulated two peroxidoxin genes (t_24_) which share 53% amino acid sequence identity with tryparedoxin peroxidase and are likely involved in detoxifying ROS as well. Additionally, two genes encoding for enzymes that produce reducing equivalents as NADPH also appeared upregulated: glucose-6-phosphate dehydrogenase (G6PD) gene (t_4_ and t_24_) and malic enzyme gene (t_4_).

### 2b – Hemin induces changes in the expression of specific genes

To assess the specific transcriptional response to hemin, we analyzed DEGs uniquely modulated under this condition, excluding those shared with the Hb treatment or the heme-deprived samples. In this group, we identified 203 downregulated and 174 upregulated genes in the t_4_ vs. t_0_ dataset, while in the t_24_ vs. t_0_ comparison, 275 genes were downregulated and 234 were upregulated (Table 1, Table S4). In this analysis, *TcHRG* displayed a logFC of −0.45, consistent with RT-qPCR results (Figure 1C).

GO enrichment analysis (Figure 4B and Table S4) of downregulated genes highlighted a reduction in the expression of genes associated with the flagellum (t_4_) and membrane (t_24_). For example, 18 and 31 trans-sialidase (TS) genes showed decreased expression at t_4_ and t_24_, respectively. Additionally, 19 mucin-associated surface proteins (MASP) genes were downregulated at t_24_. The GO terms calmodulin binding and metal ion binding (t_4_) and mismatched DNA binding (t_24_) revealed downregulation of 11 genes encoding paraflagellar rod proteins (PRP), and 18 flagellar calcium-binding proteins (FCaBP), among others [10].

Conversely, upregulated genes were enriched for the nucleic acid binding function at t_4_, while at t_24_, genes related to lipid biosynthesis, amino acid activation, carboxylic acids, organic cyclic compound metabolism, and glycolysis were significantly overrepresented. Within the glycolysis term, we found genes encoding two fructose-bisphosphate aldolases, two 3-phosphate dehydrogenases, and one phosphoglycerate kinase.

We also found a subset of DEGs encoding proteins that may participate in signaling pathways in response to hemin (highlighted in yellow in Table S4).

Finally, apart from the genes identified in the hemin∩Hb dataset, two additional genes encoding proteins related to mitochondrial electron transport chain were downregulated in this condition: NADH-ubiquinone oxidoreductase mitochondrial (t_4_) and cytochrome c oxidase assembly protein (t_24_).

### 2c -– Hemoglobin modulates the expression of a subset of genes

To determine the specific transcriptional response to Hb, we excluded those DEGs shared with the hemin treatment or the heme-deprived samples.

The analysis of DEGs in Hb-supplemented parasites revealed 146 downregulated and 121 upregulated genes in the t_4_ vs. t_0_ dataset, whereas in the t_24_ vs. t_0_ comparison, 20 genes were downregulated and 61 were upregulated (Table 1, Table S4).

GO enrichment analysis (Figure 4C and Table S4) of downregulated genes in the t_4_ vs. t_0_ dataset identified mitochondrion components and oxidoreductase and phospholipase C activities as the most affected categories. Additionally, in the t_24_ vs. t_0_ comparison, the biological process iron ion binding, which includes a putative frataxin gene, was significantly downregulated. Among upregulated genes, the biological process proteolysis was enriched at t_4_ vs. t_0_, including 33 cysteine peptidases (cruzipain). Furthermore, genes associated with nuclear components and RNA splicing were also upregulated at this time point. At t_24_ vs. t_0_, molecular functions such as phosphatidylinositol dephosphorylation, translation regulation, calcium ion transport, and ion binding were significantly enriched. Also, the nucleic acid binding category was upregulated at both time points.

A more detailed analysis of these DEGs revealed a group of genes involved in cellular signaling (highlighted in yellow in Table S4) that might play a role in the specific cellular response to Hb.

Finally, aside from the genes found in the hemin∩Hb dataset, at t_4_ we identified additional downregulated genes related to fatty acid metabolism and the mitochondrial electron transport chain in Hb supplemented parasites. These include genes encoding fatty acyl CoA synthetase and delta-4 fatty acid desaturase, as well as cytochrome c, ubiquinone biosynthesis protein COQ7 homolog, and various subunits of Complex I (subunits NB6M, NI8M, and two copies of NADH-ubiquinone oxidoreductase complex I subunit).

### 2d – Heme deprivation results in differential gene expression

To identify transcriptional changes associated with heme starvation, we focused on genes differentially expressed exclusively in heme-deprived samples, excluding those shared with either hemin and Hb conditions (Figures 3B, 4B and 4C). The analysis of this dataset revealed a small number of starvation-specific DEGs, including 23 downregulated and 13 upregulated genes in the t_4_ vs. t_0_ dataset and 24 downregulated and 11 upregulated genes in the t_24_ vs. t_0_ analysis (Table 1, Table S4).

GO enrichment analysis (Figure 4D, Table S4) of downregulated genes at t_4_ revealed a significant decrease in transaminase activity, comprising 15 tyrosine aminotransferase (TAT) genes [11]. Also, genes encoding proteins associated with mitochondrion and cytoplasm were downregulated at this time point. By t_24_, the transcriptional response shifted, with downregulated genes including those encoding proteins of the respiratory chain complex, factors involved in oxidative stress response and carbohydrate transport, as well as proteins with cation-binding properties.

Conversely, upregulated genes showed enrichment for terms associated with actin depolymerization and oxidoreductase activity at t_4_. At t_24_, the upregulated genes were related to nucleic acid binding, glutamine biosynthesis, and nucleosome components.

### 2e - Heme induces persistent transcriptional changes in *T. cruzi*

From the DEGs identified at t_4_ across the three culture conditions, approximately 80% remained differentially expressed at t_24_. Among these, some genes maintained their expression levels, while others showed a further increase/decrease at t_24_ compared to t_4_ (Figure 5A and B, Table S5).

**Figure 5.**
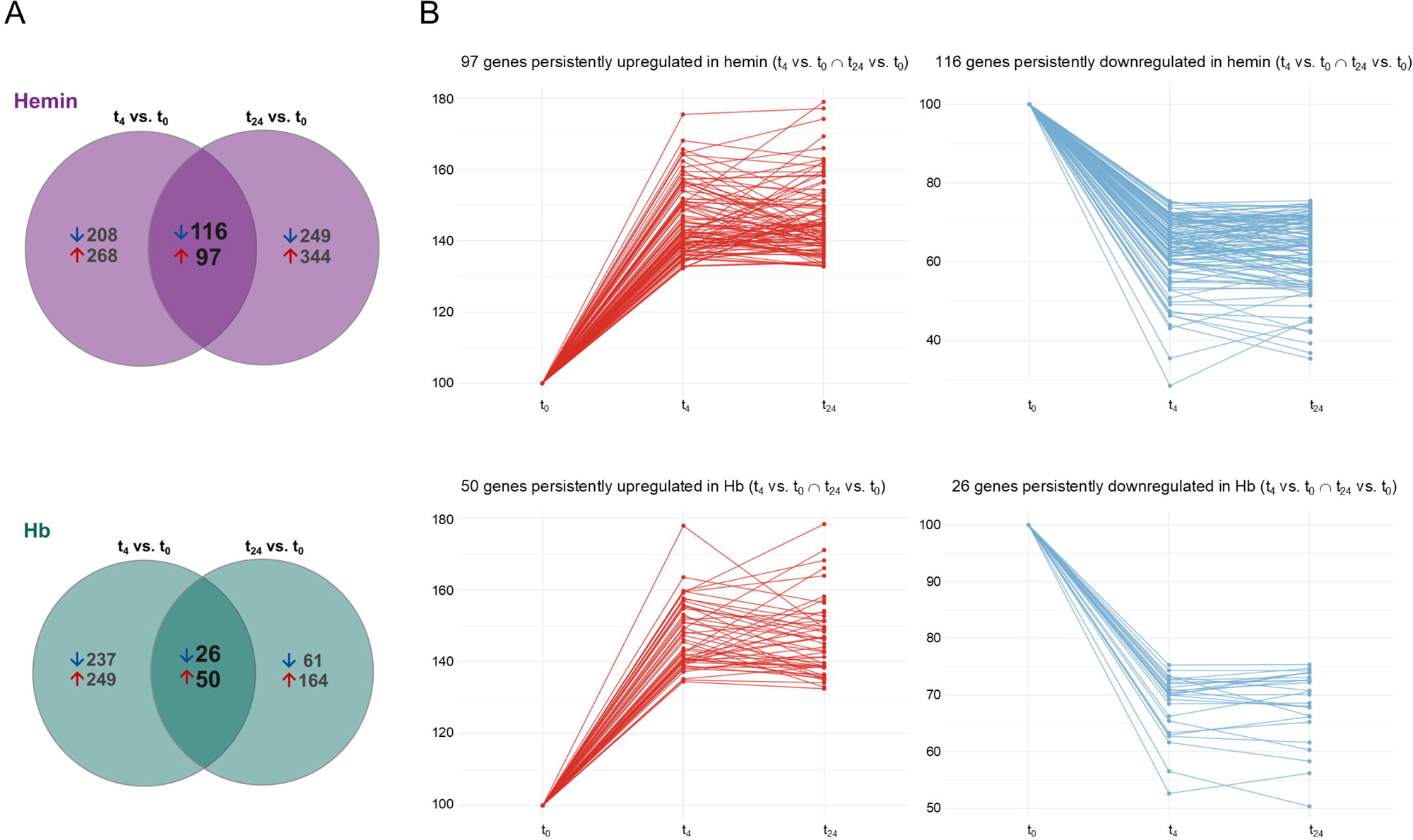
Persistent differential expression of genes in hemin and hemoglobin (Hb) conditions from t_4_ to t_24_ (A) Venn diagrams showing shared differentially expressed genes between the t_4_ vs. t_0_ and t_24_ vs. t_0_ comparisons in hemin and Hb-supplemented parasites. (B) Expression profiles over time of upregulated (red) and downregulated (blue) genes that exhibit sustained differential expression in hemin- and Hb-supplemented parasites.

Specifically, in the hemin treatment, 97 upregulated and 116 downregulated genes exhibited a sustained transcriptional response throughout the experiment. Similarly, in the Hb treatment, 50 genes remained upregulated and 26 downregulated over time (Table S5). In the heme-deprived samples, only two genes (C4B63_51g82 and C4B63_51g81, both encoding ferric reductases) were persistently found upregulated over time.

### 3 - Heme triggers specific expression changes in a small set of genes

To further explore the specific effects of heme supplementation, we performed a third analysis comparing gene expression in hemin- or Hb-supplemented cultures against the heme-deprived samples at each corresponding time point (t_4_ and t_24_) (Figure 6A). This analysis revealed a small but consistent set of DEGs.

**Figure 6.**
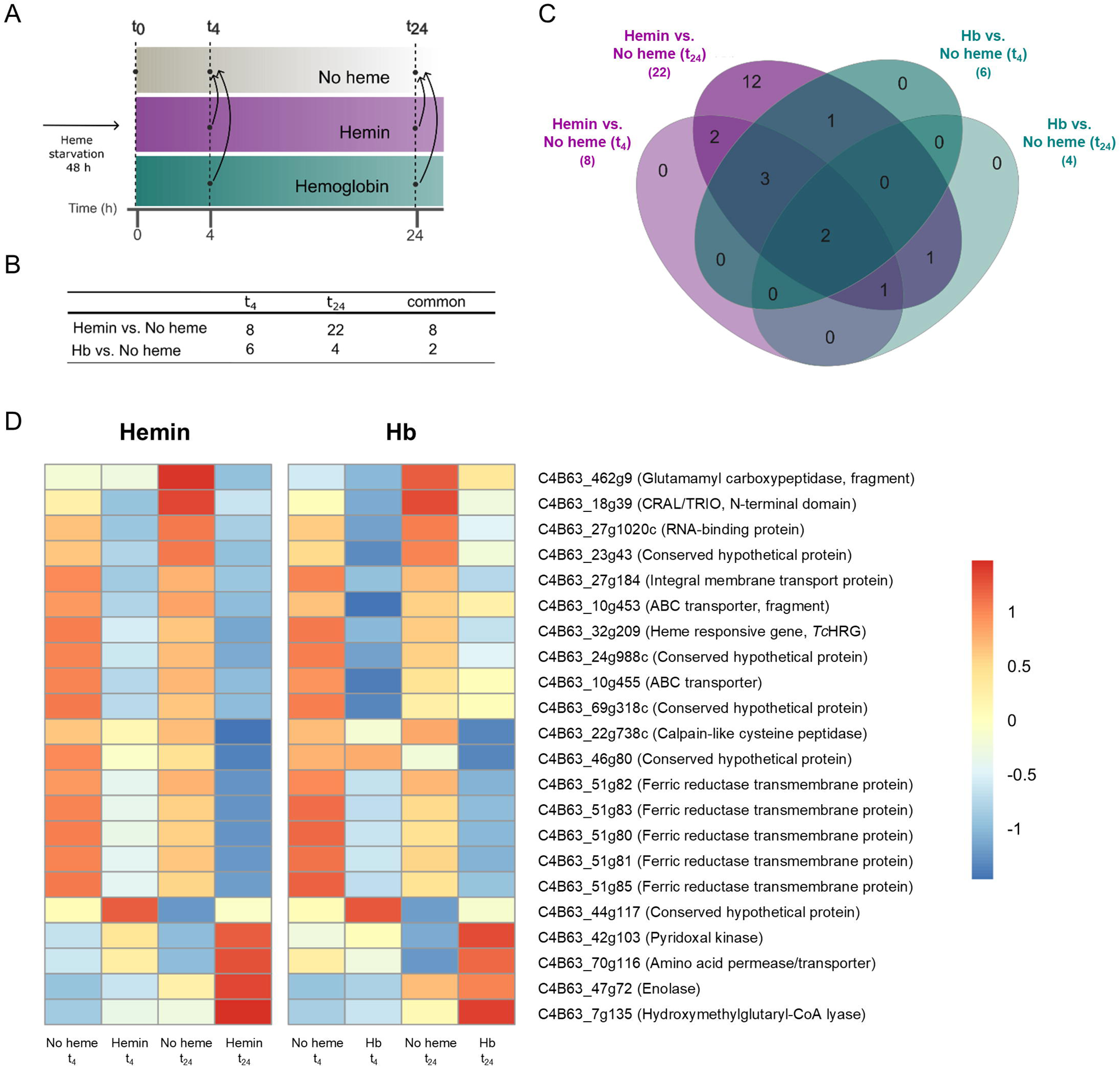
**(A)** Schematic representation of the experimental design, highlighting the comparisons between hemin- and hemoglobin (Hb)-supplemented conditions versus the no heme condition at t_4_ and t_24_. **(B)** Table summarizing the number of differentially expressed genes identified (DEGs) in each comparison. **(C)** Venn diagram showing shared DEGs in each condition. **(D)** Heatmap showing the expression profiles of 22 genes differentially expressed in both hemin vs. no heme and Hb vs. no heme conditions at t_4_ and t_24_. Gene IDs and functional annotations, shown on the right, correspond to the *Trypanosoma cruzi* Dm28c 2018 genome (*TriTrypDB*).

In hemin-supplemented parasites, only 8 downregulated genes were detected at t_4_. At t_24_, 22 genes were differentially expressed, including 17 downregulated (of which the 8 identified at t_4_ were maintained) and 5 upregulated genes (Figure 6B and Table S6).

For Hb-supplemented epimastigotes, a total of 6 downregulated genes were detected at t_4_, while at t_24_, three downregulated and one upregulated gene were identified. Only two downregulated genes were shared between t_4_ and t_24_ (Figure 6C, Table S6) and also repressed by hemin. In fact, all DEGs found in Hb-supplemented samples were also included within the 22 DEGs identified in hemin-supplemented cells at t_24_ (Figure 6B). In this analysis, *TcHRG* presented a logFC higher than |0.40| for hemin (t_24_) and Hb (t_4_).

The two downregulated genes shared between both heme sources at both time points correspond to a CRAL/TRIO domain-containing protein (C4B63_18g39) and a ferric reductase (C4B63_51g81), one of the five *TcFR* genes identified in this analysis. Additionally, an RNA-binding protein (C4B63_27g1020c) was consistently downregulated (logFC ∼ −1) across all conditions except Hb at t_24_, while an integral membrane transporter (C4B63_27g184) was downregulated exclusively in hemin-supplemented samples. Moreover, a conserved hypothetical protein (C4B63_44g117) was found upregulated at t_24_ in both hemin- and Hb-supplemented parasites (Figure 6D).

Also, the previously mentioned five genes encoding ferric reductases (*TcFR*) involved in iron metabolism [12] and containing four hallmark histidine residues predicted to bind heme molecules [13] were downregulated. The products of these genes share more than 95% amino acid sequence identity between each other.

Another ten downregulated genes with unclear roles in heme metabolism were detected. C4B63_27g1020c encodes an RNA-binding protein and C4B63_18g39 encodes a protein that contains a CRAL/TRIO domain (Interpro: IPR001251), which is commonly found in members of the SEC14 family [14]. Additionally, four of these genes may be involved in metabolite transport: two ABC transporters (C4B63_10g454 and C4B63_10g455), an integral membrane transport protein belonging to the Major Facilitator Superfamily (C4B63_27g184), and a conserved hypothetical DEG (C4B63_23g43) containing an EamA domain (InterPro: IPR000620), which is often associated to drug/metabolite transporters. Two peptidases were also downregulated, a calpain-like cysteine peptidase (C4B63_69g318c), and a glutamamyl carboxypeptidase (C4B63_462g9). On the other hand, the peptides encoded by C4B63_24g988c and C4B63_69g318c—which share 95% amino acid sequence identity—are classified as dimerization-anchoring domains from a cAMP-dependent protein kinase (Supfam: SSF47391). Finally, the analysis also identified a conserved hypothetical gene (C4B63_46g80); however, no relevant information is available for it.

The upregulated DEGs identified in this analysis include an amino acid permease transporter (C4B63_70g116) from the Amino acid-Polyamine-Organocation family, a pyridoxal kinase (C4B63_42g103), the glycolytic enzyme enolase (C4B63_47g72), a mitochondrial hydroxymethylglutaryl-CoA lyase (C4B63_7g135), associated with ketogenesis and leucine catabolism, and a conserved hypothetical gene (C4B63_44g117) containing a putative SPRY domain (InterPro: IPR003877), known to participate in key signaling pathways, such as RNA processing and regulation of histone methylation.

All DEGs identified here were also detected in the second analysis, except for hydroxymethylglutaryl-CoA lyase and the amino acid permease transporter genes. Additionally, genes encoding CRAL/TRIO domain-containing protein, *Tc*HRG, RNA-binding protein, glutamamyl carboxypeptidase, SPRY domain-containing protein, and the hypothetical dimerization-anchoring domains of cAMP-dependent protein kinases were also found in our first analysis, further highlighting the consistency and robustness of our results.

### 4 - *TcCRAL/TRIO* is a heme responsive gene encoding a hemoprotein

While SEC14 family proteins are generally known for binding small lipids such as phosphatidylinositol, Khan et al. recently characterized a Sec14-like phosphatidylinositol transfer protein from *Saccharomyces cerevisiae*, named Sec-Fourteen Homolog 5 (*Sc*Sfh5, UniProt ID: A6ZQI5) that, unexpectedly, binds heme instead [15]. It has been proposed that *Sc*Sfh5 functions in redox control and/or the regulation of heme homeostasis under stress conditions induced by organic oxidants. Considering this, we performed a series of assays to characterize the related protein encoded by the C4B63_18g39 (named here: *T. cruzi* CRAL/TRIO domain-containing protein, *Tc*CRAL/TRIO). *TcCRAL/TRIO* encodes a 183 amino acids protein and presented a logFC greater than −1.7 in hemin and −1.4 in Hb conditions compared to no heme at t_4_.

To characterize *Tc*CRAL/TRIO, we analyzed its gene expression by RT-qPCR. As shown in Figure 7A, *TcCRAL/TRIO* presented a lower expression for hemin and Hb replenished cultures compared to heme-deprived ones both at t_4_ and t_24_, thus validating RNA-seq assay results.

**Figure 7.**
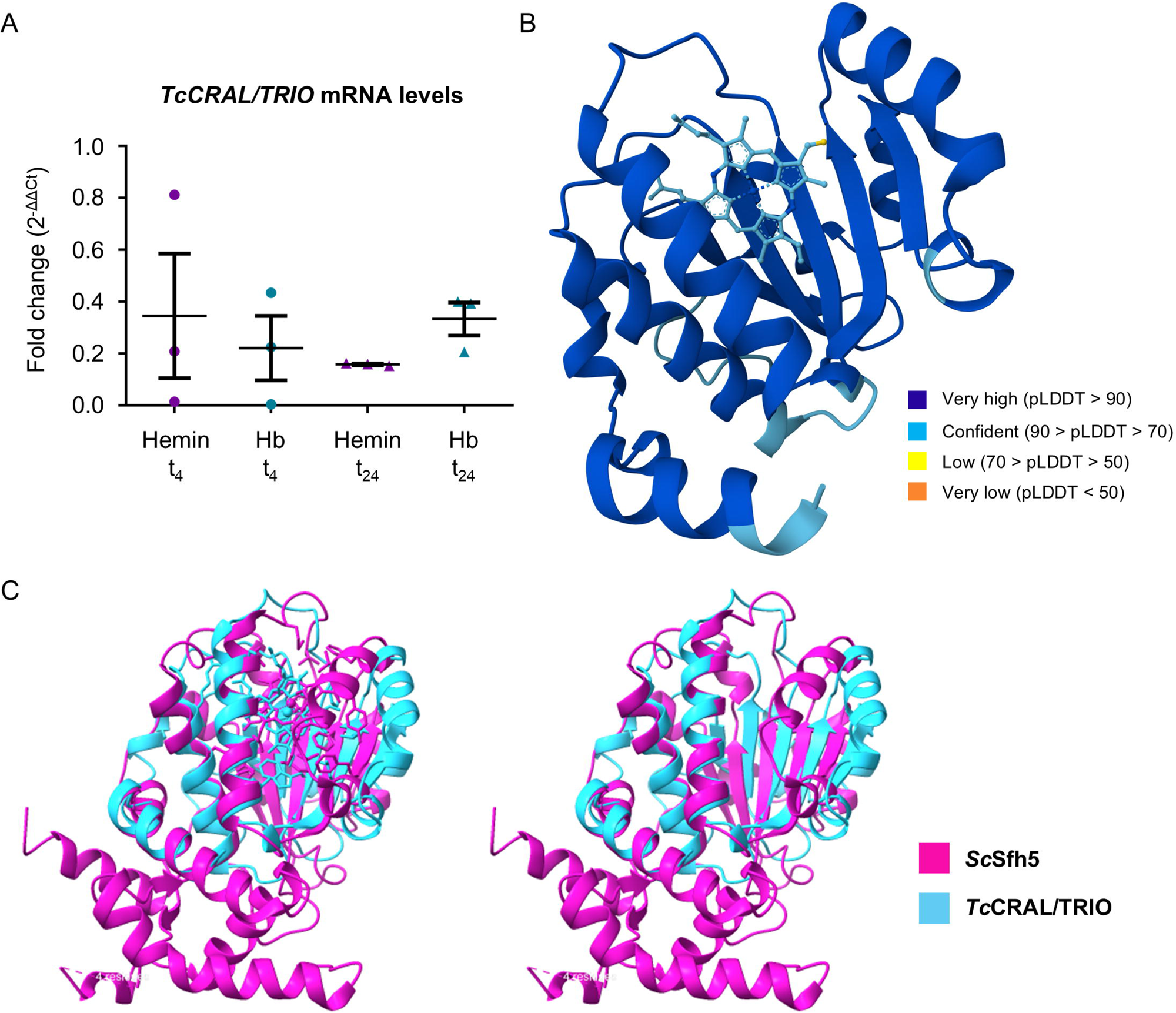
**(A)** Quantification of *TcCRAL/TRIO* mRNA levels in epimastigotes cultured in medium supplemented with 5 µM hemin or 1.25 µM hemoglobin (Hb) for 4 h (t_4_) or 24 h (t_24_). Quantification was performed by RT-qPCR, using *Tcubiquitin* as the reference gene. Data represent the mean ± SD of relative mRNA levels of three biological replicas. mRNA levels were relativized to those of no heme t_4_ sample (for hemin and Hb t_4_ samples) and no heme t_24_ sample (for hemin and Hb t_24_ samples). **(B)** Predicted 3D structure of *Tc*CRAL/TRIO in complex with heme, generated using AlphaFold. The model is color-coded by predicted Local Distance Difference Test (pLDDT) confidence scores. Most of the structure is shown in blue, indicating very high confidence (pLDDT > 90), with a few regions in cyan corresponding to confident predictions (70 < pLDDT ≤ 90). **(C)** Structural overlay of the predicted *Tc*CRAL/TRIO (cyan) with the crystal structure of *Sc*Sfh5 from *S. cerevisiae* (pink), shown in complex with heme (left panel) and without heme (right panel) to facilitate visualization.

Then, we predicted its 3D structure using AlphaFold3 [16], which suggests its potential to coordinate heme. The five AlphaFold-predicted models (Figure S2) exhibited reliable structural features, supported by high-ranking confidence scores—ranking values ranging from 0.80 to 0.86, iPTM scores between 0.78 and 0.85, and pTM scores from 0.88 to 0.89—indicating good overall model quality (Table S7). Figure 7B shows the highest-ranked model. Despite sharing only 26.7% amino acid sequence identity, the comparison with the published crystal structure of *Sc*Sfh5 (PDB ID: 6W32) [15] revealed similar tertiary structures exhibiting the typical characteristics of a CRAL/TRIO domain (Figure 7C).

Finally, we expressed *Tc*CRAL/TRIO in *Escherichia coli* as a fusion protein with Maltose Binding Protein (MBP) and a His-tag, followed by purification using Ni-NTA affinity chromatography. Both the bacterial pellet and the purified recombinant protein displayed a characteristic brownish color (Figure 8A), and UV-Vis spectrophotometric analysis of the purified recombinant *Tc*CRAL/TRIO revealed a Soret peak at 407 nm (Figure 8B) consistent with the presence of bound heme [17]. This result constitutes the first evidence of a heme responsive gene that encodes a hemoprotein in *T. cruzi*.

**Figure 8.**
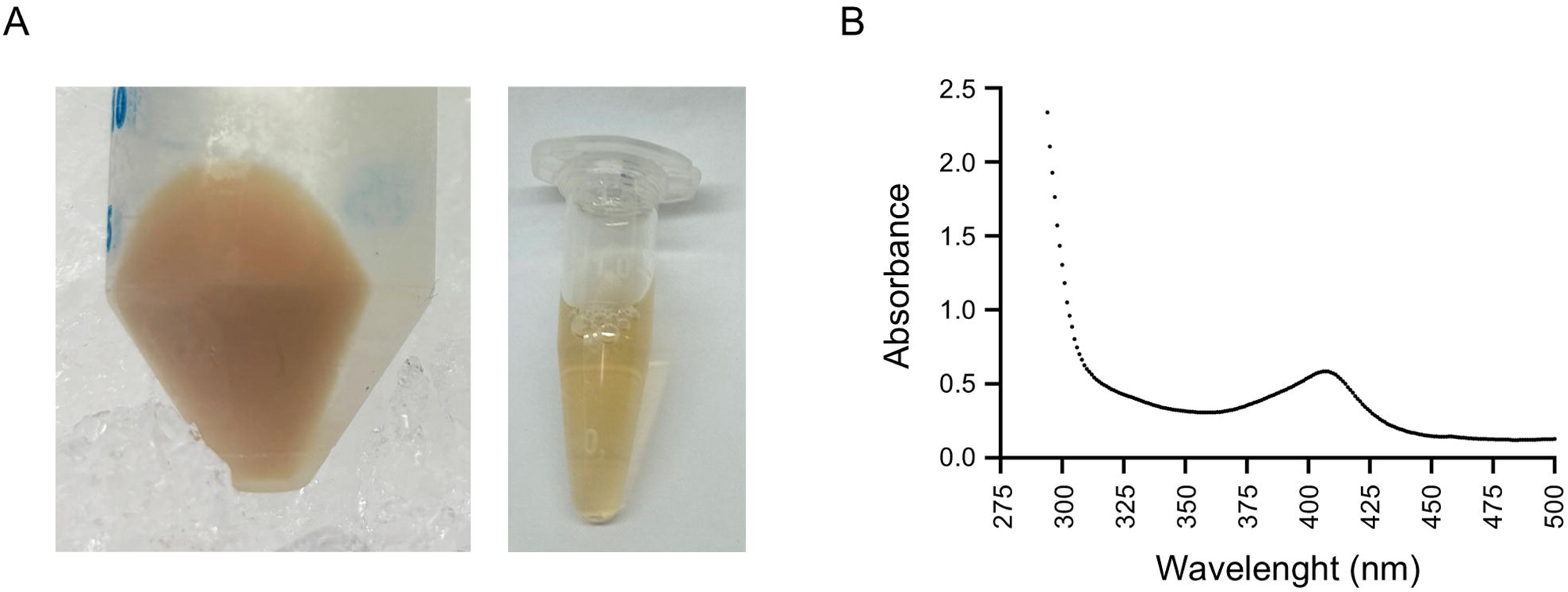
Heterologous expression and purification of *Tc*CRAL/TRIO. (A) *E. coli* cell pellet expressing *Tc*CRAL/TRIO as a fusion to Maltose Binding Protein (MBP) (left panel) and the corresponding purified *Tc*CRAL/TRIO protein (right panel). (B) UV-visible absorption spectrum of the purified *Tc*CRAL/TRIO, showing a characteristic Soret peak at 407 nm.

## DISCUSSION

Heme uptake and homeostasis have been a major focus of our research for years. We have demonstrated that *T. cruzi* acquires heme during its replicative stages, with *Tc*HRG playing a key role in this process [4, 8, 9].

To investigate how *T. cruzi* epimastigotes adapt to heme deprivation and replenishment, we performed RNA-seq after controlled supplementation (5 μM, provided as hemin or Hb equivalents) to heme-starved parasites. We performed a three-level RNA-seq analysis to fully exploit the data. First, we conducted a general overview (Section 2). Then, we compared transcriptomes against the starvation baseline (t_0_; Sections 2a–e) and finally, we compared heme-treated epimastigotes to heme-deprived ones (Section 3). This approach revealed key transcriptional shifts at early time points, offering insight into the parasite’s response to different heme sources. Across all analyses, *TcHRG* and several other genes consistently appeared among downregulated DEGs, with subtle global changes (logFC < ∣1.8∣). Interestingly, hemin modulated more gene expression than Hb, and the number of DEGs increased over time with hemin (t_4_ < t_24_). Conversely, Hb addition showed fewer DEGs at t_24_ compared to t_4_, and a subset of t_4_ DEGs maintained their expression at t_24_. Our third-level analysis identified 22 consistently regulated genes (Section 3) with overlapping subsets across all conditions, which we consider the central transcriptional program induced by heme availability in *T. cruzi* epimastigotes.

Notably, among the downregulated genes in the first global analysis, we identified seven genes encoding glutamyl carboxypeptidases and two encoding metallo-peptidases. All these genes are predicted to encode enzymes with acetylornithine deacetylase (AOD) activity, typically associated with arginine biosynthesis. Recently, it was reported that an AOD from *Arabidopsis thaliana* can function as a Cys-Gly dipeptidase implicated in the glutathione degradation pathway [18]. Although very little is known about glutathione and trypanothione degradation pathways in *T. cruzi*, the possibility that these enzymes serve a similar function in thiol metabolism is intriguing. If true, their downregulation would align with the upregulation of genes involved in trypanothione biosynthesis observed in our second analysis discussed next, suggesting a coordinated response to oxidative stress or altered redox balance.

Pathways involved in heme metabolism and oxidative stress response in *T. cruzi* arose as the core regulation as they appear in the overlapping hemin∩Hb DEGs. We observed significant upregulation of key components of the parasite’s antioxidant defense system, including enzymes involved in trypanothione biosynthesis—the primary thiol-redox buffer in trypanosomatids [19]—peroxidoxins that directly neutralize reactive oxygen species, and NADPH-producing enzymes such as G6PD and malic enzyme that maintain reducing equivalents for antioxidant systems. In fact, it was reported that G6PD of *T. cruzi* is resistant to DTT inactivation and strongly upregulated under oxidative stress (up to 46-fold; [20]). These observations highlight the critical role of NADPH regeneration in counteracting heme-induced oxidative damage.

Concurrently, we detected downregulation of genes encoding cytochromes and other mitochondrial electron transport chain components, suggesting a metabolic shift away from aerobic respiration following heme exposure. This observation aligns with previous reports that heme inhibits cytochrome c oxidase activity in *T. cruzi*, leading to reduced oxygen consumption and increased mitochondrial ROS production [6]. Another possible interpretation is that heme starvation triggers increased production of key apohemoproteins to facilitate the capture and incorporation of the limited heme into functional complexes. Upon heme repletion, the expression of these genes is downregulated, as sufficient protein reserves have already been synthesized.

The observed downregulation of arginine permease and arginine kinase genes may reflect a shift in energy storage and utilization following heme refeeding. *T. cruzi* epimastigotes convert L-arginine into L-phosphoarginine via arginine kinase, generating phosphagens that serve as energy reservoirs under nutrient-limiting conditions [21, 22], to support essential processes until metabolic pathways are restored. This finding suggests that the need for this energy-buffering system is reduced once normal metabolic activity resumes.

Similarly, the downregulation of genes encoding oleate desaturase and associated proteins suggests a reduction in desaturase activity in response to heme supplementation. A homolog of oleate desaturase has been characterized in *T. brucei*, where it contributes to the production of unsaturated fatty acids, modulating membrane fluidity [23]. It is plausible that these genes were upregulated during heme starvation to initiate metacyclogenesis and enhance membrane fluidity [24], indicating an adaptation to changes in nutrients availability. Heme refeeding may trigger a return to proliferative epimastigote form, reducing the need for such adaptations and leading to the observed transcriptional downregulation.

Altogether, these findings suggest that heme availability not only modulates redox balance and energy metabolism but also contributes to reverse starvation-induced differentiation processes, likely through signaling pathways yet to be elucidated. The hemin∩Hb dataset provides compelling evidence that *T. cruzi* employs a dual strategy to manage heme toxicity: (1) rapid induction of ROS-scavenging systems to counteract oxidative damage, and (2) metabolic adjustments to minimize further ROS production.

Complementary to this core transcriptional signature that balances heme utilization and detoxification, we also found heme source-specific metabolic adaptations and subtle yet critical variations across the three culture conditions examined. For instance, unique subsets of hemin- and Hb-exclusive DEGs revealed the presence of various kinases and phosphatases likely associated with distinct signaling pathways. This suggests a complex, source-dependent modulation of intracellular signaling mechanisms that govern the parasite’s metabolic reprogramming.

On the contrary, several observations in the hemin- or Hb-specific datasets also reinforce findings from the shared hemin∩Hb group. For example, Hb-supplemented parasites showed additional downregulation of genes related to fatty acid metabolism, while hemin-supplemented parasites exhibited upregulation of further glycolytic genes. Moreover, both conditions revealed additional downregulated genes encoding mitochondrial electron transport proteins. These findings support the model proposed by Paes et al., in which heme triggers a metabolic shift from mitochondrial respiration to aerobic fermentation in *T. cruzi* [25].

Remarkably, the rapid (t_4_) upregulation of 33 cruzipain genes after Hb exposure implies an immediate requirement for proteolytic Hb processing for heme acquisition that was not observed when providing hemin, validating a specific proteolytic response. Supporting this idea, a significant portion of cruzipain is secreted [26] or associated with the epimastigote surface [27], also reinforcing the model we proposed for Hb-derived heme uptake [9].

Regarding hemin-supplemented parasites, on the other hand, a strong downregulation of genes encoding TS and MASP was found. While most TS and MASP genes are predominantly expressed in trypomastigotes, certain members of these gene families are also active in epimastigotes [28]. Although not directly assessed here, it is possible that starvation may have triggered metacyclogenesis, inducing TS and MASP expression. Again, hemin refeeding reversed this process, promoting a return to the epimastigote stage and suppressing the expression of these surface proteins.

We also observed downregulation of genes encoding key flagellar components, particularly PRPs and FCaBPs. In trypanosomatids, certain flagellum-localized transporters may function as “transceptors”—proteins that not only transport molecules but also sense and respond to extracellular concentrations of their cargo ([29] and references therein)—and FCaBPs not only mediate flagellar function and assembly but also modulate other flagellar proteins in a Ca²⁺-dependent manner, suggesting additional signaling roles [30]. This downregulation may reflect a metabolic adaptation where active environmental sensing becomes less essential following hemin-mediated alleviation of nutritional stress.

Analysis of heme-deprived samples identified downregulation of 15 TAT genes as early as t_4_. In *Leishmania donovani*, deletion of a TAT ortholog disrupts redox homeostasis by increasing ROS and reducing tocopherol levels [31]. This evidence suggests that in trypanosomatids TATs are implicated in redox metabolism beyond their canonical role in amino acid metabolism. Consistent with redox modulation, two ascorbate peroxidase genes (essential for oxidative stress defense, [32]) were downregulated at t_24_. These coordinated transcriptional changes further demonstrate the capacity of *T. cruzi* to dynamically regulate antioxidant defenses in response to heme availability.

At the deepest level of RNAseq analysis (Figure 6), upregulated DEGs include the genes encoding pyridoxal kinase and enolase, both of which were also highlighted in the second analysis. Also, the protein encoded by C4B63_70g116, which would belong to the Amino acid-Polyamine-Organocation Family, may be involved in the transport of some of the trypanothione precursors.

Regarding downregulated DEGs, some of them encode proteins with known roles in iron and heme metabolism, such as *Tc*FR and *Tc*HRG. Others, including CRAL/TRIO domain-containing protein and RNA-binding protein, may have relevant functions as inferred from homologs in other organisms, although their specific connections to heme metabolism remain to be clarified.

*Tc*FR catalyzes the reduction of Fe^3+^ to Fe^2+^, is membrane-associated, and is upregulated under iron-limitation in absence of heme as shown by Dick et al. [12]. Moreover, the ferric reductase related protein 1 (Frp1) from *Candida albicans* is essential for heme uptake and relocalizes to the plasma membrane in the presence of heme [33]. Taken together, *Tc*FR regulation in our dataset is consistent with its proposed function and suggests it may reduce the heme Fe³⁺ prior to internalization.

Notably, the RNA-binding protein encoded by C4B63_27g1020c shares 68% amino acid sequence identity with RBP5 from *T. brucei*, reported to be upregulated under iron-starvation conditions in the bloodstream form of this parasite [34]. This similarity suggests the protein may have a conserved role in regulating stress-responsive genes under iron or heme deprivation.

The two ABC transporters (C4B63_10g454 and C4B63_10g455) identified, if related to heme uptake, could be downregulated by the parasite when heme is available, like *TcHRG*. This is supported by the reported intracellular heme decrease in the presence of ABC transporter-specific inhibitors, which was not reversed by heme supplementation [35].

As for the putative peptides encoded by the cAMP-dependent protein kinase genes C4B63_24g988c and C4B63_69g318c, which were found downregulated in the hemin-supplemented samples at t_24_, and in the first analysis across both hemin- and Hb-supplemented parasites, may be components of a signaling pathway involved in sensing and responding to heme depletion.

Finally, the reproducible downregulation of *Tc*CRAL/TRIO after heme-replenishment coupled with its confirmed heme-binding capacity, suggests an involvement in heme homeostasis. However, its precise role -whether in transport, buffering, or sensing-requires further validation.

The dual nature of heme as both essential cofactor and potential toxin requires strict homeostatic control in *T. cruzi*. Our transcriptomic analysis delineates the parasite’s response to different heme sources, identifying both conserved regulatory elements and source-specific adaptations. Heme restitution to starved epimastigotes induced rapid but moderate transcriptomic changes, consistent with a physiological rebalancing rather than a stress response. This experimental design employing standard heme concentrations [4, 8]following controlled starvation specifically captured the parasite’s adaptive mechanisms while minimizing non-physiological effects.

The identification of heme-responsive factors such as *Tc*CRAL/TRIO uncovers previously uncharacterized components of the heme management system in *T. cruzi*. These findings enhance our mechanistic understanding of the parasite’s biology and establish a foundation for exploring heme metabolism as a potential therapeutic target. Given the obligate dependence of *T. cruzi* on host-derived heme, selectively interfering with its acquisition or utilization pathways may offer novel strategies for Chagas disease intervention.

## MATERIALS AND METHODS

### Reagents

Hemin (Frontier Scientific, Logan, UT, USA) and Hb (Sigma, Burlington, MA, USA) stock solutions were prepared, quantified and stored as described [9]. Fetal Bovine Serum (FBS) (Internegocios SA, Mercedes, BA, Argentina) was subjected to heat inactivation for 30 minutes at 56 °C prior to use. Luria-Bertani broth (LB), Liver infusion broth and tryptose broth were purchased from BD Difco (Franklin Lakes, NJ, USA). All reagents used were of molecular biology grade.

### Parasite culture and experimental design

*T. cruzi* epimastigotes (Dm28c strain) were cultured at 28 °C in Liver Infusion Tryptose (LIT) medium supplemented with 10 % heat inactivated FBS (LIT-10% FBS) plus 5 μM hemin and maintained in mid-log phase by successive dilutions in fresh medium [8].

For heme starvation, epimastigotes were collected, washed twice with PBS, resuspended in fresh medium without heme (at 30×10^6^ epimastigotes/mL), and maintained for 48 hours. Then, parasites were harvested, washed twice with PBS, and resuspended at 30×10^6^ epimastigotes/mL in fresh medium without heme or containing 5 μM hemin or 1.25 μM Hb.

Samples were taken at different time points for western blotting, RT-qPCR, and RNA-seq, as specified below. Parasites were counted using Neubauer chamber and a Counter 19 Auto Hematology Analyzer (Wiener Laboratorios SAIC, Rosario, SF, Argentina) set up for parasite number measurements.

### Western blot

Samples for Western blotting were prepared and processed as previously described [9], including the use of the same primary [8] and secondary antibodies for *Tc*HRG detection and protein loading evaluation.

### Quantitative real-time PCR (RT-qPCR)

Total mRNA isolation and quantification was carried out as described previously [8, 9]. RNA was treated with DNAse (RQ1 RNAsa-Free Dnase, Promega, Madison, WS, USA) and retrotranscribed with FireScript KIT Reverse Transcriptase (Solis BioDyne, Tartu, Estonia) to obtain cDNA. qPCRs were performed with HOT FIREPol EvaGreen qPCR Mix Plus (ROX) (Solis BioDyne, Tartu, Estonia), using the primers listed below (Macrogen, Korea), in a StepOne (Applied Biosystem, ThermoFisher Scientific, Foster city, CA, USA) or, alternatively, a CFX Opus 96 Real-Time PCR System (#12011319 - Bio-Rad, Miami, FL, USA) following the protocol: 95 °C (10 min); 40 cycles of 95 °C (10 s), 60 °C (30 s), and 72 °C (20 s); followed by the denaturation curve to analyze the Tm (melting temperatures) of the products. The efficiency of each pair of primers was determined and the relative fold changes were calculated by the 2^-ΔΔCt^ method [36].

The primers used for RT-qPCR were (accession numbers, between brackets): *TcHRG* (TcCLB.511071.190) Fw: TAATTATTGGGCGGCGGCT, Rv: GAAGTACGAACTCCCCGTCC; *TcCRAL/TRIO* (C4B63_18g39) Fw: GTTGGGTGCTCTTCTTCTCT, Rv: ACCTTCTTTCGGGTACGTTT; *TcGAPDH* (TcCLB.506943.50) Fw: GTGGCAGCACCGGTAACG, Rv: CAGGTCTTTCTTTTGCGAATAGG and *TcUbiquitin* (BCY84_18965) Fw: AGGGCATTCCGGGAAAGATG, Rv: CCACCACCATGTGCAGAGTT, based on nucleotide sequences found on the annotated genomes of *T. cruzi* CL Brenner Esmeraldo-like (*TcHRG*, *TcGAPDH*), Dm28c strain 2017 (*TcUbiquitin*) and Dm28c strain 2018 (*TcCRAL/TRIO*).

For *TcHRG* mRNA quantification, *TcGAPDH* was used as reference (housekeeping) for sample normalization, and the value of starved epimastigote (t_0_) was assigned as mRNA level = 1 (reference). For *TcCRAL/TRIO* mRNA quantification, *TcUbiquitin* was used as reference for sample normalization, and cultures incubated without heme at the corresponding time points were used as the reference condition (mRNA level = 1).

### RNAseq assay

Three biological replicas were performed, from which triplicate samples were taken at the established time points for the heme-deprived, hemin and Hb conditions. Total RNA was obtained from these samples, and one complete set was subjected to RNAseq assay (Macrogen, Inc., Republic of Korea) while the other two were backed up for latter validation by RT-qPCR assays.

The transcriptome of the samples was sequenced in a Next Generation Sequencing (NGS) platform. Libraries were synthesized with Illumina TruSeq total RNA sample preparation kit with Ribo-Zero Gold. Sequencing was performed with Illumina NovaSeq6000 technology, selecting paired-end reads of 101 pb length.

### Analysis of RNA-seq data

The quality of raw reads obtained from the RNA-seq assay was assessed using *FastQC* [37]. Adapter trimming and low-quality base removal were performed using *sickle*. Reads from each library were aligned to the *Trypanosoma cruzi* Dm28c 2018 reference genome (available at *TriTrypDB*: https://tritrypdb.org) using *Bowtie2* [38], with the “end-to-end” alignment mode. The resulting SAM alignment files were converted to BAM format and indexed using *SAMtools* [39] for downstream processing and to obtain mapping statistics.

Gene-level quantification was performed using *featureCounts* [40], restricting counting to coding sequences (CDS). A global count matrix was generated, and differential gene expression analysis (DEG) was conducted in R using the *DESeq2* package [41].

Normalized counts were visualized using heatmaps generated with the *pheatmap* package. Additional data exploration and visualization were carried out using R packages including *dplyr*, *ggplot2*, *RColorBrewer*, *ggrepel*, *stringr*, *hrbrthemes*, *GGally*, and *viridis*.

Gene Ontology (GO) enrichment analysis was conducted using the tools available on the TriTrypDB platform, with statistical significance determined by a Bonferroni-adjusted p-value threshold of < 0.1. Redundant GO terms were removed by manual curation to enhance result clarity. The results were visualized as bar plots generated in R with custom scripts.

### *Tc*CRAL/TRIO expression and purification

A pET32 plasmid construct [42] encoding the 6His–MBP–TEV–*Tc*CRAL/TRIO fusion protein was transformed into *E. coli* BL21 (DE3) pLysS cells. Expression was induced with 100 μM IPTG and carried out overnight at 20 °C. Cells were resuspended in Lysis Buffer (100 mM HEPES, 1 M NaCl, 2 % (v/v) Triton X-100, 2 mM PMSF, pH 7.4) and lysed using a 750 W ultrasonic processor (Cole Parmer, Vernon Hills, IL, USA). The clarified lysate was incubated with Ni Sepharose™ 6 Fast Flow resin (GE Healthcare, Chicago, IL, USA) in the presence of Binding Buffer (100 mM HEPES, 1 M NaCl, 30 mM imidazole, pH 7.4), followed by a washing step using the same buffer. The His-tagged protein was eluted with buffer containing 250 mM imidazole and analyzed by SDS-PAGE with Coomassie staining. UV-visible spectra of the eluates were recorded using a GeneQuant 1300 spectrophotometer in quartz cuvettes.

### In silico analysis

Amino acid sequence alignments were performed using Clustal Omega [43]. The predicted 3D structure of *Tc*CRAL/TRIO bound to heme was generated using the AlphaFold algorithm via the AlphaFoldServer web service (https://alphafoldserver.com/), following the method described by Jumper et al. [16]. Structural visualization, model overlay, and comparison with the crystal structure of *S. cerevisiae* Sfh5 were performed using ChimeraX [44].

### Statistical analysis

Western blot analysis to assess *Tc*HRG protein levels were independently reproduced two times. RT-qPCR analysis was independently reproduced three times. Statistically significant differences between groups in the RT-qPCR data were determined using GraphPad Prism version 6.00 for Windows (GraphPad Software, San Diego, CA).

Transcriptomic analysis was performed using three independent biological replicas. Quantification of *Tc*CRAL/TRIO mRNA was conducted on samples taken from the same cultures used for the transcriptomic assay. *Tc*CRAL/TRIO protein purification experiments were independently reproduced at least three times.

## Data Availability

All relevant data supporting this study are included in the main article and supplementary material. The complete set of raw data has been deposited into the Repositorio de Datos Académicos UNR (RDA UNR) https://dataverse.unr.edu.ar/, the institutional data repository of the Universidad Nacional de Rosario. This open-access platform enables the dissemination and long-term preservation of academic research data, adheres to FAIR principles and is built on Dataverse Project technology. Additionally, the dataset is assigned a persistent and unique Digital Object Identifier (DOI) provided by RDA UNR number (https://doi.org/10.57715/UNR/BPKC2V).

## ABBREVIATIONS

ANOVA: analysis of variance;
DEG: differentially expressed gene;
FBS: fetal bovine serum;
GO: gene ontology;
Hb: hemoglobin;
LIT: liver infusion tryptose;
MBP: maltose binding protein;
PBS: phosphate buffered saline;
RNA-Seq: RNA sequencing;
ROS: reactive oxygen species;
RT-qPCR: reverse transcription followed by quantitative real- time PCR;
*Tc*HRG: *Trypanosoma cruzi* Heme Responsive Gene.

## ACKNOWLEDGMENTS

We thank Javier Jalife, for his valuable assistance with informatics support.

This research was partially supported by funding from UK Research and Innovation through the Global Challenges Research Fund under grant agreement ‘A Global Network for Neglected Tropical Diseases’ (grant number MR/P027989/1) awarded to JAC (2018–2022), and by the National Agency for Scientific and Technological Promotion (ANPCYT) under grant number PICT I A 2021-00274 awarded to JAC (2023–2027).

ET was supported by a post-doc research fellowship from ANPCYT associated with PICT I A 2021-00274. JAC and MGM are researchers of National Scientific and Technical Research Council (CONICET). MLM was supported by a post-doc research fellowship from CONICET. CBDC was supported by a post-doc research fellow associated to grant agreement ‘A Global Network for Neglected Tropical Diseases’ (grant number MR/P027989/1). The Institut Pasteur de Montevideo is funded by FOCEM – Fondo para la Convergencia Estructural del Mercosur (grant number COF 03/11). LB and CR are researchers of PEDECIBA and the national system of researchers (SNI-ANII).

Molecular graphics and analyses were performed with UCSF ChimeraX, developed by the Resource for Biocomputing, Visualization, and Informatics at the University of California, San Francisco, with support from National Institutes of Health R01- GM129325 and the Office of Cyber Infrastructure and Computational Biology, National Institute of Allergy and Infectious Diseases.

---

**Figure S1.** Volcano plots showing differentially expressed genes (DEGs) across various culture conditions and time points. Each plot displays log₂(fold change) vs. – log₁₀(adjusted p-value) for genes identified under hemin and hemoglobin (Hb) conditions. Timepoint comparisons include t_4_ vs. t_0_ and t_24_ vs. t_0_. Genes with |log₂(fold change)| < 0.4 and –log₁₀(adjusted p-value) < 1.3 are shown in gray (not significantly regulated). Genes with 0.4 ≤ |log₂(fold change)| ≤ 0.7 and –log₁₀(adjusted p-value) > 1.3 are shown in yellow. Genes with |log₂(fold change)| > 0.7 and –log₁₀(adjusted p-value) > 1.3 are shown in red.

**Figure S2.** Structural overlay of the five 3D structures of *Tc*CRAL/TRIO predicted by Alphafold.

**Table S1.** List of 51 DEGs in first global analysis.

**Table S2.** List and Gene Ontology analysis of DEGs common to hemin, hemoglobin and no heme conditions in the t_4_ vs. t_0_ and t_24_ vs. t_0_ analysis.

**Table S3**. List and Gene Ontology analysis of DEGs common to hemin and hemoglobin conditions in the t_4_ vs. t_0_ and t_24_ vs. t_0_ analysis.

**Table S4.** List and Gene Ontology analysis of DEGs exclusively modulated in hemin, Hb or no heme conditions in the t_4_ vs. t_0_ and t_24_ vs. t_0_ analysis. Highlighted in yellow genes probably involved in signaling pathways.

**Table S5.** List of genes with sustained differential expression through the t_4_ vs. t_0_ and t_24_ vs. t_0_ analysis in hemin and hemoglobin (Hb) conditions.

**Table S6**. List of DEGs identified in the heme-supplemented conditions vs. no heme condition analysis at t_4_ and t_24_.

**Table S7**. Confidence scores of the five AlphaFold predicted 3D structures of *Tc*CRAL/TRIO. Tabulated version of the JSON data from Alphafold for ease of viewing.

**Supplementary Files 1-5**. Raw output of confidence scores from AlphaFold for the five predicted 3D models of *Tc*CRAL/TRIO.

