## Supplementary material for "Transcriptomic response to different heme sources in *Trypanosoma cruzi*"

### Table S1

| DEGs (heme-supplemented vs. heme-starved) | | | | |
|---|---|---|---|---|
| ID | Annotation | log2FC | pvalue | padj |
| C4B63_3g227 | mucin TcMUCII | -0.32993493357209 | 0.00011690440045204 | 0.0405268588233739 |
| C4B63_462g11 | glutamamyl carboxypeptidase | -0.607368478302711 | 4.24019511270686e-05 | 0.0229309751695187 |
| C4B63_322g15 | amino acid transporter | 0.234546086764434 | 3.58640665365175e-05 | 0.0229309751695187 |
| C4B63_27g1020c | RNA-binding protein | -0.895786305604263 | 4.14698768510786e-14 | 2.80336367513291e-10 |
| C4B63_70g88 | ATG16/SAP18/CVT11/APG16 | -0.249169217702849 | 1.02047991183225e-06 | 0.00197098405828172 |
| C4B63_462g9 | glutamamyl carboxypeptidase(fragment) | -0.662535100092884 | 5.90494491750712e-05 | 0.0262631409519877 |
| C4B63_313g13 | amino acid transporter | 0.203116115381241 | 0.00010988706767567 | 0.0405268588233739 |
| C4B63_27g184 | integral membrane transport protein | -0.51462531309972 | 1.20892719481177e-14 | 1.63446956738551e-10 |
| C4B63_462g10 | glutamamyl carboxypeptidase | -0.602732341729182 | 0.000105610670822974 | 0.0405268588233739 |
| C4B63_313g15 | amino acid transporter | 0.239765565807958 | 1.40756608262442e-05 | 0.0126868622913881 |
| C4B63_350g5 | glutamamyl carboxypeptidase(fragment) | -0.542200227815286 | 0.000155766739039081 | 0.0438742981626743 |
| C4B63_124g19 | surface protease GP63 | 0.339239427118475 | 0.000136428336729589 | 0.0434440241343718 |
| C4B63_257g5 | amino acid transporter | 0.192040528159319 | 5.73521667723818e-05 | 0.0262631409519877 |
| C4B63_282g15 | cation transporter | 0.354507535145914 | 3.33892826515104e-07 | 0.000902846202896841 |
| C4B63_313g11 | amino acid transporter | 0.224776052719636 | 0.000159036340427043 | 0.0438810473994617 |
| C4B63_4g180 | serine/threonine protein kinase | -0.326450422207995 | 6.02187403484926e-05 | 0.0262631409519877 |
| C4B63_6g236 | cation transporter | 0.344007513437361 | 2.72284475063006e-06 | 0.0040903178920576 |
| C4B63_51g83 | ferric reductase transmembrane protein | -0.387411987873575 | 4.22180308878258e-05 | 0.0229309751695187 |
| C4B63_2g799 | tRNA wybutosine-synthesizing protein 4 | 0.202922242722498 | 6.35602707145551e-05 | 0.0268542143768995 |
| C4B63_336g4 | glutamamyl carboxypeptidase | -0.592696848892484 | 5.52795498147318e-05 | 0.0262631409519877 |
| C4B63_98g22 | trans-sialidase | -0.529646115656052 | 0.000127349481582203 | 0.0419942680729605 |
| C4B63_550g1 | metallo-peptidase, clan MH, family M18 | -0.619000120633786 | 0.00014449547263427 | 0.0434440241343718 |
| C4B63_34g375 | Soluble NSF attachment protein, SNAP (paraflagelar rod component) | -0.225841931522567 | 4.12696463479452e-05 | 0.0229309751695187 |
| C4B63_20g314 | serine/threonine-protein phosphatase PP1 | -0.193009470011326 | 0.000140162109283095 | 0.0434440241343718 |
| C4B63_42g103 | pyridoxal kinase | 0.341502061627782 | 0.000123297045199435 | 0.0416744012774091 |
| C4B63_19g205 | glycosomal transporter (GAT3) | 0.28984712801925 | 3.65675309403849e-06 | 0.00426790262536025 |
| C4B63_257g6 | amino acid transporter | 0.216358150969246 | 1.38970948346949e-05 | 0.0126868622913881 |
| C4B63_40g93 | conserved hypothetical protein | -0.231767513706 | 0.000155321486507511 | 0.0438742981626743 |
| C4B63_97g13 | conserved hypothetical protein | -0.276476125660578 | 0.000185029265017814 | 0.0490508953537422 |
| C4B63_313g14 | amino acid transporter | 0.272532299551567 | 2.08137954740411e-06 | 0.00351753143511294 |
| C4B63_258g25 | lipase | -0.415942368533025 | 8.78493093583167e-05 | 0.034933019486013 |
| C4B63_25g220 | 5'-3' exoribonuclease B | -0.315531752492431 | 0.00011152169768915 | 0.0405268588233739 |
| C4B63_51g81 | ferric reductase transmembrane protein | -0.377068931146649 | 0.000150804550091205 | 0.0438742981626743 |
| C4B63_51g82 | ferric reductase transmembrane protein | -0.375293796747228 | 5.42215317315641e-05 | 0.0262631409519877 |
| C4B63_322g14 | amino acid transporter | 0.247598942062281 | 1.71663653409714e-05 | 0.0145055787131208 |
| C4B63_44g117 | conserved hypothetical protein | 0.433996675126999 | 2.86095811455705e-05 | 0.0203579756362165 |
| C4B63_3g1092 | alanine racemase | -0.322947119951655 | 1.9330081790921e-05 | 0.015373100341956 |
| C4B63_69g318c | conserved hypothetical protein | -0.48388155806861 | 8.03778715482471e-06 | 0.0083592986410177 |
| C4B63_313g10 | amino acid transporter | 0.230611418184654 | 2.76686277567821e-05 | 0.0203579756362165 |
| C4B63_18g39 | CRAL/TRIO, N-terminal domain/CRAL/TRIO domain containing protein | -1.3228074513183 | 6.81009516320881e-08 | 0.000230181216516458 |
| C4B63_6g397 | cation transporter | 0.315973108034771 | 3.38440211331048e-05 | 0.0228785582859788 |
| C4B63_550g3 | metallo-peptidase, clan MH, family M18 | -0.601216854207726 | 0.00011538420281901 | 0.0405268588233739 |
| C4B63_336g1 | glutamamyl carboxypeptidase(fragment) | -0.48642113836089 | 0.0001685381192624 | 0.0455727074485531 |
| C4B63_354g1 | electron transfer flavoprotein | -0.414503719480781 | 3.77909817901243e-06 | 0.00426790262536025 |
| C4B63_51g85 | ferric reductase transmembrane protein | -0.379931647551028 | 3.86816852250706e-05 | 0.0229309751695187 |
| C4B63_31g178 | Zinc phosphodiesterase ELAC protein 2 | 0.27326856751695 | 5.95020306729887e-05 | 0.0262631409519877 |
| C4B63_294g12 | glutamamyl carboxypeptidase | -0.580367898754649 | 0.000144599192754936 | 0.0434440241343718 |
| C4B63_32g209 | Heme Responsive Gene (*Tc* HRG) | -0.487727859236573 | 1.21478070041169e-08 | 5.47461168985533e-05 |
| C4B63_19g141 | electron transfer flavoprotein | -0.402413527517146 | 4.29916698829893e-07 | 0.000968745628030025 |
| C4B63_24g988c | conserved hypothetical protein | -0.497007686289135 | 3.78807925327833e-06 | 0.00426790262536025 |
| C4B63_51g80 | ferric reductase transmembrane protein | -0.355989571436148 | 6.67571892839377e-05 | 0.0273502181551163 |

### Table S2

#### Hemin, Hb and No heme common DEGs (t4 vs. t0)

##### Downregulated

| ID | Annotation |
|---|---|
| C4B63_102g376c | conserved hypothetical protein |
| C4B63_10g157 | Double RNA binding domain protein 6A |
| C4B63_112g40 | surface protease GP63 |
| C4B63_129g44 | cytochrome c oxidase VIII (COX VIII) |
| C4B63_12g242 | SKP1-like protein |
| C4B63_12g405 | cytochrome P450 |
| C4B63_137g16 | cytochrome c oxidase assembly protein |
| C4B63_13g200 | Enriched in surface-labeled proteome protein 18 |
| C4B63_13g327 | conserved hypothetical protein |
| C4B63_140g32 | cytochrome c oxidase VIII (COX VIII) |
| C4B63_14g1 | hexose transporter |
| C4B63_14g2 | hexose transporter(fragment) |
| C4B63_14g28 | hexose transporter |
| C4B63_14g3 | hexose transporter |
| C4B63_14g30 | hexose transporter |
| C4B63_14g5 | hexose transporter |
| C4B63_155g10 | trans-sialidase |
| C4B63_163g21 | deoxyribose-phosphate aldolase |
| C4B63_16g136 | conserved hypothetical protein |
| C4B63_17g48 | ubiquitin-conjugating enzyme E2 |
| C4B63_18g1130c | FUN14 family |
| C4B63_196g32 | Mucin-associated surface protein (MASP) |
| C4B63_19g86 | CYC2-like cyclin 6 |
| C4B63_1g756 | surface protease GP63 |
| C4B63_212g5 | protein associated with differentiation 4(fragment) |
| C4B63_21g10 | arginine permease |
| C4B63_21g14 | arginine permease |
| C4B63_21g16 | arginine permease |
| C4B63_21g18 | arginine permease |
| C4B63_21g20 | arginine permease |
| C4B63_21g21 | arginine permease |
| C4B63_21g5 | arginine permease |
| C4B63_21g56 | nucleoside phosphorylase |
| C4B63_21g65 | conserved hypothetical protein |
| C4B63_21g9 | arginine permease |
| C4B63_228g46 | glutamamyl carboxypeptidase |
| C4B63_228g47 | glutamamyl carboxypeptidase |
| C4B63_228g48 | glutamamyl carboxypeptidase |
| C4B63_228g49 | glutamamyl carboxypeptidase |
| C4B63_228g50 | glutamamyl carboxypeptidase |
| C4B63_228g51 | glutamamyl carboxypeptidase(fragment) |
| C4B63_228g52 | glutamamyl carboxypeptidase(fragment) |
| C4B63_22g743c | calpain-like cysteine peptidase |
| C4B63_23g16 | Battenin |
| C4B63_23g195 | 3', 5'-cyclic nucleotide phosphodiesterase |
| C4B63_23g226 | conserved hypothetical protein |
| C4B63_23g270 | conserved hypothetical protein |
| C4B63_259g12 | conserved hypothetical protein |
| C4B63_260g20 | trans-sialidase, Group II |
| C4B63_261g5 | conserved hypothetical protein |
| C4B63_26g240 | nucleobase transporter |
| C4B63_26g278 | arginine kinase |
| C4B63_26g282 | arginine kinase |
| C4B63_26g286 | arginine kinase |
| C4B63_277g1 | glutamamyl carboxypeptidase(fragment) |
| C4B63_277g10 | glutamamyl carboxypeptidase |
| C4B63_277g11 | glutamamyl carboxypeptidase |
| C4B63_277g12 | glutamamyl carboxypeptidase |
| C4B63_277g13 | glutamamyl carboxypeptidase |
| C4B63_277g14 | glutamamyl carboxypeptidase |
| C4B63_277g15 | glutamamyl carboxypeptidase |
| C4B63_277g16 | glutamamyl carboxypeptidase |
| C4B63_277g17 | glutamamyl carboxypeptidase |
| C4B63_277g18 | glutamamyl carboxypeptidase(fragment) |
| C4B63_277g2 | glutamamyl carboxypeptidase |
| C4B63_277g20 | glutamamyl carboxypeptidase(fragment) |
| C4B63_277g21 | glutamamyl carboxypeptidase |
| C4B63_277g3 | glutamamyl carboxypeptidase |
| C4B63_277g4 | glutamamyl carboxypeptidase |
| C4B63_277g5 | glutamamyl carboxypeptidase |
| C4B63_277g6 | glutamamyl carboxypeptidase |
| C4B63_277g7 | glutamamyl carboxypeptidase |
| C4B63_277g8 | glutamamyl carboxypeptidase |
| C4B63_277g9 | glutamamyl carboxypeptidase |
| C4B63_27g1015c | conserved hypothetical protein |
| C4B63_294g1 | glutamamyl carboxypeptidase |
| C4B63_294g10 | glutamamyl carboxypeptidase |
| C4B63_294g11 | glutamamyl carboxypeptidase |
| C4B63_294g12 | glutamamyl carboxypeptidase |
| C4B63_294g13 | glutamamyl carboxypeptidase |
| C4B63_294g2 | glutamamyl carboxypeptidase(fragment) |
| C4B63_294g5 | glutamamyl carboxypeptidase |
| C4B63_294g6 | glutamamyl carboxypeptidase |
| C4B63_294g7 | glutamamyl carboxypeptidase |
| C4B63_294g8 | glutamamyl carboxypeptidase |
| C4B63_294g9 | glutamamyl carboxypeptidase |
| C4B63_2g435 | Flagellum attachment zone protein 12 |
| C4B63_2g549 | receptor-type adenylate cyclase |
| C4B63_2g702 | Gim5A protein |
| C4B63_2g704 | Gim5A protein |
| C4B63_2g706 | Gim5A protein |
| C4B63_30g122 | surface protease GP63 |
| C4B63_30g150 | cystinosin |
| C4B63_30g182 | Putative adenylate cyclase regulatory protein |
| C4B63_30g210 | serine-threonine dehydratase |
| C4B63_30g261 | deoxyribose-phosphate aldolase |
| C4B63_31g119 | trans-sialidase, Group II |
| C4B63_31g93 | conserved hypothetical protein |
| C4B63_32g206 | fatty acid desaturase |
| C4B63_336g1 | glutamamyl carboxypeptidase(fragment) |
| C4B63_336g2 | glutamamyl carboxypeptidase(fragment) |
| C4B63_336g3 | metallo-peptidase, clan MH, family M18(fragment) |
| C4B63_336g4 | glutamamyl carboxypeptidase |
| C4B63_336g5 | glutamamyl carboxypeptidase |
| C4B63_336g6 | glutamamyl carboxypeptidase |
| C4B63_350g1 | glutamamyl carboxypeptidase |
| C4B63_350g2 | glutamamyl carboxypeptidase(fragment) |
| C4B63_350g3 | glutamamyl carboxypeptidase(fragment) |
| C4B63_350g4 | metallo-peptidase, clan MH, family M18 |
| C4B63_350g5 | glutamamyl carboxypeptidase(fragment) |
| C4B63_351g27c | small myristoylated protein-1 |
| C4B63_351g3 | calpain-like cysteine peptidase |
| C4B63_37g382 | unspecified product |
| C4B63_37g383 | unspecified product |
| C4B63_38g174 | conserved hypothetical protein |
| C4B63_38g83 | conserved hypothetical protein |
| C4B63_38g94 | conserved hypothetical protein |
| C4B63_39g346 | nucleic acid binding protein |
| C4B63_3g644 | mucin TcMUCII |
| C4B63_40g186 | Gamma carbonic anhydrase 3, mitochondrial |
| C4B63_41g1149c | conserved hypothetical protein |
| C4B63_431g1 | glutamamyl carboxypeptidase(fragment) |
| C4B63_431g10 | glutamamyl carboxypeptidase |
| C4B63_431g3 | glutamamyl carboxypeptidase |
| C4B63_431g4 | glutamamyl carboxypeptidase |
| C4B63_431g5 | glutamamyl carboxypeptidase |
| C4B63_431g6 | glutamamyl carboxypeptidase |
| C4B63_431g7 | glutamamyl carboxypeptidase |
| C4B63_431g8 | glutamamyl carboxypeptidase |
| C4B63_431g9 | glutamamyl carboxypeptidase |
| C4B63_43g166 | conserved hypothetical protein |
| C4B63_44g228 | amino acid permease |
| C4B63_462g10 | glutamamyl carboxypeptidase |
| C4B63_462g11 | glutamamyl carboxypeptidase |
| C4B63_462g12 | glutamamyl carboxypeptidase |
| C4B63_462g13 | glutamamyl carboxypeptidase |
| C4B63_46g90 | Ring finger domain containing protein |
| C4B63_47g75 | splicing factor ptsr1 interacting protein |
| C4B63_485g5 | glutamamyl carboxypeptidase(fragment) |
| C4B63_4g189 | alcohol dehydrogenase |
| C4B63_54g61 | conserved hypothetical protein |
| C4B63_550g1 | metallo-peptidase, clan MH, family M18 |
| C4B63_550g2 | glutamamyl carboxypeptidase |
| C4B63_550g3 | metallo-peptidase, clan MH, family M18 |
| C4B63_550g6 | glutamamyl carboxypeptidase |
| C4B63_55g177 | conserved hypothetical protein |
| C4B63_58g70 | hexokinase |
| C4B63_5g44 | trans-sialidase, Group V |
| C4B63_63g102 | hexose transporter |
| C4B63_63g104 | hexose transporter |
| C4B63_63g106 | hexose transporter |
| C4B63_63g108 | hexose transporter |
| C4B63_63g110 | hexose transporter |
| C4B63_67g26 | trans-sialidase, Group II |
| C4B63_6g2366c | UPF0382 membrane protein HI 1073 |
| C4B63_7g143 | Tubulin polymerization-promoting protein family member 2 |
| C4B63_7g145 | conserved hypothetical protein |
| C4B63_7g361 | phosphomannomutase-like protein |
| C4B63_7g98 | Eukaryotic translation initiation factor 2-alpha kinase 2 |
| C4B63_84g160c | paraflagellar rod component |
| C4B63_87g8 | glutamamyl carboxypeptidase |
| C4B63_93g25 | unspecified product |
| C4B63_98g22 | trans-sialidase |
| C4B63_98g34 | trans-sialidase |

##### Upregulated

| ID | Annotation |
|---|---|
| C4B63_101g36 | inositol-3-phosphate synthase |
| C4B63_102g44 | hypoxanthine-guanine phosphoribosyltransferase |
| C4B63_104g78 | calreticulin |
| C4B63_10g439 | heat shock protein 90 |
| C4B63_10g449 | Triple RNA binding domain protein 3 |
| C4B63_10g517 | tryptophanyl-tRNA synthetase |
| C4B63_113g22 | heat shock protein 85(fragment) |
| C4B63_113g23 | heat shock protein 85(fragment) |
| C4B63_113g24 | heat shock protein 85(fragment) |
| C4B63_113g25 | heat shock protein 85 |
| C4B63_113g26 | heat shock protein 85 |
| C4B63_113g27 | heat shock protein 85(fragment) |
| C4B63_113g28 | heat shock protein 85 |
| C4B63_113g29 | heat shock protein 85 |
| C4B63_113g30 | heat shock protein 85 |
| C4B63_113g31 | heat shock protein 85(fragment) |
| C4B63_113g32 | heat shock protein 85 |
| C4B63_113g33 | heat shock protein 85 |
| C4B63_11g29 | 2,4-dihydroxyhept-2-ene-1,7-dioic acid aldolase |
| C4B63_11g40 | urocanate hydratase |
| C4B63_11g8 | succinyl-CoA synthetase alpha subunit |
| C4B63_11g9 | eukaryotic translation initiation factor 3 subunit j |
| C4B63_123g35 | Uncharacterized protein At3g06530 |
| C4B63_123g36 | O-6 methyl-guanine alkyl transferase |
| C4B63_127g35 | peptidase |
| C4B63_12g153 | metallo-peptidase, Clan MF, Family M17 |
| C4B63_12g195 | conserved hypothetical protein |
| C4B63_12g237 | eukaryotic release factor 3 |
| C4B63_12g295 | GTP-binding nuclear protein rtb2 |
| C4B63_133g56 | heat shock protein 70 (hsp70) |
| C4B63_133g57 | heat shock protein 70 (hsp70) |
| C4B63_133g58 | heat shock protein 70 (hsp70) |
| C4B63_133g59 | heat shock protein 70 (hsp70) |
| C4B63_133g60 | heat shock protein 70 (hsp70) |
| C4B63_133g61 | heat shock protein 70 (hsp70) |
| C4B63_133g62 | heat shock protein 70 (hsp70)(fragment) |
| C4B63_133g63 | heat shock protein 70 (hsp70) |
| C4B63_137g25 | conserved hypothetical protein |
| C4B63_137g32 | conserved hypothetical protein |
| C4B63_137g34 | heat shock protein 70 (hsp70)(fragment) |
| C4B63_137g36 | heat shock protein 70 (hsp70)(fragment) |
| C4B63_137g37 | heat shock protein 70 (hsp70)(fragment) |
| C4B63_137g38 | heat shock protein 70 (hsp70)(fragment) |
| C4B63_137g39 | heat shock protein 70 (hsp70) |
| C4B63_137g40 | heat shock protein 70 (hsp70)(fragment) |
| C4B63_137g41 | heat shock protein 70 (hsp70) |
| C4B63_138g4 | elongation factor 1-gamma (EF-1-gamma) |
| C4B63_13g130 | ER membrane protein complex subunit 3 |
| C4B63_13g145 | eukaryotic translation initiation factor 3 subunit l |
| C4B63_143g20 | glutamine synthetase |
| C4B63_143g49 | cystathionine beta-synthase |
| C4B63_148g35 | zinc-finger protein 2 |
| C4B63_149g20 | cyclophilin |
| C4B63_14g165 | N-lysine methyltransferase SETD6 |
| C4B63_14g204 | 60S acidic ribosomal protein |
| C4B63_14g220c | kinetoplast DNA-associated protein |
| C4B63_157g28 | FK506-binding protein (FKBP)-type peptidyl-prolyl isomerase |
| C4B63_16g326 | conserved hypothetical protein |
| C4B63_171g30 | FK506-binding protein (FKBP)-type peptidyl-prolyl isomerase |
| C4B63_175g9 | heat shock protein DNAJ |
| C4B63_17g101 | mitochondrial carrier protein |
| C4B63_17g179 | Importin subunit beta-3 |
| C4B63_17g65 | polypyrimidine tract-binding protein |
| C4B63_180g3 | conserved hypothetical protein |
| C4B63_183g12 | mitochondrial carrier protein |
| C4B63_184g39 | heat shock protein DNAJ(fragment) |
| C4B63_18g164 | S-adenosylmethionine synthetase |
| C4B63_18g204 | 3-hydroxy-3-methylglutaryl-CoA reductase |
| C4B63_18g212 | valyl-tRNA synthetase |
| C4B63_18g260 | small glutamine-rich tetratricopeptide repeat protein |
| C4B63_18g287 | heat shock 70 kDa protein, mitochondrial precursor |
| C4B63_18g288 | heat shock 70 kDa protein, mitochondrial precursor |
| C4B63_18g293 | heat shock 70 kDa protein, mitochondrial precursor |
| C4B63_192g3 | AAA ATPase |
| C4B63_193g9 | metallo-peptidase, Clan MF, Family M17 |
| C4B63_196g51 | CDP-diacylglycerol--inositol 3-phosphatidyltransferase |
| C4B63_207g7 | conserved hypothetical protein |
| C4B63_207g9 | ribosome biogenesis protein |
| C4B63_20g172 | chaperonin alpha subunit |
| C4B63_20g213 | enoyl-CoA hydratase/isomerase family protein |
| C4B63_210g35 | 3,2-trans-enoyl-CoA isomerase, mitochondrial precursor |
| C4B63_212g26 | conserved hypothetical protein |
| C4B63_214g14 | UDP-Gal or UDP-GlcNAc-dependent glycosyltransferase |
| C4B63_220g13 | cyclophilin |
| C4B63_22g25 | Major Facilitator Superfamily |
| C4B63_23g109 | UDP-glucose:glycoprotein glucosyltransferase |
| C4B63_23g110 | UDP-glucose:glycoprotein glucosyltransferase |
| C4B63_23g248 | inorganic pyrophosphatase |
| C4B63_23g275 | ATP-dependent DEAD/H RNA helicase |
| C4B63_24g172 | Uncharacterized protein At3g06530 |
| C4B63_24g298 | CDP-diacylglycerol--inositol 3-phosphatidyltransferase |
| C4B63_24g325 | conserved hypothetical protein |
| C4B63_252g22 | T-complex protein 1, beta subunit |
| C4B63_255g21 | calreticulin |
| C4B63_25g192 | NAA35 |
| C4B63_25g228 | Putative uridine kinase C227.14 |
| C4B63_25g249 | Protein SDA1 homolog |
| C4B63_25g279 | J Biosynthesis Protein 2 |
| C4B63_25g294 | Thioredoxin-like |
| C4B63_261g22c | conserved hypothetical protein |
| C4B63_26g204 | phosphomevalonate kinase protein |
| C4B63_26g205 | proliferative cell nuclear antigen (PCNA) |
| C4B63_26g252 | nucleosome assembly protein-like protein |
| C4B63_26g279 | ABC transporter |
| C4B63_26g283 | ABC transporter |
| C4B63_26g287 | ABC transporter |
| C4B63_276g16 | conserved hypothetical protein |
| C4B63_28g106 | malic enzyme |
| C4B63_297g10 | polypyrimidine tract-binding protein |
| C4B63_297g14 | phenylalanyl-tRNA synthetase alpha chain |
| C4B63_2g174 | polyadenylate-binding protein 1 |
| C4B63_2g223 | conserved hypothetical protein |
| C4B63_2g235 | Co-chaperone protein P23 |
| C4B63_2g305 | Hsp70-Hsp90 organizing protein 3 |
| C4B63_2g311 | conserved hypothetical protein |
| C4B63_2g430 | heat shock protein |
| C4B63_2g485 | T-complex protein 1, beta subunit |
| C4B63_2g509 | arginyl-tRNA synthetase |
| C4B63_2g510 | arginyl-tRNA synthetase |
| C4B63_2g525 | ribosome biogenesis protein MAK21 |
| C4B63_2g690 | chaperonin alpha subunit |
| C4B63_2g90 | Nop53 (60S ribosomal biogenesis) |
| C4B63_300g102c | mucin TcSMUGL |
| C4B63_304g12 | conserved hypothetical protein |
| C4B63_304g13 | nucleosome assembly protein |
| C4B63_30g161 | glutamine synthetase |
| C4B63_30g290 | mercaptopyruvate sulfurtransferase |
| C4B63_31g105 | signal peptidase type I |
| C4B63_32g205 | notchless homolog |
| C4B63_32g217 | 2-amino-3-ketobutyrate coenzyme A ligase |
| C4B63_32g22 | Uncharacterized WD repeat-containing protein C17D11.16 |
| C4B63_34g107 | Nop14-like family |
| C4B63_34g1460c | mucin TcSMUGL |
| C4B63_34g287 | glycerate kinase |
| C4B63_34g288 | glycerate kinase |
| C4B63_34g289 | glycerate kinase |
| C4B63_34g290 | glycerate kinase |
| C4B63_35g189 | eukaryotic translation initiation factor 1A |
| C4B63_360g21 | unspecified product |
| C4B63_38g144 | mitochondrial carrier protein |
| C4B63_38g320c | conserved hypothetical protein |
| C4B63_39g333 | eukaryotic translation initiation factor 3 subunit 2 |
| C4B63_3g1120 | conserved hypothetical protein |
| C4B63_41g251 | arginine N-methyltransferase |
| C4B63_42g103 | pyridoxal kinase |
| C4B63_42g178 | histidyl-tRNA synthetase |
| C4B63_42g220 | transmembrane/endomembrane-like protein |
| C4B63_42g251 | conserved hypothetical protein |
| C4B63_42g63 | adenosine kinase |
| C4B63_42g70 | p22 protein precursor |
| C4B63_43g125 | centromere/microtubule binding protein cbf5 |
| C4B63_43g161 | ATP-dependent RNA helicase SUB2 |
| C4B63_44g215 | tRNA:m(4)X modification enzyme TRM13 |
| C4B63_44g246 | nucleolar protein |
| C4B63_45g126 | RNA recognition motif (a.k.a. RRM, RBD, or RNP domain) |
| C4B63_45g239 | lysyl-tRNA synthetase |
| C4B63_46g11 | chaperonin TCP20 |
| C4B63_46g91 | leucyl-tRNA synthetase |
| C4B63_47g52 | deoxyhypusine synthase |
| C4B63_47g53 | deoxyhypusine synthase |
| C4B63_48g132 | universal minicircle sequence binding protein |
| C4B63_48g168 | AAA ATPase |
| C4B63_48g25 | conserved hypothetical protein |
| C4B63_48g85 | chaperonin HSP60, mitochondrial precursor |
| C4B63_48g86 | chaperonin HSP60, mitochondrial precursor |
| C4B63_48g87 | chaperonin HSP60, mitochondrial precursor |
| C4B63_48g88 | chaperonin HSP60, mitochondrial precursor |
| C4B63_48g89 | chaperonin HSP60, mitochondrial precursor |
| C4B63_48g90 | chaperonin HSP60, mitochondrial precursor |
| C4B63_48g91 | chaperonin HSP60, mitochondrial precursor |
| C4B63_48g92 | chaperonin HSP60, mitochondrial precursor |
| C4B63_4g131 | mitochondrial carrier protein |
| C4B63_4g200 | conserved hypothetical protein |
| C4B63_4g201 | Protein ILITYHIA |
| C4B63_4g267 | mitochondrial carrier protein |
| C4B63_4g357 | Activator of 90 kDa heat shock protein ATPase homolog 1 |
| C4B63_4g453 | conserved hypothetical protein |
| C4B63_4g454 | nucleosome assembly protein |
| C4B63_4g64 | Mitochondrial ADP/ATP carrier protein 5 |
| C4B63_4g68 | metallo-peptidase, Clan MG, Family M24 |
| C4B63_4g77 | conserved hypothetical protein |
| C4B63_4g79 | ribosome biogenesis protein |
| C4B63_55g62 | Ribosomal RNA small subunit methyltransferase NEP1 |
| C4B63_57g88 | fibrillarin |
| C4B63_58g135 | heat shock protein DNAJ |
| C4B63_58g143 | Lupus La protein homolog |
| C4B63_59g115 | stress-inducible protein STI1-like(fragment) |
| C4B63_60g172 | conserved hypothetical protein |
| C4B63_63g12 | protein disulfide isomerase 2 |
| C4B63_63g15 | ribosomal protein L1-like protein |
| C4B63_63g16 | chaperonin alpha subunit |
| C4B63_63g35 | histidine ammonia-lyase |
| C4B63_63g58 | Pre-rRNA-processing protein esf1 |
| C4B63_69g26 | endo-beta-N-acetylglucosaminidase |
| C4B63_6g118 | Mitochondrial import receptor subunit ATOM46 |
| C4B63_6g119 | oxidoreductase-like protein |
| C4B63_6g129 | heat shock protein 70 (hsp70) |
| C4B63_6g130 | heat shock protein 70 (hsp70)(fragment) |
| C4B63_6g132 | heat shock protein 70 (hsp70) |
| C4B63_6g136 | heat shock protein 70 (hsp70) |
| C4B63_6g238 | cation transporter |
| C4B63_6g239 | cation transporter |
| C4B63_6g240 | cation transporter |
| C4B63_6g241 | cation transporter |
| C4B63_6g242 | cation transporter |
| C4B63_6g243 | cation transporter |
| C4B63_6g245 | cation transporter |
| C4B63_6g246 | cation transporter |
| C4B63_6g247 | cation transporter |
| C4B63_6g248 | cation transporter |
| C4B63_6g297 | conserved hypothetical protein |
| C4B63_6g305 | heat shock protein 70 (hsp70) |
| C4B63_6g306 | heat shock protein 70 (hsp70)(fragment) |
| C4B63_6g307 | heat shock protein 70 (hsp70)(fragment) |
| C4B63_6g308 | heat shock protein 70 (hsp70) |
| C4B63_6g309 | heat shock protein 70 (hsp70) |
| C4B63_6g310 | heat shock protein 70 (hsp70)(fragment) |
| C4B63_6g312 | heat shock protein 70 (hsp70) |
| C4B63_6g313 | heat shock protein 70 (hsp70)(fragment) |
| C4B63_6g314 | heat shock protein 70 (hsp70) |
| C4B63_6g315 | heat shock protein 70 (hsp70) |
| C4B63_6g316 | heat shock protein 70 (hsp70) |
| C4B63_6g317 | heat shock protein 70 (hsp70) |
| C4B63_6g403 | palmitoyl acyltransferase 4 |
| C4B63_6g404 | Ribosome biogenesis protein bms1 |
| C4B63_6g405 | Ribosome biogenesis protein BMS1 homolog |
| C4B63_6g489 | Something about silencing protein 10 |
| C4B63_6g495 | Ankyrin repeat-containing protein P16F5.05c |
| C4B63_6g516 | WD repeat-containing protein 75 |
| C4B63_6g550 | NAD(P)-dependent steroid dehydrogenase protein |
| C4B63_6g556 | NAD(P)-dependent steroid dehydrogenase protein |
| C4B63_70g129 | metallo-peptidase, Clan MF, Family M17 |
| C4B63_70g61 | RNA recognition motif. (a.k.a. RRM, RBD, or RNP domain) |
| C4B63_70g78 | PA domain containing protein |
| C4B63_72g114 | conserved hypothetical protein |
| C4B63_78g20 | nucleolar RNA-binding protein |
| C4B63_78g23 | nucleolar RNA-binding protein |
| C4B63_78g25 | nucleolar RNA-binding protein |
| C4B63_79g54 | hypoxanthine-guanine phosphoribosyltransferase |
| C4B63_7g210 | Ribonuclease P protein subunit p25-like protein |
| C4B63_7g212 | ruvB-like DNA helicase |
| C4B63_7g223 | eukaryotic translation initiation factor 3 subunit g |
| C4B63_7g400 | ER membrane protein complex subunit 1 |
| C4B63_81g100 | chaperonin HSP60, mitochondrial precursor |
| C4B63_81g101 | chaperonin HSP60, mitochondrial precursor |
| C4B63_81g102 | chaperonin HSP60, mitochondrial precursor |
| C4B63_81g103 | chaperonin HSP60, mitochondrial precursor |
| C4B63_81g104 | chaperonin HSP60, mitochondrial precursor |
| C4B63_81g105 | chaperonin HSP60, mitochondrial precursor |
| C4B63_81g106 | chaperonin HSP60, mitochondrial precursor |
| C4B63_81g40 | ATP-dependent DEAD/H RNA helicase |
| C4B63_83g36 | ppnN |
| C4B63_83g73 | ribonucleoprotein p18, mitochondrial precursor |
| C4B63_83g80 | cytoplasmic l-asparaginase i-like protein |
| C4B63_84g21 | conserved hypothetical protein |
| C4B63_84g87 | heat shock protein 85 |
| C4B63_84g88 | heat shock protein 85 |
| C4B63_84g89 | heat shock protein 85 |
| C4B63_84g90 | heat shock protein 85(fragment) |
| C4B63_8g2656c | 10 kDa heat shock protein |
| C4B63_8g467 | NMD3 family |
| C4B63_8g559 | Adenine phosphoribosyltransferase |
| C4B63_93g64 | amidohydrolase |
| C4B63_94g51 | Mucin-1 |
| C4B63_97g18 | heat shock protein 110 |
| C4B63_9g246 | FtsJ cell division protein |
| C4B63_9g263 | trypanothione synthetase |
| C4B63_9g396 | sedoheptulose-1,7-bisphosphatase |
| C4B63_9g453 | conserved hypothetical protein |
| C4B63_9g484 | 5-methylcytosine rRNA methyltransferase NSUN4 |

#### Hemin, Hb and No heme common DEGs (t24 vs. t0)

##### Downregulated

| ID | Annotation |
|---|---|
| C4B63_101g12 | sterol 24-c-methyltransferase |
| C4B63_109g41 | Trypomastigote, Alanine, Serine and Valine rich protein (TASV), subfamily C |
| C4B63_109g61 | trans-sialidase, Group II |
| C4B63_112g40 | surface protease GP63 |
| C4B63_112g66 | Gamma carbonic anhydrase 2, mitochondrial |
| C4B63_113g41 | thiol-dependent reductase 1 |
| C4B63_113g42 | thiol-dependent reductase 1 |
| C4B63_129g44 | cytochrome c oxidase VIII (COX VIII) |
| C4B63_12g242 | SKP1-like protein |
| C4B63_130g39 | trans-sialidase, Group II |
| C4B63_137g16 | cytochrome c oxidase assembly protein |
| C4B63_140g32 | cytochrome c oxidase VIII (COX VIII) |
| C4B63_14g1 | hexose transporter |
| C4B63_14g2 | hexose transporter(fragment) |
| C4B63_155g10 | trans-sialidase |
| C4B63_156g18 | trans-sialidase |
| C4B63_156g6 | trans-sialidase |
| C4B63_163g33 | protein disulfide isomerase |
| C4B63_16g136 | conserved hypothetical protein |
| C4B63_170g37 | conserved hypothetical protein |
| C4B63_170g5 | conserved hypothetical protein |
| C4B63_17g48 | ubiquitin-conjugating enzyme E2 |
| C4B63_18g271 | conserved hypothetical protein |
| C4B63_196g32 | Mucin-associated surface protein (MASP) |
| C4B63_19g86 | CYC2-like cyclin 6 |
| C4B63_1g94 | trans-sialidase, Group II |
| C4B63_1g953 | trans-sialidase, Group V |
| C4B63_21g10 | arginine permease |
| C4B63_21g13 | arginine permease |
| C4B63_21g14 | arginine permease |
| C4B63_21g20 | arginine permease |
| C4B63_21g21 | arginine permease |
| C4B63_21g25 | arginine permease |
| C4B63_21g33 | small GTP-binding protein Rab7 |
| C4B63_21g7 | arginine permease |
| C4B63_21g9 | arginine permease |
| C4B63_228g46 | glutamamyl carboxypeptidase |
| C4B63_228g47 | glutamamyl carboxypeptidase |
| C4B63_228g48 | glutamamyl carboxypeptidase |
| C4B63_228g49 | glutamamyl carboxypeptidase |
| C4B63_228g50 | glutamamyl carboxypeptidase |
| C4B63_228g51 | glutamamyl carboxypeptidase(fragment) |
| C4B63_228g52 | glutamamyl carboxypeptidase(fragment) |
| C4B63_22g167 | calpain-like cysteine peptidase |
| C4B63_22g743c | calpain-like cysteine peptidase |
| C4B63_232g21 | trans-sialidase, Group I |
| C4B63_23g17 | Battenin |
| C4B63_244g9 | trans-sialidase, Group I |
| C4B63_248g14 | glucose-6-phosphate isomerase, glycosomal |
| C4B63_24g165 | protein kinase |
| C4B63_24g199 | conserved hypothetical protein |
| C4B63_259g12 | conserved hypothetical protein |
| C4B63_25g29 | trans-sialidase, Group I |
| C4B63_25g331 | trans-sialidase, Group I |
| C4B63_260g20 | trans-sialidase, Group II |
| C4B63_26g311 | ABC transporter(fragment) |
| C4B63_277g1 | glutamamyl carboxypeptidase(fragment) |
| C4B63_277g10 | glutamamyl carboxypeptidase |
| C4B63_277g12 | glutamamyl carboxypeptidase |
| C4B63_277g13 | glutamamyl carboxypeptidase |
| C4B63_277g14 | glutamamyl carboxypeptidase |
| C4B63_277g15 | glutamamyl carboxypeptidase |
| C4B63_277g16 | glutamamyl carboxypeptidase |
| C4B63_277g17 | glutamamyl carboxypeptidase |
| C4B63_277g18 | glutamamyl carboxypeptidase(fragment) |
| C4B63_277g2 | glutamamyl carboxypeptidase |
| C4B63_277g20 | glutamamyl carboxypeptidase(fragment) |
| C4B63_277g21 | glutamamyl carboxypeptidase |
| C4B63_277g4 | glutamamyl carboxypeptidase |
| C4B63_277g5 | glutamamyl carboxypeptidase |
| C4B63_277g6 | glutamamyl carboxypeptidase |
| C4B63_277g8 | glutamamyl carboxypeptidase |
| C4B63_277g9 | glutamamyl carboxypeptidase |
| C4B63_27g185 | conserved hypothetical protein |
| C4B63_294g1 | glutamamyl carboxypeptidase |
| C4B63_294g10 | glutamamyl carboxypeptidase |
| C4B63_294g11 | glutamamyl carboxypeptidase |
| C4B63_294g12 | glutamamyl carboxypeptidase |
| C4B63_294g13 | glutamamyl carboxypeptidase |
| C4B63_294g2 | glutamamyl carboxypeptidase(fragment) |
| C4B63_294g5 | glutamamyl carboxypeptidase |
| C4B63_294g6 | glutamamyl carboxypeptidase |
| C4B63_294g7 | glutamamyl carboxypeptidase |
| C4B63_294g8 | glutamamyl carboxypeptidase |
| C4B63_294g9 | glutamamyl carboxypeptidase |
| C4B63_29g223 | trans-sialidase, Group II |
| C4B63_2g547 | receptor-type adenylate cyclase |
| C4B63_2g548 | receptor-type adenylate cyclase |
| C4B63_2g549 | receptor-type adenylate cyclase |
| C4B63_30g122 | surface protease GP63 |
| C4B63_30g150 | cystinosin |
| C4B63_30g275 | protein disulfide isomerase |
| C4B63_312g7 | conserved hypothetical protein |
| C4B63_31g119 | trans-sialidase, Group II |
| C4B63_322g5 | glutamamyl carboxypeptidase |
| C4B63_336g2 | glutamamyl carboxypeptidase(fragment) |
| C4B63_336g3 | metallo-peptidase, clan MH, family M18(fragment) |
| C4B63_336g4 | glutamamyl carboxypeptidase |
| C4B63_336g5 | glutamamyl carboxypeptidase |
| C4B63_336g6 | glutamamyl carboxypeptidase |
| C4B63_33g163 | unspecified product |
| C4B63_33g164 | unspecified product |
| C4B63_33g167 | trans-sialidase, Group II |
| C4B63_34g117 | cytochrome b5-dependent oleate desaturase |
| C4B63_34g140 | trans-sialidase, Group IV |
| C4B63_350g1 | glutamamyl carboxypeptidase |
| C4B63_350g2 | glutamamyl carboxypeptidase(fragment) |
| C4B63_350g4 | metallo-peptidase, clan MH, family M18 |
| C4B63_350g5 | glutamamyl carboxypeptidase(fragment) |
| C4B63_37g382 | unspecified product |
| C4B63_37g383 | unspecified product |
| C4B63_38g178 | UDP-Gal or UDP-GlcNAc-dependent glycosyltransferase |
| C4B63_38g180 | UDP-Gal or UDP-GlcNAc-dependent glycosyltransferase |
| C4B63_38g184 | UDP-Gal or UDP-GlcNAc-dependent glycosyltransferase |
| C4B63_39g346 | nucleic acid binding protein |
| C4B63_39g66 | trans-sialidase, Group V |
| C4B63_40g30 | conserved hypothetical protein |
| C4B63_431g1 | glutamamyl carboxypeptidase(fragment) |
| C4B63_431g3 | glutamamyl carboxypeptidase |
| C4B63_431g4 | glutamamyl carboxypeptidase |
| C4B63_431g5 | glutamamyl carboxypeptidase |
| C4B63_431g6 | glutamamyl carboxypeptidase |
| C4B63_431g7 | glutamamyl carboxypeptidase |
| C4B63_431g8 | glutamamyl carboxypeptidase |
| C4B63_431g9 | glutamamyl carboxypeptidase |
| C4B63_43g225 | conserved hypothetical protein |
| C4B63_43g83 | trans-sialidase, Group II |
| C4B63_44g106 | cytochrome c oxidase VIII (COX VIII) |
| C4B63_44g228 | amino acid permease |
| C4B63_459g7 | UDP-Gal or UDP-GlcNAc-dependent glycosyltransferase |
| C4B63_459g9 | UDP-Gal or UDP-GlcNAc-dependent glycosyltransferase |
| C4B63_462g10 | glutamamyl carboxypeptidase |
| C4B63_462g11 | glutamamyl carboxypeptidase |
| C4B63_462g12 | glutamamyl carboxypeptidase |
| C4B63_462g13 | glutamamyl carboxypeptidase |
| C4B63_46g58 | conserved hypothetical protein |
| C4B63_46g62 | conserved hypothetical protein |
| C4B63_47g128 | trans-sialidase |
| C4B63_47g75 | splicing factor ptsr1 interacting protein |
| C4B63_485g5 | glutamamyl carboxypeptidase(fragment) |
| C4B63_4g496 | transporter |
| C4B63_550g1 | metallo-peptidase, clan MH, family M18 |
| C4B63_550g3 | metallo-peptidase, clan MH, family M18 |
| C4B63_550g6 | glutamamyl carboxypeptidase |
| C4B63_5g44 | trans-sialidase, Group V |
| C4B63_63g102 | hexose transporter |
| C4B63_63g104 | hexose transporter |
| C4B63_63g106 | hexose transporter |
| C4B63_63g108 | hexose transporter |
| C4B63_63g110 | hexose transporter |
| C4B63_68g134 | trans-sialidase, Group II |
| C4B63_7g143 | Tubulin polymerization-promoting protein family member 2 |
| C4B63_87g8 | glutamamyl carboxypeptidase |
| C4B63_93g25 | unspecified product |
| C4B63_98g22 | trans-sialidase |
| C4B63_98g34 | trans-sialidase |
| C4B63_9g382 | conserved hypothetical protein |

##### Upregulated

| ID | Annotation |
|---|---|
| C4B63_101g36 | inositol-3-phosphate synthase |
| C4B63_10g406 | High mobility group protein TDP1 |
| C4B63_10g492 | conserved hypothetical protein |
| C4B63_10g517 | tryptophanyl-tRNA synthetase |
| C4B63_112g84c | Histone H2B variant V |
| C4B63_113g22 | heat shock protein 85(fragment) |
| C4B63_113g23 | heat shock protein 85(fragment) |
| C4B63_113g24 | heat shock protein 85(fragment) |
| C4B63_113g25 | heat shock protein 85 |
| C4B63_113g26 | heat shock protein 85 |
| C4B63_113g28 | heat shock protein 85 |
| C4B63_113g29 | heat shock protein 85 |
| C4B63_113g30 | heat shock protein 85 |
| C4B63_113g31 | heat shock protein 85(fragment) |
| C4B63_113g32 | heat shock protein 85 |
| C4B63_113g33 | heat shock protein 85 |
| C4B63_11g29 | 2,4-dihydroxyhept-2-ene-1,7-dioic acid aldolase |
| C4B63_11g36 | kinesin |
| C4B63_11g37 | kinesin |
| C4B63_11g8 | succinyl-CoA synthetase alpha subunit |
| C4B63_127g35 | peptidase |
| C4B63_127g54 | squalene synthase |
| C4B63_128g301c | histone H4 |
| C4B63_128g305c | histone H4 |
| C4B63_128g308c | histone H4 |
| C4B63_128g316c | histone H4 |
| C4B63_12g153 | metallo-peptidase, Clan MF, Family M17 |
| C4B63_12g237 | eukaryotic release factor 3 |
| C4B63_12g295 | GTP-binding nuclear protein rtb2 |
| C4B63_133g56 | heat shock protein 70 (hsp70) |
| C4B63_133g57 | heat shock protein 70 (hsp70) |
| C4B63_133g58 | heat shock protein 70 (hsp70) |
| C4B63_133g59 | heat shock protein 70 (hsp70) |
| C4B63_133g61 | heat shock protein 70 (hsp70) |
| C4B63_133g62 | heat shock protein 70 (hsp70)(fragment) |
| C4B63_133g63 | heat shock protein 70 (hsp70) |
| C4B63_137g34 | heat shock protein 70 (hsp70)(fragment) |
| C4B63_137g37 | heat shock protein 70 (hsp70)(fragment) |
| C4B63_137g41 | heat shock protein 70 (hsp70) |
| C4B63_13g145 | eukaryotic translation initiation factor 3 subunit l |
| C4B63_13g263 | GPI alpha-mannosyltransferase III |
| C4B63_13g34 | 21 kDa cyclophilin |
| C4B63_142g5 | conserved hypothetical protein |
| C4B63_147g41 | conserved hypothetical protein |
| C4B63_149g20 | cyclophilin |
| C4B63_14g165 | N-lysine methyltransferase SETD6 |
| C4B63_14g220c | kinetoplast DNA-associated protein |
| C4B63_14g241 | trypanothione reductase |
| C4B63_14g249 | structural maintenance of chromosome 3 |
| C4B63_14g62 | kinetoplast DNA-associated protein |
| C4B63_14g96 | isoleucil-tRNA synthetase |
| C4B63_14g97 | isoleucyl-tRNA synthetase |
| C4B63_157g28 | FK506-binding protein (FKBP)-type peptidyl-prolyl isomerase |
| C4B63_15g14 | oligosaccharyl transferase subunit |
| C4B63_160g6 | poly(A)-specific ribonuclease PARN |
| C4B63_161g18 | UDP-Gal or UDP-GlcNAc-dependent glycosyltransferase |
| C4B63_162g33 | chaperonin TCP20 |
| C4B63_168g33 | kinesin |
| C4B63_16g315 | mitochondrial DNA primase |
| C4B63_16g326 | conserved hypothetical protein |
| C4B63_171g30 | FK506-binding protein (FKBP)-type peptidyl-prolyl isomerase |
| C4B63_175g9 | heat shock protein DNAJ |
| C4B63_17g102 | conserved hypothetical protein |
| C4B63_17g179 | Importin subunit beta-3 |
| C4B63_17g268 | PSP1 C-terminal conserved region |
| C4B63_17g90 | T-complex protein 1, beta subunit |
| C4B63_180g2 | conserved hypothetical protein |
| C4B63_180g3 | conserved hypothetical protein |
| C4B63_180g4 | conserved hypothetical protein |
| C4B63_184g15 | conserved hypothetical protein |
| C4B63_188g129c | histone H4 |
| C4B63_188g130c | histone H4 |
| C4B63_188g132c | histone H4 |
| C4B63_188g135c | histone H4 |
| C4B63_188g137c | histone H4 |
| C4B63_188g138c | histone H4 |
| C4B63_188g140c | histone H4 |
| C4B63_188g141c | histone H4 |
| C4B63_188g144c | histone H4 |
| C4B63_188g145c | histone H4 |
| C4B63_188g146c | histone H4 |
| C4B63_188g147c | histone H4 |
| C4B63_188g148c | histone H4 |
| C4B63_188g149c | histone H4 |
| C4B63_188g152c | histone H4 |
| C4B63_188g153c | histone H4 |
| C4B63_188g154c | histone H4 |
| C4B63_188g157c | histone H4 |
| C4B63_188g159c | histone H4 |
| C4B63_188g160c | histone H4 |
| C4B63_188g161c | histone H4 |
| C4B63_188g162c | histone H4 |
| C4B63_188g163c | histone H4 |
| C4B63_188g164c | histone H4 |
| C4B63_18g164 | S-adenosylmethionine synthetase |
| C4B63_18g19 | ESAG8-associated protein |
| C4B63_18g204 | 3-hydroxy-3-methylglutaryl-CoA reductase |
| C4B63_18g212 | valyl-tRNA synthetase |
| C4B63_18g229 | conserved hypothetical protein |
| C4B63_18g260 | small glutamine-rich tetratricopeptide repeat protein |
| C4B63_18g287 | heat shock 70 kDa protein, mitochondrial precursor |
| C4B63_18g292 | conserved hypothetical protein |
| C4B63_18g293 | heat shock 70 kDa protein, mitochondrial precursor |
| C4B63_192g3 | AAA ATPase |
| C4B63_193g9 | metallo-peptidase, Clan MF, Family M17 |
| C4B63_194g18 | Histone-lysine N-methyltransferase, H3 lysine-76 specific |
| C4B63_196g51 | CDP-diacylglycerol--inositol 3-phosphatidyltransferase |
| C4B63_19g195 | Nuclear pore complex protein 158 |
| C4B63_207g13 | prostaglandin F synthase |
| C4B63_207g15 | prostaglandin F synthase |
| C4B63_208g38 | kinetoplastid kinetochore protein 8 |
| C4B63_20g213 | enoyl-CoA hydratase/isomerase family protein |
| C4B63_20g219 | kinetoplast-associated protein 3 |
| C4B63_214g13 | UDP-Gal or UDP-GlcNAc-dependent glycosyltransferase |
| C4B63_214g14 | UDP-Gal or UDP-GlcNAc-dependent glycosyltransferase |
| C4B63_21g238 | conserved hypothetical protein |
| C4B63_227g2 | glucan 1,3-alpha-glucosidase |
| C4B63_22g175 | pumilio/PUF RNA binding protein 9 |
| C4B63_22g22 | ARM-like helical domain-containing protein |
| C4B63_22g23 | ARM-like helical domain-containing protein |
| C4B63_238g17 | conserved hypothetical protein |
| C4B63_23g200 | Flagellum attachment zone protein 19 |
| C4B63_23g209 | conserved hypothetical protein |
| C4B63_23g234 | inosine-adenosine-guanosine-nucleosidehydrolase |
| C4B63_23g275 | ATP-dependent DEAD/H RNA helicase |
| C4B63_23g83 | Aminopeptidase M1 |
| C4B63_24g298 | CDP-diacylglycerol--inositol 3-phosphatidyltransferase |
| C4B63_24g325 | conserved hypothetical protein |
| C4B63_252g22 | T-complex protein 1, beta subunit |
| C4B63_25g294 | Thioredoxin-like |
| C4B63_25g317 | threonine synthase |
| C4B63_261g22c | conserved hypothetical protein |
| C4B63_26g204 | phosphomevalonate kinase protein |
| C4B63_26g271 | conserved hypothetical protein |
| C4B63_26g279 | ABC transporter |
| C4B63_26g287 | ABC transporter |
| C4B63_271g12 | thymidine kinase |
| C4B63_276g16 | conserved hypothetical protein |
| C4B63_28g106 | malic enzyme |
| C4B63_28g107 | malic enzyme |
| C4B63_28g64 | aspartate aminotransferase |
| C4B63_297g10 | polypyrimidine tract-binding protein |
| C4B63_2g235 | Co-chaperone protein P23 |
| C4B63_2g293 | conserved hypothetical protein |
| C4B63_2g294 | 40 kDa cyclophilin |
| C4B63_2g412 | R27-2 protein |
| C4B63_2g422 | cell division cycle protein 45 (CDC45) |
| C4B63_2g445 | Metallocarboxypeptidase 1 |
| C4B63_2g446 | Metallocarboxypeptidase 1 |
| C4B63_2g485 | T-complex protein 1, beta subunit |
| C4B63_2g594 | Snake venom metalloprotease inhibitor 02D01 |
| C4B63_2g690 | chaperonin alpha subunit |
| C4B63_2g829 | conserved hypothetical protein |
| C4B63_302g42c | histone H3 |
| C4B63_302g52c | histone H3 |
| C4B63_302g56c | histone H3 |
| C4B63_302g62c | histone H3 |
| C4B63_302g64c | histone H3 |
| C4B63_302g72c | histone H3 |
| C4B63_302g78c | histone H3 |
| C4B63_303g16 | trypanothione reductase |
| C4B63_303g6 | Histone H2B variant V |
| C4B63_303g7 | Histone H2B variant V |
| C4B63_30g140 | conserved hypothetical protein |
| C4B63_30g290 | mercaptopyruvate sulfurtransferase |
| C4B63_30g940c | Histone H2B variant V |
| C4B63_32g214 | 2-amino-3-ketobutyrate coenzyme A ligase |
| C4B63_32g217 | 2-amino-3-ketobutyrate coenzyme A ligase |
| C4B63_32g225 | 3-hydroxy-3-methylglutaryl-CoA synthase |
| C4B63_32g326 | cell division cycle protein |
| C4B63_340g1 | cell division cycle protein |
| C4B63_34g288 | glycerate kinase |
| C4B63_34g290 | glycerate kinase |
| C4B63_355g19 | inositol-3-phosphate synthase(fragment) |
| C4B63_37g366 | thimet oligopeptidase |
| C4B63_38g320c | conserved hypothetical protein |
| C4B63_38g38 | actin 3 |
| C4B63_38g39 | actin 3 |
| C4B63_39g333 | eukaryotic translation initiation factor 3 subunit 2 |
| C4B63_41g271 | conserved hypothetical protein |
| C4B63_42g178 | histidyl-tRNA synthetase |
| C4B63_42g242 | 2-hydroxy-3-oxopropionate reductase |
| C4B63_42g251 | conserved hypothetical protein |
| C4B63_42g63 | adenosine kinase |
| C4B63_42g70 | p22 protein precursor |
| C4B63_43g204 | thymidine kinase |
| C4B63_43g3 | kinesin |
| C4B63_44g192 | mitochondrial RNA binding protein |
| C4B63_45g99 | conserved hypothetical protein |
| C4B63_46g11 | chaperonin TCP20 |
| C4B63_46g91 | leucyl-tRNA synthetase |
| C4B63_47g53 | deoxyhypusine synthase |
| C4B63_48g101 | kinetoplastid kinetochore protein 1 |
| C4B63_48g85 | chaperonin HSP60, mitochondrial precursor |
| C4B63_48g87 | chaperonin HSP60, mitochondrial precursor |
| C4B63_4g199 | ethanolamine phosphotransferase |
| C4B63_4g200 | conserved hypothetical protein |
| C4B63_4g201 | Protein ILITYHIA |
| C4B63_4g220 | conserved hypothetical protein |
| C4B63_4g357 | Activator of 90 kDa heat shock protein ATPase homolog 1 |
| C4B63_4g375 | glucan 1,3-alpha-glucosidase |
| C4B63_4g484 | kinesin |
| C4B63_4g485 | kinesin |
| C4B63_4g527 | mitochondrial carrier protein |
| C4B63_4g558 | phosphatidic acid phosphatase protein |
| C4B63_4g64 | Mitochondrial ADP/ATP carrier protein 5 |
| C4B63_4g68 | metallo-peptidase, Clan MG, Family M24 |
| C4B63_4g77 | conserved hypothetical protein |
| C4B63_51g117 | Uncharacterized protein PB7E8.02 |
| C4B63_52g41 | mitochondrial DNA polymerase I protein C |
| C4B63_55g28 | kinesin |
| C4B63_55g29 | kinesin |
| C4B63_59g115 | stress-inducible protein STI1-like(fragment) |
| C4B63_61g127 | histone H2A variant Z |
| C4B63_62g82 | thimet oligopeptidase |
| C4B63_63g16 | chaperonin alpha subunit |
| C4B63_63g35 | histidine ammonia-lyase |
| C4B63_63g46 | 2,3-bisphosphoglycerate-independent phosphoglycerate mutase |
| C4B63_63g91 | GTPase activating protein |
| C4B63_69g25 | DNA excision/repair protein SNF2 |
| C4B63_69g26 | endo-beta-N-acetylglucosaminidase |
| C4B63_6g113 | ribonucleoside-diphosphate reductase large chain |
| C4B63_6g2482c | histone H3 |
| C4B63_6g2496c | histone H3 |
| C4B63_6g2498c | histone H3 |
| C4B63_6g2502c | histone H3 |
| C4B63_6g2504c | histone H3 |
| C4B63_6g2508c | histone H3 |
| C4B63_6g2516c | histone H3 |
| C4B63_6g2520c | histone H3 |
| C4B63_6g2522c | histone H3 |
| C4B63_6g2524c | histone H3 |
| C4B63_6g2526c | histone H3 |
| C4B63_6g2530c | histone H3 |
| C4B63_6g2546c | histone H3 |
| C4B63_6g2548c | histone H3 |
| C4B63_6g2556c | histone H3 |
| C4B63_6g2558c | histone H3 |
| C4B63_6g2564c | histone H3 |
| C4B63_6g2566c | histone H3 |
| C4B63_6g307 | heat shock protein 70 (hsp70)(fragment) |
| C4B63_6g308 | heat shock protein 70 (hsp70) |
| C4B63_6g310 | heat shock protein 70 (hsp70)(fragment) |
| C4B63_6g312 | heat shock protein 70 (hsp70) |
| C4B63_6g313 | heat shock protein 70 (hsp70)(fragment) |
| C4B63_6g315 | heat shock protein 70 (hsp70) |
| C4B63_6g509 | protein kinase |
| C4B63_6g532 | Putative trans-acting enoyl reductase MT2525 |
| C4B63_6g578 | histone H3 |
| C4B63_70g129 | metallo-peptidase, Clan MF, Family M17 |
| C4B63_70g94 | PIF1 helicase-like protein |
| C4B63_76g31 | conserved hypothetical protein,zinc finger protein family member |
| C4B63_78g20 | nucleolar RNA-binding protein |
| C4B63_78g25 | nucleolar RNA-binding protein |
| C4B63_79g54 | hypoxanthine-guanine phosphoribosyltransferase |
| C4B63_7g103 | conserved hypothetical protein |
| C4B63_7g210 | Ribonuclease P protein subunit p25-like protein |
| C4B63_7g400 | ER membrane protein complex subunit 1 |
| C4B63_81g101 | chaperonin HSP60, mitochondrial precursor |
| C4B63_81g104 | chaperonin HSP60, mitochondrial precursor |
| C4B63_81g105 | chaperonin HSP60, mitochondrial precursor |
| C4B63_81g19 | chaperonin HSP60, mitochondrial precursor |
| C4B63_81g40 | ATP-dependent DEAD/H RNA helicase |
| C4B63_83g73 | ribonucleoprotein p18, mitochondrial precursor |
| C4B63_84g21 | conserved hypothetical protein |
| C4B63_84g22 | conserved hypothetical protein |
| C4B63_84g87 | heat shock protein 85 |
| C4B63_84g88 | heat shock protein 85 |
| C4B63_84g89 | heat shock protein 85 |
| C4B63_84g90 | heat shock protein 85(fragment) |
| C4B63_85g60 | tricarboxylate carrier |
| C4B63_87g42 | conserved hypothetical protein |
| C4B63_8g196 | DNA ligase |
| C4B63_8g2646c | 10 kDa heat shock protein |
| C4B63_8g2656c | 10 kDa heat shock protein |
| C4B63_8g483 | asparagine synthetase a |
| C4B63_8g559 | Adenine phosphoribosyltransferase |
| C4B63_94g51 | Mucin-1 |
| C4B63_97g18 | heat shock protein 110 |
| C4B63_97g35 | P-type H+-ATPase |
| C4B63_9g209 | conserved hypothetical protein |
| C4B63_9g263 | trypanothione synthetase |
| C4B63_9g516 | heat shock protein Hsp20 |
| C4B63_9g524 | eukaryotic translation initiation factor 3 subunit k |

| Gene Ontology analysis of Hemin, Hb and No heme common DEGs (downregulated, t4 vs. t0) | | | | | | | | | | |
|---|---|---|---|---|---|---|---|---|---|---|
| | **BIOLOGICAL PROCESS** | | | | | | | | | |
| ID | Name | Bgd count | Result count | Result gene list | Pct of bgd | Fold enrichm | Odds ratio | P-value | Benjamini | Bonferroni |
| GO:0006525 | arginine metabolic process | 38 | 36 | C4B63_228g46,C | 94.7 | 58.39 | 1472.5 | 1,88E-64 | 1,68E-62 | 3,36E-62 |
| GO:0009064 | glutamine family amino acid metabolic | 55 | 36 | C4B63_228g46,C | 65.5 | 40.34 | 154.69 | 6,14E-53 | 2,75E-51 | 1,10E-50 |
| GO:0006599 | phosphagen metabolic process | 3 | 3 | C4B63_26g278,C | 100.0 | 61.63 | inf | 4,18E-06 | 3,25E-05 | 0.000748399141299 |
| GO:0016559 | peroxisome fission | 6 | 3 | C4B63_2g702,C4 | 50.0 | 30.82 | 61.95 | 8,07E-05 | 0.000577595 | 0.0144398783747 |
| GO:0046386 | deoxyribose phosphate catabolic proc | 2 | 2 | C4B63_163g21,C | 100.0 | 61.63 | inf | 0.000261388 | 0.001671021 | 0.0467886032493 |
| | **CELLULAR COMPONENT** | | | | | | | | | |
| ID | Name | Bgd count | Result count | Result gene list | Pct of bgd | Fold enrichm | Odds ratio | P-value | Benjamini | Bonferroni |
| GO:0005777 | peroxisome | 19 | 3 | C4B63_2g702,C4 | 15.8 | 9.73 | 11.6 | 0.003348108 | 0.016182523 | 0.0970951413332 |
| | **MOLECULAR FUNCTION** | | | | | | | | | |
| ID | Name | Bgd count | Result count | Result gene list | Pct of bgd | Fold enrichm | Odds ratio | P-value | Benjamini | Bonferroni |
| GO:0004180 | carboxypeptidase activity | 94 | 64 | C4B63_228g46,C | 68.1 | 41.96 | 238.88 | 2,43E-98 | 1,99E-96 | 1,99E-96 |
| GO:0016787 | hydrolase activity | 1860 | 78 | C4B63_112g40,C | 4.2 | 2.58 | 4.77 | 5,74E-19 | 7,85E-18 | 4,71E-17 |
| GO:0016775 | phosphotransferase activity, nitrogeno | 3 | 3 | C4B63_26g278,C | 100.0 | 61.63 | inf | 4,18E-06 | 3,81E-05 | 0.000342842064729 |

| Gene Ontology analysis of Hemin, Hb and No heme common DEGs (upregulated, t4 vs. t0) | | | | | | | | | | |
|---|---|---|---|---|---|---|---|---|---|---|
| | **BIOLOGICAL PROCESS** | | | | | | | | | |
| ID | Name | Bgd count | Result count | Result gene list | Pct of bgd | Fold enrichm | Odds ratio | P-value | Benjamini | Bonferroni |
| GO:0006457 | protein folding | 91 | 49 | C4B63_104g78,C | 53.8 | 20.14 | 53.77 | 2,47E-54 | 9,39E-52 | 9,39E-52 |
| GO:0030001 | metal ion transport | 36 | 10 | C4B63_6g238,C4 | 27.8 | 10.39 | 14.6 | 2,12E-08 | 2,01E-06 | 8,04E-06 |
| GO:0042254 | ribosome biogenesis | 76 | 12 | C4B63_123g35,C | 15.8 | 5.91 | 7.15 | 6,83E-07 | 3,71E-05 | 0.000259405458758 |
| | **CELLULAR COMPONENT** | | | | | | | | | |
| ID | Name | Bgd count | Result count | Result gene list | Pct of bgd | Fold enrichm | Odds ratio | P-value | Benjamini | Bonferroni |
| GO:0005737 | cytoplasm | 813 | 50 | C4B63_104g78,C | 6.2 | 2.3 | 2.77 | 1,16E-08 | 1,85E-07 | 9,26E-07 |
| GO:0005788 | endoplasmic reticulum lumen | 4 | 3 | C4B63_104g78,C | 75.0 | 28.06 | 110.67 | 7,39E-05 | 0.000591466 | 0.00591466477824 |
| GO:0031981 | nuclear lumen | 94 | 10 | C4B63_123g35,C | 10.6 | 3.98 | 4.49 | 0.000193793 | 0.001354101 | 0.0155034768978 |
| GO:0101031 | chaperone complex | 2 | 2 | C4B63_252g22,C | 100.0 | 37.41 | inf | 0.000711481 | 0.003954588 | 0.0569185043213 |
| GO:0005739 | mitochondrion | 131 | 11 | C4B63_11g8,C4B | 8.4 | 3.14 | 3.46 | 0.000741485 | 0.003954588 | 0.0593188206697 |
| | **MOLECULAR FUNCTION** | | | | | | | | | |
| ID | Name | Bgd count | Result count | Result gene list | Pct of bgd | Fold enrichm | Odds ratio | P-value | Benjamini | Bonferroni |
| GO:0005524 | ATP binding | 1110 | 96 | C4B63_10g439,C | 8.6 | 3.24 | 5.21 | 1,65E-28 | 1,90E-26 | 3,96E-26 |
| GO:0043167 | ion binding | 2086 | 113 | C4B63_104g78,C | 5.4 | 2.03 | 3.14 | 9,35E-17 | 1,40E-15 | 2,24E-14 |
| GO:0008887 | glycerate kinase activity | 4 | 4 | C4B63_34g287,C | 100.0 | 37.41 | inf | 4,98E-07 | 6,29E-06 | 0.000119427682399 |

| Gene Ontology analysis of Hemin, Hb and No heme common DEGs (downregulated, t24 vs. t0) | | | | | | | | | | |
|---|---|---|---|---|---|---|---|---|---|---|
| | **BIOLOGICAL PROCESS** | | | | | | | | | |
| ID | Name | Bgd count | Result count | Result gene list | Pct of bgd | Fold enrichm | Odds ratio | P-value | Benjamini | Bonferroni |
| GO:0006525 | arginine metabolic process | 38 | 33 | C4B63_228g46,C | 86.8 | 54.7 | 539.92 | 3,06E-56 | 2,36E-54 | 4,72E-54 |
| GO:0009064 | glutamine family amino acid metabolic | 55 | 33 | C4B63_228g46,C | 60.0 | 37.8 | 122.46 | 6,49E-47 | 2,50E-45 | 1,00E-44 |
| GO:1901564 | organonitrogen compound metabolic p | 1615 | 49 | C4B63_112g40,C | 3.0 | 1.91 | 2.47 | 1,52E-06 | 1,17E-05 | 0.000234481029763 |
| | **CELLULAR COMPONENT** | | | | | | | | | |
| ID | Name | Bgd count | Result count | Result gene list | Pct of bgd | Fold enrichm | Odds ratio | P-value | Benjamini | Bonferroni |
| GO:0016021 | integral component of membrane | 2487 | 51 | C4B63_112g40,C | 2.1 | 1.29 | 1.48 | 0.019632002 | 0.186504025 | 0.373008051728 |
| | **MOLECULAR FUNCTION** | | | | | | | | | |
| ID | Name | Bgd count | Result count | Result gene list | Pct of bgd | Fold enrichm | Odds ratio | P-value | Benjamini | Bonferroni |
| GO:0004180 | carboxypeptidase activity | 94 | 58 | C4B63_228g46,C | 61.7 | 38.87 | 173.4 | 2,77E-85 | 1,85E-83 | 1,85E-83 |
| GO:0016997 | alpha-sialidase activity | 711 | 23 | C4B63_109g61,C | 3.2 | 2.04 | 2.29 | 0.000748563 | 0.005572642 | 0.0501537786633 |

| Gene Ontology analysis of Hemin, Hb and No heme common DEGs (upregulated, t24 vs. t0) | | | | | | | | | | |
|---|---|---|---|---|---|---|---|---|---|---|
| | **BIOLOGICAL PROCESS** | | | | | | | | | |
| ID | Name | Bgd count | Result count | Result gene list | Pct of bgd | Fold enrichm | Odds ratio | P-value | Benjamini | Bonferroni |
| GO:0006457 | protein folding | 91 | 36 | C4B63_113g22,C | 39.6 | 14.86 | 28.24 | 9,88E-34 | 3,65E-31 | 3,65E-31 |
| | **CELLULAR COMPONENT** | | | | | | | | | |
| ID | Name | Bgd count | Result count | Result gene list | Pct of bgd | Fold enrichm | Odds ratio | P-value | Benjamini | Bonferroni |
| GO:0005694 | chromosome | 168 | 60 | C4B63_112g84c, | 35.7 | 13.42 | 27.22 | 2,09E-53 | 2,80E-52 | 1,40E-51 |
| GO:0005634 | nucleus | 424 | 40 | C4B63_10g406,C | 9.4 | 3.54 | 4.41 | 1,33E-12 | 8,11E-12 | 8,92E-11 |
| GO:0005737 | cytoplasm | 813 | 38 | C4B63_113g25,C | 4.7 | 1.76 | 1.95 | 0.000388189 | 0.001529922 | 0.0260086900955 |
| GO:0101031 | chaperone complex | 2 | 2 | C4B63_252g22,C | 100.0 | 37.57 | inf | 0.000705267 | 0.002486995 | 0.0472529220645 |
| | **MOLECULAR FUNCTION** | | | | | | | | | |
| ID | Name | Bgd count | Result count | Result gene list | Pct of bgd | Fold enrichm | Odds ratio | P-value | Benjamini | Bonferroni |
| GO:0005515 | protein binding | 1216 | 104 | C4B63_112g84c, | 8.6 | 3.21 | 5.45 | 3,47E-31 | 1,34E-29 | 8,05E-29 |
| GO:0005524 | ATP binding | 1110 | 73 | C4B63_10g517,C | 6.6 | 2.47 | 3.32 | 2,71E-14 | 5,71E-13 | 6,28E-12 |
| GO:0043167 | ion binding | 2086 | 91 | C4B63_10g517,C | 4.4 | 1.64 | 2.11 | 1,01E-07 | 1,02E-06 | 2,34E-05 |
| GO:0008235 | metalloexopeptidase activity | 16 | 5 | C4B63_12g153,C | 31.3 | 11.74 | 16.98 | 4,39E-05 | 0.000424503 | 0.0101880839521 |

### Figure S1

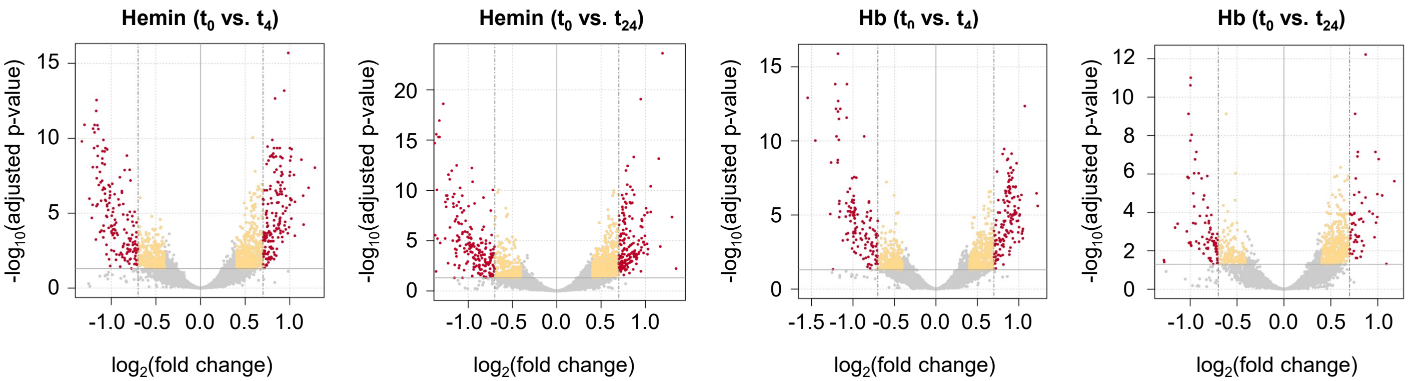

Hemin ($t_0$ vs. $t_4$)
Hemin ($t_0$ vs. $t_{24}$)
Hb ($t_0$ vs. $t_4$)
Hb ($t_0$ vs. $t_{24}$)
-log10(adjusted p-value)
log2(fold change)

### Table S3

#### Hemin and Hb common DEGs (t4 vs t0)

##### Downregulated genes

| ID | Annotation | Hemin | | Hb | |
|---|---|---|---|---|---|
| | | log2FC | padj | log2FC | padj |
| C4B63_18g39 | CRAL/TRIO, N-terminal domain/CRAL/TRIO domain c | -1,808774747 | 1,21E-18 | -1,5460513 | 1,24E-13 |
| C4B63_49g168 | conserved hypothetical protein | -1,171980321 | 0,00057483 | -0,80297 | 0,020838019 |
| C4B63_27g1020c | RNA-binding protein | -0,994646682 | 0,00021044 | -1,0598988 | 0,0001686 |
| C4B63_43g225 | conserved hypothetical protein | -0,815397296 | 0,01641791 | -1,0107327 | 0,001827284 |
| C4B63_24g199 | conserved hypothetical protein | -0,808096617 | 0,00261519 | -0,9946761 | 0,000288074 |
| C4B63_22g780c | calpain-like cysteine peptidase | -0,792715519 | 0,00349498 | -0,7770126 | 0,024259522 |
| C4B63_14g158 | serine/threonine kinase | -0,789239268 | 0,00070193 | -0,8141541 | 0,006764196 |
| C4B63_13g223 | zinc-finger protein family member | -0,783410929 | 0,01104509 | -0,8159978 | 0,021248307 |
| C4B63_2g547 | receptor-type adenylate cyclase | -1,021736703 | 0,00000206 | -0,6383037 | 0,009906963 |
| C4B63_2g548 | receptor-type adenylate cyclase | -1,01278119 | 6,44E-07 | -0,6685913 | 0,019117321 |
| C4B63_24g313 | protein kinase | -0,892294129 | 0,00438955 | -0,6949108 | 0,048874265 |
| C4B63_156g18 | trans-sialidase | -0,820056863 | 0,00128107 | -0,5831029 | 0,023175727 |
| C4B63_2g309 | NIMA-related kinase | -0,73843064 | 0,00321431 | -0,6542999 | 0,018612444 |
| C4B63_14g101 | conserved hypothetical protein | -0,73001501 | 0,00711696 | -0,5522618 | 0,044157687 |
| C4B63_47g128 | trans-sialidase | -0,722077618 | 0,00449876 | -0,6172614 | 0,030082599 |
| C4B63_37g368 | Proline dehydrogenase, mitochondrial | -0,719624519 | 0,00038991 | -0,6562838 | 0,004168137 |
| C4B63_17g61 | conserved hypothetical protein | -0,712900888 | 0,0030949 | -0,663958 | 0,006473053 |
| C4B63_55g86 | peptidyl-prolyl cis-trans isomerase | -0,692982205 | 0,00101689 | -0,5493152 | 0,030677799 |
| C4B63_130g39 | trans-sialidase, Group II | -0,692761653 | 0,00380615 | -0,6601727 | 0,004984789 |
| C4B63_156g6 | trans-sialidase | -0,692091407 | 0,00961706 | -0,634388 | 0,021248307 |
| C4B63_19g141 | electron transfer flavoprotein | -0,681873252 | 0,00017366 | -0,6525703 | 0,000156788 |
| C4B63_30g222 | conserved hypothetical protein | -0,675207601 | 9,35E-07 | -0,4475514 | 0,003576183 |
| C4B63_52g153 | conserved hypothetical protein | -0,67396387 | 0,01384785 | -0,6660266 | 0,030701785 |
| C4B63_8g495 | Regulator of chromosome condensation (RCC1) repe | -0,672881236 | 0,00028299 | -0,5601292 | 0,025263938 |
| C4B63_82g448c | conserved hypothetical protein | -0,648960027 | 0,00123418 | -0,6228363 | 0,005429819 |
| C4B63_4g511 | zinc-finger protein | -0,643850614 | 0,00185179 | -0,5390746 | 0,033548808 |
| C4B63_134g54 | trans-sialidase, Group V | -0,642876138 | 0,00183035 | -0,4698703 | 0,032182865 |
| C4B63_134g49 | trans-sialidase, Group V | -0,623373149 | 0,00673586 | -0,4752388 | 0,020329456 |
| C4B63_157g36 | trans-sialidase, Group II | -0,620882028 | 0,03034162 | -0,5483791 | 0,03207292 |
| C4B63_9g387 | conserved hypothetical protein | -0,618701592 | 0,00076891 | -0,4105115 | 0,03207292 |
| C4B63_10g494 | thymine-7-hydroxylase | -0,614017026 | 0,00382912 | -0,5652601 | 0,013757536 |
| C4B63_9g448 | superoxide dismutase | -0,607178566 | 0,00028299 | -0,4963374 | 0,013034904 |
| C4B63_354g1 | electron transfer flavoprotein | -0,605761908 | 0,00586106 | -0,6533645 | 0,005697135 |
| C4B63_34g117 | cytochrome b5-dependent oleate desaturase | -0,605479999 | 0,00460831 | -0,5302052 | 0,04066761 |
| C4B63_22g96 | protein kinase | -0,60258246 | 0,01238382 | -0,6543142 | 0,006359919 |
| C4B63_9g445 | conserved hypothetical protein | -0,599061375 | 0,02285797 | -0,5586467 | 0,03518388 |
| C4B63_42g100 | calcium-binding protein | -0,598267766 | 0,00479273 | -0,5089338 | 0,021645331 |
| C4B63_192g2 | amino acid permease 24 | -0,597546613 | 0,00472846 | -0,4979387 | 0,035045199 |
| C4B63_23g17 | Battenin | -0,59721217 | 0,00069049 | -0,4584686 | 0,014625786 |
| C4B63_193g27 | protein associated with differentiation 4 | -0,593366171 | 0,0328493 | -0,6147894 | 0,021842808 |
| C4B63_113g47 | Generative cell specific 1 protein | -0,589224099 | 0,02400521 | -0,6057219 | 0,021162993 |
| C4B63_40g146 | Gamma carbonic anhydrase 3, mitochondrial | -0,582182177 | 0,00639445 | -0,636335 | 0,001944739 |
| C4B63_21g33 | small GTP-binding protein Rab7 | -0,57016298 | 0,00054685 | -0,6077617 | 0,0007303 |
| C4B63_1g480 | trans-sialidase, Group V | -0,568699562 | 0,00625476 | -0,49473 | 0,019741116 |
| C4B63_77g33 | conserved hypothetical protein | -0,56505283 | 0,00422767 | -0,5444212 | 0,005515167 |
| C4B63_14g92 | conserved hypothetical protein | -0,563674405 | 0,01765838 | -0,6464864 | 0,007395037 |
| C4B63_16g1800c | conserved hypothetical protein | -0,557125719 | 0,01412477 | -0,6092664 | 0,004428084 |
| C4B63_278g3 | conserved hypothetical protein | -0,55609634 | 0,01218422 | -0,5302308 | 0,023791923 |
| C4B63_21g64 | conserved hypothetical protein | -0,55449461 | 0,00453296 | -0,5905586 | 0,023791923 |
| C4B63_13g317 | vesicle-associated membrane protein | -0,553362947 | 0,0017064 | -0,506283 | 0,0071948 |
| C4B63_63g52 | conserved hypothetical protein | -0,543715991 | 0,01657612 | -0,6062091 | 0,003597327 |
| C4B63_31g61 | dual-specificity protein phosphatase | -0,541297729 | 0,01511715 | -0,5065708 | 0,044078245 |
| C4B63_18g308 | sphingosine 1-phosphate lyase | -0,521463368 | 0,00228593 | -0,5420295 | 0,001740963 |
| C4B63_21g27 | arginine permease(fragment) | -0,519748902 | 0,04767094 | -0,5205113 | 0,043951003 |
| C4B63_21g112 | conserved hypothetical protein | -0,516757842 | 0,01389474 | -0,5847305 | 0,010158898 |
| C4B63_45g82 | conserved hypothetical protein | -0,514932996 | 0,00864866 | -0,4458068 | 0,015174583 |
| C4B63_459g4 | conserved hypothetical protein | -0,513981065 | 0,01236402 | -0,5382728 | 0,031620556 |
| C4B63_4g505 | ADP-ribosylation factor-like protein | -0,509282325 | 0,00381269 | -0,4550279 | 0,021969701 |
| C4B63_70g572c | cytochrome b5 | -0,507647754 | 0,04832744 | -0,5460632 | 0,025181875 |
| C4B63_25g207 | hydrolase-like protein | -0,506710476 | 0,02157845 | -0,5497422 | 0,033160367 |
| C4B63_4g180 | serine/threonine protein kinase | -0,500578733 | 0,00026607 | -0,5055941 | 0,000427798 |
| C4B63_43g74 | retrotransposon hot spot (RHS) protein | -0,498587976 | 0,01585039 | -0,4907133 | 0,008901829 |
| C4B63_4g143 | conserved hypothetical protein | -0,498571243 | 0,03195312 | -0,4970781 | 0,025865722 |
| C4B63_43g134 | cytochrome c oxidase subunit VI | -0,49686699 | 0,00464391 | -0,5255918 | 0,016838085 |
| C4B63_4g283 | conserved hypothetical protein | -0,495554516 | 0,00335186 | -0,6127837 | 0,000407948 |
| C4B63_25g236 | conserved hypothetical protein | -0,49363397 | 0,0303703 | -0,4934296 | 0,041572205 |
| C4B63_4g496 | transporter | -0,492842334 | 0,00270345 | -0,6362727 | 0,0000584 |
| C4B63_173g6 | Soluble NSF attachment protein, SNAP (paraflagelar | -0,488869117 | 0,00357026 | -0,4648574 | 0,006068708 |
| C4B63_21g25 | arginine permease | -0,486813727 | 0,01297888 | -0,5232119 | 0,007929772 |
| C4B63_22g167 | calpain-like cysteine peptidase | -0,485824991 | 0,00614085 | -0,456573 | 0,01273885 |
| C4B63_13g146 | conserved hypothetical protein | -0,485593523 | 0,02543275 | -0,4840782 | 0,040147495 |
| C4B63_309g8 | retrotransposon hot spot (RHS) protein | -0,478151858 | 0,03991761 | -0,4812183 | 0,006338857 |
| C4B63_47g59 | conserved hypothetical protein | -0,476746018 | 0,02871472 | -0,517646 | 0,018593091 |
| C4B63_44g231 | amino acid transporter | -0,47649774 | 0,03792391 | -0,5205787 | 0,006338857 |
| C4B63_60g198 | retrotransposon hot spot (RHS) protein | -0,474491569 | 0,01729889 | -0,482989 | 0,03222943 |
| C4B63_43g174 | Cytochrome b5 domain-containing protein 1 | -0,473822678 | 0,00690065 | -0,4484889 | 0,015893671 |
| C4B63_2g156 | protein kinase A catalytic subunit 3 | -0,470032629 | 0,00365284 | -0,4478492 | 0,033160367 |
| C4B63_14g53 | Basal body-orientation factor 1 | -0,464910439 | 0,00818288 | -0,5072042 | 0,003814788 |
| C4B63_20g39 | conserved hypothetical protein | -0,461761816 | 0,04195449 | -0,5302818 | 0,004162844 |
| C4B63_48g693c | conserved hypothetical protein | -0,46127976 | 0,01487736 | -0,4874072 | 0,027698415 |
| C4B63_16g104 | conserved hypothetical protein | -0,460622112 | 0,04748063 | -0,5088029 | 0,033289423 |
| C4B63_175g11 | Homologous-pairing protein 2 homolog | -0,44889857 | 0,03601283 | -0,5557569 | 0,02478701 |
| C4B63_44g106 | cytochrome c oxidase VIII (COX VIII) | -0,441974089 | 0,0000861 | -0,5109784 | 0,000986354 |
| C4B63_3g1092 | alanine racemase | -0,440397024 | 0,01480092 | -0,4608152 | 0,006117138 |
| C4B63_35g90 | flagellar calcium-binding protein | -0,438905177 | 0,00941958 | -0,4157322 | 0,013805357 |
| C4B63_61g147 | conserved hypothetical protein | -0,437378777 | 0,04490242 | -0,5221927 | 0,016117594 |
| C4B63_180g10 | retrotransposon hot spot (RHS) protein | -0,436787792 | 0,02605464 | -0,4231829 | 0,040794699 |
| C4B63_30g247 | glucosamine-fructose-6-phosphate aminotransferase | -0,429795757 | 0,01243976 | -0,4355021 | 0,008059083 |
| C4B63_27g185 | conserved hypothetical protein | -0,421891782 | 0,03069738 | -0,4147184 | 0,034520192 |
| C4B63_32g242 | Phosphorylated CTD interacting factor 1 WW domai | -0,421384575 | 0,00485976 | -0,4469063 | 0,003637157 |
| C4B63_99g112 | alanine racemase | -0,415768718 | 0,0321732 | -0,4197702 | 0,041403185 |
| C4B63_2g712 | galactokinase | -0,414605743 | 0,02205543 | -0,5168173 | 0,004123273 |
| C4B63_20g35 | conserved hypothetical protein | -0,410970776 | 0,0209229 | -0,4284884 | 0,022938251 |
| C4B63_70g88 | ATG16/SAP18/CVT11/APG16 | -0,410957882 | 0,0093311 | -0,4054939 | 0,016190048 |
| C4B63_6g511 | Hsp70-Hsp90 organizing protein 3 | -0,409799709 | 0,02699253 | -0,4855274 | 0,007159305 |
| C4B63_18g322 | conserved hypothetical protein | -0,409452141 | 0,01839807 | -0,5189421 | 0,003282909 |
| C4B63_13g310 | NAD(P)-dependent oxidoreductase | -0,405850812 | 0,04192059 | -0,4078745 | 0,038304207 |
| C4B63_14g43 | centrin-3 | -0,404864269 | 0,04810396 | -0,4338507 | 0,006932193 |
| C4B63_17g91 | Tyrosine-protein phosphatase DSP2 | -0,403673477 | 0,04233963 | -0,5008877 | 0,019518808 |
| C4B63_14g60 | conserved hypothetical protein | -0,402056758 | 0,008501 | -0,4081772 | 0,013386637 |
| C4B63_20g26 | conserved hypothetical protein | -0,40204042 | 0,04483122 | -0,4274802 | 0,016586616 |
| C4B63_47g60 | conserved hypothetical protein | -0,400716189 | 0,00499329 | -0,4592238 | 0,001345146 |
| C4B63_4g242 | protein kinase A catalytic subunit 3 | -0,688121206 | 0,00673586 | -0,7733653 | 0,001176318 |
| C4B63_19g16 | Ring finger domain containing protein | -0,675426331 | 0,03130577 | -0,7262018 | 0,019518808 |
| C4B63_21g63 | conserved hypothetical protein | -0,667304955 | 0,00156322 | -0,7542612 | 0,002876946 |
| C4B63_13g256 | conserved hypothetical protein | -0,662083269 | 0,00888957 | -0,7556758 | 0,002972805 |
| C4B63_112g66 | Gamma carbonic anhydrase 2, mitochondrial | -0,6026594 | 0,00225237 | -0,7087577 | 0,000886895 |
| C4B63_351g33c | calpain-like cysteine peptidase | -0,59539363 | 0,03837682 | -0,703454 | 0,020072185 |

##### Upregulated genes

| ID | Annotation | Hemin | | Hb | |
|---|---|---|---|---|---|
| | | log2FC | padj | log2FC | padj |
| C4B63_9g516 | heat shock protein Hsp20 | 1,00565677 | 0,00000103 | 0,8758497 | 0,0000124 |
| C4B63_61g94 | conserved hypothetical protein | 0,85504432 | 0,00240857 | 0,81983049 | 0,00055612 |
| C4B63_35g168 | 3-Beta-hydroxysteroid-delta(8), delta(7)-isomerase | 0,81617516 | 0,00072982 | 0,57398802 | 0,04283621 |
| C4B63_62g90 | ribosome biogenesis protein | 0,79171957 | 0,00025571 | 0,67343914 | 0,00102674 |
| C4B63_77g26 | conserved hypothetical protein | 0,78304588 | 0,01218422 | 0,60049583 | 0,03079491 |
| C4B63_34g340 | mucin TcSMUGL | 0,74058842 | 0,02682232 | 0,81464619 | 0,02189048 |
| C4B63_300g99c | mucin TcSMUGL | 0,73458697 | 0,01122026 | 0,80646254 | 0,00502762 |
| C4B63_97g35 | P-type H+-ATPase | 0,72736986 | 0,00049114 | 0,76098308 | 0,00015099 |
| C4B63_303g16 | trypanothione reductase | 0,72570056 | 0,00183416 | 0,59731379 | 0,01894475 |
| C4B63_6g562 | NAD(P)-dependent steroid dehydrogenase protein | 0,72159197 | 0,00034885 | 0,61910085 | 0,00338041 |
| C4B63_44g117 | conserved hypothetical protein | 0,7175753 | 2,97E-07 | 0,73774399 | 1,18E-08 |
| C4B63_8g483 | asparagine synthetase a | 0,71569028 | 0,003819 | 0,60375666 | 0,02124831 |
| C4B63_34g336 | mucin TcSMUGL | 0,71310984 | 0,00323706 | 0,66131359 | 0,0154586 |
| C4B63_80g17 | ATP binding protein-like protein | 0,70780564 | 0,00467336 | 0,47778155 | 0,04239551 |
| C4B63_180g2 | conserved hypothetical protein | 0,7021827 | 0,02523343 | 0,68791672 | 0,04833888 |
| C4B63_210g28 | Shk1 kinase-binding protein 15 | 0,69180991 | 0,01995118 | 0,52386448 | 0,03866332 |
| C4B63_61g93 | conserved hypothetical protein | 0,68931569 | 0,00110568 | 0,70136851 | 0,00011968 |
| C4B63_2g209 | Present in the outer mitochondrial membrane proteom | 0,685954 | 0,00068582 | 0,58746996 | 0,00506595 |
| C4B63_34g1506c | mucin TcSMUGL | 0,68194644 | 0,0109567 | 0,69220719 | 0,00702638 |
| C4B63_42g242 | 2-hydroxy-3-oxopropionate reductase | 0,68156526 | 0,00025571 | 0,50133803 | 0,01808554 |
| C4B63_79g25 | conserved hypothetical protein | 0,6780471 | 0,0034413 | 0,65142486 | 0,00549148 |
| C4B63_81g18 | chaperonin HSP60, mitochondrial precursor(fragment | 0,67365943 | 0,00872053 | 0,66277033 | 0,00986659 |
| C4B63_34g329 | mucin TcSMUGL | 0,67192231 | 0,01185577 | 0,68360139 | 0,03207292 |
| C4B63_6g19 | NAD(P)-dependent steroid dehydrogenase protein | 0,66648721 | 0,00076277 | 0,65986918 | 0,00110491 |
| C4B63_207g2 | fibrillarin | 0,66541027 | 0,01209556 | 0,57307728 | 0,03316037 |
| C4B63_34g1490c | mucin TcSMUGL | 0,66481496 | 0,02814741 | 0,74646015 | 0,01048459 |
| C4B63_111g29 | conserved hypothetical protein | 0,66444132 | 0,00134636 | 0,58522015 | 0,00464395 |
| C4B63_61g149 | eukaryotic translation initiation factor 3 subunit a | 0,66197769 | 0,00089882 | 0,59241558 | 0,00384603 |
| C4B63_34g1494c | mucin TcSMUGL | 0,66177485 | 0,01908327 | 0,63183823 | 0,03871905 |
| C4B63_11g83 | Eukaryotic translation initiation factor 3 subunit F | 0,65658775 | 0,0050611 | 0,54332214 | 0,03676837 |
| C4B63_6g244 | cation transporter | 0,65310657 | 0,02111985 | 0,66958391 | 0,02305951 |
| C4B63_300g96c | mucin TcSMUGL | 0,65214362 | 0,01598036 | 0,6595291 | 0,02402572 |
| C4B63_28g84 | conserved hypothetical protein | 0,65050749 | 0,00266558 | 0,68058183 | 0,00131705 |
| C4B63_25g113 | GMP reductase | 0,64791169 | 0,03056201 | 0,60953006 | 0,03435989 |
| C4B63_55g67 | DNA-directed rna polymerase I largest subunit | 0,64739513 | 0,00031023 | 0,6400291 | 2,81E-07 |
| C4B63_85g60 | tricarboxylate carrier | 0,64717185 | 0,0009647 | 0,58648819 | 0,0023785 |
| C4B63_396g14 | Present in the outer mitochondrial membrane proteom | 0,64524201 | 0,00030982 | 0,67458806 | 0,00036816 |
| C4B63_20g310 | Nucleolar complex protein 4 homolog | 0,64391367 | 0,00980129 | 0,60174773 | 0,02683434 |
| C4B63_227g9 | mevalonate-diphosphate decarboxylase | 0,64304655 | 0,00216126 | 0,5321928 | 0,02083802 |
| C4B63_40g25 | Biotin--acetyl-CoA-carboxylase ligase | 0,64167702 | 0,00178396 | 0,62154638 | 0,0034175 |
| C4B63_162g33 | chaperonin TCP20 | 0,63891244 | 0,00025571 | 0,49928785 | 0,00935275 |
| C4B63_34g1453c | mucin TcSMUGL | 0,63364335 | 0,0045584 | 0,6646673 | 0,0496854 |
| C4B63_300g106c | mucin TcSMUGL | 0,63219342 | 0,0480428 | 0,72255894 | 0,04018165 |
| C4B63_6g532 | Putative trans-acting enoyl reductase MT2525 | 0,62986732 | 0,01344982 | 0,5209624 | 0,04977837 |
| C4B63_8g510 | conserved hypothetical protein | 0,62952073 | 0,00823687 | 0,45477568 | 0,04446288 |
| C4B63_17g90 | T-complex protein 1, beta subunit | 0,62880306 | 0,00096261 | 0,60303708 | 0,00133323 |
| C4B63_6g227 | Eukaryotic translation initiation factor 3 subunit E | 0,62590605 | 0,00690065 | 0,60722401 | 0,00747062 |
| C4B63_30g162 | cystathionine beta-synthase | 0,62568419 | 0,00820089 | 0,67161852 | 0,00279626 |
| C4B63_44g104 | diphthine synthase | 0,62088082 | 0,00375278 | 0,67000289 | 0,00345095 |
| C4B63_19g82 | RNA polymerase I second largest subunit | 0,61978041 | 0,00188651 | 0,60301829 | 0,00394033 |
| C4B63_23g116 | Importin beta-like SAD2 | 0,61898145 | 0,00085953 | 0,48885278 | 0,01492251 |
| C4B63_34g1510c | mucin TcSMUGL | 0,61576722 | 0,01553482 | 0,60991118 | 0,0302182 |
| C4B63_37g373 | ribosome biogenesis protein | 0,61265563 | 0,01031193 | 0,54708999 | 0,01671189 |
| C4B63_143g50 | cystathionine beta-synthase | 0,6104597 | 0,00342453 | 0,66297337 | 0,00018468 |
| C4B63_25g240 | Macro domain containing protein | 0,5999245 | 0,02479219 | 0,47213445 | 0,00906554 |
| C4B63_13g156 | elongation factor 2 | 0,59852154 | 0,01250699 | 0,63839638 | 0,00781696 |
| C4B63_13g154 | elongation factor 2 | 0,59849943 | 0,01657612 | 0,65539508 | 0,0061175 |
| C4B63_6g236 | cation transporter | 0,59842951 | 0,00044033 | 0,55585436 | 0,00120102 |
| C4B63_13g155 | elongation factor 2 | 0,59436395 | 0,02673852 | 0,62872533 | 0,01794337 |
| C4B63_90g55 | TPR-repeat protein(fragment) | 0,58881753 | 0,00889475 | 0,5524831 | 0,01492493 |
| C4B63_13g153 | elongation factor 2 | 0,58657142 | 0,01906923 | 0,64886722 | 0,00764529 |
| C4B63_34g1464c | mucin TcSMUGL | 0,58587889 | 0,04721713 | 0,66169769 | 0,01655506 |
| C4B63_143g47 | cystathionine beta-synthase | 0,58558689 | 0,01262083 | 0,56432212 | 0,00477393 |
| C4B63_134g37 | orotidine-5-phosphate decarboxylase/orotate phospho | 0,58540324 | 0,0003152 | 0,53215102 | 0,00147212 |
| C4B63_19g205 | glycosomal transporter (GAT3) | 0,58511246 | 8,99E-11 | 0,45356899 | 0,0000149 |
| C4B63_30g163 | cystathionine beta-synthase | 0,58485866 | 0,01050773 | 0,60372353 | 0,00097809 |
| C4B63_128g49 | nucleolar RNA helicase II | 0,58360426 | 0,01774239 | 0,53862384 | 0,03142691 |
| C4B63_38g147 | spermidine synthase | 0,58240998 | 0,00170398 | 0,4547762 | 0,0234845 |
| C4B63_34g1477c | mucin TcSMUGL | 0,57991892 | 0,02038178 | 0,66748214 | 0,00967332 |
| C4B63_112g72 | cystathionine beta-synthase | 0,57959669 | 0,00438955 | 0,53654692 | 0,00364659 |
| C4B63_13g152 | elongation factor 2 | 0,57683919 | 0,02230005 | 0,60632438 | 0,01725144 |
| C4B63_48g103 | Nucleolar complex-associated protein 3 | 0,5756966 | 0,03760552 | 0,61475548 | 0,01750424 |
| C4B63_57g65 | membrane-bound acid phosphatase 2 | 0,57517147 | 0,00250675 | 0,50875166 | 0,00441221 |
| C4B63_35g130 | orotidine-5-phosphate decarboxylase/orotate phospho | 0,57296011 | 0,0000147 | 0,53407402 | 0,0002612 |
| C4B63_112g68 | cystathionine beta-synthase | 0,57160383 | 0,01242776 | 0,6156237 | 0,00341898 |
| C4B63_112g69 | cystathionine beta-synthase | 0,5687613 | 0,02676297 | 0,64621355 | 0,00249158 |
| C4B63_8g191 | JmjC domain-containing protein E | 0,56635473 | 0,0480428 | 0,60590944 | 0,04367625 |
| C4B63_70g55 | cystathionine beta-synthase | 0,5660676 | 0,01024739 | 0,64949846 | 0,0001682 |
| C4B63_143g51 | cystathionine beta-synthase | 0,56340737 | 0,00506352 | 0,6446021 | 0,00018851 |
| C4B63_214g13 | UDP-Gal or UDP-GlcNAc-dependent glycosyltransfera | 0,55886857 | 0,01878425 | 0,61468403 | 0,00863432 |
| C4B63_70g51 | cystathionine beta-synthase | 0,55761122 | 0,03686757 | 0,69145061 | 0,00093842 |
| C4B63_34g1519c | mucin TcSMUGL | 0,55555292 | 0,03896009 | 0,67908185 | 0,02643896 |
| C4B63_28g107 | malic enzyme | 0,55542601 | 0,00012244 | 0,4421036 | 0,01487071 |
| C4B63_4g220 | conserved hypothetical protein | 0,55072293 | 0,00271303 | 0,40129803 | 0,0423981 |
| C4B63_70g50 | cystathionine beta-synthase | 0,54867342 | 0,02363591 | 0,62189795 | 0,00155221 |
| C4B63_83g68 | protein phosphatase 2C | 0,54776652 | 0,04617105 | 0,70481942 | 0,00427412 |
| C4B63_19g195 | Nuclear pore complex protein 158 | 0,54690978 | 0,00018509 | 0,46788112 | 0,00173695 |
| C4B63_1g57 | AAA ATPase | 0,54671301 | 0,00404143 | 0,50514117 | 0,0097805 |
| C4B63_9g446 | Probable methyltransferase TARBP1 | 0,545777 | 0,02941617 | 0,58045248 | 0,01581239 |
| C4B63_70g53 | cystathionine beta-synthase | 0,54437063 | 0,01002424 | 0,67283396 | 0,0000763 |
| C4B63_81g19 | chaperonin HSP60, mitochondrial precursor | 0,54294315 | 0,03799918 | 0,57413201 | 0,02175757 |
| C4B63_157g32 | inosine-5'-monophosphate dehydrogenase | 0,54151834 | 0,0033574 | 0,4964959 | 0,00745929 |
| C4B63_70g49 | cystathionine beta-synthase | 0,5400358 | 0,03435513 | 0,55437834 | 0,01795306 |
| C4B63_70g52 | cystathionine beta-synthase | 0,53866209 | 0,02313629 | 0,64086544 | 0,00037867 |
| C4B63_13g157 | elongation factor 2 | 0,53811482 | 0,0224764 | 0,56718163 | 0,0136712 |
| C4B63_192g26 | conserved hypothetical protein | 0,53517449 | 0,01789637 | 0,48863487 | 0,02829124 |
| C4B63_22g292 | deoxyhypusine synthase | 0,53298327 | 0,03977727 | 0,50937501 | 0,03207292 |
| C4B63_112g70 | cystathionine beta-synthase | 0,53294212 | 0,01657612 | 0,59619668 | 0,00168277 |
| C4B63_73g1 | arginine N-methyltransferase | 0,53232929 | 0,01252195 | 0,42023489 | 0,04501288 |
| C4B63_70g54 | cystathionine beta-synthase | 0,52921401 | 0,04268918 | 0,64174066 | 0,00221407 |
| C4B63_20g219 | kinetoplast-associated protein 3 | 0,52533366 | 0,01598036 | 0,45222937 | 0,04140318 |
| C4B63_48g44 | conserved hypothetical protein | 0,52474805 | 0,0361839 | 0,4827289 | 0,04053149 |
| C4B63_112g71 | cystathionine beta-synthase | 0,52356398 | 0,01214417 | 0,55057426 | 0,00180985 |
| C4B63_4g558 | phosphatidic acid phosphatase protein | 0,52250194 | 0,00025571 | 0,40679963 | 0,03861318 |
| C4B63_28g80 | DnaJ chaperone protein | 0,52099635 | 0,00254618 | 0,41987528 | 0,03782627 |
| C4B63_1g121c | cytochrome b5-like protein | 0,52071391 | 0,00535749 | 0,43253747 | 0,04018165 |
| C4B63_73g16 | conserved hypothetical protein | 0,52033553 | 0,00818288 | 0,45193215 | 0,02478701 |
| C4B63_22g23 | ARM-like helical domain-containing protein | 0,51792457 | 0,00590987 | 0,42782919 | 0,03537106 |
| C4B63_15g14 | oligosaccharyl transferase subunit | 0,51744498 | 0,0224764 | 0,65355214 | 0,0022754 |
| C4B63_38g159 | conserved hypothetical protein | 0,51680753 | 0,00292613 | 0,50827272 | 0,00657051 |
| C4B63_87g17 | pitrilysin-like metalloprotease | 0,51418074 | 0,00039421 | 0,46093435 | 0,00068217 |
| C4B63_21g255 | Nucleolar protein 56 | 0,51408525 | 0,0002601 | 0,44173325 | 0,00575627 |
| C4B63_143g48 | cystathionine beta-synthase | 0,50699345 | 0,04827331 | 0,58638098 | 0,00536874 |
| C4B63_79g37 | methionyl-tRNA synthetase | 0,50697877 | 0,02907851 | 0,45640591 | 0,04680232 |
| C4B63_41g271 | conserved hypothetical protein | 0,50679149 | 0,00238113 | 0,4584076 | 0,00256085 |
| C4B63_3g1116 | Uncharacterized membrane protein C365.16 | 0,50583211 | 0,02116135 | 0,43176812 | 0,04415769 |
| C4B63_10g436 | SPFH domain / Band 7 family | 0,50561827 | 0,02223769 | 0,60518696 | 0,00054743 |
| C4B63_52g54 | conserved hypothetical protein | 0,50458155 | 0,01192978 | 0,50658563 | 0,01636689 |
| C4B63_6g397 | cation transporter | 0,50424596 | 0,00673586 | 0,46859541 | 0,01995044 |
| C4B63_44g114 | conserved hypothetical protein | 0,50376441 | 0,00205669 | 0,4395097 | 0,01491705 |
| C4B63_4g527 | mitochondrial carrier protein | 0,50341476 | 0,02919287 | 0,49071057 | 0,02203029 |
| C4B63_16g315 | mitochondrial DNA primase | 0,5033871 | 0,00377228 | 0,45885987 | 0,01434078 |
| C4B63_31g229 | orotidine-5-phosphate decarboxylase/orotate phospho | 0,5013394 | 0,00214838 | 0,52403937 | 0,00168277 |
| C4B63_81g44 | tRNA (cytosine(34)-C(5))-methyltransferase | 0,50008642 | 0,01261366 | 0,50369468 | 0,01183407 |
| C4B63_143g21 | cystathionine beta-synthase | 0,49995494 | 0,02146504 | 0,58492292 | 0,00089212 |
| C4B63_23g260 | conserved hypothetical protein | 0,49916823 | 0,01520746 | 0,51817964 | 0,00882209 |
| C4B63_57g98 | succinyl-CoA ligase | 0,49365512 | 0,03332186 | 0,52324247 | 0,01991792 |
| C4B63_61g92 | conserved hypothetical protein | 0,49358519 | 0,0229219 | 0,5786633 | 0,00000419 |
| C4B63_87g21 | conserved hypothetical protein | 0,49132026 | 0,01262083 | 0,44376032 | 0,02132118 |
| C4B63_6g113 | ribonucleoside-diphosphate reductase large chain | 0,49029777 | 0,00000209 | 0,44247962 | 0,0000721 |
| C4B63_200g32 | 3,2-trans-enoyl-CoA isomerase, mitochondrial precurs | 0,48762559 | 0,00356342 | 0,42484329 | 0,03940485 |
| C4B63_28g64 | aspartate aminotransferase | 0,486138 | 0,01213982 | 0,46943569 | 0,01476379 |
| C4B63_15g16 | oligosaccharyl transferase subunit | 0,48532529 | 0,04257318 | 0,62755077 | 0,00579132 |
| C4B63_12g235 | protein transport protein Sec31 | 0,4849963 | 0,02085645 | 0,49194733 | 0,02113329 |
| C4B63_183g15 | spermidine synthase | 0,48395941 | 0,00290123 | 0,432845 | 0,0086069 |
| C4B63_25g232 | Tetratricopeptide repeat protein 27 homolog | 0,48372352 | 0,00600191 | 0,43847064 | 0,02995353 |
| C4B63_282g15 | cation transporter | 0,48340099 | 0,00477139 | 0,49285965 | 0,00391881 |
| C4B63_15g12 | conserved hypothetical protein | 0,48333567 | 0,02777324 | 0,49535917 | 0,01970951 |
| C4B63_97g36 | P-type H+-ATPase | 0,4759758 | 0,00159451 | 0,48078805 | 0,0020481 |
| C4B63_104g103 | zinc transporter | 0,47272608 | 0,02515954 | 0,81702449 | 0,0000458 |
| C4B63_97g39 | P-type H+-ATPase | 0,47023749 | 0,00127129 | 0,45453318 | 0,00085777 |
| C4B63_8g62 | Nucleolar protein 10 | 0,46837475 | 0,02491629 | 0,49206505 | 0,0237951 |
| C4B63_61g95 | conserved hypothetical protein | 0,46821428 | 0,03545349 | 0,54246357 | 0,0001405 |
| C4B63_4g478 | DNA-directed RNA polymerases I and III subunit RPA | 0,46638678 | 0,00351702 | 0,42665773 | 0,02880343 |
| C4B63_78g37 | cation-transporting ATPase | 0,45976909 | 0,01806238 | 0,51145767 | 0,00716037 |
| C4B63_41g139 | conserved hypothetical protein | 0,45825912 | 0,03537463 | 0,46076169 | 0,03451891 |
| C4B63_138g7 | unspecified product | 0,45810501 | 0,02623562 | 0,52131362 | 0,0102638 |
| C4B63_25g317 | threonine synthase | 0,45451885 | 0,00525016 | 0,48558048 | 0,00904096 |
| C4B63_98g4 | glucose-6-phosphate 1-dehydrogenase | 0,45384891 | 0,01592939 | 0,42438913 | 0,02708413 |
| C4B63_97g34 | P-type H+-ATPase | 0,44733032 | 0,00418055 | 0,46140309 | 0,00192629 |
| C4B63_25g234 | RNA cytidine acetyltransferase | 0,4445443 | 0,01241995 | 0,46739182 | 0,01298396 |
| C4B63_63g91 | GTPase activating protein | 0,44281184 | 0,01338903 | 0,42515017 | 0,02995353 |
| C4B63_42g179 | ribosome biogenesis regulatory protein (RRS1) | 0,44234566 | 0,01592939 | 0,41775503 | 0,02842148 |
| C4B63_30g129 | rrp44p homologue | 0,43959459 | 0,00418055 | 0,41797321 | 0,00708454 |
| C4B63_25g117 | conserved hypothetical protein | 0,43743262 | 0,02546399 | 0,45470176 | 0,03920482 |
| C4B63_30g208 | C2 domain/Ankyrin repeats (3 copies) | 0,43642244 | 0,00418596 | 0,41773989 | 0,01494838 |
| C4B63_13g173 | beta galactofuranosyl glycosyltransferase | 0,4359951 | 0,00601812 | 0,42762103 | 0,0068588 |
| C4B63_142g6 | GMP synthase | 0,4345848 | 0,00794111 | 0,42432082 | 0,02195102 |
| C4B63_39g342 | phosphoenolpyruvate mutase | 0,42955274 | 0,0222241 | 0,45636507 | 0,01463087 |
| C4B63_31g178 | Zinc phosphodiesterase ELAC protein 2 | 0,42751047 | 0,03315248 | 0,48629507 | 0,02137856 |
| C4B63_34g118 | conserved hypothetical protein | 0,42513133 | 0,03633405 | 0,53965518 | 0,00463506 |
| C4B63_16g293 | inositol 1,4,5-trisphosphate receptor | 0,42481488 | 0,02795553 | 0,52058878 | 0,00541714 |
| C4B63_87g34 | conserved hypothetical protein | 0,42023305 | 0,03545349 | 0,44484174 | 0,04237166 |
| C4B63_194g21 | DNA-directed RNA polymerase III | 0,41644481 | 0,01206038 | 0,43221535 | 0,01322718 |
| C4B63_18g229 | conserved hypothetical protein | 0,41008184 | 0,00268317 | 0,40045274 | 0,0101589 |
| C4B63_322g14 | amino acid transporter | 0,40495851 | 0,01868402 | 0,43292788 | 0,01508389 |
| C4B63_95g32 | conserved hypothetical protein | 0,40407833 | 0,01153498 | 0,44499248 | 0,00579132 |
| C4B63_11g63 | D-alanine--D-alanine ligase | 0,40245483 | 0,023319 | 0,45124729 | 0,01387081 |
| C4B63_12g234 | protein transport protein Sec31 | 0,40195016 | 0,03773879 | 0,51405996 | 0,01201982 |
| C4B63_112g84c | Histone H2B variant V | 0,4005283 | 0,03435513 | 0,41623633 | 0,02810283 |

#### Hemin and Hb common DEGs (t24 vs t0)

##### Downregulated genes

| ID | Annotation | Hemin | | Hb | |
|---|---|---|---|---|---|
| | | log2FC | padj | log2FC | padj |
| C4B63_49g168 | conserved hypothetical protein | -1,4820465 | 4,9246E-06 | -0,9923625 | 0,004767 |
| C4B63_462g9 | glutamamyl carboxypeptidase(fragment) | -1,4450477 | 3,808E-12 | -0,8097054 | 0,00709218 |
| C4B63_431g10 | glutamamyl carboxypeptidase | -0,9316064 | 3,8466E-05 | -0,8216085 | 0,00242018 |
| C4B63_24g313 | protein kinase | -0,9065267 | 0,00310202 | -0,7862406 | 0,03118586 |
| C4B63_113g47 | Generative cell specific 1 protein | -0,8760475 | 0,000173 | -0,7254681 | 0,00603908 |
| C4B63_244g7 | trans-sialidase, Group I | -0,8742289 | 0,00256955 | -0,8005797 | 0,02415316 |
| C4B63_38g182 | UDP-Gal or UDP-GlcNAc-dependent glycosy | -0,8708623 | 0,00012786 | -0,7484477 | 0,00335613 |
| C4B63_46g90 | Ring finger domain containing protein | -0,8505314 | 0,00713798 | -0,7717359 | 0,03505683 |
| C4B63_37g189 | trans-sialidase, Group VI | -0,8424785 | 0,00013523 | -0,942598 | 0,00038415 |
| C4B63_1g941 | trans-sialidase, Group V | -0,832428 | 0,00024837 | -0,7110851 | 0,02539752 |
| C4B63_336g1 | glutamamyl carboxypeptidase(fragment) | -0,9848533 | 1,6036E-05 | -0,692245 | 0,00814607 |
| C4B63_277g11 | glutamamyl carboxypeptidase | -0,9194587 | 0,00025793 | -0,5715871 | 0,03097346 |
| C4B63_18g330 | zinc-finger protein ZFP1 | -0,8838695 | 0,00070028 | -0,6614627 | 0,04688151 |
| C4B63_67g26 | trans-sialidase, Group II | -0,8623343 | 6,2322E-06 | -0,5158722 | 0,03586942 |
| C4B63_38g186 | UDP-Gal or UDP-GlcNAc-dependent glycosy | -0,8553927 | 0,00029553 | -0,637564 | 0,02465589 |
| C4B63_68g79 | surface protease GP63 | -0,841546 | 1,609E-05 | -0,5748173 | 0,01934367 |
| C4B63_30g222 | conserved hypothetical protein | -0,82799 | 5,1856E-10 | -0,5997656 | 8,9473E-05 |
| C4B63_170g1 | conserved hypothetical protein | -0,8080193 | 0,00071953 | -0,597052 | 0,03776192 |
| C4B63_2g309 | NIMA-related kinase | -0,7745527 | 0,00148491 | -0,6132738 | 0,04086727 |
| C4B63_194g17 | phosphate-repressible phosphate permease | -0,7626285 | 0,0014678 | -0,5696705 | 0,01770816 |
| C4B63_21g18 | arginine permease | -0,7457621 | 0,00020112 | -0,6864553 | 0,00311501 |
| C4B63_73g11 | trans-sialidase, Group IV | -0,7361813 | 2,7288E-06 | -0,5749244 | 0,00091143 |
| C4B63_22g322 | phosphate-repressible phosphate permease | -0,722235 | 0,0026325 | -0,5542708 | 0,02234822 |
| C4B63_26g286 | arginine kinase | -0,7173933 | 6,9338E-05 | -0,5553001 | 0,00784888 |
| C4B63_351g3 | calpain-like cysteine peptidase | -0,6667155 | 3,4856E-06 | -0,5673555 | 0,00267818 |
| C4B63_26g282 | arginine kinase | -0,6474766 | 1,481E-05 | -0,4742744 | 0,03471292 |
| C4B63_5g393 | trans-sialidase, Group V | -0,6413421 | 0,00099043 | -0,4921271 | 0,02257767 |
| C4B63_23g195 | 3', 5'-cyclic nucleotide phosphodiesterase | -0,6274947 | 0,00485219 | -0,6173805 | 0,01298472 |
| C4B63_5g381 | mucin TcMUCII | -0,6268937 | 0,01460375 | -0,5796701 | 0,01147814 |
| C4B63_194g16 | phosphate-repressible phosphate permease | -0,6199089 | 0,00275003 | -0,4997368 | 0,02415316 |
| C4B63_11g60 | CHAP domain containing protein | -0,6183338 | 1,71E-05 | -0,4118007 | 0,04309845 |
| C4B63_21g11 | arginine permease | -0,6162036 | 0,00076292 | -0,5559931 | 0,00699831 |
| C4B63_22g321 | phosphate-repressible phosphate permease | -0,6001904 | 0,00278047 | -0,4752943 | 0,01776837 |
| C4B63_194g15 | phosphate-repressible phosphate permease | -0,5975142 | 0,00482966 | -0,4719681 | 0,02792077 |
| C4B63_21g6 | arginine permease | -0,597169 | 0,00356621 | -0,5571367 | 0,01476544 |
| C4B63_21g5 | arginine permease | -0,5917919 | 0,00230408 | -0,5978739 | 0,00450003 |
| C4B63_3g1092 | alanine racemase | -0,5909575 | 0,00035992 | -0,4353996 | 0,01608889 |
| C4B63_13g146 | conserved hypothetical protein | -0,5899994 | 0,00356034 | -0,5558323 | 0,0217109 |
| C4B63_14g28 | hexose transporter | -0,5820395 | 0,00295405 | -0,4670699 | 0,04606434 |
| C4B63_14g30 | hexose transporter | -0,5588023 | 0,00393427 | -0,4545523 | 0,04795767 |
| C4B63_23g226 | conserved hypothetical protein | -0,5481731 | 0,01108199 | -0,4739495 | 0,03657006 |
| C4B63_1g480 | trans-sialidase, Group V | -0,5412182 | 0,0094045 | -0,4664758 | 0,04118501 |
| C4B63_9g387 | conserved hypothetical protein | -0,541077 | 0,00387348 | -0,4137201 | 0,04077069 |
| C4B63_14g5 | hexose transporter | -0,5385089 | 0,00208192 | -0,4467362 | 0,03423827 |
| C4B63_10g157 | Double RNA binding domain protein 6A | -0,5343551 | 0,00025745 | -0,4387388 | 0,02172544 |
| C4B63_44g230 | amino acid transporter | -0,5337096 | 1,494E-05 | -0,4214039 | 0,00807509 |
| C4B63_21g23 | arginine permease | -0,5317464 | 0,0376242 | -0,6165957 | 0,01934367 |
| C4B63_21g27 | arginine permease(fragment) | -0,5288013 | 0,04083064 | -0,5486328 | 0,0449145 |
| C4B63_1g473 | trans-sialidase | -0,5252563 | 0,02813247 | -0,5418481 | 0,0302265 |
| C4B63_2g73 | syntaxin12 (t-snare) | -0,5192405 | 0,03430865 | -0,4861094 | 0,03377429 |
| C4B63_155g17 | trans-sialidase, Group II | -0,5089716 | 0,03567896 | -0,5920842 | 0,01848201 |
| C4B63_3g1089 | Alpha-ketoglutarate-dependent dioxygenase | -0,5059747 | 0,04522976 | -0,5792047 | 0,03355794 |
| C4B63_134g49 | trans-sialidase, Group V | -0,4842843 | 0,04960968 | -0,465359 | 0,0315107 |
| C4B63_27g1015c | conserved hypothetical protein | -0,4352029 | 0,04065166 | -0,5676481 | 0,00273583 |
| C4B63_70g65 | conserved hypothetical protein | -0,4307023 | 0,0316066 | -0,4449862 | 0,03763091 |
| C4B63_13g317 | vesicle-associated membrane protein | -0,4148283 | 0,02755419 | -0,4306619 | 0,04102151 |
| C4B63_19g130 | Dual specificity phosphatase, catalytic doma | -0,4107456 | 0,03180994 | -0,4871743 | 0,02762162 |
| C4B63_16g183 | surface protease GP63 | -0,6983272 | 0,00181378 | -0,6148869 | 0,03082793 |
| C4B63_4g180 | serine/threonine protein kinase | -0,6958293 | 5,3112E-08 | -0,4340573 | 0,00606314 |
| C4B63_109g42 | conserved hypothetical protein | -0,6932399 | 0,00021605 | -0,447963 | 0,02651241 |
| C4B63_21g16 | arginine permease | -0,6892532 | 0,00051401 | -0,5702117 | 0,00915015 |
| C4B63_26g278 | arginine kinase | -0,6762975 | 0,00018654 | -0,4819603 | 0,0124197 |
| C4B63_59g207 | mucin TcMUCII | -0,6683352 | 0,00560751 | -0,6326276 | 0,04795767 |
| C4B63_34g137 | trans-sialidase, Group IV | -0,6674387 | 0,00457838 | -0,5151104 | 0,02247693 |

##### Upregulated genes

| ID | Annotation | Hemin | | Hb | |
|---|---|---|---|---|---|
| | | log2FC | padj | log2FC | padj |
| C4B63_86g57 | conserved hypothetical protein | 0,84682464 | 0,00112086 | 0,67742002 | 0,01928453 |
| C4B63_128g306c | histone H4 | 0,84021584 | 0,00104004 | 0,57959669 | 0,0315107 |
| C4B63_47g72 | enolase | 0,83629988 | 1,9314E-09 | 0,43897637 | 0,01564897 |
| C4B63_128g303c | histone H4 | 0,83380038 | 2,8641E-05 | 0,61211345 | 0,01164414 |
| C4B63_35g168 | 3-Beta-hydroxysteroid-delta(8), delta(7)-isome | 0,83194139 | 0,00045782 | 0,6294171 | 0,03291456 |
| C4B63_188g139c | histone H4 | 0,8075546 | 0,00025053 | 0,53118865 | 0,01518836 |
| C4B63_188g158c | histone H4 | 0,79790263 | 0,00194645 | 0,6085607 | 0,01383811 |
| C4B63_42g103 | pyridoxal kinase | 0,79473904 | 2,7434E-11 | 0,54221352 | 0,00333084 |
| C4B63_15g12 | conserved hypothetical protein | 0,75667068 | 7,3023E-05 | 0,55439804 | 0,01116528 |
| C4B63_188g156c | histone H4 | 0,74017191 | 0,00020369 | 0,58455985 | 0,00715299 |
| C4B63_34g287 | glycerate kinase | 0,73841352 | 0,00104996 | 0,61826589 | 0,01098709 |
| C4B63_6g309 | heat shock protein 70 (hsp70) | 0,73135285 | 0,00108363 | 0,6126736 | 0,03256411 |
| C4B63_128g304c | histone H4 | 0,72492085 | 0,00633283 | 0,57295188 | 0,02098887 |
| C4B63_81g18 | chaperonin HSP60, mitochondrial precursor(fr | 0,72158957 | 0,00354295 | 0,70216073 | 0,00814607 |
| C4B63_48g92 | chaperonin HSP60, mitochondrial precursor | 0,72124751 | 0,00789907 | 0,69847339 | 0,01656392 |
| C4B63_78g23 | nucleolar RNA-binding protein | 0,71814994 | 0,00497453 | 0,62941545 | 0,0201758 |
| C4B63_113g27 | heat shock protein 85(fragment) | 0,71402098 | 2,1268E-05 | 0,55311182 | 0,0119577 |
| C4B63_24g243 | conserved hypothetical protein | 0,69934733 | 0,00154859 | 0,59761924 | 0,01838273 |
| C4B63_116g15 | membrane transporter protein | 0,69852597 | 1,8805E-05 | 0,41231402 | 0,02575516 |
| C4B63_81g100 | chaperonin HSP60, mitochondrial precursor | 0,69135537 | 0,01642763 | 0,72875163 | 0,01442013 |
| C4B63_303g8 | unspecified product | 0,68696833 | 0,02441798 | 0,68142274 | 0,01319114 |
| C4B63_34g289 | glycerate kinase | 0,68504229 | 0,00547084 | 0,59465911 | 0,02903767 |
| C4B63_4g383 | mevalonate-diphosphate decarboxylase | 0,68396111 | 0,00054761 | 0,53200734 | 0,02568702 |
| C4B63_133g60 | heat shock protein 70 (hsp70) | 0,67958987 | 0,00043907 | 0,58015773 | 0,00949918 |
| C4B63_80g17 | ATP binding protein-like protein | 0,67908623 | 0,00642933 | 0,52706602 | 0,03047809 |
| C4B63_48g86 | chaperonin HSP60, mitochondrial precursor | 0,67813 | 0,00731075 | 0,66772817 | 0,01250366 |
| C4B63_302g50c | histone H3 | 0,67486621 | 0,00767506 | 0,61154645 | 0,01814599 |
| C4B63_227g7 | transporter | 0,67465931 | 1,298E-06 | 0,4524177 | 0,01070649 |
| C4B63_137g40 | heat shock protein 70 (hsp70)(fragment) | 0,66519229 | 0,00145635 | 0,6455262 | 0,00412809 |
| C4B63_111g29 | conserved hypothetical protein | 0,66319046 | 0,00116542 | 0,54210318 | 0,01449531 |
| C4B63_102g44 | hypoxanthine-guanine phosphoribosyltransfera | 0,66193172 | 0,00103229 | 0,65406208 | 0,00860162 |
| C4B63_48g88 | chaperonin HSP60, mitochondrial precursor | 0,65091831 | 0,0103198 | 0,63499458 | 0,02037429 |
| C4B63_200g32 | 3,2-trans-enoyl-CoA isomerase, mitochondrial | 0,65087413 | 2,0771E-05 | 0,47456201 | 0,02642601 |
| C4B63_81g102 | chaperonin HSP60, mitochondrial precursor | 0,64048212 | 0,00625541 | 0,64221835 | 0,01025684 |
| C4B63_43g125 | centromere/microtubule binding protein cbf5 | 0,63922269 | 0,00138183 | 0,48585402 | 0,02969068 |
| C4B63_137g38 | heat shock protein 70 (hsp70)(fragment) | 0,63233175 | 0,0008588 | 0,58577639 | 0,00707817 |
| C4B63_55g66 | Fatty-acid amide hydrolase 2 | 0,62737586 | 1,8473E-06 | 0,42269866 | 0,01389839 |
| C4B63_44g246 | nucleolar protein | 0,62671318 | 0,0101438 | 0,56627159 | 0,02762162 |
| C4B63_48g90 | chaperonin HSP60, mitochondrial precursor | 0,62478258 | 0,01826435 | 0,6124909 | 0,02969068 |
| C4B63_184g39 | heat shock protein DNAJ(fragment) | 0,62451893 | 0,00049085 | 0,58217227 | 0,01105635 |
| C4B63_7g313 | amidinotransferase | 0,62214283 | 0,00024549 | 0,48466031 | 0,0221592 |
| C4B63_302g22 | histone H3 | 0,61772611 | 0,01155795 | 0,63339896 | 0,00311501 |
| C4B63_137g39 | heat shock protein 70 (hsp70) | 0,61488824 | 0,0014939 | 0,55438106 | 0,01107133 |
| C4B63_137g32 | conserved hypothetical protein | 0,61470577 | 0,00335476 | 0,54866181 | 0,01959927 |
| C4B63_35g189 | eukaryotic translation initiation factor 1A | 0,61364643 | 0,0003331 | 0,4430004 | 0,03134249 |
| C4B63_81g103 | chaperonin HSP60, mitochondrial precursor | 0,61341298 | 0,01035614 | 0,61879589 | 0,01683944 |
| C4B63_10g436 | SPFH domain / Band 7 family | 0,61285198 | 0,00282871 | 0,42409363 | 0,04043512 |
| C4B63_70g93 | DNA repair and recombination helicase protei | 0,61171389 | 0,00118684 | 0,6386604 | 0,00236946 |
| C4B63_207g9 | ribosome biogenesis protein | 0,60513568 | 0,00265919 | 0,50986388 | 0,03532622 |
| C4B63_48g91 | chaperonin HSP60, mitochondrial precursor | 0,60505214 | 0,00993604 | 0,62691435 | 0,01382321 |
| C4B63_61g149 | eukaryotic translation initiation factor 3 subuni | 0,60482749 | 0,00260084 | 0,51263869 | 0,02179193 |
| C4B63_45g239 | lysyl-tRNA synthetase | 0,60175777 | 0,03080979 | 0,60290945 | 0,03924539 |
| C4B63_79g21 | ABC transporter | 0,60157753 | 0,00177924 | 0,46027655 | 0,0295281 |
| C4B63_15g15 | oligosaccharyl transferase subunit | 0,60090255 | 0,00713798 | 0,59013954 | 0,01678208 |
| C4B63_57g88 | fibrillarin | 0,59906387 | 0,00022858 | 0,4694768 | 0,01515891 |
| C4B63_28g56 | TFIIF-stimulated CTD phosphatase | 0,59831342 | 0,00112428 | 0,60737358 | 0,00040229 |
| C4B63_48g89 | chaperonin HSP60, mitochondrial precursor | 0,59760154 | 0,01248609 | 0,66808236 | 0,00603908 |
| C4B63_2g430 | heat shock protein | 0,59731084 | 0,01241885 | 0,55770794 | 0,02308202 |
| C4B63_81g106 | chaperonin HSP60, mitochondrial precursor | 0,5928271 | 0,01192435 | 0,62726577 | 0,01333415 |
| C4B63_6g314 | heat shock protein 70 (hsp70) | 0,59145697 | 0,00734902 | 0,60089417 | 0,00475061 |
| C4B63_47g73 | importin beta-1 subunit | 0,5911002 | 0,00031888 | 0,49081484 | 0,00764991 |
| C4B63_4g79 | ribosome biogenesis protein | 0,58684392 | 0,00319295 | 0,5831513 | 0,00357173 |
| C4B63_45g126 | RNA recognition motif (a.k.a. RRM, RBD, or R | 0,58634314 | 0,03361262 | 0,55689867 | 0,03637643 |
| C4B63_6g316 | heat shock protein 70 (hsp70) | 0,5842611 | 0,00282871 | 0,52708884 | 0,01403013 |
| C4B63_42g52 | mitochondrial oligo U binding protein TBRGG1 | 0,58382582 | 0,00656178 | 0,54900829 | 0,02096594 |
| C4B63_137g36 | heat shock protein 70 (hsp70)(fragment) | 0,58382505 | 0,0018685 | 0,54379795 | 0,01574229 |
| C4B63_47g52 | deoxyhypusine synthase | 0,5814406 | 0,00767506 | 0,57918062 | 0,01043056 |
| C4B63_6g550 | NAD(P)-dependent steroid dehydrogenase pro | 0,58124862 | 0,00032466 | 0,41349721 | 0,03811443 |
| C4B63_4g454 | nucleosome assembly protein | 0,57972879 | 0,00159965 | 0,46819451 | 0,03343498 |
| C4B63_23g116 | Importin beta-like SAD2 | 0,57843616 | 0,00187284 | 0,47054967 | 0,02831737 |
| C4B63_6g236 | cation transporter | 0,5782527 | 0,00066914 | 0,48306605 | 0,01095188 |
| C4B63_263g7 | ruvB-like DNA helicase | 0,57576938 | 5,8478E-05 | 0,44953968 | 0,01127586 |
| C4B63_80g21 | Nucleoside diphosphate kinase | 0,57321125 | 0,00133996 | 0,47089207 | 0,03337729 |
| C4B63_6g306 | heat shock protein 70 (hsp70)(fragment) | 0,56737041 | 0,00284087 | 0,55186797 | 0,0051483 |
| C4B63_255g21 | calreticulin | 0,56621925 | 0,02058724 | 0,98140098 | 0,00034384 |
| C4B63_40g87 | WD40 repeat-containing protein | 0,56609794 | 0,0213207 | 0,56532353 | 0,01807531 |
| C4B63_98g4 | glucose-6-phosphate 1-dehydrogenase | 0,56183659 | 0,00130596 | 0,41479165 | 0,04309845 |
| C4B63_6g530 | conserved hypothetical protein | 0,55908969 | 0,0040822 | 0,42137761 | 0,03134249 |
| C4B63_114g17 | conserved hypothetical protein | 0,55832666 | 0,003484 | 0,49596946 | 0,02689726 |
| C4B63_11g9 | eukaryotic translation initiation factor 3 subuni | 0,5533089 | 1,6929E-05 | 0,42048176 | 0,02533924 |
| C4B63_104g78 | calreticulin | 0,55175309 | 0,00734902 | 0,83196756 | 0,00307421 |
| C4B63_31g181 | mitochondrial DNA polymerase beta | 0,54485447 | 0,04689631 | 0,63802442 | 0,01867475 |
| C4B63_6g317 | heat shock protein 70 (hsp70) | 0,54326611 | 0,00184098 | 0,50085813 | 0,01070649 |
| C4B63_27g204 | kinetoplastid kinetochore protein 10 | 0,54090836 | 0,00016826 | 0,52543146 | 0,00292964 |
| C4B63_7g212 | ruvB-like DNA helicase | 0,54067799 | 0,000167 | 0,43472643 | 0,01002591 |
| C4B63_18g288 | heat shock 70 kDa protein, mitochondrial prec | 0,53863403 | 0,00845981 | 0,49342213 | 0,02941505 |
| C4B63_26g252 | nucleosome assembly protein-like protein | 0,53845026 | 0,00967836 | 0,59548368 | 0,01599704 |
| C4B63_304g13 | nucleosome assembly protein | 0,53480676 | 0,0065431 | 0,48422879 | 0,03118304 |
| C4B63_20g137 | leucine-rich repeat protein (LRRP) | 0,53409499 | 0,00440613 | 0,51440489 | 0,01147814 |
| C4B63_2g786 | ubiquitin-activating enzyme E1 | 0,53312506 | 0,00591041 | 0,49802033 | 0,01160387 |
| C4B63_15g16 | oligosaccharyl transferase subunit | 0,53215426 | 0,02058034 | 0,56243101 | 0,0225059 |
| C4B63_81g17 | DNA mismatch repair protein MSH2(fragment) | 0,5310749 | 0,00239454 | 0,43888977 | 0,02603642 |
| C4B63_25g313 | Putrescine-cadaverine transporter | 0,52971223 | 0,02869056 | 0,7791029 | 0,02392037 |
| C4B63_97g36 | P-type H+-ATPase | 0,52842661 | 0,00026321 | 0,47083401 | 0,00423688 |
| C4B63_73g16 | conserved hypothetical protein | 0,52838614 | 0,00610066 | 0,5074506 | 0,01317969 |
| C4B63_25g234 | RNA cytidine acetyltransferase | 0,52402265 | 0,00167781 | 0,43301553 | 0,03274585 |
| C4B63_1g57 | AAA ATPase | 0,52226581 | 0,00579738 | 0,43032324 | 0,04606434 |
| C4B63_297g9 | conserved hypothetical protein | 0,52139197 | 0,00272641 | 0,46829146 | 0,00851324 |
| C4B63_207g7 | conserved hypothetical protein | 0,51678979 | 0,0003331 | 0,41562229 | 0,01382321 |
| C4B63_8g61 | heat shock protein 70 (hsp70) | 0,51634374 | 0,01104523 | 0,64273822 | 0,00835562 |
| C4B63_9g201 | GTP-binding protein | 0,51518441 | 0,02510703 | 0,47296577 | 0,02987447 |
| C4B63_97g34 | P-type H+-ATPase | 0,51298784 | 0,00054761 | 0,44537571 | 0,00490735 |
| C4B63_7g209 | Ribonuclease P protein subunit p25-like protei | 0,51284848 | 0,00068153 | 0,7243265 | 0,00013651 |
| C4B63_25g192 | NAA35 | 0,50587109 | 0,00048238 | 0,52218014 | 0,00102429 |
| C4B63_396g14 | Present in the outer mitochondrial membrane | 0,50387812 | 0,00807801 | 0,6263755 | 0,00268131 |
| C4B63_9g396 | sedoheptulose-1,7-bisphosphatase | 0,49918438 | 0,01016985 | 0,48049656 | 0,03374928 |
| C4B63_19g200 | conserved hypothetical protein | 0,49825817 | 0,0006872 | 0,45855259 | 0,01608889 |
| C4B63_23g110 | UDP-glucose:glycoprotein glucosyltransferase | 0,49607403 | 0,00691897 | 0,63163925 | 0,00098442 |
| C4B63_113g39 | MRB1-associated protein | 0,49344124 | 0,04595972 | 0,65378096 | 0,03586942 |
| C4B63_2g394 | kinesin | 0,49324471 | 0,04382442 | 0,5927005 | 0,01683944 |
| C4B63_2g209 | Present in the outer mitochondrial membrane | 0,49252957 | 0,02586587 | 0,62064137 | 0,00465422 |
| C4B63_102g79 | conserved hypothetical protein | 0,48977455 | 0,00286164 | 0,44523118 | 0,01476544 |
| C4B63_20g19 | Nucleoporin | 0,48938366 | 0,00447898 | 0,51095361 | 0,00539407 |
| C4B63_7g347 | V-type ATPase, A subunit | 0,48758259 | 0,0168315 | 0,49913714 | 0,02371205 |
| C4B63_97g19 | pyruvate dehydrogenase E1 component alpha | 0,48428019 | 0,01842826 | 0,5224913 | 0,01382321 |
| C4B63_17g64 | conserved hypothetical protein | 0,48134186 | 0,00164001 | 0,4409603 | 0,04795767 |
| C4B63_63g7 | eukaryotic translation initiation factor 3 subuni | 0,47951804 | 0,00893282 | 0,45437061 | 0,00980138 |
| C4B63_148g30 | conserved hypothetical protein | 0,47066775 | 0,04757371 | 0,52811726 | 0,04368362 |
| C4B63_43g1 | kinesin | 0,46654911 | 0,03002361 | 0,54668294 | 0,04032818 |
| C4B63_22g18 | conserved hypothetical protein | 0,46618146 | 0,00826681 | 0,41109959 | 0,03811443 |
| C4B63_7g410 | conserved hypothetical protein | 0,46590124 | 0,00180754 | 0,42621902 | 0,00530589 |
| C4B63_23g109 | UDP-glucose:glycoprotein glucosyltransferase | 0,46289862 | 0,00087483 | 0,58826518 | 0,00131096 |
| C4B63_9g246 | FtsJ cell division protein | 0,46135062 | 0,00556788 | 0,45589766 | 0,01500857 |
| C4B63_11g74 | pyruvate dehydrogenase E1 beta subunit | 0,46106904 | 0,00770536 | 0,43609408 | 0,0302265 |
| C4B63_220g12 | peroxidoxin | 0,46080869 | 0,00150125 | 0,46963438 | 0,00131096 |
| C4B63_16g98 | conserved hypothetical protein | 0,45908295 | 0,0001834 | 0,42678657 | 0,00210526 |
| C4B63_21g80 | conserved hypothetical protein | 0,45719755 | 0,00039076 | 0,40287295 | 0,00344409 |
| C4B63_43g161 | ATP-dependent RNA helicase SUB2 | 0,45623742 | 0,00211271 | 0,53157775 | 0,00412809 |
| C4B63_78g28 | RNA recognition motif (a.k.a. RRM, RBD, or R | 0,45603759 | 0,00012786 | 0,4820143 | 0,00359377 |
| C4B63_142g6 | GMP synthase | 0,45527172 | 0,00412689 | 0,40518391 | 0,04233645 |
| C4B63_70g112 | pumilio-repeat, RNA-binding protein(fragment) | 0,45475765 | 0,01651673 | 0,48200361 | 0,02906063 |
| C4B63_26g283 | ABC transporter | 0,45366683 | 0,00207594 | 0,40904223 | 0,0217109 |
| C4B63_80g38 | conserved hypothetical protein | 0,45341322 | 0,02362519 | 0,40956287 | 0,04105754 |
| C4B63_10g439 | heat shock protein 90 | 0,44843282 | 0,00770536 | 0,64168709 | 0,00015844 |
| C4B63_183g14 | conserved hypothetical protein | 0,4463395 | 0,00551154 | 0,40126005 | 0,04246147 |
| C4B63_30g177 | condensin complex subunit 2 | 0,44389126 | 0,02337148 | 0,62681857 | 0,0049972 |
| C4B63_80g37 | conserved hypothetical protein | 0,44281318 | 0,03629951 | 0,49304175 | 0,03256411 |
| C4B63_76g58 | Tripartite attachment complex protein 102 | 0,44020909 | 0,01599337 | 0,40354579 | 0,02263471 |
| C4B63_63g145 | retrotransposon hot spot (RHS) protein | 0,4389956 | 0,03002361 | 0,47926528 | 0,01787311 |
| C4B63_149g19 | peroxidoxin | 0,4369691 | 0,00494596 | 0,40661044 | 0,00814871 |
| C4B63_23g212 | serine/threonine-protein kinase | 0,43565473 | 0,00286164 | 0,48098168 | 0,00780624 |
| C4B63_76g32 | guide RNA associated protein, GAP2 | 0,43474393 | 0,00130124 | 0,4096014 | 0,00771335 |
| C4B63_61g131 | protein kinase | 0,43046415 | 0,00408025 | 0,48861544 | 0,02182967 |
| C4B63_2g174 | polyadenylate-binding protein 1 | 0,42687384 | 0,03877962 | 0,61903636 | 0,01328588 |
| C4B63_48g168 | AAA ATPase | 0,42526537 | 0,03466712 | 0,50894465 | 0,01176158 |
| C4B63_24g172 | Uncharacterized protein At3g06530 | 0,41656066 | 0,01864706 | 0,4421163 | 0,0066879 |
| C4B63_48g134 | clathrin heavy chain | 0,4158769 | 0,02692887 | 0,50782765 | 0,0205812 |
| C4B63_76g57 | Tripartite attachment complex protein 102 | 0,41485808 | 0,01554026 | 0,42927216 | 0,00599651 |
| C4B63_137g10 | Coatomer subunit gamma | 0,4012839 | 0,03991797 | 0,41620521 | 0,04417031 |

**Gene Ontology analysis of Hemin and Hb common DEGs (Downregulated, t4 vs t0)**

| ID | Name | Bgd count | Result count | Result gene list | Pct of bgd | Fold enrichment | Odds ratio | P-value | Benjamini | Bonferroni |
|---|---|---|---|---|---|---|---|---|---|---|
| | **BIOLOGICAL PROCESS** | | | | | | | | | |
| GO:0006522 | alanine metabolic process | 2 | 2 | C4B63_3g1092,C4 | 100.0 | 115.77 | inf | 7,36E-05 | 0.00537366517879 | 0.0107473303576 |
| GO:0009078 | pyruvate family amino acid metabolic process | 2 | 2 | C4B63_3g1092,C4 | 100.0 | 115.77 | inf | 7,36E-05 | 0.00537366517879 | 0.0107473303576 |
| | **MOLECULAR FUNCTION** | | | | | | | | | |
| ID | Name | Bgd count | Result count | Result gene list | Pct of bgd | Fold enrichment | Odds ratio | P-value | Benjamini | Bonferroni |
| GO:0008784 | alanine racemase activity | 2 | 2 | C4B63_3g1092,C4 | 100.0 | 115.77 | inf | 7,36E-05 | 0.00191390814587 | 0.00765563258349 |
| GO:0009055 | electron transfer activity | 22 | 3 | C4B63_19g141,C4 | 13.6 | 15.79 | 18.85 | 0.000846756 | 0.0176125297502 | 0.088062648751 |

**Gene Ontology analysis of Hemin and Hb common DEGs (Upregulated, t4 vs t0)**

| ID | Name | Bgd count | Result count | Result gene list | Pct of bgd | Fold enrichment | Odds ratio | P-value | Benjamini | Bonferroni |
|---|---|---|---|---|---|---|---|---|---|---|
| | **BIOLOGICAL PROCESS** | | | | | | | | | |
| GO:0006563 | L-serine metabolic process | 23 | 18 | C4B63_112g68,C4 | 78.3 | 55.87 | 297.95 | 3,64E-30 | 2,20E-28 | 1,10E-27 |
| GO:0006534 | cysteine metabolic process | 23 | 18 | C4B63_112g68,C4 | 78.3 | 55.87 | 297.95 | 3,64E-30 | 2,20E-28 | 1,10E-27 |
| GO:0000096 | sulfur amino acid metabolic process | 26 | 18 | C4B63_112g68,C4 | 69.2 | 49.42 | 186.15 | 1,64E-28 | 7,05E-27 | 4,94E-26 |
| GO:0042254 | ribosome biogenesis | 76 | 8 | C4B63_207g2,C4E | 10.5 | 7.51 | 8.8 | 9,98E-06 | 7,54E-05 | 0.00301466282566 |
| GO:0006177 | GMP biosynthetic process | 6 | 3 | C4B63_142g6,C4E | 50.0 | 35.7 | 72.17 | 5,20E-05 | 0.000356820362851 | 0.0157000959654 |
| GO:0008202 | steroid metabolic process | 18 | 4 | C4B63_227g9,C4E | 22.2 | 15.86 | 20.77 | 9,63E-05 | 0.000645962075026 | 0.0290682933762 |
| GO:0034654 | nucleobase-containing compound biosynthetic proce | 173 | 10 | C4B63_134g37,C4 | 5.8 | 4.13 | 4.62 | 0.000148645 | 0.000919140810392 | 0.0448909849601 |
| GO:0022618 | ribonucleoprotein complex assembly | 20 | 4 | C4B63_34g118,C4 | 20.0 | 14.28 | 18.17 | 0.000149132 | 0.000919140810392 | 0.0450378997092 |
| GO:0055085 | transmembrane transport | 256 | 12 | C4B63_104g103,C | 4.7 | 3.35 | 3.74 | 0.000235169 | 0.0013152092505 | 0.0710212995268 |
| GO:0019856 | pyrimidine nucleobase biosynthetic process | 10 | 3 | C4B63_134g37,C4 | 30.0 | 21.42 | 30.92 | 0.000299366 | 0.0015587718904 | 0.090408769643 |
| | **CELLULAR COMPONENT** | | | | | | | | | |
| ID | Name | Bgd count | Result count | Result gene list | Pct of bgd | Fold enrichment | Odds ratio | P-value | Benjamini | Bonferroni |
| GO:0005730 | nucleolus | 66 | 6 | C4B63_21g255,C4 | 9.1 | 6.49 | 7.36 | 0.000305012 | 0.00899786413862 | 0.0179957282772 |
| | **MOLECULAR FUNCTION** | | | | | | | | | |
| ID | Name | Bgd count | Result count | Result gene list | Pct of bgd | Fold enrichment | Odds ratio | P-value | Benjamini | Bonferroni |
| GO:0016616 | oxidoreductase activity, acting on the CH-OH group c | 36 | 6 | C4B63_157g32,C4 | 16.7 | 11.9 | 14.77 | 9,21E-06 | 0.00066865739191 | 0.00173978052279 |
| GO:0045182 | translation regulator activity | 128 | 9 | C4B63_11g83,C4E | 7.0 | 5.02 | 5.67 | 7,24E-05 | 0.00136834043967 | 0.0136834043967 |
| GO:0003723 | RNA binding | 425 | 17 | C4B63_11g83,C4E | 4.0 | 2.86 | 3.25 | 8,19E-05 | 0.00140708439298 | 0.0154779283228 |
| GO:0016879 | ligase activity, forming carbon-nitrogen bonds | 21 | 4 | C4B63_11g63,C4E | 19.0 | 13.6 | 17.1 | 0.000182237 | 0.00264945063734 | 0.0344428582854 |
| GO:0003746 | translation elongation factor activity | 65 | 6 | C4B63_13g152,C4 | 9.2 | 6.59 | 7.48 | 0.000280455 | 0.00333281700779 | 0.0530060486611 |
| GO:0016831 | carboxy-lyase activity | 24 | 4 | C4B63_134g37,C4 | 16.7 | 11.9 | 14.53 | 0.000313214 | 0.00348220440444 | 0.0591974748755 |

**Gene Ontology analysis of Hemin and Hb common DEGs (Downregulated, t24 vs t0)**

| ID | Name | Bgd count | Result count | Result gene list | Pct of bgd | Fold enrichment | Odds ratio | P-value | Benjamini | Bonferroni |
|---|---|---|---|---|---|---|---|---|---|---|
| | **BIOLOGICAL PROCESS** | | | | | | | | | |
| GO:0006817 | phosphate ion transport | 7 | 5 | C4B63_194g15,C4 | 71.4 | 113.32 | 434.23 | 1,71E-10 | 2,00E-08 | 2,00E-08 |
| GO:0006575 | cellular modified amino acid metabolic process | 25 | 3 | C4B63_26g278,C4 | 12.0 | 19.04 | 22.7 | 0.000493618 | 0.00641703521128 | 0.0577533169015 |
| | **CELLULAR COMPONENT** | | | | | | | | | |
| ID | Name | Bgd count | Result count | Result gene list | Pct of bgd | Fold enrichment | Odds ratio | P-value | Benjamini | Bonferroni |
| GO:0016020 | membrane | 2776 | 34 | C4B63_109g42,C4 | 1.2 | 1.94 | 3.58 | 3,71E-06 | 0.000100137408347 | 0.000100137408347 |
| | **MOLECULAR FUNCTION** | | | | | | | | | |
| ID | Name | Bgd count | Result count | Result gene list | Pct of bgd | Fold enrichment | Odds ratio | P-value | Benjamini | Bonferroni |
| GO:0016775 | phosphotransferase activity, nitrogenous group as ac | 3 | 3 | C4B63_26g278,C4 | 100.0 | 158.65 | inf | 2,37E-07 | 4,08E-06 | 1,63E-05 |
| GO:0005215 | transporter activity | 295 | 8 | C4B63_14g28,C4E | 2.7 | 4.3 | 4.98 | 0.000470774 | 0.00406042666327 | 0.0324834133061 |

**Gene Ontology analysis of Hemin and Hb common DEGs (Upregulated, t24 vs t0)**

| ID | Name | Bgd count | Result count | Result gene list | Pct of bgd | Fold enrichment | Odds ratio | P-value | Benjamini | Bonferroni |
|---|---|---|---|---|---|---|---|---|---|---|
| | **BIOLOGICAL PROCESS** | | | | | | | | | |
| GO:0006457 | protein folding | 91 | 16 | C4B63_104g78,C4 | 17.6 | 12.55 | 17.17 | 8,57E-14 | 2,70E-11 | 2,70E-11 |
| GO:0006084 | acetyl-CoA metabolic process | 6 | 3 | C4B63_11g74,C4E | 50.0 | 35.7 | 72.17 | 5,20E-05 | 0.00456985737375 | 0.016375927911 |
| GO:0006323 | DNA packaging | 17 | 4 | C4B63_26g252,C4 | 23.5 | 16.8 | 22.37 | 7,57E-05 | 0.00476780231525 | 0.0238390115762 |
| | **CELLULAR COMPONENT** | | | | | | | | | |
| ID | Name | Bgd count | Result count | Result gene list | Pct of bgd | Fold enrichment | Odds ratio | P-value | Benjamini | Bonferroni |
| GO:0043231 | intracellular membrane-bounded organelle | 841 | 28 | C4B63_104g78,C4 | 3.3 | 2.38 | 2.86 | 9,94E-06 | 0.000216789325111 | 0.00108394662556 |
| GO:0005694 | chromosome | 168 | 11 | C4B63_128g303c, | 6.5 | 4.67 | 5.33 | 2,18E-05 | 0.000395237711443 | 0.00237142626866 |
| GO:0005759 | mitochondrial matrix | 27 | 5 | C4B63_48g86,C4E | 18.5 | 13.22 | 16.65 | 3,13E-05 | 0.00048718760019 | 0.00341031320133 |
| GO:0005634 | nucleus | 424 | 17 | C4B63_128g303c, | 4.0 | 2.86 | 3.26 | 7,95E-05 | 0.000867058068221 | 0.00867058068221 |
| GO:0032991 | protein-containing complex | 630 | 21 | C4B63_11g9,C4B( | 3.3 | 2.38 | 2.73 | 0.000155310 | 0.00153898491766 | 0.0169288340942 |
| GO:1902493 | acetyltransferase complex | 10 | 3 | C4B63_25g192,C4 | 30.0 | 21.42 | 30.92 | 0.000299366 | 0.00191946939834 | 0.0326309797718 |
| GO:0005852 | eukaryotic translation initiation factor 3 complex | 10 | 3 | C4B63_11g9,C4B( | 30.0 | 21.42 | 30.92 | 0.000299366 | 0.00191946939834 | 0.0326309797718 |
| | **MOLECULAR FUNCTION** | | | | | | | | | |
| ID | Name | Bgd count | Result count | Result gene list | Pct of bgd | Fold enrichment | Odds ratio | P-value | Benjamini | Bonferroni |
| GO:1901363 | heterocyclic compound binding | 2258 | 74 | C4B63_10g439,C4 | 3.3 | 2.34 | 4.61 | 2,95E-16 | 2,60E-14 | 5,19E-14 |
| GO:0000166 | nucleotide binding | 1411 | 49 | C4B63_10g439,C4 | 3.5 | 2.48 | 3.59 | 1,46E-10 | 2,14E-09 | 2,56E-08 |
| GO:0043167 | ion binding | 2086 | 55 | C4B63_104g78,C4 | 2.6 | 1.88 | 2.67 | 1,96E-07 | 2,15E-06 | 3,44E-05 |
| GO:0051082 | unfolded protein binding | 56 | 7 | C4B63_104g78,C4 | 12.5 | 8.92 | 10.62 | 1,17E-05 | 0.000120729191527 | 0.00205239625596 |
| GO:0003676 | nucleic acid binding | 987 | 28 | C4B63_113g39,C4 | 2.8 | 2.03 | 2.38 | 0.000178162 | 0.00142699613415 | 0.0313566751858 |
| GO:0004738 | pyruvate dehydrogenase activity | 2 | 2 | C4B63_11g74,C4E | 100.0 | 71.39 | inf | 0.000194590 | 0.00142699613415 | 0.0342479072195 |
| GO:0046527 | glucosyltransferase activity | 2 | 2 | C4B63_23g109,C4 | 100.0 | 71.39 | inf | 0.000194590 | 0.00142699613415 | 0.0342479072195 |

### Table S4

**Hemin exclusive DEGs (t4 vs. t0)**

| Downregulated genes | | | | Upregulated genes | | | |
|---|---|---|---|---|---|---|---|
| ID | Annotation | log2FC | padj | ID | Annotation | log2FC | padj |
| C4B63_113g62 | kinetoplast DNA-associated protein | -1,491272998 | 0,006597676 | C4B63_1g991 | mucin TcMUCII | 1,056685652 | 0,000319915 |
| C4B63_277g19 | metallo-peptidase, clan MH, family M18 | -1,125696703 | 0,0001823 | C4B63_34g1444c | mucin TcSMUGL | 0,848644158 | 0,006850887 |
| C4B63_550g5 | glutamamyl carboxypeptidase(fragment) | -1,080526501 | 0,000157584 | C4B63_340g11 | conserved hypothetical protein | 0,798725144 | 0,009805255 |
| C4B63_26g34 | mucin TcMUCII | -1,052872168 | 0,005590545 | C4B63_34g1468c | mucin TcSMUGL | 0,760880185 | 0,019128352 |
| C4B63_196g33 | unspecified product | -1,024338014 | 0,011929776 | C4B63_84g22 | conserved hypothetical protein | 0,754999243 | 0,0000682 |
| C4B63_16g295 | RNA-binding protein | -1,011468762 | 0,033283029 | C4B63_32g335 | conserved hypothetical protein | 0,752660374 | 0,005420783 |
| C4B63_39g66 | trans-sialidase, Group V | -0,998673656 | 0,002978564 | C4B63_300g110c | mucin TcSMUGL | 0,750633404 | 0,011856407 |
| C4B63_251g10 | lipase | -0,941622889 | 0,00569034 | C4B63_66g155 | mucin TcMUCII | 0,750196933 | 0,010834936 |
| C4B63_294g3 | metallo-peptidase, clan MH, family M18 | -0,928607714 | 0,000281783 | C4B63_37g169 | mucin TcMUCII | 0,742790592 | 0,030493062 |
| C4B63_59g50 | mucin TcMUCII | -0,909064577 | 0,006001915 | C4B63_104g104 | Shk1 kinase-binding protein 15 | 0,730721588 | 0,030558868 |
| C4B63_24g236 | microtubule-associated protein Gb4 | -0,899693504 | 0,03686757 | C4B63_2g717 | conserved hypothetical protein | 0,718599066 | 0,045833354 |
| C4B63_22g60 | homocysteine S-methyltransferase | -0,875274723 | 0,008576744 | C4B63_143g34 | condensin complex subunit 2 | 0,703768919 | 0,005680765 |
| C4B63_323g3 | unspecified product | -0,866529977 | 0,001530751 | C4B63_157g25 | conserved hypothetical protein | 0,694494366 | 0,012936307 |
| C4B63_93g11 | unspecified product | -0,856930551 | 0,001078579 | C4B63_2g293 | conserved hypothetical protein | 0,688502397 | 0,018803208 |
| C4B63_37g189 | trans-sialidase, Group VI | -0,853893005 | 0,000123727 | C4B63_235g11c | conserved hypothetical protein | 0,684189874 | 0,00411543 |
| C4B63_76g66 | conserved hypothetical protein | -0,850441886 | 0,009913188 | C4B63_14g241 | trypanothione reductase | 0,68365927 | 0,005063519 |
| C4B63_5g262 | retrotransposon hot spot protein (RHS) | -0,846985965 | 0,013833633 | C4B63_71g130 | mucin TcMUCII | 0,68129353 | 0,030341622 |
| C4B63_62g85 | Proline dehydrogenase, mitochondrial | -0,825991432 | 0,004973215 | C4B63_34g331 | unspecified product | 0,67494903 | 0,009858092 |
| C4B63_18g330 | zinc-finger protein ZFP1 | -0,818825829 | 0,002318746 | C4B63_360g13 | unspecified product | 0,669991258 | 0,00586106 |
| C4B63_42g559c | conserved hypothetical protein | -0,817766071 | 0,00393794 | C4B63_230g118c | DNA-dependent RNA polymerases | 0,666718759 | 0,024710912 |
| C4B63_52g133 | kinesin-like protein | -0,798216479 | 0,023321142 | C4B63_2g486 | Serine-threonine kinase receptor-associated protein | 0,648091033 | 0,002062963 |
| C4B63_156g3 | unspecified product | -0,791084441 | 0,01219801 | C4B63_252g23 | Serine-threonine kinase receptor-associated protein | 0,64690548 | 0,002875611 |
| C4B63_244g8 | trans-sialidase, Group I | -0,774940085 | 0,049028737 | C4B63_109g37 | Gar1/Naf1 RNA binding region containing protein | 0,641919407 | 0,002878955 |
| C4B63_9g376 | Present in the outer mitochondrial membrane proteome 22 | -0,740594834 | 0,024916294 | C4B63_84g49 | RNA-binding protein | 0,63847623 | 0,001059232 |
| C4B63_322g4 | glutamamyl carboxypeptidase(fragment) | -0,736899039 | 0,010674259 | C4B63_34g1482c | mucin TcSMUGL | 0,634373452 | 0,007326816 |
| C4B63_10g453 | ABC transporter | -0,732037826 | 0,001145357 | C4B63_59g110 | mucin TcMUCII | 0,627143817 | 0,026762975 |
| C4B63_58g126 | ubiquitin-conjugating enzyme variant Kua homologue | -0,729538818 | 0,002168935 | C4B63_300g114c | mucin TcSMUGL | 0,609905037 | 0,011929776 |
| C4B63_59g45 | mucin TcMUCII | -0,721508881 | 0,023635906 | C4B63_143g4 | conserved hypothetical protein | 0,606587012 | 0,045308219 |
| C4B63_156g4 | unspecified product | -0,720004159 | 0,01392578 | C4B63_34g1450c | mucin TcSMUGL | 0,605194506 | 0,015262481 |
| C4B63_24g194 | microtubule-associated protein | -0,714920541 | 0,027428349 | C4B63_210g33 | p-nitrophenylphosphatase | 0,603373903 | 0,03231057 |
| C4B63_96g38 | trans-sialidase, Group V | -0,71124491 | 0,027024937 | C4B63_86g57 | conserved hypothetical protein | 0,596096694 | 0,041920586 |
| C4B63_127g46 | conserved hypothetical protein | -0,700795782 | 0,002496736 | C4B63_34g1434c | mucin TcSMUGL | 0,593241939 | 0,022370828 |
| C4B63_70g135 | protein associated with differentiation 4(fragment) | -0,699226625 | 0,009967466 | C4B63_220g11 | Beta propeller protein 1 | 0,592239267 | 0,02141336 |
| C4B63_30g275 | protein disulfide isomerase | -0,694352114 | 0,0000251 | C4B63_149g18 | Beta propeller protein 1 | 0,591214018 | 0,043560334 |
| C4B63_6g1868c | DNA-binding protein STK 13740 | -0,692719099 | 0,001683369 | C4B63_42g121 | L-threonine 3-dehydrogenase | 0,590242402 | 0,012393814 |
| C4B63_47g122 | trans-sialidase | -0,692636437 | 0,010520518 | C4B63_184g8 | GTP-binding protein | 0,588993473 | 0,006773874 |
| C4B63_63g111 | hexose transporter(fragment) | -0,69239724 | 0,015896957 | C4B63_84g67 | conserved hypothetical protein | 0,585718638 | 0,030562008 |
| C4B63_65g5 | unspecified product | -0,689101 | 0,017502581 | C4B63_34g1486c | mucin TcSMUGL | 0,58439393 | 0,012184223 |
| C4B63_281g9 | Ring finger domain containing protein | -0,688110415 | 0,01590677 | C4B63_42g127 | L-threonine 3-dehydrogenase | 0,580873932 | 0,009244919 |
| C4B63_133g46 | cytosolic malate dehydrogenase | -0,683750214 | 0,01216936 | C4B63_38g39 | actin 3 | 0,576593174 | 0,001852102 |
| C4B63_24g274 | conserved hypothetical protein | -0,683612142 | 0,002695177 | C4B63_34g1437c | mucin TcSMUGL | 0,568505126 | 0,045866402 |
| C4B63_66g68 | trans-sialidase, Group V | -0,675641984 | 0,020840385 | C4B63_2g564 | mRNA export factor MEX67 | 0,568430043 | 0,000000866 |
| C4B63_1g473 | trans-sialidase | -0,673702844 | 0,002926133 | C4B63_124g19 | surface protease GP63 | 0,567983869 | 0,027230336 |
| C4B63_459g7 | UDP-Gal or UDP-GlcNAc-dependent glycosyltransferase | -0,669215088 | 0,010391738 | C4B63_42g114 | L-threonine 3-dehydrogenase | 0,566495214 | 0,016196089 |
| C4B63_4g132 | NLI interacting factor-like phosphatase | -0,66685925 | 0,006126201 | C4B63_34g1498c | mucin TcSMUGL | 0,566259148 | 0,034198536 |
| C4B63_45g145 | WD and tetratricopeptide repeats protein 1 | -0,665187644 | 0,026822319 | C4B63_18g2 | L-threonine 3-dehydrogenase | 0,563220635 | 0,013602762 |
| C4B63_590g2 | unspecified product | -0,661990274 | 0,015921928 | C4B63_369g1 | conserved hypothetical protein | 0,562189688 | 0,049198405 |
| C4B63_33g147 | NADH-ubiquinone oxidoreductase, mitochondrial | -0,659932987 | 0,010892998 | C4B63_42g129 | L-threonine 3-dehydrogenase | 0,561918177 | 0,029615322 |
| C4B63_66g170 | Mucin-associated surface protein (MASP) | -0,657922994 | 0,021861596 | C4B63_18g130 | glutamyl-tRNA synthetase | 0,561626078 | 0,000375376 |
| C4B63_2g317 | unspecified product | -0,651132466 | 0,023635906 | C4B63_4g383 | mevalonate-diphosphate decarboxylase | 0,559633154 | 0,008313589 |
| C4B63_59g207 | mucin TcMUCII | -0,650992214 | 0,008355943 | C4B63_38g38 | actin 3 | 0,559620101 | 0,008889572 |
| C4B63_25g209 | conserved hypothetical protein | -0,647865422 | 0,023635906 | C4B63_42g111 | L-threonine 3-dehydrogenase | 0,558934223 | 0,023321142 |
| C4B63_13g328 | conserved hypothetical protein | -0,645262507 | 0,020448846 | C4B63_9g201 | GTP-binding protein | 0,558135972 | 0,013760194 |
| C4B63_19g266 | conserved hypothetical protein | -0,644365748 | 0,02875939 | C4B63_151g36 | notchless homolog | 0,558066567 | 0,004031696 |
| C4B63_258g25 | lipase | -0,637348544 | 0,015207458 | C4B63_47g35 | lysosomal alpha-mannosidase precursor | 0,543716829 | 0,027276024 |
| C4B63_45g123 | chaperone protein DNAj | -0,631265699 | 0,004283349 | C4B63_44g192 | mitochondrial RNA binding protein | 0,540547316 | 0,00118148 |
| C4B63_10g472 | conserved hypothetical protein | -0,626480822 | 0,019492486 | C4B63_9g220 | Thioredoxin domain-containing protein PLP3A | 0,540198007 | 0,042836092 |
| C4B63_9g377 | Present in the outer mitochondrial membrane proteome 22 | -0,626025055 | 0,007304114 | C4B63_129g42 | diphthine synthase | 0,538907073 | 0,021465037 |
| C4B63_13g208 | conserved hypothetical protein,zinc finger protein family member | -0,62364446 | 0,006175825 | C4B63_28g50 | ATP-dependent DEAD/H RNA helicase(fragment) | 0,537753105 | 0,022117697 |
| C4B63_419g2 | trans-sialidase | -0,620712433 | 0,019253706 | C4B63_4g144 | protein transport protein SEC13 | 0,536622565 | 0,005424956 |
| C4B63_45g132 | conserved hypothetical protein | -0,619956649 | 0,014533057 | C4B63_20g747c | cyclin-dependent kinases regulatory subunit | 0,536556229 | 0,002377663 |
| C4B63_212g4 | protein associated with differentiation 4 | -0,619174128 | 0,043790207 | C4B63_171g4 | phosphatidic acid phosphatase protein | 0,531081048 | 0,015840078 |
| C4B63_173g8 | small GTP-binding protein RAB6 | -0,618785099 | 0,031792661 | C4B63_22g17 | Leucine Rich Repeat | 0,527557744 | 0,048273311 |
| C4B63_4g545 | RNA-binding protein 4 | -0,615196022 | 0,027253468 | C4B63_42g118 | L-threonine 3-dehydrogenase | 0,52674916 | 0,013171673 |
| C4B63_9g249 | conserved hypothetical protein | -0,612428614 | 0,007350121 | C4B63_63g95 | poly(A)-specific ribonuclease PARN | 0,525293232 | 0,004163347 |
| C4B63_68g103 | trans-sialidase, Group II | -0,610409768 | 0,001697954 | C4B63_161g18 | UDP-Gal or UDP-GlcNAc-dependent glycosyltransferase | 0,523741555 | 0,015980364 |
| C4B63_47g125 | trans-sialidase | -0,609634333 | 0,007005061 | C4B63_18g8 | L-threonine 3-dehydrogenase | 0,519095114 | 0,027215696 |
| C4B63_12g151 | conserved hypothetical protein | -0,608122017 | 0,002190287 | C4B63_42g124 | L-threonine 3-dehydrogenase | 0,518611069 | 0,015241811 |
| C4B63_89g19 | retrotransposon hot spot (RHS) protein | -0,604264899 | 0,003987376 | C4B63_38g146 | conserved hypothetical protein | 0,517304357 | 0,002483695 |
| C4B63_1g702 | trans-sialidase, Group V | -0,60055567 | 0,026762975 | C4B63_40g159 | translation initiation factor 2 subunit | 0,516665266 | 0,020618613 |
| C4B63_15g33c | SELENOT | -0,599380248 | 0,021735987 | C4B63_7g423 | proteasome beta 7 subunit | 0,51082011 | 0,040109891 |
| C4B63_301g7 | conserved hypothetical protein | -0,597693821 | 0,005707409 | C4B63_56g31 | conserved hypothetical protein | 0,5087836 | 0,001932478 |
| C4B63_69g318c | conserved hypothetical protein | -0,595896853 | 0,009748183 | C4B63_2g43 | short-chain dehydrogenase | 0,507804916 | 0,01774239 |
| C4B63_97g15 | Zinc finger CCCH domain-containing protein 35 | -0,593215481 | 0,001504756 | C4B63_7g266 | conserved hypothetical protein | 0,507570945 | 0,000416235 |
| C4B63_163g33 | protein disulfide isomerase | -0,589596931 | 0,000459489 | C4B63_17g46 | U3 small nucleolar ribonucleoprotein protein IMP3 | 0,504119692 | 0,033152461 |
| C4B63_53g165 | conserved hypothetical protein | -0,58608106 | 0,003725051 | C4B63_34g1514c | mucin TcSMUGL | 0,504055747 | 0,028387565 |
| C4B63_27g184 | integral membrane transport protein | -0,584932706 | 0,000832912 | C4B63_286g18 | eukaryotic translation initiation factor 3 subunit c | 0,503554465 | 0,007245376 |
| C4B63_455g9 | trans-sialidase, Group VII | -0,583034363 | 0,026955096 | C4B63_160g6 | poly(A)-specific ribonuclease PARN | 0,500982479 | 0,000862697 |
| C4B63_132g20 | trans-sialidase, Group V | -0,582654756 | 0,024710912 | C4B63_18g152 | conserved hypothetical protein | 0,500017375 | 0,01392578 |
| C4B63_10g454 | ABC transporter(fragment) | -0,582439804 | 0,033295977 | C4B63_52g161 | tyrosyl-tRNA synthetase | 0,499786807 | 0,006603861 |
| C4B63_208g27 | conserved hypothetical protein | -0,575002805 | 0,02661247 | C4B63_44g207 | C-5 sterol desaturase | 0,499730532 | 0,002377609 |
| C4B63_9g234 | dolichyl-P-Man:GDP-Man7GlcNAc2-PP-dolichyl alpha-1,6-mannosyl | -0,574410051 | 0,000233382 | C4B63_2g294 | 40 kDa cyclophilin | 0,495373267 | 0,0000133 |
| C4B63_234g7 | WD and tetratricopeptide repeats protein 1 | -0,570087186 | 0,015718876 | C4B63_18g11 | L-threonine 3-dehydrogenase | 0,49158537 | 0,030312503 |
| C4B63_10g115 | conserved hypothetical protein | -0,569718493 | 0,014140775 | C4B63_37g366 | thimet oligopeptidase | 0,491397312 | 0,001333412 |
| C4B63_24g275 | conserved hypothetical protein | -0,564707684 | 0,012144168 | C4B63_22g22 | ARM-like helical domain-containing protein | 0,490895744 | 0,002657255 |
| C4B63_44g243 | conserved hypothetical protein | -0,560557058 | 0,010457873 | C4B63_227g2 | glucan 1,3-alpha-glucosidase | 0,488735006 | 0,002445833 |
| C4B63_10g455 | ABC transporter | -0,55692899 | 0,022228807 | C4B63_34g1502c | mucin TcSMUGL | 0,488011657 | 0,02391789 |
| C4B63_92g56 | conserved hypothetical protein | -0,550577734 | 0,030312503 | C4B63_28g56 | TFIIF-stimulated CTD phosphatase | 0,485537395 | 0,01392578 |
| C4B63_55g34 | Paraflagellar rod protein 2 | -0,545408859 | 0,0000165 | C4B63_12g362 | UDP-Gal or UDP-GlcNAc-dependent glycosyltransferase | 0,484058728 | 0,033680118 |
| C4B63_48g7 | NLI interacting factor-like phosphatase | -0,544613437 | 0,002533081 | C4B63_9g524 | eukaryotic translation initiation factor 3 subunit k | 0,482928577 | 0,007199187 |
| C4B63_93g32 | unspecified product | -0,543081853 | 0,044290358 | C4B63_4g375 | glucan 1,3-alpha-glucosidase | 0,482693629 | 0,008198134 |
| C4B63_51g112 | conserved hypothetical protein | -0,54244053 | 0,036545688 | C4B63_4g72 | fibrillarin | 0,480818342 | 0,030558868 |
| C4B63_16g93 | Uncharacterized protein in mobD 3'region | -0,539062038 | 0,007546807 | C4B63_200g53 | CorA-like Mg2+ transporter protein | 0,48053743 | 0,018191323 |
| C4B63_44g125 | conserved hypothetical protein | -0,530216991 | 0,001455235 | C4B63_1g70 | mitochondrial RNA-binding protein 2 | 0,479500526 | 0,016172345 |
| C4B63_20g203 | Zinc finger protein, C3H1 type-like | -0,526769702 | 0,025089488 | C4B63_4g118 | conserved hypothetical protein | 0,4762187 | 0,017014025 |
| C4B63_61g145 | conserved hypothetical protein | -0,52426101 | 0,035453493 | C4B63_62g82 | thimet oligopeptidase | 0,475666229 | 0,002650891 |
| C4B63_6g587 | N-glycosidase YbiA | -0,523617678 | 0,048042797 | C4B63_114g25 | conserved hypothetical protein | 0,474925834 | 0,025463989 |
| C4B63_12g334 | cytochrome oxidase subunit VII | -0,522476252 | 0,002081925 | C4B63_39g315 | CLP1 | 0,473055613 | 0,046197417 |
| C4B63_55g40 | Paraflagellar rod protein 2 | -0,52016133 | 0,000843714 | C4B63_257g3 | amino acid transporter | 0,47223583 | 0,046277832 |
| C4B63_18g326 | zinc-finger protein ZFP1 | -0,519429976 | 0,00432494 | C4B63_48g110 | ATP-dependent DEAD/H RNA helicase | 0,471871354 | 0,001997805 |
| C4B63_55g55 | Paraflagellar rod protein 2 | -0,519386479 | 0,000138598 | C4B63_127g54 | squalene synthase | 0,469365124 | 0,000103291 |
| C4B63_102g335c | conserved hypothetical protein | -0,517492983 | 0,035557232 | C4B63_7g229 | Probable 18S rRNA (guanine-N(7))-methyltransferase | 0,469109268 | 0,048103956 |
| C4B63_44g235 | Kinesin-13 4 | -0,516394857 | 0,003874503 | C4B63_70g98 | Optic atrophy 3 protein (OPA3) | 0,468903107 | 0,010466564 |
| C4B63_40g103 | conserved hypothetical protein | -0,516293444 | 0,023635906 | C4B63_6g2548c | histone H3 | 0,46850994 | 0,046882791 |
| C4B63_2g816 | conserved hypothetical protein | -0,516247792 | 0,002475525 | C4B63_7g318 | peroxisome biogenesis factor 1 | 0,467281774 | 0,017658376 |
| C4B63_9g382 | conserved hypothetical protein | -0,515215123 | 0,001583736 | C4B63_5g413 | mitochondrial RNA-binding protein 2 | 0,466389742 | 0,022476404 |
| C4B63_55g38 | Paraflagellar rod protein 2 | -0,515158696 | 0,001234613 | C4B63_183g14 | conserved hypothetical protein | 0,465933536 | 0,003949475 |
| C4B63_40g30 | conserved hypothetical protein | -0,51448305 | 0,030562008 | C4B63_4g219 | nucleolar RNA-associated proteins (Nrap) | 0,464448514 | 0,004163347 |
| C4B63_40g102 | conserved hypothetical protein | -0,514381695 | 0,014192706 | C4B63_26g212 | Transmembrane protein 164 homolog | 0,464173265 | 0,009551305 |
| C4B63_44g183 | conserved hypothetical protein | -0,512968853 | 0,003237062 | C4B63_260g15 | inosine-5'-monophosphate dehydrogenase | 0,463562844 | 0,02919287 |
| C4B63_16g183 | surface protease GP63 | -0,512342258 | 0,037779166 | C4B63_7g347 | V-type ATPase, A subunit | 0,460504505 | 0,027903449 |
| C4B63_221g9 | unspecified product | -0,512170223 | 0,048751822 | C4B63_91g34 | conserved hypothetical protein | 0,460125847 | 0,001645365 |
| C4B63_35g53 | flagellar calcium-binding protein | -0,512027792 | 0,001147703 | C4B63_22g788c | calpain-like cysteine peptidase | 0,455104804 | 0,032381712 |
| C4B63_98g10 | unspecified product | -0,51006962 | 0,038958985 | C4B63_49g65 | farnesyl pyrophosphate synthase | 0,45496421 | 0,002957664 |
| C4B63_55g46 | Paraflagellar rod protein 2 | -0,508065577 | 0,00072982 | C4B63_23g83 | Aminopeptidase M1 | 0,4545205 | 0,010247388 |
| C4B63_109g41 | Trypomastigote, Alanine, Serine and Valine rich protein (TASV), sub | -0,507041981 | 0,011755777 | C4B63_28g180 | WASH complex subunit 4 | 0,454491894 | 0,020480743 |
| C4B63_154g8 | retrotransposon hot spot protein | -0,505952344 | 0,030856374 | C4B63_63g145 | retrotransposon hot spot (RHS) protein | 0,450542072 | 0,025731754 |
| C4B63_16g317 | oxidoreductase | -0,505669698 | 0,0408403 | C4B63_262g21 | retrotransposon hot spot (RHS) protein | 0,450188847 | 0,047217126 |
| C4B63_58g132 | conserved hypothetical protein | -0,503698405 | 0,011445117 | C4B63_116g15 | membrane transporter protein | 0,449914443 | 0,015241811 |
| C4B63_2g530 | conserved hypothetical protein | -0,502452006 | 0,014124773 | C4B63_2g331 | polyadenylate-binding protein 1 | 0,447024029 | 0,0000767 |
| C4B63_45g115 | Protein kinase domain | -0,501420741 | 0,002062963 | C4B63_17g69 | phenylalanyl-tRNA synthetase alpha chain | 0,445004993 | 0,017283846 |
| C4B63_37g395 | retrotransposon hot spot protein (RHS) | -0,50098391 | 0,031604693 | C4B63_227g7 | transporter | 0,444896347 | 0,00440306 |
| C4B63_55g51 | Paraflagellar rod protein 2 | -0,499817778 | 0,001016888 | C4B63_51g842c | Uncharacterized protein At2g34160 | 0,444774557 | 0,000690495 |
| C4B63_24g301 | phosphatidylinositol-4-phosphate 5-kinase | -0,499524547 | 0,002851551 | C4B63_73g19 | conserved hypothetical protein | 0,443708369 | 0,036939816 |
| C4B63_127g44 | conserved hypothetical protein | -0,497132765 | 0,014724406 | C4B63_10g406 | High mobility group protein TDP1 | 0,441978859 | 0,002978564 |
| C4B63_75g67 | Probable ubiquitin carboxyl-terminal hydrolase MINDY-4 | -0,4969427 | 0,014124773 | C4B63_45g97 | eukaryotic translation initiation factor 3 subunit h | 0,439818384 | 0,015850394 |
| C4B63_24g276 | conserved hypothetical protein | -0,49659444 | 0,028145633 | C4B63_52g55 | Repetitive proline-rich cell wall protein 2 | 0,438953418 | 0,012824198 |
| C4B63_98g42 | retrotransposon hot spot (RHS) protein | -0,496409519 | 0,03775155 | C4B63_260g14 | inosine-5'-monophosphate dehydrogenase | 0,438549213 | 0,01216936 |
| C4B63_2g828 | ankyrin repeat protein | -0,495719634 | 0,047154589 | C4B63_2g744 | ALA-interacting subunit 3 | 0,438083821 | 0,006845812 |
| C4B63_45g139 | serine/threonine protein kinase | -0,495099321 | 0,024846646 | C4B63_38g167 | spermidine synthase | 0,435744064 | 0,005892579 |
| C4B63_40g40 | conserved hypothetical protein | -0,494353284 | 0,003819003 | C4B63_19g110 | Transducin beta-like protein 3 | 0,435639332 | 0,007485755 |
| C4B63_12g249 | Vacuolar protein 8 | -0,494236388 | 0,027201556 | C4B63_253g8 | NADPH--cytochrome P450 reductase | 0,435024323 | 0,011262752 |
| C4B63_26g310 | ABC transporter | -0,489227653 | 0,022316819 | C4B63_20g1417c | conserved hypothetical protein | 0,434039022 | 0,037111123 |
| C4B63_76g22 | cytoskeleton associated protein | -0,487692351 | 0,012184223 | C4B63_63g46 | 2,3-bisphosphoglycerate-independent phosphoglycerate mutase | 0,434029702 | 0,004668484 |
| C4B63_25g220 | 5'-3' exoribonuclease B | -0,485715323 | 0,005892579 | C4B63_7g74 | centrin | 0,43264393 | 0,009178261 |
| C4B63_14g20 | hexose transporter | -0,478689438 | 0,005273356 | C4B63_87g15 | Golgi/lysosome glycoprotein 1 | 0,432459333 | 0,004859761 |
| C4B63_55g50 | Paraflagellar rod protein 2 | -0,47771273 | 0,001916615 | C4B63_18g223 | eukaryotic translation initiation factor 3 subunit d | 0,43199132 | 0,001798226 |
| C4B63_11g51 | Oocyte-specific histone RNA stem-loop-binding protein 2 | -0,476909839 | 0,00225237 | C4B63_150g36 | membrane-bound acid phosphatase 2 | 0,431064631 | 0,040440389 |
| C4B63_97g13 | conserved hypothetical protein | -0,475334409 | 0,000264294 | C4B63_11g260 | conserved hypothetical protein | 0,430863355 | 0,048042797 |
| C4B63_4g99 | conserved hypothetical protein | -0,47505247 | 0,006735859 | C4B63_114g17 | conserved hypothetical protein | 0,430436159 | 0,041005498 |
| C4B63_20g162 | conserved hypothetical protein | -0,474346224 | 0,031056965 | C4B63_13g174 | beta galactofuranosyl glycosyltransferase(fragment) | 0,430169443 | 0,037051781 |
| C4B63_44g214 | conserved hypothetical protein | -0,474176522 | 0,007939695 | C4B63_4g223 | mannosyltransferase-II | 0,430164748 | 0,014047756 |
| C4B63_21g6 | arginine permease | -0,473980627 | 0,033205876 | C4B63_2g289 | Hypoxia up-regulated protein 1 (Fragment) | 0,429379479 | 0,014046149 |
| C4B63_19g230 | calpain-like cysteine peptidase | -0,473616676 | 0,012691725 | C4B63_93g70 | conserved hypothetical protein | 0,428108949 | 0,030804814 |
| C4B63_5g393 | trans-sialidase, Group V | -0,473529466 | 0,027096203 | C4B63_6g586 | proteasome beta 6 subunit | 0,427350565 | 0,028289309 |
| C4B63_272g23 | retrotransposon hot spot (RHS) protein | -0,473475756 | 0,033205876 | C4B63_2g219 | conserved hypothetical protein | 0,426745342 | 0,02655658 |
| C4B63_44g240 | Coiled-coil domain-containing protein 39 | -0,472903071 | 0,018302268 | C4B63_14g39 | conserved hypothetical protein | 0,426113572 | 0,025092956 |
| C4B63_9g383 | conserved hypothetical protein | -0,471299593 | 0,019253706 | C4B63_23g262 | choline/ethanolamine phosphotransferase | 0,425146955 | 0,022224099 |
| C4B63_76g17 | conserved hypothetical protein | -0,467463982 | 0,004608309 | C4B63_137g10 | Coatomer subunit gamma | 0,423310506 | 0,028394012 |
| C4B63_1g953 | trans-sialidase, Group V | -0,465817944 | 0,042450294 | C4B63_32g1134c | BolA-like protein 3 | 0,421650813 | 0,005411319 |
| C4B63_35g69 | flagellar calcium-binding protein | -0,465514979 | 0,001199326 | C4B63_89g45 | Mitochondrial SSU ribosomal protein 29 | 0,421646181 | 0,034186507 |
| C4B63_20g24 | conserved hypothetical protein | -0,464850899 | 0,023867085 | C4B63_52g41 | mitochondrial DNA polymerase I protein C | 0,419007164 | 0,000587101 |
| C4B63_6g284 | cytochrome c oxidase assembly protein | -0,463833031 | 0,020746865 | C4B63_81g17 | DNA mismatch repair protein MSH2(fragment) | 0,418564433 | 0,028096771 |
| C4B63_55g36 | Paraflagellar rod protein 2 | -0,460306228 | 0,001919913 | C4B63_21g53 | RNA-binding protein | 0,418007738 | 0,005892261 |
| C4B63_9g267 | paraflagellar rod protein 5 | -0,457254072 | 0,02655658 | C4B63_58g87 | elongation factor 1-alpha | 0,415349381 | 0,003804811 |
| C4B63_52g170 | cytoskeleton associated protein | -0,455708181 | 0,036694693 | C4B63_4g508 | peptidyl-prolyl cis-trans isomerase | 0,41430609 | 0,012348363 |
| C4B63_42g257 | cyclophilin | -0,455293239 | 0,004163347 | C4B63_60g133 | protein tyrosine phosphatase-like protein | 0,413456657 | 0,004877143 |
| C4B63_70g84 | serine/threonine protein kinase | -0,454792186 | 0,022237695 | C4B63_4g381 | transporter | 0,412251902 | 0,00491455 |
| C4B63_12g230 | palmitoyl acyltransferase 7 | -0,454712068 | 0,022921903 | C4B63_8g47 | telomerase component(fragment) | 0,411992342 | 0,030558868 |
| C4B63_70g86 | conserved hypothetical protein | -0,448162239 | 0,048882077 | C4B63_19g111 | WD domain, G-beta repeat/Utp13 specific WD40 associated domain | 0,411577578 | 0,031898699 |
| C4B63_17g119 | Phosphatidylinositol/phosphatidylcholine transfer protein SFH14 | -0,447353191 | 0,024502938 | C4B63_58g89 | elongation factor 1-alpha | 0,410267764 | 0,005141512 |
| C4B63_55g42 | Paraflagellar rod protein 2 | -0,446516876 | 0,003339381 | C4B63_78g38 | lipoate-protein ligase | 0,408966569 | 0,037923906 |
| C4B63_35g62 | flagellar calcium-binding protein | -0,445394321 | 0,010466564 | C4B63_2g728 | pre-rRNA-processing protein PNO1 | 0,407126318 | 0,01392578 |
| C4B63_32g259 | Cyclin-U4-1 | -0,445012962 | 0,024455455 | C4B63_42g1177c | conserved hypothetical protein | 0,40647616 | 0,029926484 |
| C4B63_267g18 | conserved hypothetical protein | -0,444492612 | 0,041376971 | C4B63_13g189 | ubiquitin-like protein | 0,406106416 | 0,000413066 |
| C4B63_75g60 | NADH-ubiquinone oxidoreductase complex I subunit,FT | -0,443910329 | 0,01112487 | C4B63_2g367 | nucleolar GTP-binding protein 1 | 0,405621897 | 0,003761897 |
| C4B63_55g53 | paraxonemal rod protein PAR2 | -0,443186479 | 0,014121606 | C4B63_117g55 | inosine-5'-monophosphate dehydrogenase | 0,404653432 | 0,023635906 |
| C4B63_35g51 | flagellar calcium-binding protein | -0,439723678 | 0,019116099 | C4B63_9g209 | conserved hypothetical protein | 0,404319561 | 0,018155469 |
| C4B63_35g89 | flagellar calcium-binding protein | -0,439315465 | 0,011454237 | C4B63_38g299 | inosine-5'-monophosphate dehydrogenase | 0,403438784 | 0,014364544 |
| C4B63_35g70 | flagellar calcium-binding protein | -0,438596628 | 0,025703392 | C4B63_42g192 | N-alpha-acetyltransferase 25, NatB auxiliary subunit | 0,402897209 | 0,007456436 |
| C4B63_35g68 | flagellar calcium-binding protein | -0,434666318 | 0,01769851 | C4B63_7g316 | ruvB-like DNA helicase | 0,402142239 | 0,009462316 |
| C4B63_69g76 | conserved hypothetical protein | -0,433470848 | 0,048103956 | C4B63_52g42 | glutathione synthetase | 0,401116231 | 0,004715258 |
| C4B63_30g214 | conserved hypothetical protein | -0,432576876 | 0,000718111 | C4B63_57g75 | dihydrolipoamide acetyltransferase precursor | 0,400774516 | 0,00569034 |
| C4B63_134g59 | trans-sialidase, Group V | -0,432544015 | 0,033233247 | C4B63_45g230 | ER membrane protein complex subunit 8 | 0,400135981 | 0,010520518 |
| C4B63_150g38 | glyceraldehyde 3-phosphate dehydrogenase, cytosolic | -0,431419891 | 0,043066751 | | | | |
| C4B63_35g82 | flagellar calcium-binding protein | -0,429035936 | 0,021960253 | | | | |
| C4B63_231g31 | flagellar calcium-binding protein | -0,428371294 | 0,012000612 | | | | |
| C4B63_51g183 | Cilia- and flagella-associated protein 73 | -0,427139204 | 0,017867172 | | | | |
| C4B63_26g201 | cAMP specific phosphodiesterase | -0,426616536 | 0,006001915 | | | | |
| C4B63_35g97 | flagellar calcium-binding protein | -0,424666891 | 0,004549684 | | | | |
| C4B63_35g71 | flagellar calcium-binding protein | -0,423366036 | 0,005517684 | | | | |
| C4B63_17g196 | calpain-like cysteine peptidase | -0,423293131 | 0,01906923 | | | | |
| C4B63_55g48 | Paraflagellar rod protein 2 | -0,422094472 | 0,008236867 | | | | |
| C4B63_35g81 | flagellar calcium-binding protein | -0,421951986 | 0,019285399 | | | | |
| C4B63_35g84 | flagellar calcium-binding protein | -0,420876196 | 0,022141478 | | | | |
| C4B63_35g75 | flagellar calcium-binding protein | -0,419683251 | 0,005357488 | | | | |
| C4B63_167g40 | trans-sialidase, Group I | -0,419425503 | 0,045560957 | | | | |
| C4B63_80g15 | cAMP response protein 1 | -0,418010319 | 0,023146503 | | | | |
| C4B63_54g35 | protein kinase | -0,417547864 | 0,010362352 | | | | |
| C4B63_4g98 | conserved hypothetical protein | -0,416407471 | 0,004389548 | | | | |
| C4B63_18g16 | GTPase activating protein | -0,415425802 | 0,007637706 | | | | |
| C4B63_231g29 | flagellar calcium-binding protein | -0,41410061 | 0,044556175 | | | | |
| C4B63_35g85 | flagellar calcium-binding protein | -0,412549173 | 0,017283846 | | | | |
| C4B63_35g72 | flagellar calcium-binding protein | -0,410888687 | 0,038311039 | | | | |
| C4B63_95g22 | actin-like protein | -0,410831469 | 0,025871188 | | | | |
| C4B63_51g192 | protein kinase A regulatory subunit | -0,410552367 | 0,02437719 | | | | |
| C4B63_35g80 | flagellar calcium-binding protein | -0,409432312 | 0,014157161 | | | | |
| C4B63_26g198 | cAMP specific phosphodiesterase | -0,4078985 | 0,025485141 | | | | |
| C4B63_259g15 | Paladin | -0,405634516 | 0,032421161 | | | | |
| C4B63_26g243 | nucleobase transporter | -0,405460202 | 0,00894204 | | | | |
| C4B63_145g23 | trans-sialidase, Group III | -0,405368026 | 0,036224661 | | | | |
| C4B63_24g277 | conserved hypothetical protein | -0,404361663 | 0,011474439 | | | | |
| C4B63_9g378 | conserved hypothetical protein | -0,400017952 | 0,039421601 | | | | |

**Hemin exclusive DEGs (t24 vs. t0)**

| Downregulated genes | | | | Upregulated genes | | | |
|---|---|---|---|---|---|---|---|
| ID | Annotation | log2FC | padj | ID | Annotation | log2FC | padj |
| C4B63_476nc10 | unspecified product | -7,524046267 | 0,00037481 | C4B63_10g404 | guanine deaminase | 0,980741141 | 1,32797E-05 |
| C4B63_522nc5 | unspecified product | -2,757626096 | 0,039063192 | C4B63_348g24 | guanine deaminase | 0,949438924 | 0,011176378 |
| C4B63_79g110 | conserved hypothetical protein | -1,558581495 | 0,026583072 | C4B63_11g259 | membrane transporter protein | 0,890016899 | 0,000414773 |
| C4B63_277g19 | metallo-peptidase, clan MH, family M18 | -1,372377488 | 2,87907E-06 | C4B63_188g155c | histone H4 | 0,801843951 | 0,004253111 |
| C4B63_550g5 | glutamamyl carboxypeptidase(fragment) | -1,260698885 | 6,084E-06 | C4B63_183g12 | mitochondrial carrier protein | 0,793495723 | 0,000332054 |
| C4B63_24g236 | microtubule-associated protein Gb4 | -1,227852864 | 0,001743648 | C4B63_128g313c | histone H4 | 0,791409959 | 0,002562355 |
| C4B63_113g62 | kinetoplast DNA-associated protein | -1,156067027 | 0,049378282 | C4B63_63g4 | PSP1 C-terminal conserved region | 0,782456752 | 0,007325919 |
| C4B63_431g2 | glutamamyl carboxypeptidase(fragment) | -1,152316274 | 0,002562355 | C4B63_133g1 | NAD-specific glutamate dehydrogenase | 0,773958435 | 0,008792691 |
| C4B63_18g39 | CRAL/TRIO, N-terminal domain/CRAL/TRIO domain containing protein | -1,140302488 | 7,52354E-08 | C4B63_50g198 | Cold-inducible RNA-binding protein | 0,76377039 | 0,00414049 |
| C4B63_76g64 | conserved hypothetical protein | -1,106142128 | 0,005336865 | C4B63_6g297 | conserved hypothetical protein | 0,762987463 | 0,000224026 |
| C4B63_26g34 | mucin TcMUCII | -1,099813999 | 0,002944967 | C4B63_72g114 | conserved hypothetical protein | 0,735058565 | 0,000442397 |
| C4B63_60g128 | Right handed beta helix region containing protein | -1,065240017 | 0,049474565 | C4B63_485g4 | conserved hypothetical protein | 0,731161205 | 0,007080075 |
| C4B63_196g33 | unspecified product | -1,047449848 | 0,008641097 | C4B63_38g144 | mitochondrial carrier protein | 0,727571658 | 0,000353295 |
| C4B63_46g95 | conserved hypothetical protein | -1,025398505 | 2,54541E-05 | C4B63_188g150c | histone H4 | 0,718128427 | 0,005779114 |
| C4B63_44g132 | multidrug resistance protein E | -1,015255565 | 0,000106681 | C4B63_188g142c | histone H4 | 0,71551909 | 0,001836795 |
| C4B63_277g7 | glutamamyl carboxypeptidase | -0,988644358 | 6,32048E-05 | C4B63_8g510 | conserved hypothetical protein | 0,700486786 | 0,00206211 |
| C4B63_114g5 | conserved hypothetical protein | -0,955079632 | 0,032651482 | C4B63_13g305 | conserved hypothetical protein | 0,695786822 | 0,000913131 |
| C4B63_46g80 | conserved hypothetical protein | -0,951085299 | 1,39727E-11 | C4B63_12g220 | pyruvate phosphate dikinase | 0,695443693 | 0,026898793 |
| C4B63_58g126 | ubiquitin-conjugating enzyme variant Kua homologue | -0,945836349 | 2,54541E-05 | C4B63_315g8 | amino acid transporter | 0,688710121 | 0,008414612 |
| C4B63_27g1020c | RNA-binding protein | -0,941771858 | 0,000447645 | C4B63_25g228 | Putative uridine kinase C227.14 | 0,686161059 | 0,019679226 |
| C4B63_341g21 | lipase | -0,939401502 | 0,007277508 | C4B63_12g221 | pyruvate phosphate dikinase | 0,663578964 | 0,032565759 |
| C4B63_294g3 | metallo-peptidase, clan MH, family M18 | -0,922130514 | 0,000274856 | C4B63_286g15 | PSP1 C-terminal conserved region | 0,663198107 | 0,010414819 |
| C4B63_56g41 | protein kinase | -0,921589155 | 0,004011107 | C4B63_143g34 | condensin complex subunit 2 | 0,660624507 | 0,009933383 |
| C4B63_46g96 | conserved hypothetical protein | -0,913237196 | 0,000785887 | C4B63_25g113 | GMP reductase | 0,659598859 | 0,025756666 |
| C4B63_550g4 | glutamamyl carboxypeptidase(fragment) | -0,904984543 | 6,36279E-05 | C4B63_49g106 | histone H2A | 0,656189228 | 0,005368603 |
| C4B63_10g453 | ABC transporter | -0,891333282 | 2,71074E-05 | C4B63_227g9 | mevalonate-diphosphate decarboxylase | 0,651319742 | 0,001517938 |
| C4B63_5g426 | trans-sialidase, Group II | -0,886445942 | 0,016336897 | C4B63_39g336 | S-adenosylhomocysteine hydrolase | 0,648810588 | 0,020445396 |
| C4B63_9g377 | Present in the outer mitochondrial membrane proteome 22 | -0,88266273 | 3,15289E-05 | C4B63_6g397 | cation transporter | 0,642594427 | 0,000199291 |
| C4B63_9g376 | Present in the outer mitochondrial membrane proteome 22 | -0,876850919 | 0,004570017 | C4B63_48g25 | conserved hypothetical protein | 0,642347015 | 2,17218E-05 |
| C4B63_104g98 | lipase | -0,873614871 | 0,003115083 | C4B63_282g15 | cation transporter | 0,641061325 | 4,63169E-05 |
| C4B63_341g17 | lipase | -0,86681571 | 0,019587962 | C4B63_214g4 | amino acid transporter | 0,63960715 | 0,01970888 |
| C4B63_76g66 | conserved hypothetical protein | -0,864033364 | 0,007490529 | C4B63_17g257 | chromosomal passenger protein | 0,639411386 | 0,008141978 |
| C4B63_71g133 | trans-sialidase, Group VI | -0,861793045 | 0,025898256 | C4B63_210g35 | 3,2-trans-enoyl-CoA isomerase, mitochondrial precursor | 0,623425809 | 4,63169E-05 |
| C4B63_258g25 | lipase | -0,8458152 | 0,000547544 | C4B63_13g190 | cdc2-related kinase 3 | 0,623420169 | 0,009933801 |
| C4B63_244g6 | trans-sialidase, Group I | -0,843414883 | 0,018745063 | C4B63_2g223 | conserved hypothetical protein | 0,620508131 | 0,015705403 |
| C4B63_322g4 | glutamamyl carboxypeptidase(fragment) | -0,831332072 | 0,00260717 | C4B63_18g233 | glyceraldehyde 3-phosphate dehydrogenase | 0,619366059 | 0,026898793 |
| C4B63_170g26 | conserved hypothetical protein | -0,825300924 | 4,14849E-07 | C4B63_10g449 | Triple RNA binding domain protein 3 | 0,618327253 | 0,003267656 |
| C4B63_331g5 | UDP-Gal or UDP-GlcNAc-dependent glycosyltransferase | -0,823396301 | 0,036001703 | C4B63_302g60c | histone H3 | 0,61629756 | 0,027109777 |
| C4B63_139g11 | Mucin-associated surface protein (MASP) | -0,817057809 | 0,010669581 | C4B63_25g287 | conserved hypothetical protein | 0,616162041 | 0,031292014 |
| C4B63_65g5 | unspecified product | -0,810183196 | 0,003286696 | C4B63_147g266c | conserved hypothetical protein | 0,615857775 | 0,043445984 |
| C4B63_44g243 | conserved hypothetical protein | -0,805491511 | 6,19392E-05 | C4B63_197g16 | Nucleolar protein 10 | 0,614933133 | 0,03894059 |
| C4B63_5g689 | Mucin-associated surface protein (MASP) | -0,800731848 | 0,045768344 | C4B63_4g381 | transporter | 0,610914143 | 3,30738E-06 |
| C4B63_27g201 | conserved hypothetical protein | -0,798649471 | 0,041938564 | C4B63_263g4 | amidinotransferase | 0,601219422 | 0,000510127 |
| C4B63_46g66 | conserved hypothetical protein | -0,786083326 | 0,001534936 | C4B63_60g172 | conserved hypothetical protein | 0,598632177 | 0,031179217 |
| C4B63_10g454 | ABC transporter(fragment) | -0,769080501 | 0,002112707 | C4B63_18g232 | glyceraldehyde 3-phosphate dehydrogenase | 0,59858398 | 0,031230087 |
| C4B63_96g38 | trans-sialidase, Group V | -0,761739029 | 0,015242092 | C4B63_58g135 | heat shock protein DNAJ | 0,590218303 | 0,011081989 |
| C4B63_46g177 | conserved hypothetical protein | -0,756410473 | 1,7413E-05 | C4B63_102g80 | kinesin | 0,589966662 | 0,040075561 |
| C4B63_39g146 | Mucin-associated surface protein (MASP) | -0,756208729 | 0,026176204 | C4B63_6g562 | NAD(P)-dependent steroid dehydrogenase protein | 0,589567117 | 0,00497123 |
| C4B63_21g22 | arginine permease(fragment) | -0,754385443 | 0,001585546 | C4B63_210g33 | p-nitrophenylphosphatase | 0,587618112 | 0,037423017 |
| C4B63_10g455 | ABC transporter | -0,748775059 | 0,000735618 | C4B63_220g11 | Beta propeller protein 1 | 0,585021556 | 0,022015125 |
| C4B63_251g10 | lipase | -0,748479878 | 0,03612984 | C4B63_6g2514c | histone H3 | 0,583989763 | 0,035006462 |
| C4B63_69g318c | conserved hypothetical protein | -0,74394346 | 0,000560111 | C4B63_17g80 | Staphylococcal nuclease domain-containing protein 1 | 0,582910667 | 3,51903E-08 |
| C4B63_2g318 | conserved hypothetical protein | -0,740422658 | 0,019587962 | C4B63_91g43 | conserved hypothetical protein | 0,580169023 | 0,021876261 |
| C4B63_22g32 | conserved hypothetical protein | -0,735978625 | 0,000331377 | C4B63_302g76c | histone H3 | 0,578943117 | 0,018596166 |
| C4B63_354g1 | electron transfer flavoprotein | -0,729021385 | 0,00045011 | C4B63_76g52 | pyrroline-5-carboxylate reductase | 0,573909517 | 4,10915E-05 |
| C4B63_24g194 | microtubule-associated protein | -0,728587294 | 0,022633885 | C4B63_7g316 | ruvB-like DNA helicase | 0,573603579 | 3,85402E-05 |
| C4B63_109g67 | conserved hypothetical protein | -0,725447073 | 0,000282911 | C4B63_2g740 | conserved hypothetical protein | 0,572743864 | 0,023048591 |
| C4B63_22g37 | conserved hypothetical protein | -0,723623538 | 0,000224545 | C4B63_6g305 | heat shock protein 70 (hsp70) | 0,572268981 | 0,017917356 |
| C4B63_17g329 | retrotransposon hot spot protein (RHS) | -0,722266815 | 0,031923171 | C4B63_137g25 | conserved hypothetical protein | 0,567400826 | 0,015650146 |
| C4B63_11g199 | Mucin-associated surface protein (MASP) | -0,713262826 | 0,003689901 | C4B63_84g67 | conserved hypothetical protein | 0,566932472 | 0,036955575 |
| C4B63_323g3 | unspecified product | -0,709228562 | 0,011542212 | C4B63_83g36 | ppnN | 0,556545324 | 0,010888526 |
| C4B63_36g184 | trans-sialidase, Group V | -0,7046605 | 0,006203924 | C4B63_80g43 | conserved hypothetical protein | 0,555326233 | 0,00866125 |
| C4B63_1g702 | trans-sialidase, Group V | -0,703233611 | 0,006118657 | C4B63_19g45 | isovaleryl-coA dehydrogenase | 0,553881395 | 1,18064E-05 |
| C4B63_10g472 | conserved hypothetical protein | -0,703028112 | 0,006210407 | C4B63_17g81 | Staphylococcal nuclease domain-containing protein 1 | 0,553307191 | 7,08758E-06 |
| C4B63_22g40 | conserved hypothetical protein | -0,698675162 | 0,000365312 | C4B63_25g279 | J Biosynthesis Protein 2 | 0,552109013 | 0,001384542 |
| C4B63_34g124 | conserved hypothetical protein | -0,698524597 | 2,61393E-07 | C4B63_156g14 | Succinate-acetate transporter protein/GPR1/FUN34/yaaH family | 0,551574906 | 0,009932793 |
| C4B63_51g215 | unspecified product | -0,695417503 | 0,033510301 | C4B63_12g254 | conserved hypothetical protein | 0,55157336 | 0,01473025 |
| C4B63_22g28 | conserved hypothetical protein | -0,695394428 | 0,000529199 | C4B63_171g4 | phosphatidic acid phosphatase protein | 0,548256283 | 0,010805506 |
| C4B63_11g57 | CHAP domain containing protein | -0,693551706 | 4,13934E-08 | C4B63_252g23 | Serine-threonine kinase receptor-associated protein | 0,548046412 | 0,014899909 |
| C4B63_95g4 | unspecified product | -0,692674219 | 0,041674481 | C4B63_2g219 | conserved hypothetical protein | 0,545781744 | 0,00193534 |
| C4B63_24g988c | conserved hypothetical protein | -0,685757136 | 0,008568773 | C4B63_26g272 | conserved hypothetical protein | 0,544108305 | 0,000468722 |
| C4B63_17g328 | retrotransposon hot spot protein (RHS) | -0,684236805 | 0,041713463 | C4B63_51g842c | Uncharacterized protein At2g34160 | 0,543486057 | 1,17172E-05 |
| C4B63_21g246 | conserved hypothetical protein | -0,682023336 | 0,011595774 | C4B63_43g2 | kinesin | 0,540619936 | 0,039335856 |
| C4B63_17g61 | conserved hypothetical protein | -0,681595232 | 0,004411358 | C4B63_184g8 | GTP-binding protein | 0,536639201 | 0,015376685 |
| C4B63_156g4 | unspecified product | -0,681276864 | 0,020242584 | C4B63_2g726 | CRAL/TRIO domain containing protein | 0,536236175 | 0,010184858 |
| C4B63_152g45 | unspecified product | -0,680126757 | 0,030970188 | C4B63_7g340 | beta-fructofuranosidase-like protein | 0,534773214 | 0,019587962 |
| C4B63_13g223 | zinc-finger protein family member | -0,676653382 | 0,033612624 | C4B63_4g440 | conserved hypothetical protein | 0,533839385 | 0,039093201 |
| C4B63_96g34 | trans-sialidase, Group V | -0,676154381 | 0,042950925 | C4B63_13g226 | conserved hypothetical protein | 0,530950392 | 0,002861643 |
| C4B63_168g180c | conserved hypothetical protein | -0,674389845 | 0,013963358 | C4B63_32g205 | notchless homolog | 0,530409619 | 0,00275003 |
| C4B63_5g369 | unspecified product | -0,67225794 | 0,027878506 | C4B63_2g218 | PGAP1-like protein/Alpha/beta hydrolase family | 0,529546389 | 0,004144332 |
| C4B63_157g36 | trans-sialidase, Group II | -0,667734994 | 0,016336897 | C4B63_52g161 | tyrosyl-tRNA synthetase | 0,528101333 | 0,00303343 |
| C4B63_22g738c | calpain-like cysteine peptidase | -0,666649282 | 1,66337E-10 | C4B63_17g101 | mitochondrial carrier protein | 0,526943063 | 0,022838453 |
| C4B63_44g179 | RNA-binding protein | -0,665658482 | 0,036533312 | C4B63_2g43 | short-chain dehydrogenase | 0,52630464 | 0,011749943 |
| C4B63_1g1255 | trans-sialidase, Group V | -0,664202507 | 0,017625445 | C4B63_41g251 | arginine N-methyltransferase | 0,524972825 | 0,0178062 |
| C4B63_417g21 | unspecified product | -0,659228428 | 0,035607276 | C4B63_79g20 | conserved hypothetical protein | 0,522924594 | 0,017625445 |
| C4B63_39g138 | surface protease GP63 | -0,658175321 | 0,018449635 | C4B63_8g467 | NMD3 family | 0,519458137 | 0,00065138 |
| C4B63_351g27c | small myristoylated protein-1 | -0,655559327 | 8,97075E-11 | C4B63_81g21 | Cell division cycle 20.4, cofactor of APC complex | 0,517729381 | 0,013283186 |
| C4B63_211g28 | conserved hypothetical protein | -0,652634654 | 0,019536057 | C4B63_86g2 | chaperone protein DNAj | 0,516095688 | 0,00206211 |
| C4B63_67g15 | trans-sialidase, Group VI | -0,651575213 | 0,048698031 | C4B63_2g251 | chaperone protein DNAj | 0,512563204 | 0,008791658 |
| C4B63_3g766 | Mucin-associated surface protein (MASP) | -0,650292246 | 0,028568534 | C4B63_112g19 | proteasome beta 3 subunit | 0,511073068 | 0,034200193 |
| C4B63_1g792 | unspecified product | -0,649102664 | 0,0280409 | C4B63_10g167 | conserved hypothetical protein | 0,509960221 | 0,027216143 |
| C4B63_8g240 | trans-sialidase, Group VI | -0,648632256 | 0,035687063 | C4B63_49g166 | alanyl-tRNA synthetase | 0,509604378 | 7,37971E-05 |
| C4B63_9g383 | conserved hypothetical protein | -0,645497274 | 0,000408049 | C4B63_20g747c | cyclin-dependent kinases regulatory subunit | 0,507569221 | 0,003926179 |
| C4B63_66g170 | Mucin-associated surface protein (MASP) | -0,643845448 | 0,023991967 | C4B63_40g25 | Biotin--acetyl-CoA-carboxylase ligase | 0,504234664 | 0,020247783 |
| C4B63_29g79 | unspecified product | -0,639421072 | 0,023194212 | C4B63_42g187 | conserved hypothetical protein | 0,503695039 | 0,016698569 |
| C4B63_70g82 | protein kinase | -0,633638637 | 0,000178439 | C4B63_51g209 | nucleoside 2-deoxyribosyltransferase | 0,502615965 | 0,013048836 |
| C4B63_15g5 | zinc-finger CCCH domain containing protein 11 | -0,629522983 | 0,000813155 | C4B63_79g37 | methionyl-tRNA synthetase | 0,498496659 | 0,031724256 |
| C4B63_13g208 | conserved hypothetical protein,zinc finger protein family member | -0,628165956 | 0,005031804 | C4B63_21g281 | DNA repair and recombination helicase protein PIF5 | 0,498269516 | 0,021762787 |
| C4B63_34g122 | Trypomastigote, Alanine, Serine and Valine rich protein (TASV), subfamily C | -0,627936083 | 0,003581637 | C4B63_52g28 | catalytic subunit of the vacuolar transporter chaperone 4(fragmen | 0,498133577 | 0,003326697 |
| C4B63_52g153 | conserved hypothetical protein | -0,624241045 | 0,024265716 | C4B63_56g42 | conserved hypothetical protein | 0,497312013 | 0,04684336 |
| C4B63_29g448 | Mucin-associated surface protein (MASP) | -0,623432148 | 0,027554188 | C4B63_2g486 | Serine-threonine kinase receptor-associated protein | 0,496876605 | 0,027357346 |
| C4B63_4g242 | protein kinase A catalytic subunit 3 | -0,622576763 | 0,015999194 | C4B63_20g212 | enoyl-CoA hydratase/isomerase family protein | 0,496650355 | 0,00045011 |
| C4B63_181g36 | mismatch repair protein MSH4 | -0,621635062 | 0,001983533 | C4B63_9g326 | ubiquitin-conjugating enzyme | 0,496462046 | 0,034308654 |
| C4B63_323g5 | unspecified product | -0,621612344 | 0,046068546 | C4B63_18g61 | conserved hypothetical protein | 0,493433683 | 0,03451358 |
| C4B63_397g11 | zinc-finger CCCH domain containing protein 11 | -0,620718374 | 0,000585906 | C4B63_18g141 | MORN repeat-containing protein 1 | 0,490577449 | 0,002326654 |
| C4B63_66g68 | trans-sialidase, Group V | -0,61925679 | 0,036409727 | C4B63_113g34 | ABC transporter | 0,488832743 | 0,00569126 |
| C4B63_16g181 | trans-sialidase, Group VIII | -0,617961914 | 0,014873661 | C4B63_114g25 | conserved hypothetical protein | 0,4880448 | 0,019274678 |
| C4B63_5g648 | unspecified product | -0,617748552 | 0,049378282 | C4B63_84g49 | RNA-binding protein | 0,487288063 | 0,019247764 |
| C4B63_47g122 | trans-sialidase | -0,614710119 | 0,026708131 | C4B63_308g4 | kinetoplastid kinetochore protein 8 | 0,487017706 | 0,014562048 |
| C4B63_4g545 | RNA-binding protein 4 | -0,613587711 | 0,026745416 | C4B63_70g104 | Ribosome biogenesis protein BOP1 homolog | 0,485963173 | 0,002147315 |
| C4B63_181g49 | mismatch repair protein MSH4(fragment) | -0,61339406 | 0,049435869 | C4B63_353g8 | unspecified product | 0,485520065 | 0,018969712 |
| C4B63_3g1037 | trans-sialidase, Group V | -0,610026059 | 0,039582611 | C4B63_41g139 | conserved hypothetical protein | 0,485163312 | 0,021655529 |
| C4B63_22g320 | phosphate-repressible phosphate permease | -0,608253765 | 0,002065098 | C4B63_81g22 | dolichyl-P-Man:GDP-Man5GlcNAc2-PP-dolichyl alpha-1,3-manno | 0,485103983 | 0,000474728 |
| C4B63_16g95 | conserved hypothetical protein | -0,608088265 | 0,015705403 | C4B63_40g159 | translation initiation factor 2 subunit | 0,48423633 | 0,032122551 |
| C4B63_19g141 | electron transfer flavoprotein | -0,606089601 | 0,001010325 | C4B63_19g82 | RNA polymerase I second largest subunit | 0,484088067 | 0,022049182 |
| C4B63_13g332 | RNA-binding protein | -0,604069609 | 0,035157974 | C4B63_4g144 | protein transport protein SEC13 | 0,484046006 | 0,013908372 |
| C4B63_50g227 | Mucin-associated surface protein (MASP) | -0,60128855 | 0,036143192 | C4B63_2g217 | PGAP1-like protein/Alpha/beta hydrolase family | 0,484027632 | 0,002953282 |
| C4B63_27g184 | integral membrane transport protein | -0,600300724 | 0,000479068 | C4B63_92g48 | squalene synthase | 0,483191316 | 0,018648042 |
| C4B63_6g587 | N-glycosidase YbiA | -0,599362701 | 0,017486004 | C4B63_2g510 | arginyl-tRNA synthetase | 0,481395008 | 0,002323178 |
| C4B63_419g2 | trans-sialidase | -0,597676557 | 0,024417979 | C4B63_7g421 | coatomer alpha subunit | 0,480911068 | 0,00614814 |
| C4B63_6g611 | unspecified product | -0,594107486 | 0,010184858 | C4B63_63g12 | protein disulfide isomerase 2 | 0,478142876 | 0,035717302 |
| C4B63_11g343 | Mucin-associated surface protein (MASP) | -0,590132282 | 0,016863748 | C4B63_14g204 | 60S acidic ribosomal protein | 0,47795789 | 0,039118382 |
| C4B63_147g63 | Mucin-associated surface protein (MASP) | -0,589205026 | 0,003860385 | C4B63_19g205 | glycosomal transporter (GAT3) | 0,477626853 | 1,64772E-07 |
| C4B63_427g3 | surface protease GP63 | -0,589141794 | 0,011134476 | C4B63_14g148 | NADH dehydrogenase | 0,476425301 | 6,49989E-05 |
| C4B63_13g256 | conserved hypothetical protein | -0,589025276 | 0,022838453 | C4B63_2g289 | Hypoxia up-regulated protein 1 (Fragment) | 0,476177303 | 0,004301655 |
| C4B63_1g691 | Mucin-associated surface protein (MASP) | -0,587165626 | 0,014593677 | C4B63_57g98 | succinyl-CoA ligase | 0,475200675 | 0,041713463 |
| C4B63_260g23 | trans-sialidase, Group II | -0,587068188 | 0,001983576 | C4B63_6g19 | NAD(P)-dependent steroid dehydrogenase protein | 0,474772525 | 0,028438538 |
| C4B63_60g113 | unspecified product | -0,587042482 | 0,030234176 | C4B63_4g118 | conserved hypothetical protein | 0,474138453 | 0,016473042 |
| C4B63_3g65 | Mucin-associated surface protein (MASP) | -0,582539708 | 0,045679939 | C4B63_14g46 | ABC1 family protein YPL109C, mitochondrial | 0,473541536 | 0,008586297 |
| C4B63_5g26 | trans-sialidase, Group V | -0,582069127 | 0,036035045 | C4B63_81g96 | conserved hypothetical protein | 0,473249526 | 0,028438538 |
| C4B63_149g38 | unspecified product | -0,579651127 | 0,035540043 | C4B63_45g97 | eukaryotic translation initiation factor 3 subunit h | 0,47253608 | 0,007194625 |
| C4B63_3g207 | Mucin-associated surface protein (MASP) | -0,578300302 | 0,030930299 | C4B63_39g1477c | 40S ribosomal protein S27 | 0,471933607 | 0,02731757 |
| C4B63_41g261 | trans-sialidase, Group IV | -0,576353102 | 0,00353577 | C4B63_4g196 | conserved hypothetical protein | 0,470687861 | 0,014445349 |
| C4B63_102g58 | mismatch repair protein MSH4(fragment) | -0,57630155 | 0,000388484 | C4B63_10g106 | kinesin | 0,469010572 | 0,000811106 |
| C4B63_22g773c | small myristoylated protein-1 | -0,576067431 | 5,82486E-09 | C4B63_8g485 | glutathione peroxidase-like protein | 0,46885621 | 0,000211234 |
| C4B63_1g946 | unspecified product | -0,575385566 | 0,039148457 | C4B63_114g4 | 40S ribosomal protein S15 | 0,468341086 | 0,024124458 |
| C4B63_78g32 | conserved hypothetical protein | -0,574255011 | 0,044569796 | C4B63_7g318 | peroxisome biogenesis factor 1 | 0,46813526 | 0,016239705 |
| C4B63_43g74 | retrotransposon hot spot (RHS) protein | -0,573325468 | 0,003420563 | C4B63_168g32 | conserved hypothetical protein | 0,467657032 | 0,025026692 |
| C4B63_16g35 | unspecified product | -0,572971898 | 0,027026108 | C4B63_114g16 | conserved hypothetical protein | 0,464697673 | 0,000913 |
| C4B63_20g203 | Zinc finger protein, C3H1 type-like | -0,572304943 | 0,011647401 | C4B63_132g317c | 40S ribosomal protein S27 | 0,464398461 | 0,026261386 |
| C4B63_52g65 | Protein LOL4 | -0,572061853 | 0,0013737 | C4B63_69g73 | fatty acid desaturase | 0,463628772 | 0,001144485 |
| C4B63_19g227 | calpain-like cysteine peptidase | -0,570091785 | 0,021451877 | C4B63_516g5 | retrotransposon hot spot (RHS) protein | 0,46142102 | 0,046101072 |
| C4B63_43g166 | conserved hypothetical protein | -0,569162084 | 0,027652386 | C4B63_163g18 | kinesin | 0,460880629 | 0,048340424 |
| C4B63_14g106 | conserved hypothetical protein | -0,56521654 | 6,86062E-06 | C4B63_13g169 | L-ribulokinase | 0,460080934 | 0,004179992 |
| C4B63_235g1 | rhodanese-like domain containing protein | -0,556261235 | 0,031345961 | C4B63_6g161 | Eukaryotic initiation factor 4A-1 | 0,458325682 | 0,005201504 |
| C4B63_21g17 | arginine permease(fragment) | -0,55618565 | 0,018647063 | C4B63_30g178 | conserved hypothetical protein | 0,456236869 | 0,003569673 |
| C4B63_41g248 | cyclin-like F-box protein | -0,553846716 | 0,011081989 | C4B63_30g260 | conserved hypothetical protein | 0,455861559 | 0,034612802 |
| C4B63_47g125 | trans-sialidase | -0,552355907 | 0,016336897 | C4B63_114g29 | translocon-associated protein (TRAP) alpha subunit | 0,45567202 | 0,005498099 |
| C4B63_3g355 | Mucin-associated surface protein (MASP) | -0,552202843 | 0,049417438 | C4B63_18g211 | conserved hypothetical protein | 0,454350163 | 0,017704083 |
| C4B63_25g220 | 5'-3' exoribonuclease B | -0,55169411 | 0,001083628 | C4B63_4g72 | fibrillarin | 0,453123097 | 0,044508708 |
| C4B63_203g11 | dispersed gene family protein 1 (DGF-1) | -0,550432876 | 0,003282065 | C4B63_233g13 | conserved hypothetical protein | 0,452367462 | 0,018264351 |
| C4B63_4g132 | NLI interacting factor-like phosphatase | -0,547694747 | 0,032299244 | C4B63_19g134 | predicted WD40 repeat protein | 0,451943149 | 0,00084102 |
| C4B63_45g173 | unspecified product | -0,547168646 | 9,57845E-06 | C4B63_151g36 | notchless homolog | 0,451383597 | 0,027924005 |
| C4B63_37g106 | trans-sialidase, Group VI | -0,544205493 | 0,03296846 | C4B63_116g13 | Nucleolar complex protein 2 homolog | 0,450922792 | 0,00468062 |
| C4B63_13g331 | RNA-binding protein | -0,543995174 | 0,045985567 | C4B63_18g230 | conserved hypothetical protein | 0,450705438 | 0,000208154 |
| C4B63_1g1161 | trans-sialidase, Group II | -0,543791675 | 0,021616298 | C4B63_81g41 | Suppressor of disruption of TFIIS | 0,450700667 | 0,013963358 |
| C4B63_98g20 | trans-sialidase, Group IV | -0,541916325 | 0,018651996 | C4B63_22g173 | coatomer beta subunit | 0,450162543 | 0,003677323 |
| C4B63_14g43 | centrin-3 | -0,540729099 | 0,00353577 | C4B63_235g10 | conserved hypothetical protein | 0,44967799 | 0,000439412 |
| C4B63_36g170 | unspecified product | -0,539066767 | 0,019759994 | C4B63_268g10 | conserved hypothetical protein | 0,448635692 | 0,002286846 |
| C4B63_5g424 | Mucin-associated surface protein (MASP) | -0,537234226 | 0,009233534 | C4B63_2g331 | polyadenylate-binding protein 1 | 0,448229114 | 6,25367E-05 |
| C4B63_36g84 | unspecified product | -0,537068527 | 0,013264128 | C4B63_2g349 | Present in the outer mitochondrial membrane proteome 5 | 0,447174769 | 0,002746537 |
| C4B63_166g26 | trans-sialidase, Group VIII | -0,535157469 | 0,001899571 | C4B63_353g7 | unspecified product | 0,447119306 | 0,018417804 |
| C4B63_278g3 | conserved hypothetical protein | -0,534067318 | 0,016135174 | C4B63_30g193 | Phosphoethanolamine/phosphocholine phosphatase | 0,446872782 | 0,000408049 |
| C4B63_15g218 | Mucin-associated surface protein (MASP) | -0,53174621 | 0,045782874 | C4B63_61g118 | conserved hypothetical protein | 0,446302264 | 0,028690559 |
| C4B63_122g12 | Domain of unknown function (DUF4464) | -0,52337901 | 0,048908903 | C4B63_335g2 | conserved hypothetical protein | 0,445570287 | 0,004640321 |
| C4B63_79g10 | mismatch repair protein MSH4 | -0,521465151 | 0,008526442 | C4B63_163g20 | conserved hypothetical protein | 0,443278734 | 0,022838453 |
| C4B63_4g283 | conserved hypothetical protein | -0,520793162 | 0,001507046 | C4B63_4g219 | nucleolar RNA-associated proteins (Nrap) | 0,443058321 | 0,006088918 |
| C4B63_137g3 | DNA repair and recombination helicase protein PIF1 | -0,519892372 | 0,001562913 | C4B63_116g5 | P-type H+-ATPase | 0,442992343 | 0,00206211 |
| C4B63_68g91 | trans-sialidase, Group VIII | -0,518403059 | 0,032047901 | C4B63_8g517 | conserved hypothetical protein | 0,44222381 | 0,002320656 |
| C4B63_45g139 | serine/threonine protein kinase | -0,517504917 | 0,016427633 | C4B63_14g55 | hook complex protein, conserved | 0,442002977 | 0,002200805 |
| C4B63_246g16 | conserved hypothetical protein | -0,515173843 | 0,012221673 | C4B63_7g332 | adenosine kinase | 0,441882777 | 0,019003374 |
| C4B63_35g612c | conserved hypothetical protein | -0,512708415 | 0,034359061 | C4B63_7g223 | eukaryotic translation initiation factor 3 subunit g | 0,44184057 | 0,002784143 |
| C4B63_73g74 | retrotransposon hot spot (RHS) protein | -0,512336643 | 0,011405621 | C4B63_22g312 | phosphoglycerate kinase | 0,439564434 | 0,024607989 |
| C4B63_37g122 | trans-sialidase, Group V | -0,512028401 | 0,029534677 | C4B63_8g62 | Nucleolar protein 10 | 0,438393099 | 0,038555382 |
| C4B63_10g424 | conserved hypothetical protein | -0,511987226 | 0,020412065 | C4B63_2g395 | kinesin | 0,43803712 | 0,028690559 |
| C4B63_97g13 | conserved hypothetical protein | -0,508246712 | 6,48846E-05 | C4B63_57g74 | conserved hypothetical protein | 0,437847743 | 0,046068546 |
| C4B63_53g165 | conserved hypothetical protein | -0,507604441 | 0,013899202 | C4B63_9g324 | conserved hypothetical protein | 0,436774831 | 0,002507716 |
| C4B63_109g60 | trans-sialidase, Group IV | -0,501879585 | 0,005195667 | C4B63_56g31 | conserved hypothetical protein | 0,436508571 | 0,009760243 |
| C4B63_26g309 | ABC transporter(fragment) | -0,50156996 | 0,018648042 | C4B63_52g54 | conserved hypothetical protein | 0,435788537 | 0,036143192 |
| C4B63_76g17 | conserved hypothetical protein | -0,501261035 | 0,001640006 | C4B63_23g23 | conserved hypothetical protein | 0,435665898 | 0,017657249 |
| C4B63_45g193 | unspecified product | -0,501143386 | 0,019841527 | C4B63_20g226 | Probable importin subunit beta-4 | 0,435576468 | 0,019430852 |
| C4B63_52g43 | protein phosphatase 2C | -0,500262179 | 0,005649036 | C4B63_63g37 | oligopeptidase b(fragment) | 0,434320164 | 0,000294927 |
| C4B63_99g112 | alanine racemase | -0,500163 | 0,005870281 | C4B63_286g18 | eukaryotic translation initiation factor 3 subunit c | 0,432905973 | 0,026099081 |
| C4B63_127g46 | conserved hypothetical protein | -0,497287182 | 0,047573709 | C4B63_1g70 | mitochondrial RNA-binding protein 2 | 0,432471105 | 0,034393341 |
| C4B63_15g385 | dispersed gene family protein 1 (DGF-1) | -0,496991266 | 0,026898793 | C4B63_18g152 | conserved hypothetical protein | 0,431642348 | 0,041713463 |
| C4B63_24g274 | conserved hypothetical protein | -0,495040544 | 0,046453815 | C4B63_44g234 | CRAL/TRIO domain containing protein | 0,429686227 | 0,014562362 |
| C4B63_225g35 | unspecified product | -0,49382312 | 0,010188448 | C4B63_6g489 | Something about silencing protein 10 | 0,429053307 | 0,020445396 |
| C4B63_238g9 | conserved hypothetical protein | -0,493579953 | 0,010407949 | C4B63_25g231 | conserved hypothetical protein | 0,428251336 | 0,000766972 |
| C4B63_44g235 | Kinesin-13 4 | -0,493200302 | 0,005511543 | C4B63_57g119 | conserved hypothetical protein | 0,427540145 | 0,04909789 |
| C4B63_6g572 | 5'-3' exoribonuclease A | -0,493034456 | 0,023147956 | C4B63_54g169 | conserved hypothetical protein | 0,427060184 | 0,048403964 |
| C4B63_12g151 | conserved hypothetical protein | -0,489001117 | 0,018449635 | C4B63_2g509 | arginyl-tRNA synthetase | 0,426291096 | 0,005947233 |
| C4B63_13g327 | conserved hypothetical protein | -0,488346675 | 0,033612624 | C4B63_297g14 | phenylalanyl-tRNA synthetase alpha chain | 0,424876236 | 0,003474947 |
| C4B63_39g347 | conserved hypothetical protein | -0,486668078 | 0,035282579 | C4B63_10g1870c | conserved hypothetical protein | 0,424548898 | 0,022753296 |
| C4B63_312g23 | mismatch repair protein MSH4 | -0,484651152 | 0,018427564 | C4B63_128g58 | basal body component | 0,423558989 | 0,032831512 |
| C4B63_44g231 | amino acid transporter | -0,484143227 | 0,032912013 | C4B63_6g445 | prenyltransferase | 0,423160425 | 0,005174677 |
| C4B63_94g84 | dispersed gene family protein 1 (DGF-1) | -0,482648176 | 0,045409958 | C4B63_6g279 | Coatomer subunit gamma | 0,422768378 | 0,013677847 |
| C4B63_19g218 | calpain-like cysteine peptidase | -0,481538192 | 0,010805506 | C4B63_75g68 | calcium motive p-type ATPase | 0,422321077 | 0,031849818 |
| C4B63_8g495 | Regulator of chromosome condensation (RCC1) repeat | -0,478091699 | 0,017486004 | C4B63_9g314 | DNA replication licensing factor MCM3 | 0,420020624 | 0,001461073 |
| C4B63_168g15 | mismatch repair protein MSH4 | -0,477874367 | 0,000766972 | C4B63_322g14 | amino acid transporter | 0,419529716 | 0,012624509 |
| C4B63_301g2 | conserved hypothetical protein | -0,476736644 | 0,041713463 | C4B63_14g210 | WD repeat-containing protein 55 | 0,417757661 | 0,047253514 |
| C4B63_171g31 | choline/carnitine O-acyltransferase | -0,47511537 | 0,01428227 | C4B63_2g311 | conserved hypothetical protein | 0,417063735 | 0,000309296 |
| C4B63_44g183 | conserved hypothetical protein | -0,474972141 | 0,006510894 | C4B63_348g21 | ADP-ribosylation factor-like protein 3 | 0,416318009 | 0,047573709 |
| C4B63_24g275 | conserved hypothetical protein | -0,474848749 | 0,043259471 | C4B63_32g30 | translationally controlled tumor protein (TCTP) | 0,415409061 | 0,005870281 |
| C4B63_8g466 | protein kinase | -0,474582744 | 0,004774203 | C4B63_55g171 | kinesin | 0,414983572 | 0,002215459 |
| C4B63_5g698 | Mucin-associated surface protein (MASP) | -0,473044979 | 0,048403964 | C4B63_49g78 | conserved hypothetical protein | 0,414247145 | 0,027490566 |
| C4B63_97g15 | Zinc finger CCCH domain-containing protein 35 | -0,471818972 | 0,016204479 | C4B63_11g63 | D-alanine--D-alanine ligase | 0,414238672 | 0,017447632 |
| C4B63_70g87 | NADH-cytochrome b5 reductase | -0,471470344 | 0,011783595 | C4B63_22g108 | Replication Factor C Subunit 1-related protein | 0,413618603 | 0,049520567 |
| C4B63_44g229 | amino acid permease | -0,471315434 | 5,00548E-05 | C4B63_38g147 | spermidine synthase | 0,413525397 | 0,041938564 |
| C4B63_380g14 | conserved hypothetical protein | -0,469784544 | 0,009420721 | C4B63_6g118 | Mitochondrial import receptor subunit ATOM46 | 0,413326375 | 0,039810592 |
| C4B63_18g308 | sphingosine 1-phosphate lyase | -0,469536505 | 0,006551143 | C4B63_32g239 | tubulin folding cofactor D | 0,413000563 | 0,002914934 |
| C4B63_13g388 | Mucin-associated surface protein (MASP) | -0,466427701 | 0,0280409 | C4B63_23g199 | Flagellum attachment zone protein 19 | 0,411000772 | 0,007280475 |
| C4B63_12g175 | Protein Associated with Differentiation | -0,466249843 | 0,007730001 | C4B63_12g385 | Acyl carrier protein, mitochondrial | 0,410297182 | 0,030957158 |
| C4B63_16g142 | trans-sialidase, Group II | -0,466073759 | 0,022623344 | C4B63_2g244 | conserved hypothetical protein | 0,409630016 | 0,007290437 |
| C4B63_26g310 | ABC transporter | -0,465656509 | 0,030957158 | C4B63_30g187 | deoxyuridine triphosphatase | 0,40945932 | 0,003626823 |
| C4B63_20g312 | conserved hypothetical protein | -0,465439699 | 0,001552337 | C4B63_2g566 | phenylalanyl-tRNA synthetase | 0,408407001 | 0,031706884 |
| C4B63_32g51 | Failed axon connections homolog | -0,465226692 | 8,4158E-06 | C4B63_10g498 | conserved hypothetical protein | 0,408156074 | 0,030957158 |
| C4B63_25g236 | conserved hypothetical protein | -0,464641005 | 0,044320928 | C4B63_355g21 | conserved hypothetical protein | 0,407912977 | 0,028983564 |
| C4B63_218g22 | conserved hypothetical protein | -0,463357047 | 0,035540043 | C4B63_13g271 | fructose-bisphosphate aldolase, glycosomal | 0,406926353 | 0,000438924 |
| C4B63_24g225 | serine/threonine protein phosphatase | -0,461426564 | 0,036685181 | C4B63_2g305 | Hsp70-Hsp90 organizing protein 3 | 0,406846252 | 0,024285996 |
| C4B63_32g209 | conserved hypothetical protein | -0,454983797 | 0,004179992 | C4B63_38g45 | conserved hypothetical protein | 0,405345158 | 0,031115141 |
| C4B63_10g443 | conserved hypothetical protein | -0,453798424 | 0,014394483 | C4B63_35g130 | orotidine-5-phosphate decarboxylase/orotate phosphoribosyltrans | 0,404481855 | 0,004710586 |
| C4B63_32g53 | conserved hypothetical protein | -0,453413302 | 0,006937298 | C4B63_14g39 | conserved hypothetical protein | 0,404084216 | 0,035363206 |
| C4B63_16g93 | Uncharacterized protein in mobD 3'region | -0,45243949 | 0,031682345 | C4B63_30g253 | conserved hypothetical protein | 0,403580363 | 0,037802648 |
| C4B63_47g154 | dispersed gene family protein 1 (DGF-1) | -0,450703506 | 0,023088161 | C4B63_28g180 | WASH complex subunit 4 | 0,403435344 | 0,046102101 |
| C4B63_9g313 | DnaJ chaperone protein | -0,450241293 | 0,036143192 | C4B63_19g29 | vesicular-fusion ATPase-like protein | 0,403039787 | 0,02114448 |
| C4B63_246g14 | conserved hypothetical protein | -0,449072743 | 0,028748767 | C4B63_58g143 | Lupus La protein homolog | 0,402769488 | 0,018449635 |
| C4B63_14g20 | hexose transporter | -0,449019519 | 0,009175913 | C4B63_3g1120 | conserved hypothetical protein | 0,401960892 | 0,002600842 |
| C4B63_246g15 | conserved hypothetical protein | -0,448036551 | 0,03829681 | C4B63_123g22 | fatty acid desaturase | 0,401723413 | 0,00851259 |
| C4B63_45g174 | unspecified product | -0,447265777 | 0,018836287 | C4B63_20g179 | CYC2-like cyclin | 0,40146757 | 0,002449243 |
| C4B63_25g189 | 3'5'-cyclic nucleotide phosphodiesterase | -0,447224633 | 0,017065484 | C4B63_13g270 | fructose-bisphosphate aldolase, glycosomal | 0,401454738 | 0,001279679 |
| C4B63_47g162 | retrotransposon hot spot (RHS) protein | -0,446982846 | 0,042950925 | C4B63_80g12 | RING-variant domain containing protein | 0,401406562 | 0,008107256 |
| C4B63_4g511 | zinc-finger protein | -0,446701308 | 0,048908903 | C4B63_23g255 | leucine-rich repeat protein (LRRP) | 0,401195859 | 0,026583072 |
| C4B63_16g33 | trans-sialidase, Group VIII | -0,445196026 | 0,030659328 | C4B63_23g112 | Helicase required for RNAi-mediated heterochromatin assembly 1 | 0,400812833 | 0,017459093 |
| C4B63_134g59 | trans-sialidase, Group V | -0,44484198 | 0,026137749 | C4B63_10g510 | 26S proteasome regulatory non-ATPase subunit | 0,400297642 | 0,00723211 |
| C4B63_308g6 | conserved hypothetical protein | -0,444210566 | 0,013048836 | | | | |
| C4B63_2g776 | conserved hypothetical protein | -0,442903259 | 0,034208075 | | | | |
| C4B63_45g175 | unspecified product | -0,441754216 | 0,008112206 | | | | |
| C4B63_314g7 | conserved hypothetical protein | -0,439411481 | 0,000551553 | | | | |
| C4B63_23g16 | Battenin | -0,438864703 | 0,032578847 | | | | |
| C4B63_4g509 | FKBP-type peptidylprolyl cis-trans isomerase | -0,437458055 | 0,001465609 | | | | |
| C4B63_42g60 | amastin | -0,43656745 | 0,001564837 | | | | |
| C4B63_44g227 | amino acid permease | -0,435546652 | 0,000389849 | | | | |
| C4B63_10g416 | conserved hypothetical protein | -0,435264038 | 0,033785049 | | | | |
| C4B63_57g110 | conserved hypothetical protein | -0,434949277 | 0,016176667 | | | | |
| C4B63_75g113 | retrotransposon hot spot protein (RHS) | -0,434940924 | 0,045782874 | | | | |
| C4B63_45g80 | conserved hypothetical protein | -0,434729463 | 0,04277622 | | | | |
| C4B63_18g326 | zinc-finger protein ZFP1 | -0,434084157 | 0,021762787 | | | | |
| C4B63_128g242c | dynein light chain | -0,433384672 | 0,01414948 | | | | |
| C4B63_49g109 | protein kinase | -0,431918309 | 0,045782874 | | | | |
| C4B63_26g240 | nucleobase transporter | -0,430711168 | 0,019437772 | | | | |
| C4B63_51g83 | ferric reductase transmembrane protein | -0,430435018 | 0,00405047 | | | | |
| C4B63_95g6 | trans-sialidase, Group V | -0,430190565 | 0,019143156 | | | | |
| C4B63_18g322 | conserved hypothetical protein | -0,428500285 | 0,011405621 | | | | |
| C4B63_189g32 | conserved hypothetical protein | -0,426991806 | 0,012592775 | | | | |
| C4B63_69g77 | protein kinase | -0,425185789 | 0,011558596 | | | | |
| C4B63_212g8 | conserved hypothetical protein | -0,425034461 | 0,004638435 | | | | |
| C4B63_180g10 | retrotransposon hot spot (RHS) protein | -0,423357285 | 0,031274381 | | | | |
| C4B63_70g84 | serine/threonine protein kinase | -0,421733611 | 0,036409727 | | | | |
| C4B63_51g80 | ferric reductase transmembrane protein | -0,419081955 | 0,000872311 | | | | |
| C4B63_2g33 | conserved hypothetical protein | -0,414044128 | 0,039714977 | | | | |
| C4B63_255g15 | UDP-Gal or UDP-GlcNAc-dependent glycosyltransferase | -0,413508827 | 0,034407072 | | | | |
| C4B63_44g218 | conserved hypothetical protein | -0,41311994 | 0,021040674 | | | | |
| C4B63_17g85 | conserved hypothetical protein | -0,412885743 | 0,009206136 | | | | |
| C4B63_10g420 | conserved hypothetical protein | -0,411555506 | 0,039917965 | | | | |
| C4B63_47g102 | cytochrome c oxidase assembly protein | -0,410476915 | 0,035540043 | | | | |
| C4B63_17g196 | calpain-like cysteine peptidase | -0,410180149 | 0,022838453 | | | | |
| C4B63_2g156 | protein kinase A catalytic subunit 3 | -0,409506577 | 0,013340872 | | | | |
| C4B63_6g273 | DNA repair and recombination helicase protein PIF1 | -0,409293535 | 0,01612218 | | | | |
| C4B63_57g121 | meiosis recombination protein SPO11 | -0,406988729 | 0,034308654 | | | | |
| C4B63_4g335 | conserved hypothetical protein | -0,404274081 | 0,007899072 | | | | |
| C4B63_208g40 | conserved hypothetical protein | -0,404052245 | 0,039917965 | | | | |
| C4B63_5g787 | Mucin-associated surface protein (MASP) | -0,403907225 | 0,046896312 | | | | |
| C4B63_18g169 | Zinc finger CCCH domain-containing protein 47 | -0,4020007 | 0,021120659 | | | | |
| C4B63_51g82 | ferric reductase transmembrane protein | -0,401274954 | 0,002914934 | | | | |
| C4B63_40g93 | conserved hypothetical protein | -0,400973549 | 0,005285726 | | | | |

#### Hb exclusive DEGs (t4 vs. t0)

##### Downregulated genes

| ID | Annotation | log2FC | padj |
|---|---|---|---|
| C4B63_43g863c | Protein BOLA1, chloroplastic | -1,240155828 | 0,046271108 |
| C4B63_244g7 | trans-sialidase, Group I | -0,908535133 | 0,005027624 |
| C4B63_20g344 | conserved hypothetical protein | -0,824485449 | 0,03088001 |
| C4B63_235g1 | rhodanese-like domain containing protein | -0,822341523 | 0,000669743 |
| C4B63_244g6 | trans-sialidase, Group I | -0,792707358 | 0,039427911 |
| C4B63_99g109 | 2OG-Fe(II) oxygenase superfamily | -0,763726033 | 0,00440518 |
| C4B63_84g51 | Coiled-coil domain-containing protein 40 homolog | -0,721547989 | 0,005545793 |
| C4B63_19g175 | Pleckstrin homology domain | -0,712245684 | 0,01351841 |
| C4B63_7g246 | Patatin-like phospholipase domain-containing protein 3 | -0,70464162 | 0,005758503 |
| C4B63_17g109 | BCL-6 corepressor | -0,699200892 | 0,033998151 |
| C4B63_233g27c | conserved hypothetical protein | -0,690051554 | 0,039893396 |
| C4B63_43g123 | conserved hypothetical protein | -0,683464229 | 0,009561388 |
| C4B63_112g64 | conserved hypothetical protein | -0,669165384 | 0,013804188 |
| C4B63_84g19 | conserved hypothetical protein | -0,668550004 | 0,031620556 |
| C4B63_69g19 | exopolyphosphatase | -0,665276472 | 0,039411221 |
| C4B63_442g10 | Uncharacterized protein C4orf22 homolog | -0,662668403 | 0,040386832 |
| C4B63_3g1089 | Alpha-ketoglutarate-dependent dioxygenase AlkB-like superfamily | -0,659607611 | 0,007816958 |
| C4B63_25g29 | trans-sialidase, Group I | -0,650846053 | 0,04703967 |
| C4B63_26g9 | conserved hypothetical protein | -0,646587215 | 0,009361249 |
| C4B63_170g37 | conserved hypothetical protein | -0,64408542 | 0,042412919 |
| C4B63_7g258 | conserved hypothetical protein | -0,637932517 | 0,023795098 |
| C4B63_27g3 | ribonucleoside-diphosphate reductase small chain | -0,629598731 | 0,037630308 |
| C4B63_122g15 | conserved hypothetical protein | -0,62377988 | 0,039512345 |
| C4B63_69g151 | conserved hypothetical protein | -0,614869715 | 0,004821723 |
| C4B63_16g279 | S-adenosyl-L-methionine-dependent methyltransferase | -0,612039104 | 0,017413919 |
| C4B63_124g35 | retrotransposon hot spot (RHS) protein | -0,60941608 | 0,008436548 |
| C4B63_79g19 | Calmodulin-like protein 11 | -0,6094081 | 0,011642421 |
| C4B63_18g795c | zinc-binding protein (Yippee) | -0,60657563 | 0,033160367 |
| C4B63_122g12 | Domain of unknown function (DUF4464) | -0,600072754 | 0,018913079 |
| C4B63_221g21 | conserved hypothetical protein | -0,598743846 | 0,046114862 |
| C4B63_21g22 | arginine permease(fragment) | -0,596267183 | 0,022722583 |
| C4B63_1g128c | Heat shock protein 67Bb | -0,595297904 | 0,005545793 |
| C4B63_2g335 | conserved hypothetical protein | -0,592469651 | 0,040374691 |
| C4B63_101g35 | delta-4 fatty acid desaturase | -0,591160463 | 0,032090825 |
| C4B63_280g10 | prostaglandin F2alpha synthase | -0,589545686 | 0,028605053 |
| C4B63_19g130 | Dual specificity phosphatase, catalytic domain containing protein | -0,588483857 | 0,00295146 |
| C4B63_7g251 | Calcyphosin-like protein | -0,585586871 | 0,030507577 |
| C4B63_117g53 | protein kinase | -0,581379179 | 0,035190538 |
| C4B63_2g4640c | conserved hypothetical protein | -0,578954846 | 0,045587485 |
| C4B63_7g426 | Double RNA binding domain protein 7 | -0,572482818 | 0,022722583 |
| C4B63_17g22 | Ras-related protein Rab4 | -0,56581602 | 0,030082599 |
| C4B63_80g18 | conserved hypothetical protein | -0,554672134 | 0,010158898 |
| C4B63_2g39c | dynein-associated protein | -0,553342941 | 0,03825855 |
| C4B63_4g417 | conserved hypothetical protein | -0,542094502 | 0,020868051 |
| C4B63_1g130c | Thiosulfate:glutathione sulfurtransferase | -0,538268357 | 0,009998568 |
| C4B63_22g10 | Enkuring domain-containig protein | -0,537177543 | 0,030969294 |
| C4B63_18g80 | ferric reductase transmembrane protein | -0,535739528 | 0,020771209 |
| C4B63_11g6 | conserved hypothetical protein | -0,53398733 | 0,01788802 |
| C4B63_12g9 | conserved hypothetical protein | -0,530869838 | 0,022610748 |
| C4B63_4g418 | conserved hypothetical protein | -0,526307197 | 0,01923482 |
| C4B63_23g70 | ubiquinone biosynthesis protein COQ7 homolog | -0,526278895 | 0,034799246 |
| C4B63_17g176 | serine carboxypeptidase S28 | -0,520264245 | 0,034622974 |
| C4B63_7g249 | conserved hypothetical protein | -0,516884742 | 0,010227003 |
| C4B63_49g55 | conserved hypothetical protein | -0,515236282 | 0,031387066 |
| C4B63_6g387 | NADH dehydrogenase subunit NB6M | -0,512377919 | 0,018973873 |
| C4B63_18g202 | conserved hypothetical protein | -0,512090274 | 0,034942732 |
| C4B63_184g6 | conserved hypothetical protein | -0,51180815 | 0,007423155 |
| C4B63_52g313c | calpain-like cysteine peptidase | -0,511344679 | 0,012486342 |
| C4B63_12g182 | PUF nine target 1 | -0,51100523 | 0,047311923 |
| C4B63_4g353 | conserved hypothetical protein | -0,510356299 | 0,03882038 |
| C4B63_18g345 | conserved hypothetical protein | -0,505430353 | 0,004225262 |
| C4B63_113g42 | thiol-dependent reductase 1 | -0,50353573 | 0,000000461 |
| C4B63_10g278 | conserved hypothetical protein | -0,502666865 | 0,030507577 |
| C4B63_2g326 | protein kinase | -0,502600423 | 0,006764196 |
| C4B63_27g162 | conserved hypothetical protein | -0,502502577 | 0,035191466 |
| C4B63_70g65 | conserved hypothetical protein | -0,502091689 | 0,009559678 |
| C4B63_230g10 | A/G-specific adenine glycosylase | -0,498192635 | 0,040794699 |
| C4B63_2g689 | Nucleoside diphosphate kinase 3 | -0,496155507 | 0,039136862 |
| C4B63_8g2705c | conserved hypothetical protein | -0,494716941 | 0,02478701 |
| C4B63_52g312c | calpain-like cysteine peptidase | -0,494506823 | 0,028783305 |
| C4B63_54g167 | conserved hypothetical protein | -0,4939067 | 0,045000081 |
| C4B63_4g243 | cAMP phosphodiesterase A | -0,490477286 | 0,021842808 |
| C4B63_127g27 | Cilia- and flagella-associated protein 43 | -0,490216582 | 0,014797999 |
| C4B63_9g461 | conserved hypothetical protein | -0,488721406 | 0,040957824 |
| C4B63_7g372 | dynein heavy chain | -0,487862576 | 0,010865466 |
| C4B63_7g430 | 1,2-Dihydroxy-3-keto-5-methylthiopentene dioxygenase | -0,486247686 | 0,013638521 |
| C4B63_405g4 | glutamate dehydrogenase | -0,481596514 | 0,015825719 |
| C4B63_17g97 | conserved hypothetical protein | -0,480296557 | 0,010312581 |
| C4B63_12g24 | frataxin-like, mitochondrial precursor | -0,478187229 | 0,007045433 |
| C4B63_14g59 | conserved hypothetical protein | -0,476367666 | 0,000744325 |
| C4B63_31g116 | conserved hypothetical protein | -0,47625606 | 0,020882981 |
| C4B63_20g157 | NADH dehydrogenase subunit NI8M | -0,475238491 | 0,000604385 |
| C4B63_48g142 | serine/threonine protein kinase | -0,471603767 | 0,000543223 |
| C4B63_6g230 | conserved hypothetical protein | -0,470926746 | 0,00382844 |
| C4B63_70g114 | conserved hypothetical protein | -0,470254149 | 0,024888252 |
| C4B63_70g115 | protein kinase | -0,468885336 | 0,04222028 |
| C4B63_47g115 | fatty acyl CoA synthetase | -0,466900226 | 0,001201024 |
| C4B63_83g40 | glycine dehydrogenase [decarboxylating](fragment) | -0,46633828 | 0,049293304 |
| C4B63_116g14 | conserved hypothetical protein | -0,46332118 | 0,010158898 |
| C4B63_14g63 | Haloacid dehalogenase-like hydrolase | -0,46273929 | 0,036774946 |
| C4B63_24g165 | protein kinase | -0,461426646 | 0,011748225 |
| C4B63_7g260 | ADP/ATP mitochondrial carrier protein | -0,460657085 | 0,022722583 |
| C4B63_102g69 | katanin | -0,456761908 | 0,002360843 |
| C4B63_11g265 | conserved hypothetical protein | -0,456322449 | 0,004169035 |
| C4B63_248g14 | glucose-6-phosphate isomerase, glycosomal | -0,455948857 | 0,04351471 |
| C4B63_136g35 | conserved hypothetical protein | -0,454900882 | 0,024505347 |
| C4B63_9g333 | cysteine peptidase, Clan CA, family C51 | -0,454252022 | 0,012998286 |
| C4B63_14g209 | conserved hypothetical protein | -0,454137608 | 0,029943886 |
| C4B63_8g561 | conserved hypothetical protein | -0,453691677 | 0,04349771 |
| C4B63_7g438 | glutamate dehydrogenase | -0,452746159 | 0,033548808 |
| C4B63_13g170 | pumilio/PUF RNA binding protein 1 | -0,451985706 | 0,002844022 |
| C4B63_43g190 | Putative RING-H2 finger protein ATL71 | -0,450982055 | 0,033548808 |
| C4B63_18g267 | conserved hypothetical protein | -0,450880374 | 0,046000233 |
| C4B63_6g531 | conserved hypothetical protein | -0,450821969 | 0,004274116 |
| C4B63_12g259 | phosphoinositide-specific phospholipase C | -0,450486051 | 0,003510809 |
| C4B63_14g106 | conserved hypothetical protein | -0,44698023 | 0,001212559 |
| C4B63_2g63 | conserved hypothetical protein | -0,446414405 | 0,031333477 |
| C4B63_70g547c | conserved hypothetical protein | -0,445408471 | 0,038613176 |
| C4B63_233g32 | ubiquitin hydrolase | -0,445165125 | 0,039658983 |
| C4B63_191g32 | Cilia- and flagella-associated protein 43 | -0,443350726 | 0,021797618 |
| C4B63_1g20 | conserved hypothetical protein | -0,441896937 | 0,04349771 |
| C4B63_133g32 | conserved hypothetical protein | -0,441416084 | 0,04999574 |
| C4B63_113g57 | dual specificity protein phosphatase or MAP kinase phosphatase | -0,441299408 | 0,041433691 |
| C4B63_4g376 | NADH-ubiquinone oxidoreductase complex I subunit,FT | -0,44005485 | 0,001189079 |
| C4B63_18g1035c | conserved hypothetical protein | -0,439695078 | 0,049133439 |
| C4B63_43g193 | conserved hypothetical protein | -0,437672678 | 0,041572205 |
| C4B63_227g3 | NADH-ubiquinone oxidoreductase complex I subunit,FT | -0,437663774 | 0,011344441 |
| C4B63_70g87 | NADH-cytochrome b5 reductase | -0,436191996 | 0,03797988 |
| C4B63_55g287c | cytochrome c | -0,43442792 | 0,027698415 |
| C4B63_49g161 | conserved hypothetical protein | -0,433401128 | 0,026700096 |
| C4B63_18g52 | protein kinase | -0,431108105 | 0,036352553 |
| C4B63_120g44 | phosphoinositide-specific phospholipase C | -0,429029573 | 0,015615853 |
| C4B63_41g249 | conserved hypothetical protein | -0,426512571 | 0,009665159 |
| C4B63_39g312 | conserved hypothetical protein | -0,426450687 | 0,033160367 |
| C4B63_57g131 | glutamate dehydrogenase | -0,424993219 | 0,026703969 |
| C4B63_40g74 | conserved hypothetical protein | -0,424587111 | 0,044306022 |
| C4B63_3g1125 | short chain dehydrogenase | -0,422310585 | 0,010158898 |
| C4B63_69g78 | protein kinase | -0,421349043 | 0,025373432 |
| C4B63_289g4 | glutamate dehydrogenase | -0,418895697 | 0,049934421 |
| C4B63_186g5 | conserved hypothetical protein | -0,417698373 | 0,022722583 |
| C4B63_11g480 | conserved hypothetical protein | -0,416125112 | 0,01788802 |
| C4B63_35g55 | flagellar calcium-binding protein | -0,415496632 | 0,021248307 |
| C4B63_17g60 | protein kinase | -0,413756847 | 0,013041359 |
| C4B63_4g211 | conserved hypothetical protein | -0,413510454 | 0,016366894 |
| C4B63_14g155 | Leucine-rich repeat-containing protein 74A | -0,41224649 | 0,02695635 |
| C4B63_7g151 | conserved hypothetical protein | -0,411748023 | 0,00229128 |
| C4B63_12g252 | conserved hypothetical protein | -0,411166235 | 0,003686365 |
| C4B63_47g72 | enolase | -0,410044042 | 0,021004555 |
| C4B63_2g788 | conserved hypothetical protein | -0,40932716 | 0,00277344 |
| C4B63_183g11 | conserved hypothetical protein | -0,407142273 | 0,000691651 |
| C4B63_4g517 | conserved hypothetical protein | -0,407102344 | 0,049133439 |
| C4B63_13g117 | conserved hypothetical protein | -0,405940792 | 0,002365881 |
| C4B63_18g169 | Zinc finger CCCH domain-containing protein 47 | -0,405778172 | 0,021162993 |
| C4B63_7g391 | ubiquitin hydrolase | -0,404246862 | 0,028532938 |
| C4B63_25g235 | Protein OSCP1 | -0,402202918 | 0,033160367 |
| C4B63_6g259 | nucleotidyl transferase | -0,400157258 | 0,018358864 |

##### Upregulated genes

| ID | Annotation | log2FC | padj |
|---|---|---|---|
| C4B63_225g40 | conserved hypothetical protein | 0,748386504 | 0,00510403 |
| C4B63_34g1472c | mucin TcSMUGL | 0,726181544 | 0,003597327 |
| C4B63_10g159 | hypothetical Zinc-finger containing protein | 0,722988221 | 0,012067303 |
| C4B63_46g17 | conserved hypothetical protein | 0,696284952 | 0,007419168 |
| C4B63_40g162 | exosome-associated protein 4 | 0,689623871 | 0,020435922 |
| C4B63_34g1526c | mucin TcSMUGL | 0,677343387 | 0,015410034 |
| C4B63_85g83 | Eukaryotic initiation factor 4A-1 | 0,667834988 | 0,041562037 |
| C4B63_15g15 | oligosaccharyl transferase subunit | 0,648235884 | 0,004714921 |
| C4B63_504g2 | unspecified product | 0,634033759 | 0,035623058 |
| C4B63_202g28 | cysteine peptidase(fragment) | 0,628768225 | 0,021378564 |
| C4B63_188g148c | histone H4 | 0,6274261 | 0,014545101 |
| C4B63_188g147c | histone H4 | 0,615462782 | 0,00395163 |
| C4B63_44g113 | conserved hypothetical protein | 0,615444195 | 0,007571357 |
| C4B63_9g299 | ATP-dependent Clp protease subunit, heat shock protein 104 | 0,615430167 | 0,036290641 |
| C4B63_480g17 | cysteine peptidase | 0,60499424 | 0,028739129 |
| C4B63_188g145c | histone H4 | 0,604799599 | 0,010409787 |
| C4B63_46g16 | conserved hypothetical protein | 0,600830335 | 0,02092142 |
| C4B63_9g416 | ATP-dependent Clp protease subunit, heat shock protein 104 | 0,591979759 | 0,03222943 |
| C4B63_77g10 | unspecified product | 0,587680085 | 0,022371984 |
| C4B63_188g154c | histone H4 | 0,586803851 | 0,025351461 |
| C4B63_295g15 | nucleosome assembly protein-like protein | 0,585782087 | 0,015586429 |
| C4B63_188g132c | histone H4 | 0,580859917 | 0,010983823 |
| C4B63_200g30 | p-nitrophenylphosphatase | 0,580504076 | 0,016021319 |
| C4B63_35g179 | protein tyrosine phosphatase | 0,575632998 | 0,040740429 |
| C4B63_45g84 | conserved hypothetical protein | 0,573125616 | 0,01788802 |
| C4B63_41g159 | cysteine peptidase(fragment) | 0,570377277 | 0,023003551 |
| C4B63_2g782 | ATP-dependent RNA helicase | 0,570366618 | 0,014763793 |
| C4B63_202g15 | cysteine peptidase | 0,570267867 | 0,027130166 |
| C4B63_202g24 | cysteine peptidase | 0,56781098 | 0,028964441 |
| C4B63_70g48 | cystathionine beta-synthase | 0,566823963 | 0,008269429 |
| C4B63_202g17 | cysteine peptidase | 0,56598157 | 0,048207729 |
| C4B63_202g25 | cysteine peptidase | 0,56258199 | 0,032449213 |
| C4B63_78g13 | Proteasome activator complex subunit 4 | 0,56256771 | 0,033792728 |
| C4B63_202g20 | cysteine peptidase | 0,562108552 | 0,033548808 |
| C4B63_202g27 | cysteine peptidase | 0,560811356 | 0,026438959 |
| C4B63_202g16 | cysteine peptidase | 0,558178326 | 0,03207292 |
| C4B63_275g2 | polyubiquitin | 0,557353592 | 0,04703967 |
| C4B63_302g64c | histone H3 | 0,55575431 | 0,030074277 |
| C4B63_89g71 | cysteine peptidase | 0,555518991 | 0,040789709 |
| C4B63_89g67 | cysteine peptidase | 0,553549463 | 0,035191466 |
| C4B63_89g68 | cruzipain precursor | 0,551401798 | 0,035623058 |
| C4B63_41g158 | cysteine peptidase | 0,549770358 | 0,005306485 |
| C4B63_202g29 | cysteine peptidase(fragment) | 0,548707346 | 0,033377916 |
| C4B63_89g70 | cysteine peptidase | 0,547321677 | 0,037826269 |
| C4B63_41g154 | cysteine peptidase | 0,546557898 | 0,004292901 |
| C4B63_202g22 | cysteine peptidase | 0,54490258 | 0,03825855 |
| C4B63_188g138c | histone H4 | 0,544456924 | 0,038296577 |
| C4B63_202g18 | cysteine peptidase | 0,542799488 | 0,038944112 |
| C4B63_302g68c | histone H3 | 0,541564241 | 0,044157687 |
| C4B63_202g19 | cysteine peptidase | 0,540518174 | 0,042398105 |
| C4B63_4g494 | conserved hypothetical protein | 0,536737557 | 0,043676254 |
| C4B63_202g13 | cysteine peptidase | 0,534197282 | 0,037246791 |
| C4B63_292g20 | polyubiquitin | 0,53144424 | 0,046061219 |
| C4B63_42g187 | conserved hypothetical protein | 0,531153535 | 0,015083886 |
| C4B63_178g40 | conserved hypothetical protein | 0,525218731 | 0,041562037 |
| C4B63_202g23 | cysteine peptidase | 0,524719545 | 0,045925699 |
| C4B63_77g15 | dispersed gene family protein 1 (DGF-1) | 0,524481705 | 0,043676254 |
| C4B63_202g30 | cysteine peptidase(fragment) | 0,519534851 | 0,019797475 |
| C4B63_114g34 | conserved hypothetical protein | 0,5154172 | 0,020072185 |
| C4B63_38g165 | conserved hypothetical protein | 0,514381303 | 0,003161092 |
| C4B63_188g159c | histone H4 | 0,512193169 | 0,034520192 |
| C4B63_480g15 | cysteine peptidase | 0,511778035 | 0,027245911 |
| C4B63_202g12 | cysteine peptidase | 0,509346736 | 0,040393224 |
| C4B63_7g209 | Ribonuclease P protein subunit p25-like protein | 0,507884922 | 0,009996843 |
| C4B63_89g69 | cysteine peptidase | 0,507183863 | 0,042521567 |
| C4B63_148g34 | pumilio/PUF RNA binding protein 7 | 0,504219652 | 0,045583398 |
| C4B63_2g376 | ribosome assembly protein RRB1 | 0,500684629 | 0,008634324 |
| C4B63_10g167 | conserved hypothetical protein | 0,497832722 | 0,027694612 |
| C4B63_302g52c | histone H3 | 0,496537106 | 0,030507577 |
| C4B63_188g135c | histone H4 | 0,49495545 | 0,021126449 |
| C4B63_40g87 | WD40 repeat-containing protein | 0,489002256 | 0,038213273 |
| C4B63_58g56 | conserved hypothetical protein | 0,488718751 | 0,002185923 |
| C4B63_128g70 | ATP-dependent RNA helicase | 0,487320011 | 0,031474408 |
| C4B63_47g73 | importin beta-1 subunit | 0,485964654 | 0,005491483 |
| C4B63_3g1106 | peroxisome targeting signal 1 receptor | 0,481702746 | 0,00636503 |
| C4B63_70g93 | DNA repair and recombination helicase protein PIF2 | 0,477279931 | 0,026901022 |
| C4B63_128g44 | major vault protein | 0,476863742 | 0,02840431 |
| C4B63_439g17 | cysteine peptidase(fragment) | 0,475572304 | 0,030419999 |
| C4B63_128g61 | 50S ribosome-binding GTPase | 0,474165376 | 0,005374318 |
| C4B63_19g101 | U-rich RNA-binding protein UBP-2 | 0,474065958 | 0,033998151 |
| C4B63_51g115 | Proliferation-associated protein 2G4 | 0,474010283 | 0,047311923 |
| C4B63_135g30 | unspecified product | 0,471095105 | 0,040531493 |
| C4B63_202g32 | cysteine peptidase | 0,468928734 | 0,037826269 |
| C4B63_41g78 | conserved hypothetical protein | 0,467309088 | 0,006932193 |
| C4B63_228g44 | conserved hypothetical protein | 0,466923098 | 0,026304888 |
| C4B63_188g156c | histone H4 | 0,466736018 | 0,032487206 |
| C4B63_28g51 | ATP-dependent DEAD/H RNA helicase(fragment) | 0,460530733 | 0,029696052 |
| C4B63_16g99 | cytosolic leucyl aminopeptidase | 0,460230365 | 0,01812296 |
| C4B63_188g130c | histone H4 | 0,459414542 | 0,048842294 |
| C4B63_41g153 | cysteine peptidase | 0,458053543 | 0,016737791 |
| C4B63_6g2516c | histone H3 | 0,455181289 | 0,046952736 |
| C4B63_45g85 | conserved hypothetical protein | 0,454733608 | 0,022526483 |
| C4B63_178g43 | conserved hypothetical protein | 0,453826386 | 0,03207292 |
| C4B63_371g8 | beta galactofuranosyl glycosyltransferase | 0,452192364 | 0,033998151 |
| C4B63_56g75 | Vacuolar protein sorting-associated protein VTA1 homolog | 0,449255722 | 0,03207292 |
| C4B63_81g62 | glutamine aminotransferase | 0,441906455 | 0,014340776 |
| C4B63_11g495 | retrotransposon hot spot protein (RHS) | 0,440984832 | 0,037680795 |
| C4B63_480g11 | cysteine peptidase | 0,440896523 | 0,031620556 |
| C4B63_30g940c | Histone H2B variant V | 0,435403128 | 0,022891698 |
| C4B63_308g2 | kinetoplast DNA-associated protein(fragment) | 0,433955675 | 0,048487549 |
| C4B63_16g100 | cytosolic leucyl aminopeptidase | 0,433168356 | 0,043231054 |
| C4B63_178g46 | conserved hypothetical protein | 0,428243428 | 0,02683434 |
| C4B63_87g20 | Brefeldin A-inhibited guanine nucleotide-exchange protein 1 | 0,427291904 | 0,00852017 |
| C4B63_2g829 | conserved hypothetical protein | 0,427012686 | 0,003160363 |
| C4B63_95g67 | nucleoside 2-deoxyribosyltransferase | 0,423210276 | 0,048604443 |
| C4B63_480g19 | cysteine peptidase | 0,420835658 | 0,027610436 |
| C4B63_353g3 | unspecified product | 0,418576924 | 0,023458273 |
| C4B63_116g5 | P-type H+-ATPase | 0,418139012 | 0,006858805 |
| C4B63_5g793 | elongation factor 1-gamma (EF-1-gamma) | 0,41813434 | 0,021004555 |
| C4B63_211g35 | conserved hypothetical protein | 0,416046134 | 0,04066761 |
| C4B63_361g15 | trans-sialidase, Group V | 0,414012068 | 0,012614008 |
| C4B63_51g222 | unspecified product | 0,413551587 | 0,03207292 |
| C4B63_56g94 | conserved hypothetical protein | 0,407488954 | 0,021422112 |
| C4B63_46g107 | NADH-cytochrome b5 reductase | 0,406980431 | 0,030296037 |
| C4B63_95g55 | ubiquitin-like protein | 0,406462525 | 0,040224268 |
| C4B63_313g14 | amino acid transporter | 0,406412913 | 0,017533512 |
| C4B63_28g151 | formin-like protein | 0,406343208 | 0,032356644 |
| C4B63_41g155 | cysteine peptidase | 0,405634009 | 0,031837836 |
| C4B63_89g66 | conserved hypothetical protein | 0,405541166 | 0,026304888 |
| C4B63_38g160 | ADP-ribosylation factor | 0,40348227 | 0,047888596 |
| C4B63_480g13 | cysteine peptidase | 0,400600565 | 0,020838019 |

#### Hb exclusive DEGs (t24 vs. t0)

##### Downregulated genes

| ID | Annotation | log2FC | padj |
|---|---|---|---|
| C4B63_442g10 | Uncharacterized protein C4orf22 homolog | -0,686055549 | 0,045677358 |
| C4B63_3g1044 | Mucin-associated surface protein (MASP) | -0,601749393 | 0,024298102 |
| C4B63_83g44 | retrotransposon hot spot (RHS) protein | -0,580620225 | 0,028925311 |
| C4B63_85g422c | conserved hypothetical protein | -0,564153393 | 0,02035994 |
| C4B63_39g97 | dispersed gene family protein 1 (DGF-1) | -0,562213293 | 0,015648974 |
| C4B63_77g33 | conserved hypothetical protein | -0,542503306 | 0,010595508 |
| C4B63_16g104 | conserved hypothetical protein | -0,528548577 | 0,041076864 |
| C4B63_30g183 | conserved hypothetical protein | -0,517194945 | 0,02517909 |
| C4B63_133g32 | conserved hypothetical protein | -0,483531443 | 0,040327882 |
| C4B63_134g54 | trans-sialidase, Group V | -0,465816555 | 0,045323428 |
| C4B63_5g29 | surface protease GP63 | -0,455920559 | 0,023235857 |
| C4B63_40g21 | Charged multivesicular body protein 3 | -0,450412582 | 0,041057543 |
| C4B63_47g60 | conserved hypothetical protein | -0,442203732 | 0,004128351 |
| C4B63_55g177 | conserved hypothetical protein | -0,43775076 | 0,004058654 |
| C4B63_41g249 | conserved hypothetical protein | -0,429938038 | 0,013823208 |
| C4B63_54g61 | conserved hypothetical protein | -0,425155109 | 0,015648974 |
| C4B63_70g99 | conserved hypothetical protein | -0,421432874 | 0,042176862 |
| C4B63_12g24 | frataxin-like, mitochondrial precursor | -0,421387229 | 0,032168982 |
| C4B63_4g135 | conserved hypothetical protein | -0,408392016 | 0,022275137 |
| C4B63_53g174 | surface protease GP63 | -0,403349858 | 0,008422994 |

##### Upregulated genes

| ID | Annotation | log2FC | padj |
|---|---|---|---|
| C4B63_70g95 | 14-3-3 protein 2 | 0,94624206 | 0,000117864 |
| C4B63_85g83 | Eukaryotic initiation factor 4A-1 | 0,830748488 | 0,010598413 |
| C4B63_46g16 | conserved hypothetical protein | 0,799189369 | 0,001720686 |
| C4B63_46g17 | conserved hypothetical protein | 0,778783074 | 0,003470455 |
| C4B63_46g18 | conserved hypothetical protein | 0,762512986 | 0,001486249 |
| C4B63_31g80 | Calcium ATPase SERCA-like | 0,745740077 | 0,010256839 |
| C4B63_15g18 | NADH-dependent fumarate reductase | 0,702636822 | 0,013823208 |
| C4B63_12g330 | ATP synthase subunit beta, mitochondrial | 0,673431528 | 0,007787638 |
| C4B63_114g34 | conserved hypothetical protein | 0,668635113 | 0,002105259 |
| C4B63_148g35 | zinc-finger protein 2 | 0,642550179 | 0,016088891 |
| C4B63_61g93 | conserved hypothetical protein | 0,639931614 | 0,00149312 |
| C4B63_2g507 | zinc-finger protein | 0,639767602 | 0,006296747 |
| C4B63_20g172 | chaperonin alpha subunit | 0,635889887 | 0,022348217 |
| C4B63_84g82 | DNA mismatch repair protein MSH2 | 0,626637196 | 0,007986284 |
| C4B63_4g90 | ATP-dependent RNA helicase HEL67 | 0,623204622 | 0,033557941 |
| C4B63_19g101 | U-rich RNA-binding protein UBP-2 | 0,614614711 | 0,005218417 |
| C4B63_135g38 | retrotransposon hot spot (RHS) protein | 0,605281209 | 0,02987447 |
| C4B63_252g14 | zinc-finger protein | 0,603347938 | 0,00580587 |
| C4B63_199g2 | conserved hypothetical protein | 0,596180794 | 0,033557941 |
| C4B63_61g94 | conserved hypothetical protein | 0,5919266 | 0,036988403 |
| C4B63_13g155 | elongation factor 2 | 0,581084967 | 0,041615675 |
| C4B63_20g178 | dihydrolipoyl dehydrogenase | 0,572910383 | 0,021388281 |
| C4B63_13g154 | elongation factor 2 | 0,570973797 | 0,028623579 |
| C4B63_4g188 | Elongation factor Tu, mitochondrial | 0,567398678 | 0,000618794 |
| C4B63_21g285 | conserved hypothetical protein | 0,555022963 | 0,040517462 |
| C4B63_122g16 | oligopeptidase b | 0,549852681 | 0,020019923 |
| C4B63_13g152 | elongation factor 2 | 0,546847511 | 0,047628082 |
| C4B63_13g153 | elongation factor 2 | 0,546645343 | 0,041666099 |
| C4B63_13g156 | elongation factor 2 | 0,534903112 | 0,044108218 |
| C4B63_13g157 | elongation factor 2 | 0,511955196 | 0,039403081 |
| C4B63_25g240 | Macro domain containing protein | 0,508154657 | 0,006709369 |
| C4B63_72g79 | inositol 5-phosphatase 1 | 0,496433928 | 0,010406012 |
| C4B63_153g41 | inositol 5-phosphatase 1(fragment) | 0,483424401 | 0,0013735 |
| C4B63_55g67 | DNA-directed rna polymerase I largest subunit | 0,482678336 | 0,000571227 |
| C4B63_16g293 | inositol 1,4,5-trisphosphate receptor | 0,478986079 | 0,01683944 |
| C4B63_54g186 | retrotransposon hot spot (RHS) protein | 0,47371581 | 0,036052354 |
| C4B63_192g26 | conserved hypothetical protein | 0,47212596 | 0,046417241 |
| C4B63_32g268 | lysophospholipase | 0,467707253 | 0,030478089 |
| C4B63_138g113c | calmodulin | 0,466976037 | 0,026426014 |
| C4B63_153g40 | inositol 5-phosphatase 1 | 0,46602356 | 0,021851252 |
| C4B63_12g20 | 2-oxoglutarate-dependent dioxygenase tropC | 0,463699866 | 0,029812791 |
| C4B63_41g78 | conserved hypothetical protein | 0,460635701 | 0,012294855 |
| C4B63_206g14 | delta-1-pyrroline-5-carboxylate dehydrogenase | 0,456909272 | 0,048959124 |
| C4B63_57g65 | membrane-bound acid phosphatase 2 | 0,456064019 | 0,019537051 |
| C4B63_25g185 | ATP-dependent RNA helicase | 0,453036313 | 0,025024557 |
| C4B63_133g42 | dynein heavy chain | 0,452789934 | 0,035326218 |
| C4B63_89g45 | Mitochondrial SSU ribosomal protein 29 | 0,451945363 | 0,015648974 |
| C4B63_61g92 | conserved hypothetical protein | 0,446102262 | 0,001720686 |
| C4B63_21g42 | PAB1-binding protein | 0,44188437 | 0,041439579 |
| C4B63_42g220 | transmembrane/endomembrane-like protein | 0,440905941 | 0,046614419 |
| C4B63_12g195 | conserved hypothetical protein | 0,438398571 | 0,006111694 |
| C4B63_61g95 | conserved hypothetical protein | 0,437214353 | 0,0065861 |
| C4B63_25g237 | eukaryotic translation initiation factor 3 subunit b | 0,435441248 | 0,007152993 |
| C4B63_13g344 | conserved hypothetical protein | 0,43484463 | 0,030296061 |
| C4B63_34g110 | mitochondrial carrier protein | 0,426491385 | 0,032866209 |
| C4B63_26g260 | solanesyl-diphosphate synthase | 0,423504918 | 0,01095188 |
| C4B63_6g223 | DNA topoisomerase II | 0,410844609 | 0,041888784 |
| C4B63_31g104 | conserved hypothetical protein | 0,410047997 | 0,001786674 |
| C4B63_38g167 | spermidine synthase | 0,406170396 | 0,018998446 |
| C4B63_134g37 | orotidine-5-phosphate decarboxylase/orotate phosphoribosyltra | 0,405015126 | 0,035326218 |
| C4B63_35g109 | conserved hypothetical protein | 0,403130009 | 0,031563786 |

**No heme exclusive DEGs (t4 vs. t0)**

| Downregulated genes | | | | | Upregulated genes | | | |
|---|---|---|---|---|---|---|---|---|
| ID | Annotation | log2FC | padj | | ID | Annotation | log2FC | padj |
| C4B63_101g1 | tyrosine aminotransferase | -0,7288 | 0,00511 | | C4B63_10g1811c | cofilin/actin depolymerizing factor | 0,4488 | 0,0133 |
| C4B63_101g2 | tyrosine aminotransferase | -0,7178 | 0,00323 | | C4B63_20g177 | chaperone DNAJ protein | 0,4812 | 0,0010 |
| C4B63_101g3 | tyrosine aminotransferase | -0,7669 | 0,00101 | | C4B63_23g234 | inosine-adenosine-guanosine-nucleosidehydrolase | 0,4485 | 0,0033 |
| C4B63_101g4 | tyrosine aminotransferase | -0,7288 | 0,00255 | | C4B63_252g15 | arginyl-tRNA synthetase | 0,4827 | 0,0343 |
| C4B63_101g5 | tyrosine aminotransferase | -0,7178 | 0,00072 | | C4B63_253g15 | haloacid dehalogenase-like hydrolase | 0,4921 | 0,0329 |
| C4B63_13g199 | GPI transamidase component Tta2 | -0,7669 | 0,02806 | | C4B63_45g221 | ATP-dependent DEAD/H RNA helicase | 0,4464 | 0,0272 |
| C4B63_21g103 | citrate transporter | -0,4562 | 0,04182 | | C4B63_51g80 | ferric reductase transmembrane protein | 0,4453 | 0,0077 |
| C4B63_25g331 | trans-sialidase, Group I | -1,0142 | 0,02476 | | C4B63_51g81 | ferric reductase transmembrane protein | 0,5492 | 0,0007 |
| C4B63_26g354 | tyrosine aminotransferase | -0,7442 | 0,00196 | | C4B63_51g82 | ferric reductase transmembrane protein | 0,4628 | 0,0048 |
| C4B63_33g163 | unspecified product | -0,4737 | 0,01724 | | C4B63_51g83 | ferric reductase transmembrane protein | 0,4835 | 0,0121 |
| C4B63_33g164 | unspecified product | -0,5137 | 0,04286 | | C4B63_51g85 | ferric reductase transmembrane protein | 0,5001 | 0,0026 |
| C4B63_42g164 | tyrosine aminotransferase | -0,7621 | 0,00150 | | C4B63_61g73 | Acyl-CoA-binding domain-containing protein 5 | 0,5059 | 0,0146 |
| C4B63_43g196 | adenylate kinase 1 | -0,5080 | 0,04341 | | C4B63_76g32 | guide RNA associated protein, GAP2 | 0,4001 | 0,0385 |
| C4B63_44g227 | amino acid permease | -0,5240 | 0,00100 | | | | | |
| C4B63_4g336 | CMP-sialic acid transporter | -0,4944 | 0,04398 | | | | | |
| C4B63_54g138 | tyrosine aminotransferase | -0,7707 | 0,00046 | | | | | |
| C4B63_81g64 | tyrosine aminotransferase | -0,7436 | 0,00191 | | | | | |
| C4B63_81g65 | tyrosine aminotransferase | -0,6865 | 0,00247 | | | | | |
| C4B63_81g66 | tyrosine aminotransferase | -0,7133 | 0,00250 | | | | | |
| C4B63_81g67 | tyrosine aminotransferase | -0,7029 | 0,00327 | | | | | |
| C4B63_81g68 | tyrosine aminotransferase | -0,7169 | 0,00144 | | | | | |
| C4B63_81g69 | tyrosine aminotransferase | -0,7204 | 0,00118 | | | | | |
| C4B63_81g70 | tyrosine aminotransferase | -0,7579 | 0,00450 | | | | | |

**No heme exclusive DEGs (t24 vs. t0)**

| Downregulated genes | | | | | Upregulated genes | | | |
|---|---|---|---|---|---|---|---|---|
| ID | Annotation | log2FC | padj | | ID | Annotation | log2FC | padj |
| C4B63_122g15 | conserved hypothetical protein | -0,6850 | 0,0487 | | C4B63_18g39 | CRAL/TRIO, N-terminal domain/CRAL/TRIO domair | 0,6361 | 0,0011 |
| C4B63_127g52 | serine/threonine-protein kinase A(fragment) | -0,4436 | 0,0034 | | C4B63_23g43 | conserved hypothetical protein | 0,5146 | 0,0261 |
| C4B63_135g21 | ascorbate peroxidase | -1,0347 | 0,0174 | | C4B63_28g80 | DnaJ chaperone protein | 0,4633 | 0,0421 |
| C4B63_13g200 | Enriched in surface-labeled proteome protein 18 | -0,7645 | 0,0008 | | C4B63_302g66c | histone H3 | 0,7388 | 0,0394 |
| C4B63_181g25 | conserved hypothetical protein | -0,9024 | 0,0293 | | C4B63_30g161 | glutamine synthetase | 0,4201 | 0,0474 |
| C4B63_22g121 | unspecified product | -0,6816 | 0,0215 | | C4B63_51g81 | ferric reductase transmembrane protein | 0,4310 | 0,0171 |
| C4B63_248g12 | Present in the outer mitochondrial membrane proteome | -0,7129 | 0,0470 | | C4B63_51g82 | ferric reductase transmembrane protein | 0,4176 | 0,0171 |
| C4B63_267g18 | conserved hypothetical protein | -0,5594 | 0,0430 | | C4B63_6g2538c | histone H3 | 0,7169 | 0,0407 |
| C4B63_26g51 | Mucin-associated surface protein (MASP) | -0,6166 | 0,0407 | | C4B63_6g2542c | histone H3 | 0,6173 | 0,0355 |
| C4B63_281g9 | Ring finger domain containing protein | -0,6889 | 0,0411 | | C4B63_6g2554c | histone H3 | 0,6934 | 0,0322 |
| C4B63_28g96 | sugar transporter(fragment) | -0,7890 | 0,0407 | | C4B63_6g2562c | histone H3 | 0,8380 | 0,0101 |
| C4B63_33g147 | NADH-ubiquinone oxidoreductase, mitochondrial | -0,7577 | 0,0152 | | | | | |
| C4B63_38g176 | conserved hypothetical protein | -0,6967 | 0,0337 | | | | | |
| C4B63_43g134 | cytochrome c oxidase subunit VI | -0,5408 | 0,0060 | | | | | |
| C4B63_43g98 | unspecified product | -0,6609 | 0,0408 | | | | | |
| C4B63_45g115 | Protein kinase domain | -0,4218 | 0,0484 | | | | | |
| C4B63_47g23 | ascorbate peroxidase | -0,9819 | 0,0364 | | | | | |
| C4B63_4g189 | alcohol dehydrogenase | -0,9326 | 0,0020 | | | | | |
| C4B63_55g92 | conserved hypothetical protein | -0,4839 | 0,0294 | | | | | |
| C4B63_6g259 | nucleotidyl transferase | -0,4357 | 0,0145 | | | | | |
| C4B63_7g361 | phosphomannomutase-like protein | -0,6676 | 0,0144 | | | | | |
| C4B63_82g448c | conserved hypothetical protein | -0,4751 | 0,0355 | | | | | |
| C4B63_89g64 | hydroxyacylglutathione hydrolase | -0,5887 | 0,0398 | | | | | |
| C4B63_9g408 | conserved hypothetical protein | -0,5091 | 0,0246 | | | | | |

| Gene Ontology analysis of Hemin exclusive DEGs (downregulated, t4 vs. t0) | | | | | | | | | | | |
|---|---|---|---|---|---|---|---|---|---|---|---|
| | **CELLULAR COMPONENT** | | | | | | | | | | |
| ID | Name | Bgd count | Result count | Result gene | Pct of bgd | Fold enrichm | Odds ratio | P-value | | Benjamini | Bonferroni |
| GO:0042995 | cell projection | 163 | 31 | C4B63_231g | 19.0 | 13.25 | 21.22 | 6,73E-27 | | 9,26E-26 | 3,70E-25 |
| GO:0043227 | membrane-bounded organelle | 995 | 34 | C4B63_231g | 3.4 | 2.38 | 2.97 | 8,15E-07 | | 8,97E-06 | 4,48E-05 |
| | **MOLECULAR FUNCTION** | | | | | | | | | | |
| ID | Name | Bgd count | Result count | Result gene | Pct of bgd | Fold enrichm | Odds ratio | P-value | | Benjamini | Bonferroni |
| GO:0005516 | calmodulin binding | 24 | 11 | C4B63_55g3 | 45.8 | 31.92 | 63.7 | 7,24E-15 | | 1,01E-12 | 1,01E-12 |
| GO:0046872 | metal ion binding | 792 | 31 | C4B63_13g2 | 3.9 | 2.73 | 3.4 | 1,49E-07 | | 6,69E-06 | 2,07E-05 |

| Gene Ontology analysis of Hemin exclusive DEGs (upregulated, t4 vs. t0) | | | | | | | | | | |
|---|---|---|---|---|---|---|---|---|---|---|
| | **MOLECULAR FUNCTION** | | | | | | | | | |
| ID | Name | Bgd count | Result count | Result gene | Pct of bgd | Fold enrichm | Odds ratio | P-value | Benjamini | Bonferroni |
| GO:0003723 | RNA binding | 425 | 18 | C4B63_109g | 4.2 | 3.05 | 3.52 | 2,07E-05 | 0.00394570728267 | 0.00394570728267 |
| GO:0003676 | nucleic acid binding | 987 | 29 | C4B63_109g | 2.9 | 2.12 | 2.52 | 5,98E-05 | 0.00571378513375 | 0.0114275702675 |

| Gene Ontology analysis of Hemin exclusive DEGs (downregulated, t24 vs. t0) | | | | | | | | | | |
|---|---|---|---|---|---|---|---|---|---|---|
| | **CELLULAR COMPONENT** | | | | | | | | | |
| ID | Name | Bgd count | Result count | Result gene | Pct of bgd | Fold enrichm | Odds ratio | P-value | Benjamini | Bonferroni |
| GO:0016020 | membrane | 2776 | 85 | C4B63_109g | 3.1 | 1.51 | 2.02 | 3,94E-06 | 6,70E-05 | 0.000200902193868 |
| | **MOLECULAR FUNCTION** | | | | | | | | | |
| ID | Name | Bgd count | Result count | Result gene | Pct of bgd | Fold enrichm | Odds ratio | P-value | Benjamini | Bonferroni |
| GO:0030983 | mismatched DNA binding | 16 | 6 | C4B63_102g | 37.5 | 18.46 | 29.94 | 4,36E-07 | 5,58E-05 | 5,58E-05 |
| GO:0003690 | double-stranded DNA binding | 19 | 6 | C4B63_102g | 31.6 | 15.55 | 23.02 | 1,40E-06 | 8,98E-05 | 0.000179602487712 |
| GO:0016997 | alpha-sialidase activity | 711 | 31 | C4B63_109g | 4.4 | 2.15 | 2.46 | 3,49E-05 | 0.00111708551683 | 0.00446834206732 |

| Gene Ontology analysis of Hemin exclusive DEGs (upregulated, t24 vs. t0) | | | | | | | | | | |
|---|---|---|---|---|---|---|---|---|---|---|
| | **BIOLOGICAL PROCESS** | | | | | | | | | |
| ID | Name | Bgd count | Result count | Result gene | Pct of bgd | Fold enrichm | Odds ratio | P-value | Benjamini | Bonferroni |
| GO:0019752 | carboxylic acid metabolic process | 307 | 19 | C4B63_12g2 | 6.2 | 3.27 | 3.74 | 4,90E-06 | 0.000399771837744 | 0.00223393647037 |
| GO:0006096 | glycolytic process | 22 | 5 | C4B63_13g2 | 22.7 | 12.02 | 15.71 | 4,61E-05 | 0.00140249904726 | 0.0210374857089 |
| GO:1901360 | organic cyclic compound metabolic process | 722 | 28 | C4B63_10g4 | 3.9 | 2.05 | 2.32 | 0.000184754098715 | 0.00350964222004 | 0.0842478690139 |
| GO:0008610 | lipid biosynthetic process | 85 | 8 | C4B63_123g | 9.4 | 4.98 | 5.62 | 0.000192414595397 | 0.00350964222004 | 0.0877410555011 |
| GO:0043039 | tRNA aminoacylation | 65 | 7 | C4B63_297g | 10.8 | 5.7 | 6.5 | 0.000210465679428 | 0.00355453147479 | 0.0959723498193 |
| | **CELLULAR COMPONENT** | | | | | | | | | |
| ID | Name | Bgd count | Result count | Result gene | Pct of bgd | Fold enrichm | Odds ratio | P-value | Benjamini | Bonferroni |
| GO:0005634 | nucleus | 424 | 21 | C4B63_112g | 5.0 | 2.62 | 2.96 | 4,59E-05 | 0.00140020490363 | 0.00560081961453 |
| GO:0044815 | DNA packaging complex | 124 | 10 | C4B63_128g | 8.1 | 4.26 | 4.78 | 0.000113442888295 | 0.00221903678164 | 0.013840032372 |
| GO:0030137 | COPI-coated vesicle | 7 | 3 | C4B63_22g1 | 42.9 | 22.66 | 39.63 | 0.000219718471203 | 0.00297840594297 | 0.0268056534868 |
| GO:0020015 | glycosome | 2 | 2 | C4B63_18g2 | 100.0 | 52.88 | inf | 0.000355413609349 | 0.00309717573861 | 0.0433604603406 |
| GO:0005798 | Golgi-associated vesicle | 9 | 3 | C4B63_22g1 | 33.3 | 17.63 | 26.41 | 0.000512796726598 | 0.00368007062618 | 0.062561200645 |
| GO:0005737 | cytoplasm | 813 | 29 | C4B63_112g | 3.6 | 1.89 | 2.12 | 0.000571835754917 | 0.00373538103579 | 0.0697639620999 |
| GO:0032991 | protein-containing complex | 630 | 24 | C4B63_10g5 | 3.8 | 2.01 | 2.24 | 0.000726724781263 | 0.0044330211657 | 0.0886604233141 |
| | **MOLECULAR FUNCTION** | | | | | | | | | |
| ID | Name | Bgd count | Result count | Result gene | Pct of bgd | Fold enrichm | Odds ratio | P-value | Benjamini | Bonferroni |
| GO:1901363 | heterocyclic compound binding | 2258 | 66 | C4B63_10g4 | 2.9 | 1.55 | 1.95 | 4,07E-05 | 0.00476664764343 | 0.00953329528686 |
| GO:0016705 | oxidoreductase activity, acting on paired donors, w | 14 | 4 | C4B63_123g | 28.6 | 15.11 | 21.25 | 0.000106378177048 | 0.00711262363991 | 0.0248924934292 |
| GO:0005524 | ATP binding | 1110 | 37 | C4B63_113g | 3.3 | 1.76 | 2.02 | 0.000336855217016 | 0.00711262363991 | 0.0788241207818 |
| GO:0004332 | fructose-bisphosphate aldolase activity | 2 | 2 | C4B63_13g2 | 100.0 | 52.88 | inf | 0.000355413609349 | 0.00711262363991 | 0.0831667845877 |
| GO:0050242 | pyruvate, phosphate dikinase activity | 2 | 2 | C4B63_12g2 | 100.0 | 52.88 | inf | 0.000355413609349 | 0.00711262363991 | 0.0831667845877 |

| Gene Ontology analysis of Hb exclusive DEGs (downregulated, t4 vs. t0) | | | | | | | | | | |
|---|---|---|---|---|---|---|---|---|---|---|
| | **CELLULAR COMPONENT** | | | | | | | | | |
| ID | Name | Bgd count | Result count | Result gene | Pct of bgd | Fold enrichm | Odds ratio | P-value | Benjamini | Bonferroni |
| GO:0005739 | mitochondrion | 131 | 9 | C4B63_12g2 | 6.9 | 6.07 | 7.0 | 1,59E-05 | 0.00047777 | 0.000955550395816 |
| | **MOLECULAR FUNCTION** | | | | | | | | | |
| ID | Name | Bgd count | Result count | Result gene | Pct of bgd | Fold enrichm | Odds ratio | P-value | Benjamini | Bonferroni |
| GO:0016491 | oxidoreductase activity | 402 | 19 | C4B63_101g | 4.7 | 4.17 | 5.14 | 8,89E-08 | 1,30E-05 | 1,30E-05 |
| GO:0004435 | phosphatidylinositol phospholipase C activity | 2 | 2 | C4B63_120g | 100.0 | 88.32 | inf | 0.00012689 | 0.00463157 | 0.0185263125304 |

| Gene Ontology analysis of Hb exclusive DEGs (upregulated, t4 vs. t0) | | | | | | | | | | |
|---|---|---|---|---|---|---|---|---|---|---|
| | **BIOLOGICAL PROCESS** | | | | | | | | | |
| ID | Name | Bgd count | Result count | Result gene | Pct of bgd | Fold enrichm | Odds ratio | P-value | Benjamini | Bonferroni |
| GO:0006508 | proteolysis | 318 | 31 | C4B63_16g1 | 9.7 | 9.18 | 14.74 | 2,86E-22 | 4,57E-20 | 4,57E-20 |
| GO:0008380 | RNA splicing | 63 | 5 | C4B63_178g | 7.9 | 7.47 | 8.44 | 0.00052373 | 0.00761794 | 0.0837973837498 |
| | **CELLULAR COMPONENT** | | | | | | | | | |
| ID | Name | Bgd count | Result count | Result gene | Pct of bgd | Fold enrichm | Odds ratio | P-value | Benjamini | Bonferroni |
| GO:0044815 | DNA packaging complex | 124 | 15 | C4B63_188g | 12.1 | 11.39 | 15.15 | 2,36E-12 | 2,52E-11 | 8,96E-11 |
| GO:0005634 | nucleus | 424 | 15 | C4B63_10g1 | 3.5 | 3.33 | 3.89 | 3,51E-05 | 0.00019062 | 0.00133437379986 |
| | **MOLECULAR FUNCTION** | | | | | | | | | |
| ID | Name | Bgd count | Result count | Result gene | Pct of bgd | Fold enrichm | Odds ratio | P-value | Benjamini | Bonferroni |
| GO:0003676 | nucleic acid binding | 987 | 24 | C4B63_10g1 | 2.4 | 2.29 | 2.79 | 6,70E-05 | 0.00064231 | 0.0077077624258 |

| Gene Ontology analysis of Hb exclusive DEGs (downregulated, t24 vs. t0) | | | | | | | | | | |
|---|---|---|---|---|---|---|---|---|---|---|
| | **MOLECULAR FUNCTION** | | | | | | | | | |
| ID | Name | Bgd count | Result count | Result gene | Pct of bgd | Fold enrichm | Odds ratio | P-value | Benjamini | Bonferroni |
| GO:0008199 | ferric iron binding | 1 | 1 | C4B63_12g2 | 100.0 | 778.82 | inf | 0.00128399 | 0.00770398 | 0.0308159215598 |

| Gene Ontology analysis of Hb exclusive DEGs (upregulated, t24 vs. t0) | | | | | | | | | | |
|---|---|---|---|---|---|---|---|---|---|---|
| | **BIOLOGICAL PROCESS** | | | | | | | | | |
| ID | Name | Bgd count | Result count | Result gene | Pct of bgd | Fold enrichm | Odds ratio | P-value | Benjamini | Bonferroni |
| GO:0006816 | calcium ion transport | 5 | 2 | C4B63_16g2 | 40.0 | 76.15 | 132.08 | 0.00026711 | 0.02555691 | 0.0512853154451 |
| GO:0046856 | phosphatidylinositol dephosphorylation | 6 | 2 | C4B63_153g | 33.3 | 63.46 | 99.05 | 0.00039932 | 0.02555691 | 0.07667075578 |
| | **MOLECULAR FUNCTION** | | | | | | | | | |
| ID | Name | Bgd count | Result count | Result gene | Pct of bgd | Fold enrichm | Odds ratio | P-value | Benjamini | Bonferroni |
| GO:0045182 | translation regulator activity | 128 | 9 | C4B63_13g1 | 7.0 | 13.39 | 17.65 | 1,57E-08 | 6,19E-07 | 1,86E-06 |
| GO:0003676 | nucleic acid binding | 987 | 15 | C4B63_13g1 | 1.5 | 2.89 | 3.88 | 9,13E-05 | 0.00098822 | 0.0107720144924 |
| GO:0043167 | ion binding | 2086 | 23 | C4B63_12g2 | 1.1 | 2.1 | 3.27 | 9,21E-05 | 0.00098822 | 0.0108705236749 |

| Gene Ontology analysis of No heme exclusive DEGs (downregulated, t4 vs. t0) | | | | | | | | | | |
|---|---|---|---|---|---|---|---|---|---|---|
| | **BIOLOGICAL PROCESS** | | | | | | | | | |
| ID | Name | Bgd count | Result count | Result gene | Pct of bgd | Fold enrichm | Odds ratio | P-value | Benjamini | Bonferroni |
| GO:0009072 | aromatic amino acid family metabolic process | 13 | 11 | C4B63_101g | 84.6 | 362.45 | 5221.94 | 2,88E-29 | 1,80E-28 | 1,44E-27 |
| | **CELLULAR COMPONENT** | | | | | | | | | |
| ID | Name | Bgd count | Result count | Result gene | Pct of bgd | Fold enrichm | Odds ratio | P-value | Benjamini | Bonferroni |
| GO:0005739 | mitochondrion | 131 | 9 | C4B63_101g | 6.9 | 29.43 | 56.5 | 5,05E-12 | 4,55E-11 | 9,09E-11 |
| GO:0005737 | cytoplasm | 813 | 10 | C4B63_101g | 1.2 | 5.27 | 9.64 | 4,28E-06 | 2,57E-05 | 7,70E-05 |
| | **MOLECULAR FUNCTION** | | | | | | | | | |
| ID | Name | Bgd count | Result count | Result gene | Pct of bgd | Fold enrichm | Odds ratio | P-value | Benjamini | Bonferroni |
| GO:0008483 | transaminase activity | 25 | 15 | C4B63_101g | 60.0 | 257.01 | 2561.1 | 6,79E-37 | 1,46E-35 | 2,92E-35 |
| GO:0030170 | pyridoxal phosphate binding | 36 | 15 | C4B63_101g | 41.7 | 178.48 | 1218.0 | 1,15E-33 | 1,24E-32 | 4,94E-32 |

| Gene Ontology analysis of No heme exclusive DEGs (upregulated, t4 vs. t0) | | | | | | | | | | |
|---|---|---|---|---|---|---|---|---|---|---|
| | **BIOLOGICAL PROCESS** | | | | | | | | | |
| ID | Name | Bgd count | Result count | Result gene | Pct of bgd | Fold enrichm | Odds ratio | P-value | Benjamini | Bonferroni |
| GO:0030042 | actin filament depolymerization | 1 | 1 | C4B63_10g1 | 100.0 | 713.92 | inf | 0.00140072370725 | 0.0280144741449 | 0.0840434224348 |
| | **MOLECULAR FUNCTION** | | | | | | | | | |
| ID | Name | Bgd count | Result count | Result gene | Pct of bgd | Fold enrichm | Odds ratio | P-value | Benjamini | Bonferroni |
| GO:0016491 | oxidoreductase activity | 402 | 5 | C4B63_51g8 | 1.2 | 8.88 | 14.68 | 0.000133649842961 | 0.00574694324732 | 0.00574694324732 |

| Gene Ontology analysis of No heme exclusive DEGs (downregulated, t24 vs. t0) | | | | | | | | | | |
|---|---|---|---|---|---|---|---|---|---|---|
| | **BIOLOGICAL PROCESS** | | | | | | | | | |
| ID | Name | Bgd count | Result count | Result gene | Pct of bgd | Fold enrichm | Odds ratio | P-value | Benjamini | Bonferroni |
| GO:0006979 | response to oxidative stress | 11 | 2 | C4B63_135g | 18.2 | 119.82 | 172.63 | 0.000116019957952 | 0.00359661869651 | 0.00359661869651 |
| GO:0008643 | carbohydrate transport | 2 | 1 | C4B63_28g9 | 50.0 | 329.5 | 712.75 | 0.00303277558837 | 0.0313386810798 | 0.0940160432395 |
| | **CELLULAR COMPONENT** | | | | | | | | | |
| ID | Name | Bgd count | Result count | Result gene | Pct of bgd | Fold enrichm | Odds ratio | P-value | Benjamini | Bonferroni |
| GO:0070469 | respirasome | 15 | 2 | C4B63_33g1 | 13.3 | 87.87 | 119.45 | 0.000220735212019 | 0.0057391155125 | 0.0057391155125 |
| | **MOLECULAR FUNCTION** | | | | | | | | | |
| ID | Name | Bgd count | Result count | Result gene | Pct of bgd | Fold enrichm | Odds ratio | P-value | Benjamini | Bonferroni |
| GO:0004601 | peroxidase activity | 17 | 2 | C4B63_135g | 11.8 | 77.53 | 103.5 | 0.000285415414571 | 0.0044914450695 | 0.015983263216 |
| GO:0043169 | cation binding | 801 | 6 | C4B63_135g | 0.7 | 4.94 | 8.37 | 0.000632667764084 | 0.00541458438806 | 0.0354293947887 |

| Gene Ontology analysis of No heme exclusive DEGs (upregulated, t24 vs. t0) | | | | | | | | | | |
|---|---|---|---|---|---|---|---|---|---|---|
| | **BIOLOGICAL PROCESS** | | | | | | | | | |
| ID | Name | Bgd count | Result count | Result gene | Pct of bgd | Fold enrichm | Odds ratio | P-value | Benjamini | Bonferroni |
| GO:0006542 | glutamine biosynthetic process | 2 | 1 | C4B63_30g1 | 50.0 | 475.94 | 1069.62 | 0.00210010443286 | 0.05880292412 | 0.05880292412 |
| | **CELLULAR COMPONENT** | | | | | | | | | |
| ID | Name | Bgd count | Result count | Result gene | Pct of bgd | Fold enrichm | Odds ratio | P-value | Benjamini | Bonferroni |
| GO:0044815 | DNA packaging complex | 124 | 5 | C4B63_302g | 4.0 | 38.38 | 88.64 | 7,05E-08 | 2,94E-07 | 1,13E-06 |
| GO:0000785 | chromatin | 125 | 5 | C4B63_302g | 4.0 | 38.08 | 87.9 | 7,34E-08 | 2,94E-07 | 1,17E-06 |
| | **MOLECULAR FUNCTION** | | | | | | | | | |
| ID | Name | Bgd count | Result count | Result gene | Pct of bgd | Fold enrichm | Odds ratio | P-value | Benjamini | Bonferroni |
| GO:0003677 | DNA binding | 291 | 5 | C4B63_302g | 1.7 | 16.36 | 36.15 | 4,92E-06 | 2,63E-05 | 7,88E-05 |

### Table S5

| Hemin DEGs (t4 vs. t0)∩(t24 vs. t0) | | | |
|---|---|---|---|
| Downregulated | | Upregulated | |
| ID | Annotation | ID | Annotation |
| C4B63_10g453 | ABC transporter | C4B63_10g436 | SPFH domain / Band 7 family |
| C4B63_10g454 | ABC transporter(fragment) | C4B63_111g29 | conserved hypothetical protein |
| C4B63_10g455 | ABC transporter | C4B63_114g17 | conserved hypothetical protein |
| C4B63_10g472 | conserved hypothetical protein | C4B63_114g25 | conserved hypothetical protein |
| C4B63_113g47 | Generative cell specific 1 protein | C4B63_116g15 | membrane transporter protein |
| C4B63_113g62 | kinetoplast DNA-associated protein | C4B63_11g63 | D-alanine--D-alanine ligase |
| C4B63_127g46 | conserved hypothetical protein | C4B63_133g1 | NAD-specific glutamate dehydrogenase |
| C4B63_12g151 | conserved hypothetical protein | C4B63_137g10 | Coatomer subunit gamma |
| C4B63_134g49 | trans-sialidase, Group V | C4B63_142g6 | GMP synthase |
| C4B63_134g59 | trans-sialidase, Group V | C4B63_143g34 | condensin complex subunit 2 |
| C4B63_13g146 | conserved hypothetical protein | C4B63_14g39 | conserved hypothetical protein |
| C4B63_13g208 | conserved hypothetical protein,zinc finger protein family member | C4B63_151g36 | notchless homolog |
| C4B63_13g223 | zinc-finger protein family member | C4B63_15g12 | conserved hypothetical protein |
| C4B63_13g256 | conserved hypothetical protein | C4B63_15g16 | oligosaccharyl transferase subunit |
| C4B63_13g317 | vesicle-associated membrane protein | C4B63_171g4 | phosphatidic acid phosphatase protein |
| C4B63_14g101 | conserved hypothetical protein | C4B63_183g14 | conserved hypothetical protein |
| C4B63_14g158 | serine/threonine kinase | C4B63_184g8 | GTP-binding protein |
| C4B63_14g20 | hexose transporter | C4B63_18g152 | conserved hypothetical protein |
| C4B63_14g43 | centrin-3 | C4B63_19g205 | glycosomal transporter (GAT3) |
| C4B63_156g3 | unspecified product | C4B63_19g82 | RNA polymerase I second largest subunit |
| C4B63_156g4 | unspecified product | C4B63_1g57 | AAA ATPase |
| C4B63_157g36 | trans-sialidase, Group II | C4B63_1g70 | mitochondrial RNA-binding protein 2 |
| C4B63_16g183 | surface protease GP63 | C4B63_200g32 | 3,2-trans-enoyl-CoA isomerase, mitochondrial precursor |
| C4B63_16g93 | Uncharacterized protein in mobD 3'region | C4B63_20g747c | cyclin-dependent kinases regulatory subunit |
| C4B63_17g196 | calpain-like cysteine peptidase | C4B63_210g33 | p-nitrophenylphosphatase |
| C4B63_17g61 | conserved hypothetical protein | C4B63_220g11 | Beta propeller protein 1 |
| C4B63_180g10 | retrotransposon hot spot (RHS) protein | C4B63_227g7 | transporter |
| C4B63_18g308 | sphingosine 1-phosphate lyase | C4B63_227g9 | mevalonate-diphosphate decarboxylase |
| C4B63_18g322 | conserved hypothetical protein | C4B63_23g116 | Importin beta-like SAD2 |
| C4B63_18g326 | zinc-finger protein ZFP1 | C4B63_252g23 | Serine-threonine kinase receptor-associated protein |
| C4B63_18g330 | zinc-finger protein ZFP1 | C4B63_25g113 | GMP reductase |
| C4B63_18g39 | CRAL/TRIO, N-terminal domain/CRAL/TRIO domain containing protein | C4B63_25g117 | conserved hypothetical protein |
| C4B63_196g33 | unspecified product | C4B63_25g234 | RNA cytidine acetyltransferase |
| C4B63_19g141 | electron transfer flavoprotein | C4B63_263g7 | ruvB-like DNA helicase |
| C4B63_19g257 | conserved hypothetical protein | C4B63_282g15 | cation transporter |
| C4B63_1g473 | trans-sialidase | C4B63_286g18 | eukaryotic translation initiation factor 3 subunit c |
| C4B63_1g480 | trans-sialidase, Group V | C4B63_28g180 | WASH complex subunit 4 |
| C4B63_1g702 | trans-sialidase, Group V | C4B63_28g56 | TFIIF-stimulated CTD phosphatase |
| C4B63_20g203 | Zinc finger protein, C3H1 type-like | C4B63_2g209 | Present in the outer mitochondrial membrane proteome 6 |
| C4B63_21g27 | arginine permease(fragment) | C4B63_2g219 | conserved hypothetical protein |
| C4B63_21g6 | arginine permease | C4B63_2g289 | Hypoxia up-regulated protein 1 (Fragment) |
| C4B63_244g8 | trans-sialidase, Group I | C4B63_2g331 | polyadenylate-binding protein 1 |
| C4B63_24g194 | microtubule-associated protein | C4B63_2g43 | short-chain dehydrogenase |
| C4B63_24g236 | microtubule-associated protein Gb4 | C4B63_2g486 | Serine-threonine kinase receptor-associated protein |
| C4B63_24g274 | conserved hypothetical protein | C4B63_322g14 | amino acid transporter |
| C4B63_24g275 | conserved hypothetical protein | C4B63_35g130 | orotidine-5-phosphate decarboxylase/orotate phosphoribosyltransferase |
| C4B63_24g313 | protein kinase | C4B63_35g168 | 3-Beta-hydroxysteroid-delta(8), delta(7)-isomerase |
| C4B63_251g10 | lipase | C4B63_38g147 | spermidine synthase |
| C4B63_258g25 | lipase | C4B63_396g14 | Present in the outer mitochondrial membrane proteome 6 |
| C4B63_25g207 | hydrolase-like protein | C4B63_40g159 | translation initiation factor 2 subunit |
| C4B63_25g220 | 5'-3' exoribonuclease B | C4B63_40g25 | Biotin--acetyl-CoA-carboxylase ligase |
| C4B63_25g236 | conserved hypothetical protein | C4B63_41g139 | conserved hypothetical protein |
| C4B63_26g310 | ABC transporter | C4B63_45g97 | eukaryotic translation initiation factor 3 subunit h |
| C4B63_26g34 | mucin TcMUCII | C4B63_4g118 | conserved hypothetical protein |
| C4B63_277g19 | metallo-peptidase, clan MH, family M18 | C4B63_4g144 | protein transport protein SEC13 |
| C4B63_278g3 | conserved hypothetical protein | C4B63_4g219 | nucleolar RNA-associated proteins (Nrap) |
| C4B63_27g1020c | RNA-binding protein | C4B63_4g381 | transporter |
| C4B63_27g184 | integral membrane transport protein | C4B63_4g383 | mevalonate-diphosphate decarboxylase |
| C4B63_294g3 | metallo-peptidase, clan MH, family M18 | C4B63_4g72 | fibrillarin |
| C4B63_2g156 | protein kinase A catalytic subunit 3 | C4B63_51g842c | Uncharacterized protein At2g34160 |
| C4B63_2g309 | NIMA-related kinase | C4B63_52g161 | tyrosyl-tRNA synthetase |
| C4B63_30g222 | conserved hypothetical protein | C4B63_52g28 | catalytic subunit of the vacuolar transporter chaperone 4(fragment) |
| C4B63_322g4 | glutamamyl carboxypeptidase(fragment) | C4B63_52g54 | conserved hypothetical protein |
| C4B63_323g3 | unspecified product | C4B63_56g31 | conserved hypothetical protein |
| C4B63_354g1 | electron transfer flavoprotein | C4B63_57g98 | succinyl-CoA ligase |
| C4B63_37g189 | trans-sialidase, Group VI | C4B63_61g149 | eukaryotic translation initiation factor 3 subunit a |
| C4B63_3g1092 | alanine racemase | C4B63_63g145 | retrotransposon hot spot (RHS) protein |
| C4B63_419g2 | trans-sialidase | C4B63_63g7 | eukaryotic translation initiation factor 3 subunit c |
| C4B63_43g74 | retrotransposon hot spot (RHS) protein | C4B63_6g161 | Eukaryotic initiation factor 4A-1 |
| C4B63_44g183 | conserved hypothetical protein | C4B63_6g19 | NAD(P)-dependent steroid dehydrogenase protein |
| C4B63_44g231 | amino acid transporter | C4B63_6g236 | cation transporter |
| C4B63_44g235 | Kinesin-13 4 | C4B63_6g397 | cation transporter |
| C4B63_44g243 | conserved hypothetical protein | C4B63_6g562 | NAD(P)-dependent steroid dehydrogenase protein |
| C4B63_459g4 | conserved hypothetical protein | C4B63_70g104 | Ribosome biogenesis protein BOP1 homolog |
| C4B63_45g139 | serine/threonine protein kinase | C4B63_73g16 | conserved hypothetical protein |
| C4B63_462g9 | glutamamyl carboxypeptidase(fragment) | C4B63_79g20 | conserved hypothetical protein |
| C4B63_47g122 | trans-sialidase | C4B63_79g21 | ABC transporter |
| C4B63_47g125 | trans-sialidase | C4B63_79g37 | methionyl-tRNA synthetase |
| C4B63_48g693c | conserved hypothetical protein | C4B63_7g266 | conserved hypothetical protein |
| C4B63_49g168 | conserved hypothetical protein | C4B63_7g316 | ruvB-like DNA helicase |
| C4B63_4g132 | NLI interacting factor-like phosphatase | C4B63_7g318 | peroxisome biogenesis factor 1 |
| C4B63_4g180 | serine/threonine protein kinase | C4B63_7g340 | beta-fructofuranosidase-like protein |
| C4B63_4g242 | protein kinase A catalytic subunit 3 | C4B63_7g347 | V-type ATPase, A subunit |
| C4B63_4g283 | conserved hypothetical protein | C4B63_80g17 | ATP binding protein-like protein |
| C4B63_4g511 | zinc-finger protein | C4B63_81g17 | DNA mismatch repair protein MSH2(fragment) |
| C4B63_4g545 | RNA-binding protein 4 | C4B63_81g18 | chaperonin HSP60, mitochondrial precursor(fragment) |
| C4B63_52g153 | conserved hypothetical protein | C4B63_84g49 | RNA-binding protein |
| C4B63_53g165 | conserved hypothetical protein | C4B63_84g67 | conserved hypothetical protein |
| C4B63_550g4 | glutamamyl carboxypeptidase(fragment) | C4B63_86g57 | conserved hypothetical protein |
| C4B63_550g5 | glutamamyl carboxypeptidase(fragment) | C4B63_8g510 | conserved hypothetical protein |
| C4B63_58g126 | ubiquitin-conjugating enzyme variant Kua homologue | C4B63_8g62 | Nucleolar protein 10 |
| C4B63_59g207 | mucin TcMUCII | C4B63_97g19 | pyruvate dehydrogenase E1 component alpha subunit |
| C4B63_5g393 | trans-sialidase, Group V | C4B63_97g34 | P-type H+-ATPase |
| C4B63_65g5 | unspecified product | C4B63_97g36 | P-type H+-ATPase |
| C4B63_66g170 | Mucin-associated surface protein (MASP) | C4B63_97g39 | P-type H+-ATPase |
| C4B63_66g61 | surface protease GP63 | C4B63_98g4 | glucose-6-phosphate 1-dehydrogenase |
| C4B63_66g68 | trans-sialidase, Group V | C4B63_9g201 | GTP-binding protein |
| C4B63_69g318c | conserved hypothetical protein | | |
| C4B63_6g284 | cytochrome c oxidase assembly protein | | |
| C4B63_6g587 | N-glycosidase YbiA | | |
| C4B63_70g82 | protein kinase | | |
| C4B63_70g84 | serine/threonine protein kinase | | |
| C4B63_76g17 | conserved hypothetical protein | | |
| C4B63_76g66 | conserved hypothetical protein | | |
| C4B63_89g19 | retrotransposon hot spot (RHS) protein | | |
| C4B63_8g495 | Regulator of chromosome condensation (RCC1) repeat | | |
| C4B63_92g110 | protein associated with differentiation 8 | | |
| C4B63_96g38 | trans-sialidase, Group V | | |
| C4B63_97g13 | conserved hypothetical protein | | |
| C4B63_97g15 | Zinc finger CCCH domain-containing protein 35 | | |
| C4B63_99g112 | alanine racemase | | |
| C4B63_9g234 | dolichyl-P-Man:GDP-Man7GlcNAc2-PP-dolichyl alpha-1,6-mannosyltransferase | | |
| C4B63_9g376 | Present in the outer mitochondrial membrane proteome 22 | | |
| C4B63_9g377 | Present in the outer mitochondrial membrane proteome 22 | | |
| C4B63_9g383 | conserved hypothetical protein | | |
| C4B63_9g387 | conserved hypothetical protein | | |

| Hb DEGs (t4 vs t0)∩(t24 vs t0) | | | |
|---|---|---|---|
| Downregulated | | Upregulated | |
| ID | Annotation | ID | Annotation |
| C4B63_113g47 | Generative cell specific 1 protein | C4B63_10g436 | SPFH domain / Band 7 family |
| C4B63_12g24 | frataxin-like, mitochondrial precursor | C4B63_111g29 | conserved hypothetical protein |
| C4B63_133g32 | conserved hypothetical protein | C4B63_114g34 | conserved hypothetical protein |
| C4B63_134g49 | trans-sialidase, Group V | C4B63_122g16 | oligopeptidase b |
| C4B63_134g54 | trans-sialidase, Group V | C4B63_134g37 | orotidine-5-phosphate decarboxylase/orotate phosphoribosyltransferase |
| C4B63_13g146 | conserved hypothetical protein | C4B63_13g152 | elongation factor 2 |
| C4B63_13g317 | vesicle-associated membrane protein | C4B63_13g153 | elongation factor 2 |
| C4B63_16g104 | conserved hypothetical protein | C4B63_13g154 | elongation factor 2 |
| C4B63_19g130 | Dual specificity phosphatase, catalytic domain containing protein | C4B63_13g155 | elongation factor 2 |
| C4B63_1g480 | trans-sialidase, Group V | C4B63_13g156 | elongation factor 2 |
| C4B63_21g11 | arginine permease | C4B63_13g157 | elongation factor 2 |
| C4B63_21g27 | arginine permease(fragment) | C4B63_142g6 | GMP synthase |
| C4B63_244g7 | trans-sialidase, Group I | C4B63_15g12 | conserved hypothetical protein |
| C4B63_24g313 | protein kinase | C4B63_15g15 | oligosaccharyl transferase subunit |
| C4B63_2g309 | NIMA-related kinase | C4B63_15g16 | oligosaccharyl transferase subunit |
| C4B63_30g222 | conserved hypothetical protein | C4B63_16g293 | inositol 1,4,5-trisphosphate receptor |
| C4B63_3g1089 | Alpha-ketoglutarate-dependent dioxygenase AlkB-like superfamily | C4B63_188g156c | histone H4 |
| C4B63_3g1092 | alanine racemase | C4B63_192g26 | conserved hypothetical protein |
| C4B63_41g249 | conserved hypothetical protein | C4B63_19g101 | U-rich RNA-binding protein UBP-2 |
| C4B63_442g10 | Uncharacterized protein C4orf22 homolog | C4B63_1g57 | AAA ATPase |
| C4B63_47g60 | conserved hypothetical protein | C4B63_200g32 | 3,2-trans-enoyl-CoA isomerase, mitochondrial precursor |
| C4B63_49g168 | conserved hypothetical protein | C4B63_23g116 | Importin beta-like SAD2 |
| C4B63_4g180 | serine/threonine protein kinase | C4B63_25g234 | RNA cytidine acetyltransferase |
| C4B63_70g65 | conserved hypothetical protein | C4B63_25g237 | eukaryotic translation initiation factor 3 subunit b |
| C4B63_77g33 | conserved hypothetical protein | C4B63_25g240 | Macro domain containing protein |
| C4B63_9g387 | conserved hypothetical protein | C4B63_2g209 | Present in the outer mitochondrial membrane proteome 6 |
| | | C4B63_35g168 | 3-Beta-hydroxysteroid-delta(8), delta(7)-isomerase |
| | | C4B63_396g14 | Present in the outer mitochondrial membrane proteome 6 |
| | | C4B63_40g87 | WD40 repeat-containing protein |
| | | C4B63_41g78 | conserved hypothetical protein |
| | | C4B63_46g16 | conserved hypothetical protein |
| | | C4B63_46g17 | conserved hypothetical protein |
| | | C4B63_47g73 | importin beta-1 subunit |
| | | C4B63_55g67 | DNA-directed rna polymerase I largest subunit |
| | | C4B63_57g65 | membrane-bound acid phosphatase 2 |
| | | C4B63_61g149 | eukaryotic translation initiation factor 3 subunit a |
| | | C4B63_61g92 | conserved hypothetical protein |
| | | C4B63_61g93 | conserved hypothetical protein |
| | | C4B63_61g94 | conserved hypothetical protein |
| | | C4B63_61g95 | conserved hypothetical protein |
| | | C4B63_6g236 | cation transporter |
| | | C4B63_70g93 | DNA repair and recombination helicase protein PIF2 |
| | | C4B63_73g16 | conserved hypothetical protein |
| | | C4B63_7g209 | Ribonuclease P protein subunit p25-like protein |
| | | C4B63_80g17 | ATP binding protein-like protein |
| | | C4B63_81g18 | chaperonin HSP60, mitochondrial precursor(fragment) |
| | | C4B63_85g83 | Eukaryotic initiation factor 4A-1 |
| | | C4B63_97g34 | P-type H+-ATPase |
| | | C4B63_97g36 | P-type H+-ATPase |
| | | C4B63_98g4 | glucose-6-phosphate 1-dehydrogenase |

### Table S6

| Hemin vs. No heme (t4) | | | |
|---|---|---|---|
| ID | Annotaton | log2FC | padj |
| C4B63_18g39 | CRAL/TRIO, N-terminal domain/CRAL/TRIO dom | -1,698147969 | 2,66001E-10 |
| C4B63_27g1020c | RNA-binding protein | -0,939278435 | 8,07905E-07 |
| C4B63_27g184 | integral membrane transport protein | -0,672629407 | 0,016102027 |
| C4B63_51g80 | ferric reductase transmembrane protein | -0,450166135 | 0,001306312 |
| C4B63_51g81 | ferric reductase transmembrane protein | -0,519573589 | 1,24874E-06 |
| C4B63_51g82 | ferric reductase transmembrane protein | -0,461908344 | 9,80739E-06 |
| C4B63_51g83 | ferric reductase transmembrane protein | -0,48197602 | 2,01094E-05 |
| C4B63_51g85 | ferric reductase transmembrane protein | -0,530149195 | 3,29905E-06 |

| Hemin vs. No heme (t24) | | | |
|---|---|---|---|
| ID | Annotaton | log2FC | padj |
| C4B63_18g39 | CRAL/TRIO, N-terminal domain/CRAL/TRIO domain co | -1,781020136 | 2,83772E-29 |
| C4B63_10g454 | ABC transporter(fragment) | -1,147001515 | 0,023416274 |
| C4B63_27g1020c | RNA-binding protein | -1,053472531 | 1,14104E-05 |
| C4B63_10g455 | ABC transporter | -1,02948615 | 0,001536276 |
| C4B63_462g9 | glutamamyl carboxypeptidase(fragment) | -0,910806702 | 0,023416274 |
| C4B63_51g82 | ferric reductase transmembrane protein | -0,822584392 | 1,46408E-15 |
| C4B63_51g81 | ferric reductase transmembrane protein | -0,799944792 | 5,08189E-14 |
| C4B63_24g988c | conserved hypothetical protein | -0,798755109 | 0,003092612 |
| C4B63_51g83 | ferric reductase transmembrane protein | -0,786642986 | 1,03938E-10 |
| C4B63_46g80 | conserved hypothetical protein | -0,750841729 | 0,025368929 |
| C4B63_51g85 | ferric reductase transmembrane protein | -0,742654566 | 1,87681E-08 |
| C4B63_51g80 | ferric reductase transmembrane protein | -0,731765312 | 4,71039E-09 |
| C4B63_69g318c | conserved hypothetical protein | -0,729726224 | 0,001060547 |
| C4B63_32g209 | *Tc* HRG | -0,729531427 | 1,87681E-08 |
| C4B63_27g184 | integral membrane transport protein | -0,623790412 | 0,000179134 |
| C4B63_23g43 | conserved hypothetical protein | -0,521299198 | 0,001035815 |
| C4B63_22g738c | calpain-like cysteine peptidase | -0,457503214 | 0,03424526 |
| C4B63_42g103 | pyridoxal kinase | 0,434936738 | 0,023416274 |
| C4B63_44g117 | conserved hypothetical protein | 0,443705889 | 0,001690098 |
| C4B63_7g135 | Hydroxymethylglutaryl-CoA lyase, mitochondrial | 0,461459445 | 0,039991998 |
| C4B63_70g116 | amino acid permease/transporter | 0,508197543 | 0,001690098 |
| C4B63_47g72 | enolase | 0,511176423 | 7,37702E-06 |

| Hb vs. No heme (t4) | | | |
|---|---|---|---|
| ID | Annotaton | log2FC | padj |
| C4B63_18g39 | CRAL/TRIO, N-terminal domain/CRAL/TRIO dom | -1,43222839 | 7,81732E-07 |
| C4B63_27g1020c | RNA-binding protein | -1,005231966 | 1,71295E-07 |
| C4B63_32g209 | *Tc* HRG | -0,676872385 | 0,001798043 |
| C4B63_51g81 | ferric reductase transmembrane protein | -0,404398408 | 0,002429377 |
| C4B63_51g83 | ferric reductase transmembrane protein | -0,441116788 | 0,000799977 |
| C4B63_51g85 | ferric reductase transmembrane protein | -0,442097131 | 0,001384228 |

| Hb vs. No heme (t24) | | | |
|---|---|---|---|
| ID | Annotaton | log2FC | padj |
| C4B63_18g39 | CRAL/TRIO, N-terminal domain/CRAL/TRIO domain co | -1,009685067 | 1,41308E-07 |
| C4B63_44g117 | conserved hypothetical protein | 0,426749137 | 0,034246414 |
| C4B63_51g81 | ferric reductase transmembrane protein | -0,41343673 | 0,048118011 |
| C4B63_51g82 | ferric reductase transmembrane protein | -0,426087127 | 0,031101978 |

Figure S2

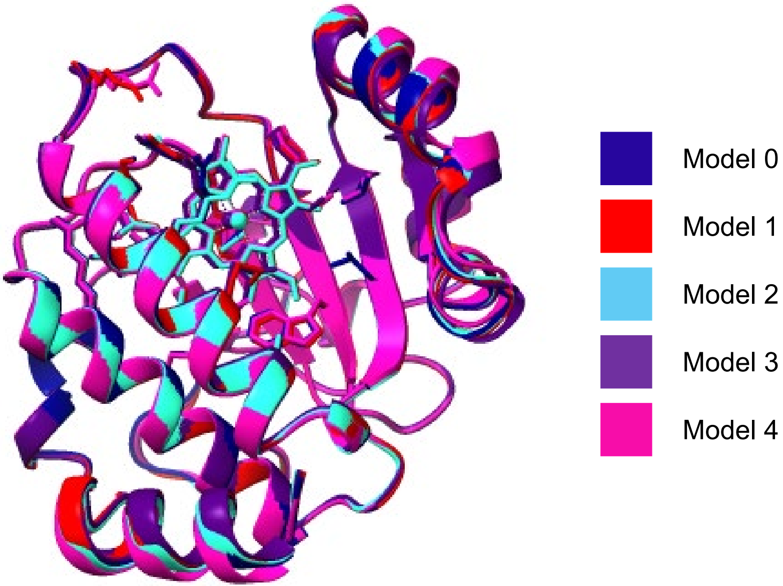

### Table S7

**Model 0**

| Name | Value |
|---|---|
| chain_iptm | [List] |
| chain_pair_iptm | [List] |
| chain_pair_pae_min | [List] |
| chain_ptm | [List] |
| fraction_disordered | 0 |
| has_clash | 0 |
| iptm | 0,85 |
| num_recycles | 10 |
| ptm | 0,89 |
| ranking_score | 0,86 |

**Model 1**

| Name | Value |
|---|---|
| chain_iptm | [List] |
| chain_pair_iptm | [List] |
| chain_pair_pae_min | [List] |
| chain_ptm | [List] |
| fraction_disordered | 0 |
| has_clash | 0 |
| iptm | 0,85 |
| num_recycles | 10 |
| ptm | 0,89 |
| ranking_score | 0,86 |

**Model 2**

| Name | Value |
|---|---|
| chain_iptm | [List] |
| chain_pair_iptm | [List] |
| chain_pair_pae_min | [List] |
| chain_ptm | [List] |
| fraction_disordered | 0 |
| has_clash | 0 |
| iptm | 0,83 |
| num_recycles | 10 |
| ptm | 0,89 |
| ranking_score | 0,84 |

**Model 3**

| Name | Value |
|---|---|
| chain_iptm | [List] |
| chain_pair_iptm | [List] |
| chain_pair_pae_min | [List] |

| | |
|---|---|
| chain_ptm | [List] |
| fraction_disordered | 0 |
| has_clash | 0 |
| iptm | 0,79 |
| num_recycles | 10 |
| ptm | 0,88 |
| ranking_score | 0,8 |

**Model 4**

| Name | Value |
|---|---|
| chain_iptm | [List] |
| chain_pair_iptm | [List] |
| chain_pair_pae_min | [List] |
| chain_ptm | [List] |
| fraction_disordered | 0 |
| has_clash | 0 |
| iptm | 0,78 |
| num_recycles | 10 |
| ptm | 0,88 |
| ranking_score | 0,8 |
